# Life history and deleterious mutation rate coevolution

**DOI:** 10.1101/2022.05.11.491530

**Authors:** Piret Avila, Laurent Lehmann

**Author notes:** **Author contributions:** All authors gave final approval for publication and each agreed to be held accountable for all the parts of the work performed therein. **Data Accessibility Statement:** The proofs of the analysis written in Mathematica and the code for individual-based simulations are available in the S.I. as a Mathematica notebook.

## Abstract

The cost of germline maintenance gives rise to a trade-off between lowering the deleterious mutation rate and investing in life history functions. Therefore, life history and the mutation rate coevolve, but this coevolution is not well understood. We develop a mathematical model to analyse the evolution of resource allocation traits, which simultaneously affect life history and the deleterious mutation rate. First, we show that the invasion fitness of such resource allocation traits can be approximated by the basic reproductive number of the least-loaded class; the expected lifetime production of offspring without deleterious mutations born to individuals without deleterious mutations. Second, we apply the model to investigate (i) the coevolution of reproductive effort and germline maintenance and (ii) the coevolution of age-at-maturity and germline maintenance. This analysis provides two resource allocation predictions when exposure to environmental mutagens is higher. First, selection favours higher allocation to germline maintenance, even if it comes at the expense of life history functions, and leads to a shift in allocation towards reproduction rather than survival. Second, life histories tend to be faster, characterized by individuals with shorter lifespans and smaller body sizes at maturity. Our results suggest that mutation accumulation via the cost of germline maintenance can be a major force shaping life-history traits.

## 1 Introduction

Maintaining and accurately transmitting genetically encoded information is central for every living organism. Mutations induce errors in the processing of genetic information and their effects on survival and reproduction tend to be deleterious (e.g. Eyre-Walker and Keightley, 2007; Halligan and Keightley, 2009; Charmouh et al., 2023). Therefore, it is likely that selection primarily favours a reduction of the mutation rate of organisms (Sniegowski et al., 2000). Yet, investing resources into germline maintenance is physiologically costly (Kirkwood, 1986; Maklakov and Immler, 2016; Monaghan and Metcalfe, 2019; Chen et al., 2020). Thus, the balance between selection against deleterious mutations driving for lower mutation rates and selection for reduced physiological cost increasing mutation rates is expected to lead to a positive rate of mutation. This argument has been formalised in a number of classical population genetic models assuming semelparous reproduction (e.g. Kimura, 1967; Kondrashov, 1995; Dawson, 1998, 1999; Johnson, 1999*b*; André and Godelle, 2006; Gervais and Roze, 2017) and also under iteroparous reproduction (Lesaffre, 2021) to show that evolution indeed favours an evolutionary stable non-zero mutation rate. These studies emphasise the role of the physiological cost of germline fidelity in explaining the patterns of mutation rates but do not connect the cost of germline fidelity explicitly to life-history evolution, which depends on underlying physiological trade-offs.

Indeed, a central premise made in life-history theory is that life-history trade-offs are mediated through the allocation of resources to different life-history functions, such as growth, reproduction, and maintenance of soma, or information gathering and processing (Stearns, 1992; Roff, 2008). Since germline maintenance takes a toll on available resources, mutation rate evolution and life-history evolution are tied together through a resource allocation trade-off. This implies that the rate of deleterious mutations should evolve jointly with life history and affect various life-history traits, such as reproductive effort, age-at-maturity, adult body size, and expected lifespan. Yet the bulk of models about the evolution of mutation rates, which often go under the heading of modifier models, consider physiologically neutral mutation rates (e.g. Leigh, 1970; Gillespie, 1981; Holsinger and Feldman, 1983; Liberman and Feldman, 1986; Gerrish et al., 2007; Altenberg et al., 2017; Baumdicker et al., 2020). And while the effect of fixed mutation rate on life-history trade-offs has been studied before (e.g. Charlesworth, 1990; Dańko et al., 2012), these models suggest that a relatively high level of mutation rates is needed for mutation accumulation to alter life-history trade-offs. This led to the conclusion that mutation accumulation is a minor force in shaping life-history traits (Dańko et al., 2012). But these studies view mutation rates as fixed traits acting only as a hindrance to adaptive life-history evolution. Hence, no study has yet investigated the coevolution of both life-history traits and deleterious mutation rates via allocation to germline maintenance.

The aim of this paper is to start to fill this gap by formally extending evolutionary invasion analysis– “ESS” theory–(e.g., Maynard Smith, 1982; Eshel and Feldman, 1984; Metz et al., 1992; Charlesworth, 1994), which has been routinely applied to life-history evolution (e.g., León, 1976; Michod, 1979; Schaffer, 1982; Iwasa and Roughgarden, 1984; Stearns, 1992; Perrin, 1992; Perrin and Sibly, 1993; Cichon and Kozlowski, 2000; Iwasa, 2000; Day and Taylor, 2000), to the case where life-history trait(s) evolving by selection also control the rate of deleterious mutations. This covers the situation where life-history resource allocation schedules evolve on a background where deleterious mutation accumulation can occur. Our formalisation thus aims to integrate both the standard theories of life-history evolution and deleterious mutation rate accumulation.

The rest of this paper is organised into two parts. First, we characterise the invasion process of a mutant life-history trait affecting the load of deleterious mutation accumulation in an age-structured population. We show that ascertaining the joint evolutionary stability of life-history schedules and mutation rates of deleterious mutations is usually complicated, but under certain biologically feasible conditions; in particular, when the zero mutation class (least-loaded class) dominates the population in frequency, evolutionary stability can be ascertained from the basic reproductive number of the least-loaded class alone (expected lifetime production of offspring with no mutations born to individuals with no mutations). Second, we analyse and solve two concrete biological scenarios: (i) joint evolution between reproductive effort and germline maintenance, where individuals face a trade-off between allocating resources to survival, reproduction and germline maintenance and (ii) joint evolution between age-at-maturity and germline maintenance, where individuals face a trade-off between allocating resources to growth, reproduction and germline maintenance. These scenarios allow us to illustrate how our model can be applied to analyse questions in life history and mutation rate evolution and provide predictions about how life history and mutation rate coevolves. It also allows us to verify that the analysis based on the basic reproductive number of the least-loaded class as an invasion fitness proxy is a useful approximation, as analytical predictions match closely with the results from individual-based stochastic simulations.

## 2 Model

### 2.1 Main biological assumptions

We consider a panmictic population of haploid individuals reproducing asexually. The population size is assumed to be regulated by density-dependent competition. Individuals in the population are structured into age classes and each individual undergoes birth, possibly development, reproduction, and death. We allow for discrete and continuous age classes with corresponding discrete and continuous demographic processes. In the discrete case, an individual in the age range [*a*−1, *a*] for *a* = 1, 2, 3, …, will, by convention, belong to the *a*-th age class and so we begin counting discrete age classes with 1 (one of the two possible conventions to count discrete age classes, Fig. 3.1 Case, 2000). An individual of age class *a* is characterised by a type *θ*(*a*) = (**u**(*a*), *n*_m_(*a*)), which consists of two genetically determined components (see Table 1 for a list of symbols and more formalities on functions). The first component, **u**(*a*), is the individual’s lifehistory trait expressed in age class *a*; namely, a resource allocation to different life-history functions (e.g. growth, reproduction, or somatic maintenance, see e.g. Iwasa and Roughgarden, 1984; Kozlowski, 1992; Perrin, 1992; Perrin and Sibly, 1993; Day and Taylor, 2000). We denote by **u** = {**u**(*a*)}_*a*∈𝒯F_ the whole life-history schedule or path of resource allocation over all possible age classes 𝒯F an individual can be in (formally, **u** ∈ 𝒰[𝒯], where 𝒰[𝒯] is the set of all admissible life-history schedules with domain 𝒯, which for discrete age classes is 𝒯 = {1, 2, .., *T* } and for continuous age classes is 𝒯 = [0, *T*] when the maximum lifespan is *T*). The second component, *n*_m_(*a*), represents the number of deleterious germline mutations of an individual belonging to the *a*-th age class, which, by definition, negatively affects the viability traits (e.g. physiology, reproduction). Hence, *n*_m_(*a*) can be thought of as the load of deleterious mutations as considered in classical population genetic models of mutation accumulation (e.g., Kimura and Maruyama, 1966; Haigh, 1978; Dawson, 1999; Bürger, 2000), but here extended to an age-structured model (see also Steinsaltz et al., 2005 for aged-structured mutation accumulation model). As such, *n*_m_ = {*n*_m_(*a*)}_*a*∈𝒯_ represents the number of deleterious mutations an individual has acquired across its lifespan.

**Table 1:** List of key symbols of the general model.

| Key symbols of the model |  |
| --- | --- |
| $a$ | Individual age; age $a$ can take either discrete $a \in \mathcal{T} = \{0, 1, 2, \dots\}$ or continuous $a \in \mathcal{T} = [0, \infty)$ values over all possible age classes $\mathcal{T}$ under the scenario under consideration. |
| $\mathbf{u}(a)$ | Individual life-history trait expressed at age $a$ (e.g. proportional allocation fecundity, survival, germline maintenance); formally, $\mathbf{u} : \mathcal{T} \rightarrow \mathbb{R}^n$ |
| $\mathbf{u} = \{\mathbf{u}(a)\}_{a \in \mathcal{T}}$ | Full life-history schedule over all age classes (e.g. proportional allocation of resources to fecundity from birth to death); formally, $\mathbf{u} \in \mathcal{U}[\mathcal{T}]$, where $\mathcal{U}[\mathcal{T}]$ is a set of all admissible life-history schedules; namely, a set of discrete or continuous real-valued functions over domain $\mathcal{T}$. |
| $n_m(a)$ | Number of deleterious mutations at age $a$ in the locus where deleterious mutations can accumulate; formally, $n_m : \mathcal{T} \rightarrow \mathbb{N}$. Since we assume asexual reproduction, the genetic details of the locus for trait $n_m(a)$ is irrelevant (i.e. it may consist of many underlying loci). |
| $n_m = \{n_m(a)\}_{a \in \mathcal{T}}$ | Profile of deleterious mutations across all age classes; formally, $n_m \in \mathbb{N}[\mathcal{T}]$ is an element of the space $\mathbb{N}[\mathcal{T}]$ of all possible discrete functions of range $\mathbb{N}$ over domain $\mathcal{T}$. |
| $\mathbf{p}(\mathbf{v})$ | Equilibrium probability distribution for the number of deleterious mutations in the resident population carried by individuals across the different age-classes, formally $\mathbf{p}(\mathbf{v}) \in \Delta(\mathbb{N} \times \mathcal{T})$, where $\Delta(A)$ is the set of probability measure over set $A$. In the absence of age-dependence, $\mathbf{p}(\mathbf{v}) \in \Delta(\mathbb{N})$, see eq. (B-5) for an example. |
| $\rho_0(\mathbf{u}, \mathbf{v})$ | Invasion fitness (per-capita growth ratio) of a zero-class mutant allele $\mathbf{u}$ (i.e. carriers of this mutant have zero deleterious mutations) of $a$ in a population monomorphic for resident trait $\mathbf{v}$. |
| $\tilde{R}_0(\mathbf{u}, \mathbf{v})$ | Basic reproductive number of the least-loaded class, i.e. the expected number of offspring with zero deleterious mutations produced by an individual with zero deleterious mutations |
| $\tilde{b}_0(a, \mathbf{u}, \mathbf{v})$ | Effective number of newborns with zero mutations produced by zero-class mutant individuals age $a$ in a resident population (discrete time model); effective birth rate of newborns with zero mutations of zero-class mutant individual of age $a$ in a resident population (continuous time model). |
| $d_0(a, \mathbf{u}, \mathbf{v})$ | Death rate of zero-class mutant individual at age $a$ in the resident population |
| $\tilde{l}_0(a, \mathbf{u}, \mathbf{v})$ | Probability of survival of a mutant zero-class individual to age $a$ in a resident population. |
| $\mu_f(a, \mathbf{u}, \mathbf{v})$ | Rate at which germline mutations appear in an offspring when giving birth at age $a$. |
| $\mu_s(a, \mathbf{u}, \mathbf{v})$ | Rate at which germline mutations appear in an individual at age $a$. |

We envision that the genotype determining the type *θ* = {*θ*(*a*)}_*a*∈𝒯_ = (**u**, *n*_m_) of an individual consists of two separate positions (or loci) that are necessarily linked under asexual reproduction, one locus determining **u** and the other *n*_m_ (see Fig. 1). The mutation rate at the life-history trait **u** locus is assumed to be fixed and is thus exogenously given. However, this trait is assumed to control allocation to germline maintenance and other life-history functions, and thus to control the mutation rate at the locus where the *n*_m_ deleterious mutations accumulate whose number are thus endogenously determined (Fig. 1). This separation between genes coding for the life-history trait **u** and those coding for mutation accumulation *n*_m_ (see Fig. 1) is conceptually equivalent to the separation between *modifier locus* and loci affecting vital rates of modifier theory, where the modifier locus affects the pattern of transmission of other traits (e.g., Leigh, 1970; Altenberg, 2009). Since the life-history trait **u** is the evolving trait controlling trait *n*_m_, it can be regarded as the modifier trait.

**Figure 1:**
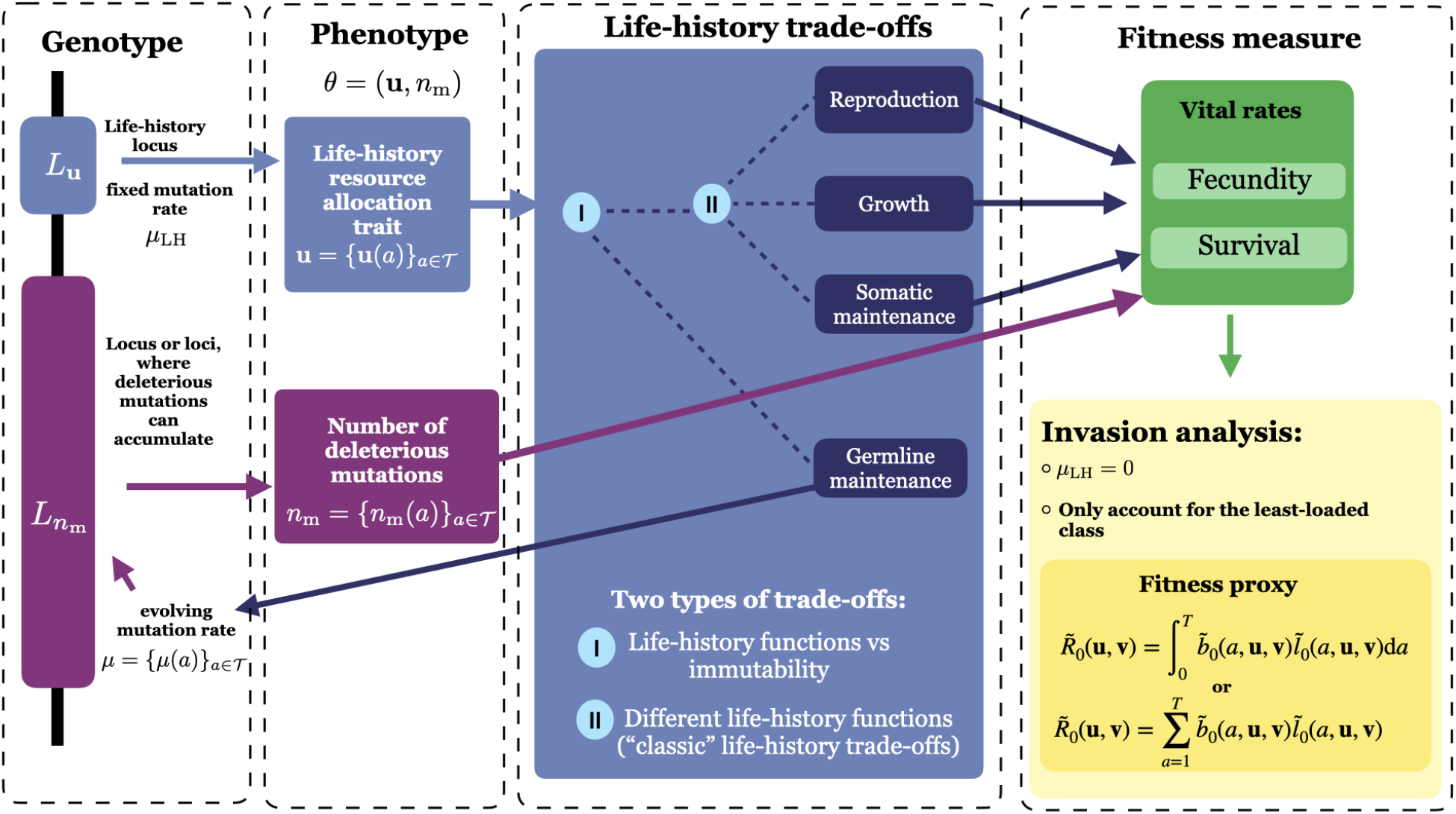
Key components of the life-history model with mutation accumulation. An individual’s genotype is characterised by a life-history locus *L*_**u**_ and a deleterious mutation locus (purple rectangle) 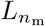 . The mutation rate at the life-history locus *μ*_LH_ is considered to be fixed, while the mutation rate *μ* at the deleterious mutation locus depends on the life-history trait **u** and is evolving. Individuals are characterised by the life-history allocation schedule **u** = {**u**(*a*)}_*a*∈𝒯_ (life-history trait) and the number *n*_m_ ={ *n*_m_(*a*) }_*a*∈𝒯_ of deleterious mutations accumulated in the germline throughout lifespan, where *a* denotes the age of an individual. The resource allocation trait leads to two different types of tradeoffs: (i) between immutability vs life-history and (ii) between different life-history functions themselves (“classic” life history trade-offs, e.g. Stearns, 1992; Roff, 2008). Hence, the life-history locus affects the vital rates and thus fitness directly via resource allocation to life-history functions and indirectly through allocation to germline maintenance since vital rates depend on the number of deleterious mutations. For the invasion analysis, we use the basic reproductive number of the least-loaded class 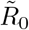 as a fitness proxy (eqs. 1 and 4 as detailed in section 2.3.2).

Let us now first focus on a continuous time process and introduce the birth *b*(*a*), death *d*(*a*), and deleterious mutations *μ*(*a*) rates of an individual of age *a*. We assume that the birth *b*(*a*) and death *d*(*a*) rates can depend on both the number of deleterious mutations *n*_m_(*a*) at age *a* and the life-history schedule **u**, while the mutation rate *μ*(*a*) depends only the life-history schedule **u** (see also Fig. 1). The vital rates may further depend on properties of the population, such as age class densities and allele frequencies, but we leave this dependence implicit. We note that when measured on an exponential scale, the death and mutation rates define a survival probability exp(−*d*(*a*)) and an immutability probability exp(−*μ*(*a*)) (probability that no mutations occur in an individual of age *a*), which allows us to handle the discrete-time process with the same basic quantities. Further, for a discrete-time process, the birth function *b*(*a*) gives the expected number of offspring produced by an individual of age *a*, while for a continuous time process, *b*(*a*) is defined as the rate at which an individual produces a single offspring. Finally, we make the following assumptions that are central to our analysis.

1. Mutations at the locus determining *n*_m_ can only be deleterious, with no specific assumption on the effect size that can range thus from neutral to lethal. The effective birth rate *b*_*i*_(*a*), survival probability exp(−*d*_*i*_(*a*)), and immutability probability exp(−*μ*_*i*_(*a*)) of an individual at age *a* with *n*_m_(*a*) = *i* mutations are non-increasing functions of the number of mutations. Formally, *b*_*i*_(*a*) ≥*b*_*i*+1_(*a*), *d*_*i*_(*a*) ≤ *d*_*i*+1_(*a*), and *μ*_*i*_(*a*) ≤ *μ*_*i*+1_(*a*).
2. Deleterious mutations can only accumulate within an individual. There are no back mutations and an individual with *i* deleterious mutations can only mutate towards having *i* + 1 such mutations.

These assumptions are standard in population genetics (e.g. Kimura, 1967; Leigh, 1970; Haigh, 1978; Dawson, 1998; Johnson, 1999*a*; Gillespie, 2004) and we are here endorsing these assumptions in formalising selection at the life-history locus. Namely, the objective of our analysis is to develop a tractable evolutionary invasion analysis to evaluate candidate evolutionary stable life-history trait *u*^∗^ ∈ 𝒰[𝒯] that will be favoured by long-term evolution.

### 2.2 Invasion analysis

For a moment, let us ignore the effect of deleterious mutations. Then, evolutionary invasion analysis (e.g., Eshel and Feldman, 1984; Parker and Maynard Smith, 1990; Metz et al., 1992; Charlesworth, 1994; Ferrière and Gatto, 1995; Eshel, 1996; Otto and Day, 2007; Lehmann et al., 2016; Avila and Mullon, 2023; Van Cleve, 2023) can be applied straightforwardly to our model as follows. First, the population is postulated to be large enough and the mutations at the life-history trait rare enough so that one can focus on the invasion fitness *ρ*(**u, v**) of a mutant trait **u** introduced as a single copy in a population monomorphic for some resident life-history trait **v** (i.e., the geometric growth ratio of the mutant) in order to characterize evolutionary stable trait values. The invasion fitness can be interpreted as the per capita number of mutant copies produced per unit of time by the mutant lineage descending from the single initial mutation when the mutant reproductive process has reached a stationary state distribution (over ages) in a resident population at its demographic attractor while the mutant remains overall rare in the population. To invade the population, the invasion fitness must exceed unity *ρ*(**u, v**) *>* 1; otherwise, it follows from the theory of branching processes that the mutant will go extinct with probability one in an (infinitely) large population (Harris, 1963; Mode, 1971 and see also Appendix A). A trait value **u**^∗^ will be called uninvadable if it is resistant to invasion by all trait values in the trait space: *ρ*(**u, u**^∗^) ≤ 1 for all **u** ∈ 𝒰[𝒯]. Hence, an uninvadable trait **u**^∗^ maximises the invasion fitness of the mutant holding the resident at the uninvadable population state; namely, **u**^∗^ solves the maximization problem: max_**u**∈𝒰[𝒯]_ *ρ*(**u, u**^∗^), whereby **u**^∗^ defines a candidate endpoint of evolution.

To be a meaningful endpoint of evolution, an uninvadable trait needs to be an attractor of the evolutionary dynamics and thus be convergence stable (Eshel and Motro, 1988; Geritz et al., 1998; Leimar, 2009*a*). When mutant and resident traits are sufficiently close to each other so that selection is weak, mutants increasing invasion fitness will not only invade the resident population but also become fixed in it (Eshel et al., 1997, a result applying to the present and more complicated demographic scenarios, Rousset, 2004; Priklopil and Lehmann, 2021). Therefore, invasion fitness alone under weak phenotypic deviations allows to determine whether gradual evolution under the constant influx of mutations will drive a population towards the uninvadable population state, regardless of patterns of frequency-dependence. An evolutionary invasion analysis thus generally consists of both using invasion fitness to (i) characterise uninvadable trait values and (ii) determine whether these are attractors of the evolutionary dynamics, thus allowing to make definite statements about joint evolutionary dynamics. Useful summaries of these concepts can be found in Geritz et al. (1998); Leimar (2009*b*) and individual-based stochastic simulations have repeatedly validated the conclusions of this approach in genetic explicit contexts (e.g., Mullon et al., 2018; Mullon and Lehmann, 2019, see also Otto and Day, 2007; Dercole and Rinaldi, 2008 for textbook discussions). This approach to characterise stable and attainable life histories has indeed been in use more or less explicitly and completely in standard life-history theory for decades (e.g., León, 1976; Michod, 1979; Schaffer, 1982; Iwasa and Roughgarden, 1984; Perrin, 1992; Perrin and Sibly, 1993; Kawecki, 1993; Charlesworth, 1994; McNamara, 1997; Charnov, 1997; Cichon and Kozlowski, 2000; Iwasa, 2000; Irie and Iwasa, 2005; Rueffler et al., 2013; Avila et al., 2019). We next push forward this approach into the realm where life-history evolution interacts with mutation accumulation and thus relax the standard life-history theory assumption that the rate of deleterious mutations is zero.

### 2.3 Invasion analysis with mutation accumulation

#### 2.3.1 Mutation-selection balance in the resident population

Let us now consider that deleterious mutations can accumulate. We still assume that mutations at the life-history locus are rare enough so that whenever a mutant trait **u** arises, it does so in a resident population monomorphic for some resident life-history trait **v**. But owing to the occurrence of deleterious mutations, the resident population will be polymorphic for the number of deleterious mutations in the *n*_m_ locus, and this polymorphism will depend on **v**. The resident population is then assumed to have reached a mutation-selection equilibrium for deleterious mutations, and the resident trait **v** thus determines a resident probability distribution **p**(**v**) over the different number of deleterious mutations carried by individuals across the different age-classes. This assumption is nothing else than the usual assumption of the internal stability of the resident population used in invasion analysis (see e.g. Altenberg et al., 2017; Eshel and Feldman, 1984; Metz et al., 1992). Here, it entails that the resident population has reached an equilibrium for both demographic and genetic processes.

In the absence of age classes, **p**(**v**) is the equilibrium probability distribution for the number of deleterious mutations in standard selection-mutation balance models (see Bürger, 2000 for a general treatment). For instance, when the number of novel (deleterious) mutations follows a Poisson distribution with the mean rate *μ* and each additional mutation decreases baseline fecundity by a constant multiplicative factor *σ* in a semelparous population, then **p**(**v**) is Poisson distributed with mean *μ/σ* (Haigh, 1978, Bürger, 2000, p. 300, and see also eq. B-5 of the Appendix). This holds in an age-structured population across age classes under a certain but limited number of conditions (Steinsaltz et al., 2005). More generally, **p**(**v**) will depend on the details of the model.

The key feature determining the invasion process of a mutant **u** in a resident population **v** is the mutational background on which it arises. This means that the invasion fitness depends on how many deleterious mutations the mutant individual carries (i.e. mutational class of an individual) and thus on the distribution **p**(**v**) in the resident population. This invasion process is derived from stochastic process considerations in Appendix A and shown to be a reducible process whereby an invasion fitness can be associated to each mutational class (see Appendix A.1 for details). However, it follows from our assumptions about deleterious mutations (section 2.1) that the invasion fitness for a mutant life history trait arising on any mutational background cannot be higher than the invasion fitness for this trait that arises in an individual belonging to the least-loaded class (see Appendix A.1 for details). Thus, as a first step, it is reasonable to consider a situation where the mutation-selection process is such that the least-loaded class–individuals with zero mutations–dominates the population in frequency (i.e. the frequency of the zero-class individuals is close to one) so that this class defines invasion fitness.

Indeed, if selection is stronger than mutation, then deleterious alleles will tend to be purged and the mutation-selection balance will be far away from the error threshold of mutation accumulation or meltdown of asexual populations (e.g., Eigen, 1971; Lynch et al., 1993; Szathmary and Maynard Smith, 1997). For instance, in the aforementioned classical mutation-selection equilibrium model (Haigh, 1978, Bürger, 2000) and its generalization to overlapping generations (see eq. B-5), the frequency of the zero mutation class is *e*^−*μ/σ*^. So when *μ* ≪ *σ*, say for definiteness, the selection coefficient is one order of magnitude larger than the mutation rate (e.g. for *μ* = 0.01 and *σ* = 0.1, *μ/σ* = 0.1), then the least-loaded class dominates in frequency (*e*^−*μ/σ*^ ≈ 0.9). Under these conditions, the click rate of Muller’s ratchet, which is the rate at which individuals with the least amount of deleterious mutations in the population become extinct in a finite population (i.e. in the presence of genetic drift), is small for finite but sufficiently large population sizes. For instance, in a population of size *N* = 1000 under a Moran process of overlapping generations, the click rate is 8.4 × 10^−34^ (obtained from 1*/τ* where 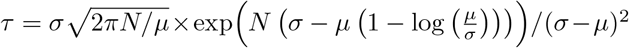 is the inverse of the click rate, see eq. 23 Metzger and Eule, 2013, where *σ* = *s* and *μ* = *u*). Hence, the click rate of Muller’s ratchet can be considered negligible compared to the scale of mutation rates. Genetic drift is thus unlikely to fully eliminate the least-loaded class from the population, even for small populations. Therefore, if the selection coefficient against a deleterious mutation is one order of magnitude larger than the mutation rate in a population resident for life-history trait **v**, then a new mutant life-history trait **u** is likely to surface in a member of the least-loaded class (i.e. zero mutation background).

#### 2.3.2 Invasion analysis for dominating least-loaded class

We now fully endorse the assumption that the least-loaded class dominates in frequency the resident population. It allows us to characterise uninvadability and convergence stability directly from the basic reproductive number of the least-loaded class 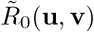, which is the expected number of offspring without mutations produced by an individual without mutations over its lifespan. This is a fitness proxy for invasion fitness whereby an uninvadable strategy **u**^∗^ solves the maximization problem 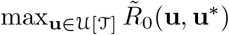 (see Appendix A.1 for a proof). In a discrete age-structured population, the basic reproductive number of the least-loaded class is given explicitly in terms of vital rates as

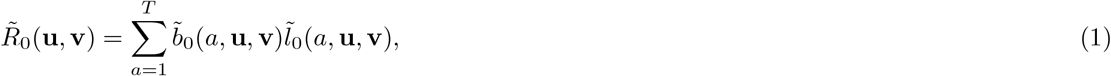

where

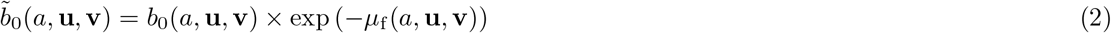

and

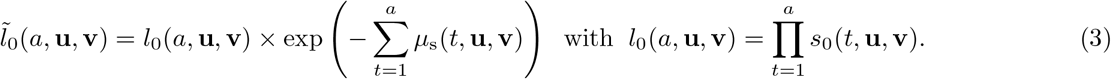

In eq. (2), *b*_0_(*a*, **u, v**) is the effective fecundity of an individual in the *a*-th age class with trait **u** and exp (−*μ*_f_ (*a*, **u, v**)) is the probability that such an individual does not produce a mutated offspring with *μ*_f_ (*a*, **u, v**) being the mutation rate during reproduction in the *a*-th age class (recall that *a* ∈ {1, 2, .., *T* }). In eq. (3), *l*_0_(*a*, **u, v**) is the probability that an individual with trait **u** survives to age *a*, which depends on its probability *s*_0_(*a*, **u, v**) = exp (−*d*_o_(*a*, **u, v**)) of survival over the age interval [*a* − 1, *a*], where *d*_0_(*a*, **u, v**) is its death rate. Finally, exp 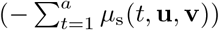 is the probability that an individual with trait **u** does not acquire any germline mutation until age *a* with *μ*_s_(*a*, **u, v**) being the germline mutation rate when being in the *a*-th age class. In eqs. (2)–(3), we have thus distinguished between the germline mutation rate *μ*_f_ (*a*, **u, v**) when an offspring is produced and that rate *μ*_s_(*a*, **u, v**) in the parent in the *a*-th age class. When *μ*_f_ (*a*, **u, v**) = *μ*_s_(*a*, **u, v**) = 0 for all *a* ∈ {1, 2, .., *T* }, eq. (1) reduces to the standard basic reproductive number for age-structured populations (e.g. Charlesworth, 1994). We emphasise that we allowed for fecundity, survival and mutation rate to be dependent on the whole life history schedule because the evolving traits may affect physiological state variables (e.g. body size). As long as there is a direct correspondence between age and physiological state (see e.g. the discussion in de Roos, 1997), then the extension of current formalisation to physiologically-structured populations is direct (see also section 3.2 for an example). Further, since all rates in eqs. (2)–(3) depend on the resident trait **v**, individuals can be affected by the trait of others and our model thus covers frequency- and density-dependent interactions.

For a continuous set of age classes 𝒯 = [0, *T*], the basic reproductive number of the least-loaded class is

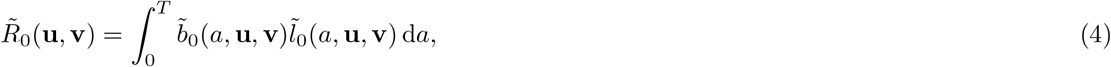

(see Appendix A.1 for a proof) where 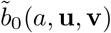 takes the same functional form as in eq. (2) but is now interpreted as the effective birth rate (of offspring with no mutations) at age *a*, and 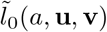 satisfies the differential equation:

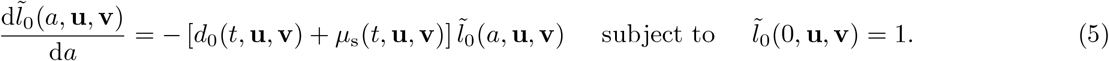

We now make four observations on the use of 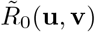 to characterise long-term coevolution for lifehistory traits and mutation rates. (1) Because 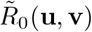 depends on the number of deleterious mutations in the population solely via **v**, the distribution **p**(**v**) is needed only under frequency-dependent selection. This makes life-history evolution in the presence of deleterious mutations tractable even if the underlying evolutionary process of mutation is not (see section eq. 3.2 for an example). The characterisation of uninvadability using 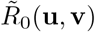 (and thus applying eqs. 1–5) generalises the results of Leigh (1970) and Dawson (1998, p. 148) to overlapping generations and an explicit life-history context. (2) Because 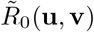 takes the standard form of the basic reproductive number, optimal control and dynamic game theory results can be applied to characterise uninvadability. This is particularly useful for reaction norm and developmental evolution and formalising different modes of trait expressions (e.g., Perrin and Sibly, 1993; Avila et al., 2021). (3) While low mutation rates relative to selection are presumed to be able to use 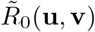 as a proxy for invasion fitness, these mutation rates are endogenously determined by the uninvadable strategy. It is thus plausible that the uninvadable mutation rate generally entails a low mutation rate. So the assumption of a low mutation rate may not appear so drastic, and the extent to which this assumption is limiting depends on investigating explicit evolutionary scenarios. (4) If deleterious mutations act in such a way that the invasion fitness of a life-history mutant appearing in any background is proportional to that of the least-loaded class (see Appendix A.1), then the invasion process is fully characterised by the invasion fitness of the least-loaded class 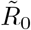. This is the case for the standard mutation accumulation models with multiplicative effects of (deleterious) mutations under semelparous life-history (e.g., Kimura, 1967; Dawson, 1998; Johnson, 1999*b*; Gervais and Roze, 2017). Then using 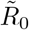 does not rely on making any assumption of low (deleterious) mutation rates relative to selection.

All this gives good reasons to use 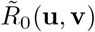 as a proxy of invasion fitness to understand life-history evolution in the context of mutation accumulation and as such, in the rest of this paper we consider two scenarios of the joint evolution of life history and mutation rate which we analyse by using 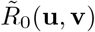. This allows us to illustrate the different concepts, demonstrate the usefulness of focusing on 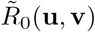 to get insights about how life-history evolution interacts with mutation accumulation, and check how closely our analysis matches with individual-based stochastic simulations.

## 3 Examples of life-history and mutation rate coevolution

### 3.1 Coevolution of reproductive effort and germline maintenance

#### 3.1.1 Biological scenario

Our first scenario considers the evolution of reproductive effort when resources can be allocated to (germline) maintenance in an iteroparous population. To that end, we assume a population with a large but fixed number *N* of individuals undergoing the following discrete-time life-cycle (see Table 2 for a summary of key symbols for this example). (1) Each of the *N* adult individuals produces a large number of juveniles and either survives or dies independently of other individuals. Juveniles and surviving adults acquire mutations at the deleterious allele locus at the same rate. (2) Density-dependent competition occurs among juveniles for the vacated breeding spots (left by the dead adults) and the population is regulated back to size *N*.

**Table 2:** List of key symbols of “Coevolution of reproductive effort and the mutation rate model”.

| Symbols for “Coevolution of reproductive effort and the mutation rate”. |  |
| --- | --- |
| $u_g, v_g$ | Proportional allocation of resources to germline maintenance of mutant and resident individuals, respectively. |
| $u_s, v_s$ | Proportional allocation of resources to the survival of mutant and resident individual, respectively. |
| $N$ | Total population size; exogeneously fixed ($N = 7500$ in simulations). |
| $f_b, s_b$ | Baseline (maximal) fecundity and probability of survival, respectively ($f_b = 5$, $s_b = 0.5$ ). |
| $\mu(\mathbf{u}) = \mu_s(\mathbf{u}) = \mu_f(\mathbf{u})$ | Rate at which germline mutations appear in an offspring when giving birth and when surviving from generation to the next. |
| $\mu_b$ | Baseline mutation rate at which germline mutations appear; mutation rate, when no resources are allocated into germline maintenance. |
| $\alpha_f, \alpha_s$ | Efficiency parameters (or scaling factors) of investing resources into fecundity and survival, respectively; lower values of $\alpha_f, \alpha_s$ corresponds to a higher efficiency of investment; analytical results obtained for $\alpha_f = \alpha_s = \alpha$ ($\alpha = 0.02$, $\alpha = 0.1$, $\alpha = 0.2$ in simulations). |
| $\alpha_\mu$ | Efficiency parameter (or scaling factor) of investing resources into germline maintenance; the higher value of $\alpha_\mu$ corresponds to a higher efficiency of investment; ($\alpha_\mu = 2$). |

We postulate that an individual has a life-history trait consisting of two components **u** = (*u*_g_, *u*_s_) (**u** ∈ 𝒰[𝒯] = [0, 1]^2^), which determines how a fixed amount of resources available to each individual is allocated between three physiological functions: (i) a proportion (1 − *u*_g_)(1 − *u*_s_) of resources is allocated to reproduction, (ii) a proportion (1 − *u*_g_)*u*_s_ of resources is allocated to survival, and (iii) a proportion *u*_g_ of resources is allocated to germline maintenance.

We assume that an individual with trait **u** and *i >* 0 deleterious mutations has the following fecundity *f*_*i*_(**u**), survival probability *s*_*i*_(**u**), and mutation rate *μ*(**u**) [at giving birth and when surviving to the next generation, i.e., *μ*(**u**) = *μ*_f_ (**u**) = *μ*_s_(**u**)]:

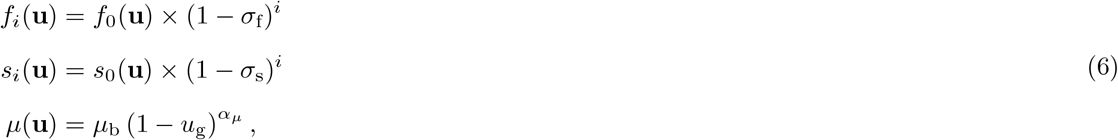

where 0 ≤ *σ*_f_ ≤ 1 and 0 ≤ *σ*_s_ ≤ 1 are, respectively, the reductions in fecundity and survival from carrying an additional deleterious mutation (that are assumed to act multiplicatively), *μ*_b_ is the baseline mutation rate (mutation rate when allocation to germline maintenance is at its minimum, *u*_g_ = 0), and *α*_*μ*_ is the maintenance scaling factor (a parameter tuning how investing a unit resource into maintenance translates into reducing the mutation rate). We assume that *α*_*μ*_ *>* 1, such that *μ*(**u**) has decreasing negative slopes in *u*_g_ and hence exhibits diminishing returns from investment into germline maintenance. The parameter *α*_*μ*_ can thus be interpreted as the “efficiency” of converting resources to lowering the mutation rates (since higher values of *α*_*μ*_ correspond to lower mutation rates). The quantities *f*_0_(**u**) and *s*_0_(**u**) are, respectively the fecundity and survival of the least-loaded class and they are written as

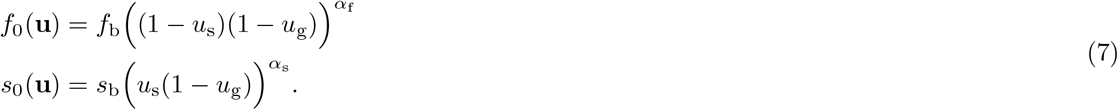

Here, *f*_b_ and *s*_b_ are, respectively, the baseline fecundity and baseline probability of survival; *α*_f_ and *α*_s_ are, respectively, the fecundity and survival scaling factors (parameters tuning how a unit resource translates into fecundity and survival). We assume that *α*_f_, *α*_s_ ≤ 1, whereby both survival and fecundity have decreasing positive slopes in the net amount of resources allocated to them and thus exhibit diminishing returns. Smaller values of *α*_f_ and *α*_s_ correspond to more efficient returns of investing resources into reproduction and survival, respectively.

In the absence of allocation to germline maintenance and deleterious mutations, the model defined by eqs. (6)–(7) reduces to the standard model of reproductive effort of life-history theory with a tradeoff between reproduction and survival (Charnov, 1993; Pen, 2000; Case, 2000). Conversely, with no overlapping generations and no life-history evolution, the model reduces to the classical model of mutation accumulation (Haigh, 1978; Bürger, 2000), and with zero survival and resource allocation evolution, it is equivalent to the asexual model of Dawson (1998). The model thus contains a trade-off between lifehistory traits (survival and reproduction) and immutability (germline maintenance) whose evolutionary consequences are unexplored.

#### 3.1.2 Uninvadable and convergence stable strategies

We now carry out the invasion analysis. To that end, we use eqs. (6)–(7) to first evaluate the basic reproductive number 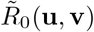 for this model (see Appendix B.1 where we also show that the equilibrium distribution **p**(**v**) of individuals with different numbers of mutations follows a Poisson distribution with mean *λ*(**v**) = *μ*(**v**)*/σ*, where *σ* = *σ*_f_ = *σ*_s_ and **v** = (*v*_g_, *v*_s_)). From eq. (B-4) we obtain that the selection pressure (e.g., Parker and Maynard Smith, 1990; Frank, 2008; Geritz et al., 1998; Rousset, 2004) on resource allocation to maintenance can be written as

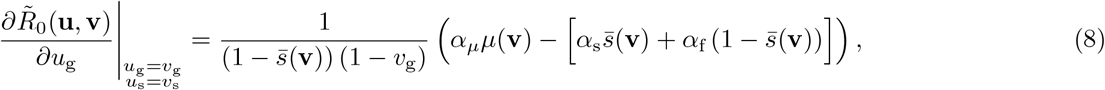

which displays a trade-off between allocating resources to maintenance vs into the two vital rates and where 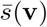 is the mean survival in the resident population (see eq. B-6 in Appendix B). All our mathematical computations can be followed and confirmed via an accompanying Supplementary Information, S.I. consisting of a Mathematica (Wolfram, 1991) notebook. The first term in eq. (8) is the marginal benefit of investment into repair, which is an increasing function of *μ*_b_ and *α*_*μ*_. The second term is the marginal cost of investment into maintenance, which depends on the weighted sum over average survival and open breeding spots. Thus we find that an increase in the baseline mutation rate and more efficient returns from investment into germline maintenance and vital rates promote allocation to maintenance (increasing *α*_*μ*_ and decreasing *α*_s_ and *α*_f_). Meanwhile, the selection pressure on resource allocation to survival can be written as

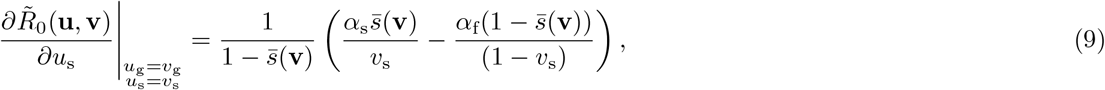

where the terms in the parenthesis display the trade-off between allocating resources into survival vs fecundity (i.e. the classical reproductive effort trade-off, e.g. Pen, 2000, eq. 4) with the difference that it is here affected by the mutation rate via 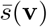. The first term in eq. (9) is the marginal benefit of investment into survival, which is a decreasing function of *v*_g_. In contrast, the second term is the marginal benefit of investment into fecundity, and it is an increasing function of *v*_g_. Thus, all else being equal, higher allocation to germline maintenance promotes fecundity over survival. We also find that an increase in the baseline mutation rate *μ*_b_(*v*_g_) favours higher allocation to survival (by increasing 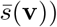).

A necessary condition for 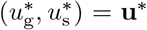 to be an evolutionary equilibrium is that the selection pressures vanish at this point (for an interior equilibrium, i.e. 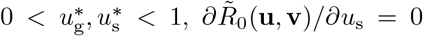 and 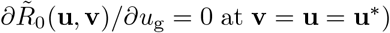. Without further assumptions on eqs. (8)–(9), we were unable to find such analytical solutions. However, when assuming that the efficiency of investing resources into fecundity and survival is the same (i.e. setting *α*_s_ = *α*_f_ = *α*), we find that the solution takes two forms, depending on the parameter values. First, when *μ*_b_ ≤ *α/α*_*μ*_, then 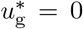 and 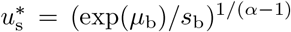. This means that when the baseline mutation rate *μ*_b_ is lower than the threshold *α/α*_*μ*_ given by the efficiency parameters, germline maintenance does not evolve. In this situation, the equilibrium mutation rate is equal to the baseline mutation rate *μ*(**u**^∗^) = *μ*_b_ and the mean number of novel mutations is *λ*(**u**^∗^) = *μ*_b_*/σ*, as expected when there is no germline maintenance. When the threshold *α/α*_*μ*_ is low, it indicates that the efficiency of investing in vital rates and germline maintenance is high. This is because smaller values of *α* and larger values of *α*_*μ*_ signal more efficient investment into vital rates and germline maintenance, respectively. So, when the efficiency of investing into vital rates and germline maintenance is sufficiently high (or, in other words, the threshold *α/α*_*μ*_ is sufficiently low) relative to the baseline mutation rate (when *μ*_b_ *> α/α*_*μ*_), then there is a unique interior equilibrium:

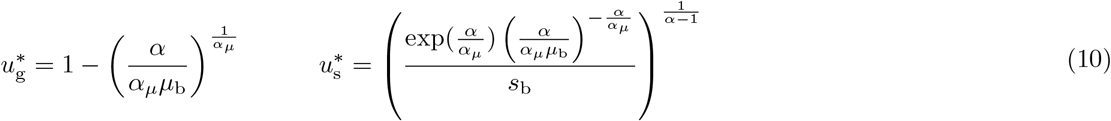

with corresponding expressions for the mutation rate *μ*(**u**^∗^) and mean number of novel (deleterious) mutations *λ*(**u**^∗^) taking the following form

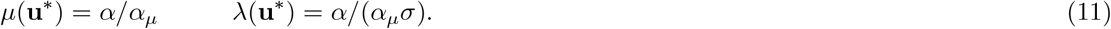

Fig. (2) illustrates these equilibrium strategies 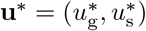 (panels a and b), the corresponding mutation rate *μ*(**u**^∗^) (panel c) and the mean number of novel mutations *λ*(**u**^∗^) (panel d) as a function of the baseline mutation rate *μ*_b_ for different values of the scaling parameter *α* of vital rates. Fig. (2) shows that investment into maintenance is higher when the baseline mutation rate *μ*_b_ increases and when investment into vital is more efficient (*α* ≪ 1).

**Figure 2:**
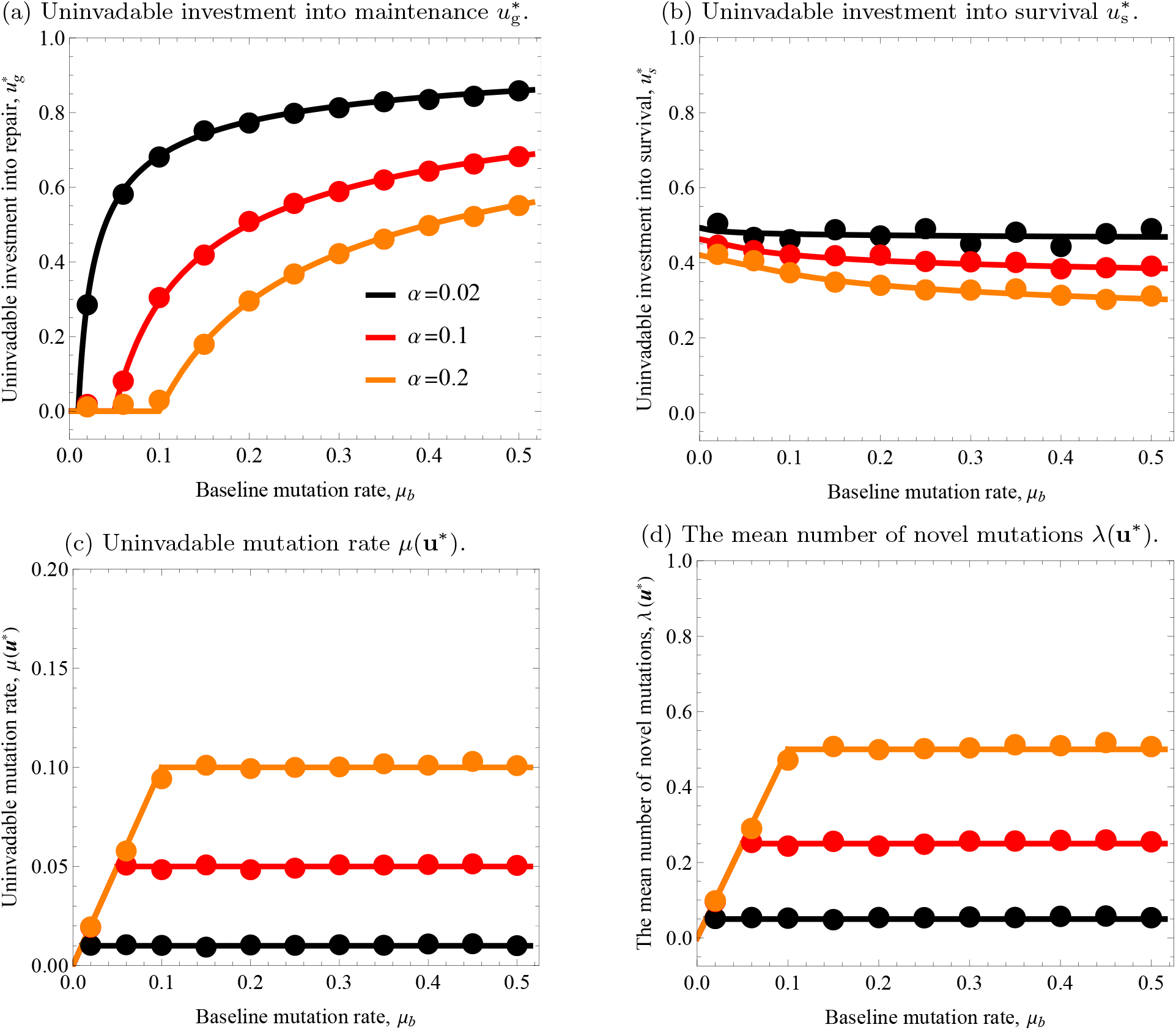
Predictions from the analytical model (solid lines) and from individual-based simulations of a finite population (circles) for the uninvadable life-history strategies 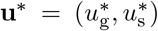 (panel a and b) and the mean number of novel mutations *λ*(**u**^∗^) (panel c) as functions of baseline mutation rate *μ*_b_. The solution for the individual-based simulations is a time-averaged mean values, measured over 7500 generations while starting the simulation at the analytically predicted equilibrium (see Appendix C for details about the simulations and Table C-4 for the time-average standard deviations from the mean; see S.I. for the simulation code). The different colours represent different values of the efficiency *α* of reproduction and survival (smaller values of *α* correspond to more efficient returns from investment into vital rates). Parameter values: *f*_b_ = 5, *α*_*μ*_ = 2, *s*_b_ = 0.5, *σ* = *σ*_f_ = *σ*_s_ = 0.2; for simulations: *N* = 7500, *f*_b_ = 5, *s*_b_ = 0.5, the mutations in the life-history locus follow a Normal distribution with zero mean and a standard deviation of 0.1.

Three main conclusions can be drawn from this analysis. First, selection favours physiologically costly germline maintenance at the expense of lowering investment into vital rates (survival and reproduction) when the baseline mutation rate is high enough (whenever *μ*_b_ *> α/α*_*μ*_). Second, when germline maintenance evolves, the mutation rate, *μ*(**u**^∗^), depends only on the efficiency parameters (*α* and *α*_*μ*_) and is independent of the baseline mutation rate *μ*_b_ (see Fig. 2c, eq. 11, and additional figures in section 1.5.5. of S.I.). This is so in this model because the effect of *μ*_b_ on the cost of germline maintenance via the expected survival 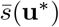 cancels out due to the nature of density-dependence (decrease in expected survival is cancelled out by the increase in the expectation of acquiring a breeding spot; see eq. (8) when taking *α*_s_ = *α*_f_ = *α*). Third, higher allocation to reproduction at the expense of survival occurs as *μ*_b_ increases (Fig. 2b). This is so because the effect of the mutation rate on fitness is similar to that of external mortality and thus decreases the value of allocating resources to survival. As a result, reproduction is prioritised when *μ*_b_ is large. This effect is more pronounced when investment into vital rates and maintenance is less efficient (*α* is larger or *α*_*μ*_ is smaller; see also section 1.5.5. in the S.I.). We also find that immortality (complete survival, 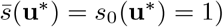) can evolve in our model only in the absence of external mortality (*s*_b_ = 1) and zero baseline mutation rate (*μ*_b_ = 0, see eq. 10). In section 1.1.4. of S.I., we numerically checked that our results are qualitatively robust when relaxing the assumption that the scaling factors of investment into reproduction and survival are not equal *α*_f_≠ *α*_s_.

Using standard local analysis (e.g., Eshel, 1983; Taylor, 1989; Geritz et al., 1998; Mullon et al., 2016 and see section 1.5.3. of the S.I.), we checked that the strategy **u**^∗^ defined by eq. (10) is also locally uninvadable for biologically realistic parameter values (e.g. for the parameter values in Fig. 2) and in which case **u**^∗^ is also necessarily locally convergence stable (see argument after eq. B-4). Since there is a unique solution, a locally uninvadable **u**^∗^ is also globally uninvadable (e.g.,(Sydsaeter et al., 2008)) and thus globally convergence stable. The results from individual-based stochastic simulations, Fig. (3) confirm that the co-evolutionary dynamics indeed converge towards the uninvadable strategy **u**^∗^ (eq. 10) predicted by the analytical model. We can observe from Fig. (2) that the analytically predicted trait values (eq. 10–11) correspond closely to the mean trait values observed in the simulations of the full evolutionary process with reasonably small population sizes, and which implement the assumptions of the biological scenarios (section 3.1.1) but allows for mutation at the life-history locus (e.g. Fig. 1 and see Appendix C.1 for the description of the simulations, Table C-4 for the standard deviations around the mean traits, and the S.I. for the Mathematica code of the simulations). We observed that simulation results generally matched well with the analytical predictions when the selection coefficient is one order of magnitude larger than the baseline mutation rate (e.g., recall the first paragraph of section 2.3.2).

**Figure 3:**
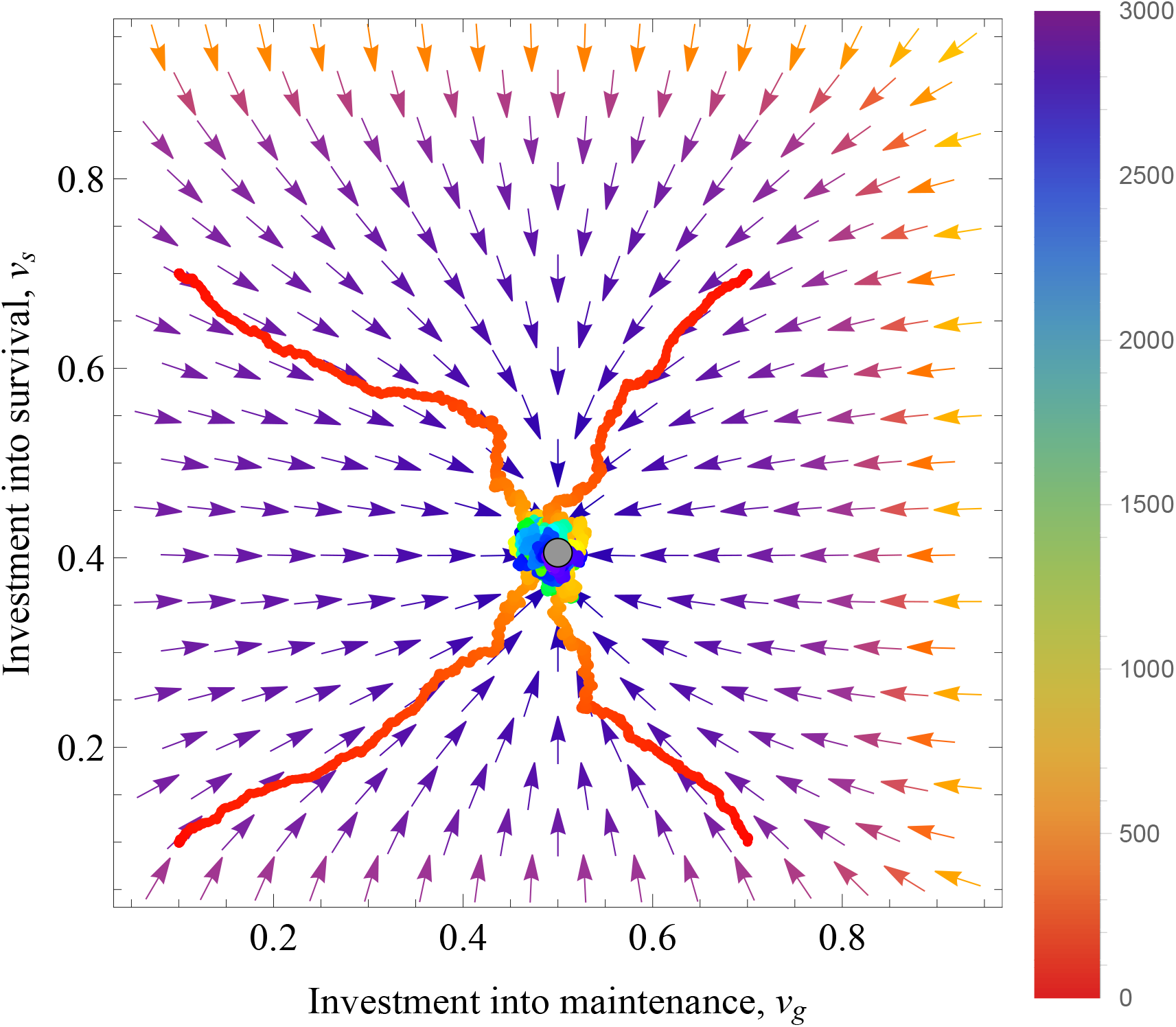
Evolutionary convergence towards the uninvadable life-history strategy 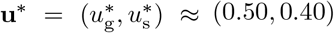 (grey circle). The arrows give the analytic direction of selection at any population state (eqs. 8 and 9) and the colourful jagged lines represent the evolution of population average trait values over evolutionary time in simulations (from initial time, up to 3000 generations). Simulation were started from four different initial conditions: (i) *v*_g_ = 0.1, *v*_s_ = 0.1, (ii) *v*_g_ = 0.1, *v*_s_ = 0.7, (iii) *v*_g_ = 0.7, *v*_s_ = 0.1, and (iv) *v*_g_ = 0.7, *v*_s_ = 0.7. The colour of jagged lines indicates the number of generations since the start of the simulation (the color bar on the right-hand-side indicates the number of generations). The simulations indicate that the population converges close to the uninvadable strategy within 3000 generations. Parameter values: *f*_b_ = 5, *α*_*μ*_ = 2, *s*_b_ = 0.5, *σ* = *σ*_f_ = *σ*_s_ = 0.2; for simulations: *N* = 6000, *f*_b_ = 5, *s*_b_ = 0.5, the mutations in the life-history locus follow a Normal distribution with zero mean and a standard deviation of 0.05.

### 3.2 Coevolution of age at maturity and germline maintenance

#### 3.2.1 Biological scenario

Our second scenario considers the evolution of age-at-maturity when mutation accumulation can occur during growth and reproduction. To that end, we consider that age is continuous, and each individual undergoes the following events. (1) An individual is born and grows in size until it reaches maturity (growth phase). (2) At maturity, an individual starts to reproduce at a constant rate, and fecundity is assumed to be density- and size-dependent (reproductive phase). (3) Throughout their lives, individuals can die at some constant rate and acquire germline mutations. We postulate that individuals have again a life-history trait consisting of two components **u** = (*u*_g_, *u*_m_), where *u*_g_ is the allocation to germline maintenance (lowering the mutation rate) and *u*_m_ is the age-at-maturity (see Table 3 for a summary of key symbols for this example). The life-history trait **u** determines how resources are allocated between three physiological functions: (i) a proportion *u*_g_ of resources is allocated to the maintenance of the germline at any age *a*, (ii) a proportion (1 − *u*_g_) of resources are allocated to growth when an individual is of age *a < u*_m_, (iii) a proportion (1 − *u*_g_) of resources is allocated to reproduction when an individual is at age *a* ≥ *u*_m_, (hence **u** ∈ 𝒰[𝒯] = [0, 1] × ℛ ^+^).

**Table 3:** List of key symbols of “Coevolution of age at maturity and germline maintenance model”.

| Symbols for “Coevolution of age at maturity and germline maintenance model”. |  |
| --- | --- |
| $u_g, v_g$ | Proportional allocation of resources to germline maintenance of mutant and resident individual, respectively. |
| $u_m, v_m$ | Proportional allocation of resources to germline maintenance of mutant and resident individual, respectively. |
| $N(\mathbf{v})$ | Total population size; endogeneously determined and thus depends on the resident trait $\mathbf{v}$. |
| $x(t)$ | Body size of a mutant individual at age $t$. |
| $x_m(\mathbf{u})$ | Body size of a mutant individual at maturity. |
| $B(x_m(\mathbf{u}))$ | Surplus energy rate, i.e., rate of energy available to be allocated to life-history functions; we assume that the surplus energy scales as the power with size, i.e. energy available to mature individuals is $B(x_m(\mathbf{u})) = ax_m(\mathbf{u})^c$. |
| $\mu_f$ | Rate at which germline mutations appear in an offspring when giving birth (fixed parameter, $\mu_f = 0$ in simulations). |
| $\mu_s(u_g)$ | Rate at which germline mutations appear in a mutant individual over time (independent of age). |
| $\mu_b$ | Baseline mutation rate at which germline mutations appear; mutation rate, when no resources are allocated into germline maintenance. |
| $d_b$ | Baseline death rate. |

We assume that the effective fecundity, death, and mutation rate of an individual with trait **u** and *i* deleterious mutations in a population with resident trait **v** = (*v*_g_, *v*_m_) is given by

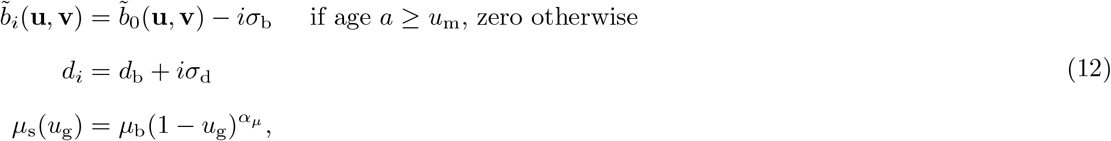

where 0 ≤ *σ*_b_ and 0 ≤ *σ*_d_ are, respectively, the additive effects on fecundity and death rate from carrying deleterious mutations. The death rate of an individual of the least-loaded class is given by the baseline death rate *d*_b_ and the effective birth rate of zero-class individuals of the least-loaded class is given by

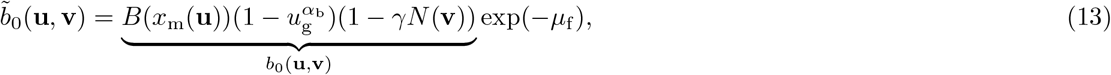

where *B*(*x*_m_(**u**)) = *ax*_m_(**u**)^*c*^ is the surplus energy rate at maturity, i.e., the rate of energy available to be invested into life-history functions and germline maintenance, where *x*_m_(**u**) is size-at-maturity and *a >* 0 and 0 *< c <* 1 are parameters. Here, 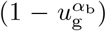 represents how reproduction depends on the allocation strategy, and 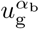 represents the cost of reproduction when allocating a proportion *u*_g_ of resources to germline maintenance. The parameter *α*_b_ is a scaling factor (*α*_b_ *>* 1 corresponds to diminishing returns of investing resources into reproduction), and *α*_b_ can be interpreted as the efficiency parameter. The term (1−*γN* (**v**)) accounts for density-dependent regulation of reproduction, where *N* (**v**) is the total population size of the resident population, which can be solved analytically regardless of the way deleterious mutations affect survival and reproduction (see eq. (B-8) of Appendix B.2), and *γ* tunes the intensity of density dependence. Finally, exp(−*μ*_f_) is the probability that the offspring do not acquire new mutations during reproduction where the mutation rate at giving birth *μ*_f_ is assumed constant. In order to close the expression for the birth rate, we need an explicit expression for size at maturity *x*_m_(**u**). During the growth phase, we postulate that size follows the differential equation

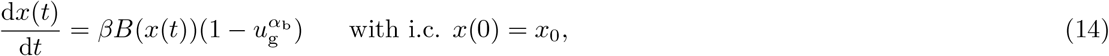

where *B*(*x*(*t*)) is the surplus energy rate at age *t* and 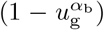 represents the proportional allocation of resources devoted towards growth (instead of repair). For tractability, we assume that 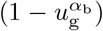 has the same functional form as the proportional allocation towards reproduction (eq. 13) and *β* allows to tune how much resources are needed to grow one unit, compared to the resources needed to produce one offspring. We further assume that the surplus energy rate is given by the power law *B*(*x*(*t*)) = *ax*(*t*)^*c*^, which is considered to be appropriate for modelling size/age-at-maturity under determinate growth (see Day and Taylor (1996) for a justification). It follows from integrating eq. (14) that the size at maturity takes the form

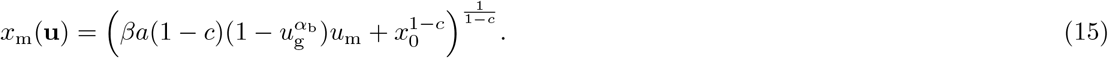

In the absence of mutation rate, the model reduces to the standard model of age-at-maturity (Kozlowski, 1992; Day and Taylor, 1997; Stearns, 1992; Roff, 2008). The model thus contains a trade-off between life-history traits (growth and reproduction) and immutability (germline maintenance and repair) whose evolutionary consequences have not been explored.

#### 3.2.2 Uninvadable and convergence stable strategies

Let us now carry out the invasion analysis for which we first evaluate the basic reproductive number for this model (see Appendix B.2). Then using eq. (B-8) along with eq. (15), we find that the selection pressure on resource allocation to germline maintenance can be written as

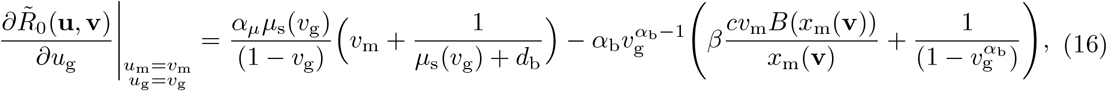

where the two terms display the trade-off between allocating resources into maintenance vs. growth and reproduction. The first term is the marginal benefit of investing into germline maintenance and it increases with the age-of-maturity and the expected lifespan. The second term is the marginal cost of investing into maintenance, which is a weighted sum of the expected loss in growth and reproduction. The marginal cost of maintenance is smaller when size-at-maturity *x*_m_(**v**) is larger (since *B*(*x*_m_(**v**))*/x*_m_(**v**) decreases with *x*_m_(**v**) for 0 *< c <* 1). This implies that all else being equal, organisms that grow larger should invest more into germline maintenance. Note that the selection gradient is independent of the mutation rate *μ*_f_ at giving birth. We find that the selection pressure on the age-at-maturity can be written as

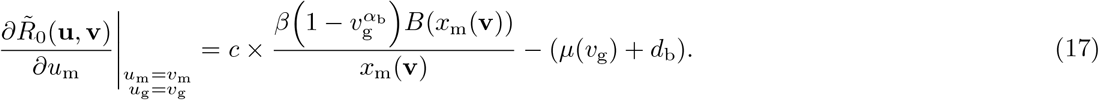

The first term is the marginal benefit of investment into growth and thus the benefit for maturing later, while the second term is the marginal cost of investment into growth and thus the benefit for maturing earlier. We can see that the increase in mutation rate will select for earlier age-at-maturity. This implies that organisms with lower germline mutation rate can grow larger.

By first solving 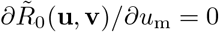 for 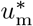 when evaluated at **v** = **u** = **u**^∗^, we obtain

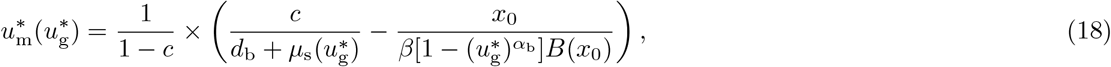

which is a function 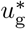. Eq. (18) says that individuals tend to mature later, when individuals growth rate at birth 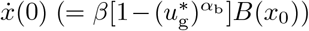 is higher and/or when death rate *d*_b_, mutation rate 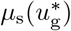, and birth size *x*_0_ are smaller (holding everything else constant). When *μ*_b_ → 0 and 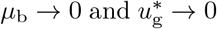, age-at-maturity reduces to 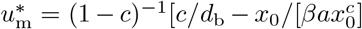, which is consistent with standard results about the optimal age/size at maturity (see e.g. Day and Taylor, 1996) and it is useful to compare how an allocation to germline maintenance affects the age-at-maturity. In order to determine the joint equilibrium 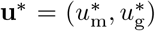, we need to substitute eq. (18) into eq. (17) and solve for 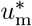 and 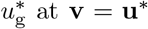 at **v** = **u**^∗^. We were unable to obtain an analytical solution for the general case. But restricting attention to *α*_*μ*_ = *α*_b_ = 2 (i.e. assuming diminishing returns of investment into germline maintenance and reproduction) and biologically feasible set of solutions (such that 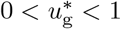 and 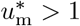 *>* 1; see section 2.1.3 of S.I. for calculations), we find that there is a unique interior equilibrium

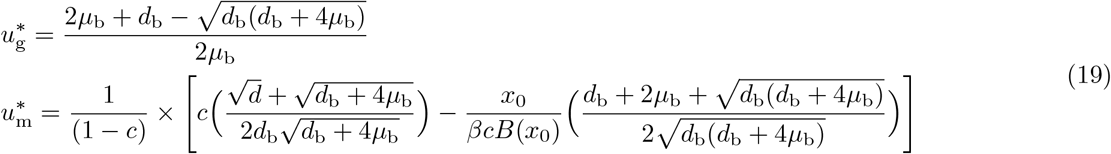

with the corresponding mutation rate given by

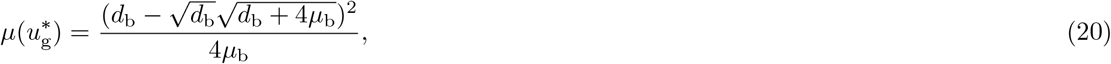

while the corresponding population size *N* (**u**^∗^) can also be explicitly expressed in terms of model parameters (see eq. B-8). Fig. 4 provides a graphical depiction of the equilibrium strategy **u**^∗^ as a function of the baseline mutation rate (panels (a) and (b)) and the corresponding equilibrium population size and mutation rate (panels (c) and (d)). Fig. (2) shows that that investment into maintenance is higher when the baseline mutation rate is high and when external mortality is low.

**Figure 4:**
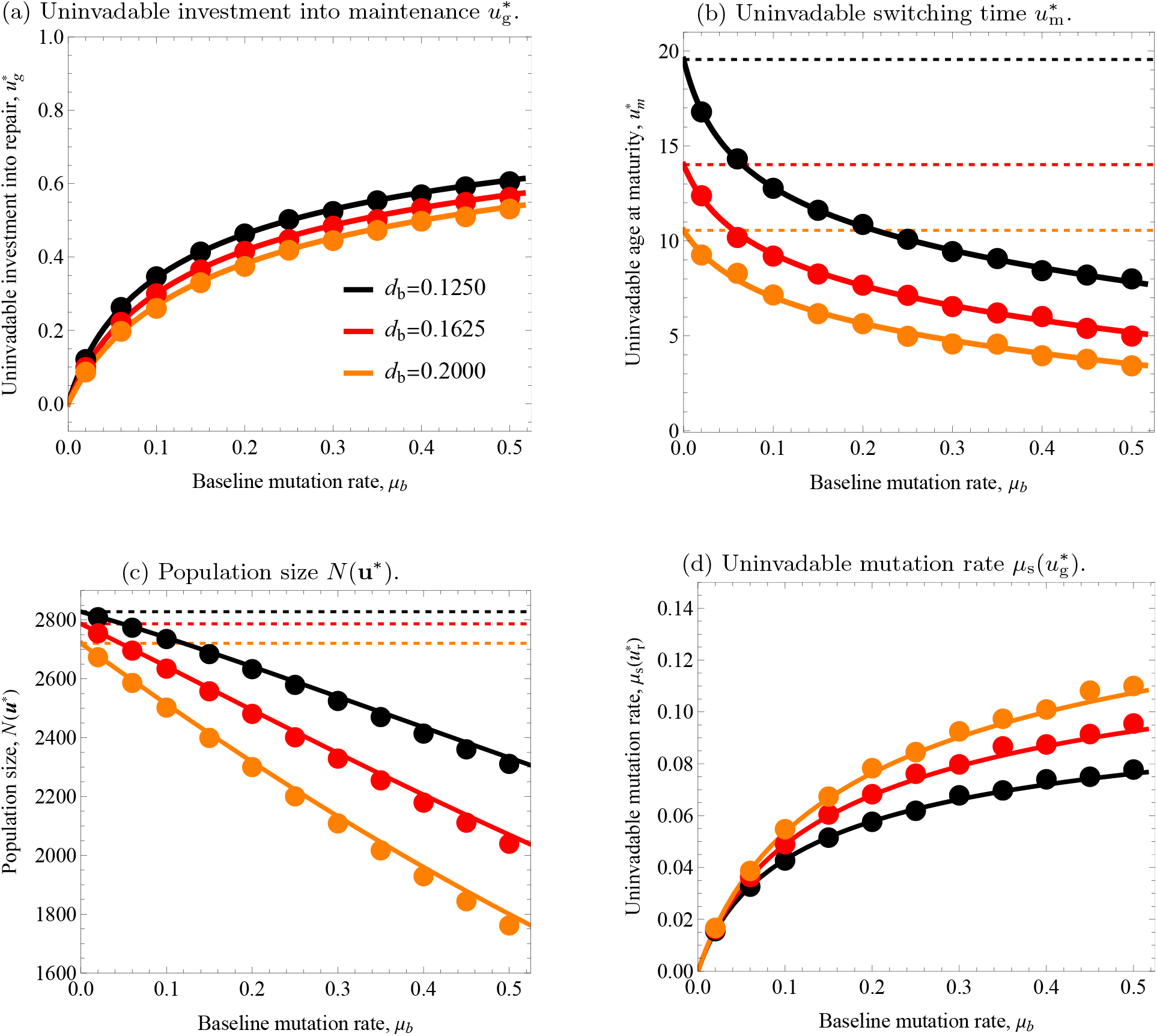
Predictions from the analytical model (solid lines) and from individual-based simulations (circles obtained as averages) for uninvadable life-history strategies 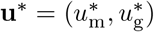 (panel a and b), population size *N* (**u**^∗^) (panel c) and mutation rate 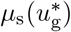 as functions of baseline mutation rate *μ*_b_ for different values of baseline mortality *d*_b_ (*d*_b_ = 0.1250 - black, *d*_b_ = 0.1625 - red, *d*_b_ = 0.2 - orange). The dashed lines represent the “classical life-history” prediction (i.e. when 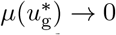 and 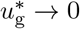), where the colours of the dashed represent the different values for baseline death rate *d*_b_ parameter and match the values of solid lines (*d*_b_ = 0.1250 - black, *d*_b_ = 0.1625 - red, *d*_b_ = 0.2 - orange). The solution for the individual-based simulations are obtained as time-averaged mean values measured over 3000 “generations” while starting the simulation at analytically predicted equilibrium for the trait values and population size (Table C-5 for the time-average standard deviations from the mean; see S.I. section 2.3 for the code and for more details). The different colours represent different values of baseline mortality rate *d*_b_. Parameter values: *σ* = *σ*_b_ = *σ*_d_ = 0.2, *x*_0_ = 1, *a* = 0.9, *c* = 0.75, *γ* = 0.00035, *β* = 1, *μ*_f_ = 0, the mutations in the life-history locus follows a Normal distribution with zero mean and a standard deviation of 0.07.

Three main results can be drawn from this analysis. First, as in the previous example, selection favours physiologically costly germline maintenance at the expense of lowering the investment into lifehistory functions (here, into growth and reproduction, see Fig. 4a). Here, the uninvadable mutation rate (*μ*(**u**^∗^)) monotonically increases with the baseline mutation rate (Fig. 4d). Second, we predict earlier age-at-maturity when the baseline mutation rate is high (Fig. 4b). In fact, the baseline mutation rate and external mortality have qualitatively similar effects on fitness as they increase the marginal cost of investment into growth (see the last term in eq. 17). Thus, we find that the shift in growth-reproduction trade-off towards reproduction is higher under: (i) high external mortality rates and (ii) high baseline mutation rates. Since maturing earlier causes the growth period to be shorter, the body size at maturity *x*_m_(**u**^∗^) will also be smaller with a higher baseline mutation rate *μ*_b_ (Fig. 6a). Smaller body size at maturity, in turn, causes the birth rate *b*_0_(**u**^∗^, **u**^∗^) to be smaller (Fig. 6b). Third, a higher baseline mutation rate causes a smaller equilibrium population size (Fig. 4c).

The equilibrium strategy 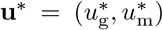 determined by eq. (19) is convergence stable and locally uninvadable (see section 2.5.4. and 2.5.5. in the S.I. for derivation for the parameter values in Fig. (4)). Since eq. (19) is a unique equilibrium for the feasible trait space (see section 2.1.3 of S.I.), then local uninvadability implies global uninvadability. Using individual-based stochastic simulations, we were able to confirm that 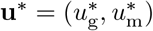 given in eq. (19) is indeed a stable attractor of the evolutionary dynamics (see Fig. 5 for a graphical depiction of convergence in the individual-based simulations for four different initial population states). Fig. (4) also illustrates the equilibrium population size *N* (**u**^∗^) (panel c), and the uninvadable mutation rate 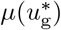 (panel d) as a function of the baseline mutation rate *μ*_b_. Fig. (6) illustrates the body size at maturity *x*_m_(**u**^∗^) (panel a) and the effective birth rate 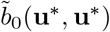 at the uninvadable population state as a function of baseline mutation rate. Overall, Fig. (4) reaffirms that the analytically predicted trait values (here using eqs. 19–20) correspond very closely to the mean trait values observed through individual-based simulations of the full evolutionary process (see Appendix C.2 for the description of the simulations, Table C-5 for the standard deviations around the mean traits, and section 2.3. of the S.I. file for the Mathematica code for the simulations).

**Figure 5:**
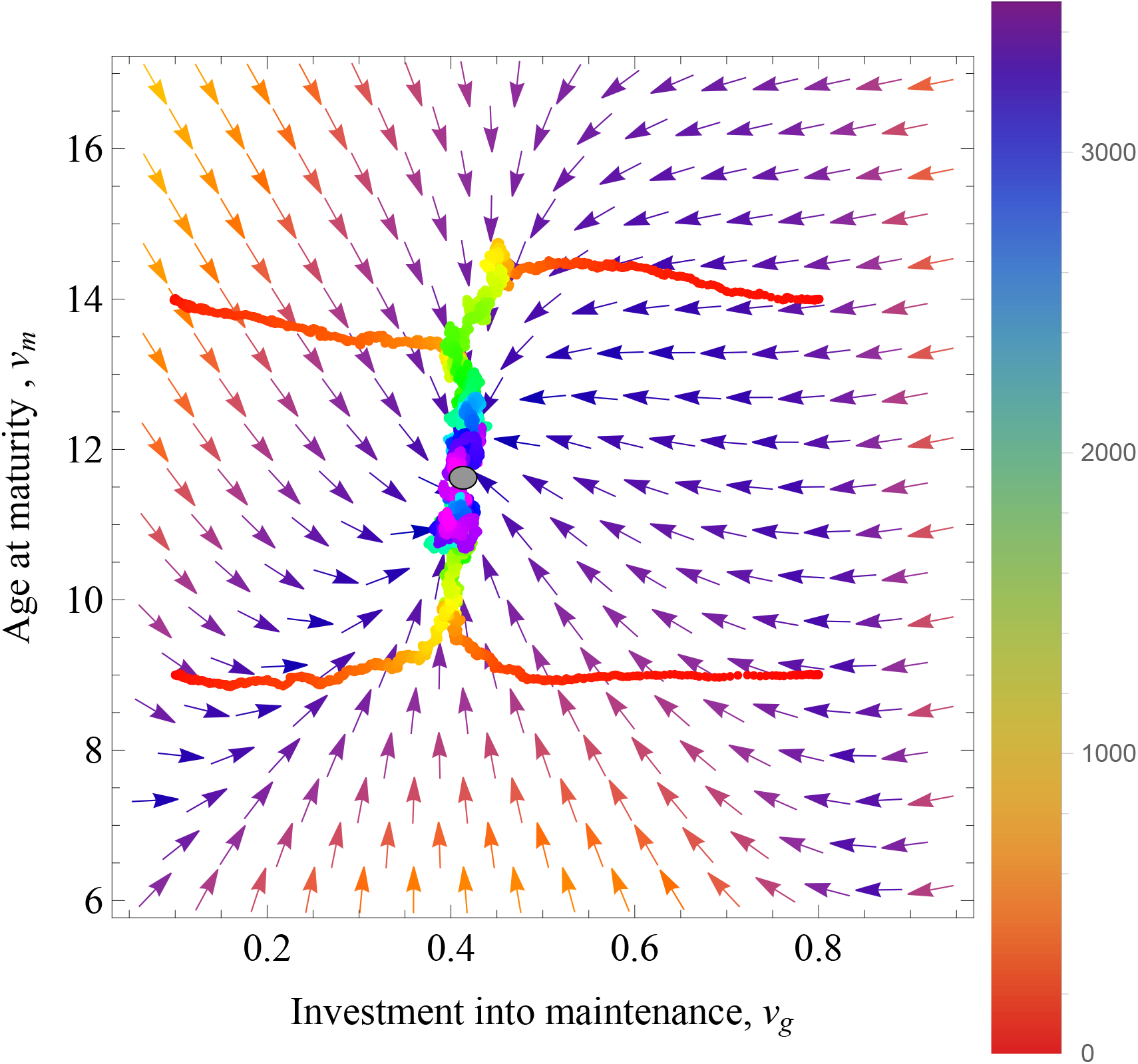
Evolutionary convergence to the uninvadable life-history strategy 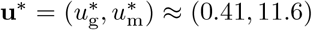 (grey circle). The arrows give the direction of selection at any resident population state (eqs. 16 and 17) and the colourful jagged lines represent the evolution of the population average trait values over evolutionary time in simulations (from initial time, up to 3500 “generations”). Simulations were started from four different initial conditions: (i) *v*_g_ = 0.1, *v*_m_ = 9, (ii) *v*_g_ = 0.1, *v*_m_ = 14, (iii) *v*_g_ = 0.8, *v*_m_ = 9, and (iv) *v*_g_ = 0.8, *v*_s_ = 14. The colour of jagged lines indicates the number of generations since the start of the simulation (the color bar on the right-hand-side indicates the number of generations). The simulations indicate that the population converges close to the uninvadable strategy within 3500 generations. Parameter values: *σ* = *σ*_b_ = *σ*_d_ = 0.2, *x*_0_ = 1, *a* = 0.9, *c* = 0.75, *γ* = 0.00035, *β* = 1, *μ*_f_ = 0, *d*_b_ = 0.1250, the mutations in the life-history locus follow a Normal distribution with zero mean and a standard deviation of 0.07.

**Figure 6:**
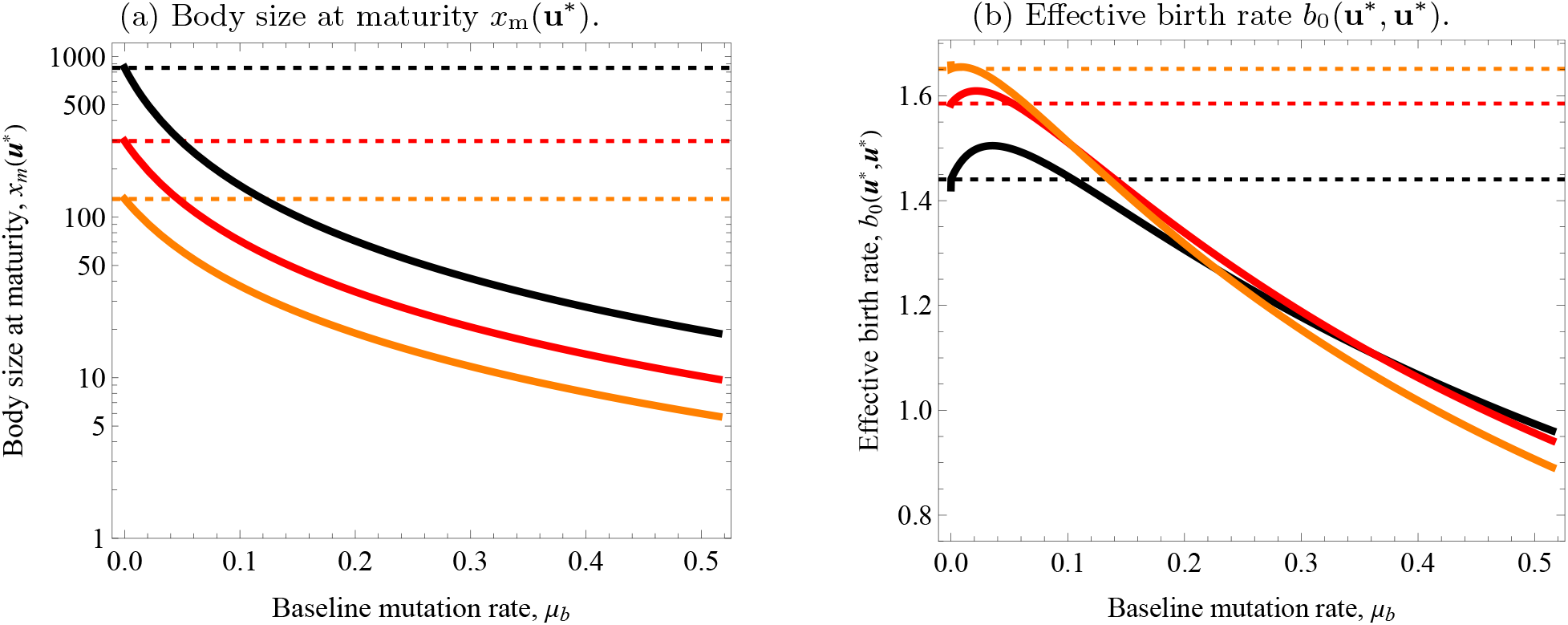
Predictions from the analytical model for the body size at maturity *x*_m_(**u**^∗^) and the effective birth rate *b*_0_(**u**^∗^, **u**^∗^) at the uninvadable population state as a function of baseline mutation rate for different values of mortality rate *d*_b_ (*d*_b_ = 0.1250 - black, *d*_b_ = 0.1625 - red, *d*_b_ = 0.2 - black). The dashed lines represent the “classical life-history” prediction (i.e. when 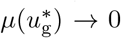 and 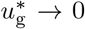), where the colours of the dashed represent the different values for *d*_b_ parameter and match the values of solid lines (*d*_b_ = 0.1250 - black, *d*_b_ = 0.1625 - red, *d*_b_ = 0.2 - black). Parameter values: *σ* = *σ*_b_ = *σ*_d_ = 0.2, *x*_0_ = 1, *a* = 0.9, *c* = 0.75, *γ* = 0.00035, *β* = 1, *μ*_f_ = 0.

## 4 Discussion

We formalised selection on resource allocation traits affecting life history and the deleterious mutation rate under asexual reproduction. When selection against deleterious mutations is high compared to the deleterious mutation rate, the class of individuals having the fewest such mutations (the least-loaded class) dominates in frequency the population and thus determines the fate of any mutant modifier allele affecting the deleterious mutation rate and life history. Then, we showed that the basic reproductive number of the least-loaded class (eq. 1 and eq. 4) allows to characterise the joint evolutionary stable lifehistory and deleterious mutation rate. We analysed two specific applications to illustrate this approach: (1) joint evolution between reproductive effort and the mutation rate and (2) joint evolution between the age-at-maturity and the mutation rate. These two models confirmed the validity of using the least-loaded class as a fitness proxy by comparing results to those obtained by individual-based stochastic simulations (Figs. 2–5) and provide several insights about the joint evolution of life-history and deleterious mutation rate.

The first model shows that a positive deleterious mutation rate evolves when selection against increasing the mutation rate is balanced by the cost of germline maintenance. This extends established results (Kimura, 1967; Kondrashov, 1995; Dawson, 1998, 1999) to an explicit life history context. We find that the trade-off between reproduction and survival shifts towards higher allocation into reproduction under a high baseline mutation rate. This confirms the numerical observation of Charlesworth (1990) that a higher level of a fixed mutation rate (no germline maintenance) causes higher allocation to reproduction over survival. We further predict that the shift in survival-reproduction trade-off towards reproduction is stronger: (i) when the conversion of resources into vital and germline maintenance is less efficient (e.g. in environments where organisms have high maintenance costs, e.g., colder climates), (ii) high external mortality rates (e.g., high predation environment), and (iii) high baseline mutation rates (e.g., induced by environmental stressors). We also find that immortality (complete survival) cannot evolve even in an environment with no external mortality because the mutation rate cannot be brought down to zero. This highlights a potentially overlooked role of mutation accumulation, which, alongside extrinsic environmental hazards (Medawar, 1952; Hamilton, 1966; Charlesworth, 1994), can prevent the evolution towards immortality. This means that the forces of direct selection on survival and reproduction, which are decreasing as a function of the death rate (Hamilton, 1966; Ronce and Promislow, 2010) will also be declining as a function of the mutation rate. Overall, this example reveals that endogenous and/or exogenous factors that increase the baseline mutation rate cause lower lifespans through higher allocation to fecundity.

The model examining the combined evolution of age-at-maturity and mutation rate also indicates that a stable positive mutation rate evolves. However, in this scenario, it emerges because of a balance between germline maintenance and investing in growth and reproduction. We predict that a higher baseline mutation rate favours smaller size-at-maturity marked by an earlier onset of the reproductive stage. This complements the results of Dańko et al. (2012), where a higher fixed mutation rate causes earlier maturity, although they report the effect of mutation accumulation as minor. In contrast, we show that mutation accumulation can significantly affect life-history trade-offs since allocation to germline maintenance co-evolves with life history. Using individual-based simulations, the joint evolution between somatic maintenance, germline maintenance, size-at-maturity, and population size has also been explored by Rozhok and DeGregori (2019) who found that selection for larger body size (by imposing size-dependent mortality) can lead to higher germline mutation rate because more resources need to be invested into somatic maintenance. Thus, they found that a higher germline mutation rate and sizeat-maturity are expected to be positively correlated, which is an opposite prediction from our result. Further studies clarifying the selection pressures involved in the trade-off between growth and investment into the maintenance of germline and soma could thus shed light on how patterns of body size, longevity and mutation rate are expected to be correlated.

Our second model also predicts that higher mutation rates correlate with smaller equilibrium population sizes, supporting previous findings (e.g. Gabriel et al., 1993). An increased baseline mutation rate can thus amplify the effect of genetic drift, especially in small populations. This could intensify genetic drift’s influence on mutation accumulation (Lynch et al., 2016), and potentially even trigger the mutational meltdown of the (asexual) population (Gabriel et al., 1993). As discussed in section 2.3.1, if selection against deleterious mutations is significantly larger than the mutation rate in models ignoring age-specific effects, the impact of this should be negligible. Thus, the deterministic approximation for the resident mutation-selection balance should then fare well, even under relatively strong genetic drift. This is the case in our stochastic individual-based simulations that further and more importantly show that the mean of the allocation traits align with the analytical predictions of the evolutionary stable trait values (e.g., Figure 4 where the population size is about 2000 individuals at the evolutionary equilibrium). This is a robust feature of quantitative trait evolution models (see references section 2.2), where even in small populations, the mean observed trait value in individual-based simulations can be well predicted by analytical approximations (e.g. even in populations with less than 10 individuals Wakano and Lehmann, 2012, Fig. 1 but where the variance in trait values can become large). This ultimately stems from the fact that the selection pressures obtained from invasion fitness are proportional to those obtained from fixation probabilities in the absence of social interactions and so population size does not qualitatively affect the direction of selection (e.g., Rousset, 2004). In summary, we thus expect that the resource allocation predictions from our two concrete applications are robust to the effect of genetic drift as long as selection against deleterious mutations is at least one order of magnitude larger than the mutation rate.

Two main findings about how life history coevolves with deleterious mutation rate emerge from these applications. First, the trade-off between lowering the rate of mutations vs investing in life-history functions affects the evolutionary outcome of life-history trade-offs (e.g. survival–vs–reproduction or growth–vs–reproduction). Hence, mutation accumulation can have a significant effect on life-history evolution through the process of joint evolution that previous models focusing on the effect of fixed mutation rates on life-history evolution have not revealed (Charlesworth, 1990; Dańko et al., 2012). Looking at the effect of fixed mutation rates on life history evolution underestimates the effect of deleterious mutation accumulation on life history evolution, as it does not consider the physiological cost of immutability on life history evolution. Second, factors that contribute to higher baseline mutation rate select for faster life histories: higher investment into current reproduction at the expense of survival and earlier age– at–maturity. Factors that could increase the baseline mutation rate include factors that increase DNA replication errors (number of germ-line cell divisions) or environmental mutagens (oxygen level, nutrition quality, see e.g. Ferenci, 2019 for a review).

Our analysis using the basic reproductive number of the least-loaded class to locate evolutionary stable resource allocation strategies under deleterious mutation accumulation relies on a number of simplifying assumptions. Most notably (i) a separation between life-history traits (e.g., the timing of reproduction, age and size at maturity, longevity) and traits that could be referred to as viability traits, such as morphology and physiology, (ii) that mutations are only deleterious at the viability traits, and (iii) that reproduction is asexual. The separation between life-history traits and viability traits allows us to focus on the life-history resource allocation trade-off and avoids modelling the viability traits mechanistically. This separation makes biological sense for organisms that have similar viability traits but differ in their life history. For instance, both annual and perennial plants can have similar cellular pathways for photosynthesis and oxidative phosphorylation, where deleterious mutations can affect the functionality of these pathways that are under strong selection. For such organisms, our model thus consists of characterising the optimal life history and how the mutation rate at the cellular machinery evolves. While other formulations are possible and could be explored in future work, our approach has analytical traction and is conceptually equivalent to the separation between modifier locus and loci affecting vital rates from modifier theory, where modifier alleles affect the pattern of transmission of other traits (e.g., Leigh, 1970; Altenberg, 2009). In our model, modifier alleles are also under direct selection owing to their effect on life history. From modifier theory, we know that regardless of the details of transmission; namely, regardless of whether reproduction is asexual or sexual or the exact pattern of mutation rates (whether they are reducible or not), a decrease in the mutation rate is always favoured in the absence of trade-off with reproduction (Altenberg, 2009; Altenberg et al., 2017). Thus, in the presence of a trade-off in any genetic system, the evolutionary stable mutation rate will be determined by the balance between the benefit of lowering the mutation rate and the benefit of increasing the mutation rate and thus reducing the physiological cost of germline maintenance (e.g. André and Godelle, 2006; Kondrashov, 1995; Dawson, 1998). Under sexual reproduction, however, fewer resources will be allocated to germline maintenance (Kondrashov, 1995; Dawson, 1998; André and Godelle, 2006; Altenberg, 2009) because deleterious mutations are linked under asexual reproduction, and this linkage is broken down by recombination under sexual reproduction whereby the benefit of lowering the mutation is smaller (Dawson, 1998; Gervais and Roze, 2017). However, any direct effect that the modifier has would be experienced in the same way regardless of the reproductive system. In summary, while genetic details might be important to compare the quantitative effect of the reproductive system on the joint evolution of mutation rate and life history, they are unlikely to qualitatively affect our main prediction that the cost of germline maintenance can be a significant force affecting life-history traits.

Our model can be further extended to study several open questions in life-history theory and mutation accumulation theory. First, our model can be applied to understand how allocation to germline maintenance affects the evolution of ageing. This can be done by explicitly applying our model in the scenario where deleterious mutations can have age-specific effects and/or allow for allocation to somatic repair. Current theories of ageing either rely on accumulation of late-acting deleterious mutations in the germline (e.g. Medawar, 1952; Lehtonen, 2020) or are based on a trade-off between reproduction and survival (disposable soma and antagonistic pleiotropy theories of ageing; Williams, 1957; Kirkwood, 1977; Cichon and Kozlowski, 2000), but no theoretical study has explored how these processes evolve jointly when germline maintenance is costly. Second, our model can also be extended to study the Lansing effect, a widely reported negative effect of parental age on offspring fitness (e.g. Monaghan et al., 2020). Third, extending our model for sexual reproduction would allow studying how sex-specific differences in germline maintenance can induce sex differences in life-history trade-offs (Maklakov and Lummaa, 2013). Our hope is that the formalisation proposed in this paper can be fruitfully used to these ends.

## Appendix A Invasion process with mutation accumulation

In this appendix, we prove that the basic reproductive number 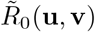 defined by eq. (1) (in discrete-time) and eq. (4) (in continuous-time) is an appropriate invasion fitness proxy when the least-loaded class dominates in frequency the resident population. To that end, following the assumptions of the main text section 2.1, we first characterise the mutant invasion process and use renewal equations for both discrete and continuous-time to show that 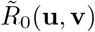 is an appropriate invasion fitness proxy (section Appendix A.1). These renewal equations are derived from full stochastic considerations on birth and death by using age-dependent branching process theory (Appendix A.2).

### Appendix A.1 Reducible mutant invasion process

Since the resident population is assumed to have reached a mutation-selection balance and is thus at a genetic-demographic equilibrium, i.e. the standard internal stability assumptions of invasion analysis (Eshel and Feldman, 1984; Altenberg et al., 2017; Metz, 2011), the mutant allele for trait **u** can arise in individuals carrying different numbers of deleterious mutations. Hence, the invasion process of **u** is contingent on the genetic background in which it arises. We refer to the initial carrier of the **u** trait as the progenitor (or ancestor) of **u**. The invasion fitness of mutant **u** is then determined by the size of the lineage of the progenitor, which consists of all of its descendants carrying **u** far into the future. Namely, the immediate descendants of the progenitor, including the surviving self, the immediate descendants of the immediate descendants, etc., covering the whole family history tree *ad infinitum*. Noteworthy, descendants may accumulate deleterious mutations during the initial growth or extinction phase of the mutant lineage when this lineage is rare, which we refer throughout as the “invasion process”. As such, the mutant invasion process, whether in continuous or discrete time, can be regarded as a multitype age-dependent branching process (Mode, 1968, 1971) since during the growth or extinction of the mutant lineage, novel genotypes are produced by mutation.

To analyse this invasion process, it is useful to organise individuals into *equivalence classes*. The defining feature of an equivalence class is that it is a collection of states of a process among which transitions eventually occur, so the states are said to communicate (Karlin and Taylor, 1975, p. 60). The equivalence class C_*i*_ will stand for all mutant individuals carrying *i* deleterious mutations and thus consist of individuals of all ages. This is an equivalence class because, through survival and reproduction, an individual of any age with *i* mutations may eventually transition to become an individual of any other age (in the absence of menopause). This follows from the fact that the process of survival and reproduction in an age-structured population in the absence of mutations (and menopause) is *irreducible* (Karlin and Taylor, 1975, p. 60), since starting in any age-class, eventually, every age class can be reached by the members of a lineage of individuals. Owing to our assumptions that mutations are deleterious and can only accumulate, however, starting in a given mutational class, it is possible to enter another class, but not transition back from that class (otherwise the two classes would form a single class). Thereby the mutation process is reducible, since a lineage of individuals cannot go from any mutational class to any other mutation class. Equivalence class C_*i*+1_ is then said to follow class C_*i*_ since individuals in equivalence class *i* can only transition to class *i* + 1 by acquiring mutations. This means that the invasion process is reducible and can be regarded as a reducible multitype age-dependent branching process (Nair and Mode, 1971; Mode, 1971). Reducibility typically arises in population genetics models without back-mutations but sometimes also occurs in models in ecology and demography, e.g. when some class of individuals do not contribute to reproduction, as is, for instance, the case under menopause (e.g. Caswell, 2000; Altenberg, 2009; Bode et al., 2006; Stott et al., 2010; McDonald, 2015).

To see why the notion of an equivalence class is useful to understand the mutant invasion process, let us first focus on a discrete-time process with *T* discrete age classes and denote by *n*_*i*_(*t*) the expected number of individuals at time *t* = 0, 1, 2, …, in class C_*i*_ that descend from a single class C_*i*_ newborn progenitor born at *t* = 0 (i.e. *n*_*i*_(0) = 1). Thus, *n*_*i*_(*t*) is the expected lineage size of class C_*i*_ individuals descending from a newborn progenitor of class C_*i*_ (including the surviving self). Accounting entails that *n*_*i*_(*t*) satisfies the renewal equation

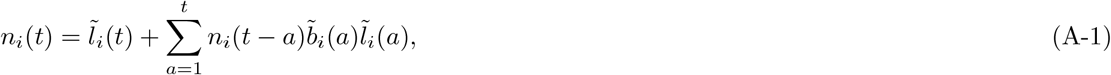

where 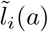 is the probability that a C_*i*_ class newborn survives to the *a*-th age class and has not acquired any new mutation, and 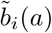 is the expected number of (newborn) offspring without mutations produced by an individual belonging to the *a*-th age class and being of mutation class 𝒞_*i*_. Eq. (A-1) is derived in Appendix A.2.1 from branching process considerations and its left-hand side can be understood as follows. First, 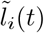 accounts for the survival and immutability of the progenitor itself until age *t*. Second, 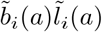 is the progenitor’s expected number of offspring of class 𝒞_*i*_ born during the time interval corresponding to the *a*-th age class and each of these newborns contribute an expected number *n*_*i*_(*t* − *a*) of class 𝒞_*i*_ individuals to the progenitor’s total lineage size at *t*. This is so because as long as the mutant is rare, each newborn starts a new independent lineage and adding all terms together, the right-hand side of eq. (A-1) thus gives the total lineage size of the progenitor (see Appendix A.2.1 for more details).

A key feature of eq. (A-1) is that it depends only on the vital rates and states of individuals of class 𝒞_*i*_. As such, eq. (A-1) is functionally equivalent to the standard renewal equation of population dynamics in discrete age-structured populations (Charlesworth, 1994, eq. 1.34), but recall that *n*_*i*_(*t*) counts total lineage size in class 𝒞_*i*_ and not zygote size as in the classical theory of age-structured populations. It then follows from the classical theory of age-structured populations (e.g., Charlesworth, 1994, p. 25-26) or the branching process formulation (Mode, 1974) that asymptotically, as *t* → ∞, the number *n*_*i*_(*t*) grows geometrically as

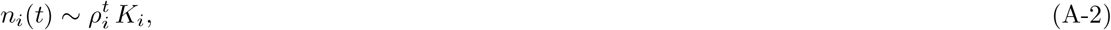

where *K*_*i*_ is a constant depending on the process and *ρ*_*i*_ is the unique root satisfying the characteristic (or Euler-Lotka) equation 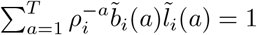.

Since individuals of class *i* contribute to individuals of class *i* + 1 through mutations, the equivalence class 𝒞_*i*+1_ follows class 𝒞_*i*_, then *n*_*i*_(*t*) does not describe the total expected lineage size of the progenitor. However, owing to the assumptions that mutations are deleterious and can only accumulate, the growth ratio *ρ*_*i*_ is at least as large as *ρ*_*i*+1_, i.e., *ρ*_*i*_ ≥ *ρ*_*i*+1_ for all *i*. This implies that when the ancestor is of type *i*, the expected lineage size is determined by the growth ratio *ρ*_*i*_, since it dominates that of any other following equivalence class. Hence, asymptotically, the total expected lineage of an 𝒞_*i*_ class progenitor has a geometric growth ratio *ρ*_*i*_. It further follows from the theory of multitype age-dependent branching processes that the realised lineage size of a single progenitor (a random variable) has growth ratio *ρ*_*i*_ if *ρ*_*i*_ *>* 1 and otherwise if *ρ*_*i*_ ≤ 1, the lineage goes extinct with probability one (Mode, 1971, Theorem 7.2 p. 245, Corrolary 6.1 p. 280, see also Mode, 1974 for the single type case). Further, *ρ*_*i*_ ≤ 1 if and only if 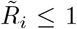, where 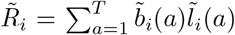 is the expected number of offspring of a progenitor of type *i* produced throughout its lifespan (i.e. Mode, 1971, Theorem 7.2 p. 245, Corrolary 6.1 p. 280, see also Karlin and Taylor, 1981, p. 424, Caswell, 2000). Hence, *ρ*_*i*_ is an appropriate measure of invasion fitness and 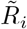 is an appropriate proxy of it for a type *i* mutant **u** arising in a resident **v** background. The same argument can be made for continuous-time processes, in which case, the invasion fitness of a mutant arising in a progenitor in class C_*i*_ is *ρ*_*i*_ = exp (*r*_*i*_), where *r*_*i*_ is the rate of natural increase of the lineage size of a progenitor of type *i*, i.e., the Malthusian growth rate and for which 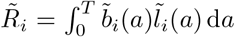 is an appropriate invasion fitness proxy (see Appendix A.2.2 for a derivation).

The key feature of the invasion process in a population with distinct mutational equivalence classes is that the invasion fitness of a mutant life history trait thus depends on the class in which it appears (*ρ*_*i*_ for class *i*), which in turn depends on the distribution **p**(**v**). This means that there are as many growth rates as equivalence classes, since the invasion process is reducible (see also Altenberg, 2009, p. 1278). Therefore characterising long-term evolution using a single representation of invasion fitness (or proxy thereof) is at first glance unattainable as it requires to make the process irreducible, which can be achieved by introducing back mutations or recombination. Reaching irreducibility in this way, however, would make the model much more complicated. It also follows from these considerations that when the least-loaded class dominates in frequency the population, the face of mutant **u** appearing in a resident **v** population is determined from the growth ratio *ρ*_0_(**u, v**) of the least-loaded class only, since any mutant will appear on the C_0_ background and so this is an appropriate overall measure of invasion fitness (*ρ*(**u, v**) = *ρ*_0_(**u, v**)). Further, since 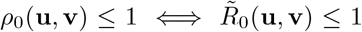, where 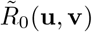 is the basic reproductive number of the least-loaded class, i.e. the expected number of class C_0_ offspring produced by a class individual C_0_ individual over its lifespan, is sufficient to characterise the fate of the mutant. This then justifies using 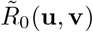 as an invasion fitness proxy defined by eq. (1) and eq. (4) for discrete and continuous time, respectively. Further, if deleterious mutations are such that all the *ρ*_*i*_’s are proportional to *ρ*_0_’s, which is the case for the standard mutation accumulation models with a multiplicative effect of (deleterious) mutations, then using 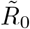 does no rely on making the assumption of low mutation rates relative to selection, since regardless in which background the mutation appears, it will grow proportionally to *ρ*_0_.

#### Appendix A.2 Renewal equations from age-dependent branching process

We here derive the renewal eq. (A-1) for a discrete-time process and its continuous-time analogue from underlying age-dependent branching process assumptions (Crump and Mode, 1968; Mode, 1968, 1971). As such, we consider a full stochastic model of survival and reproduction to describe the fate, invasion or extinction, of a mutation introduced as a single copy in a resident population. The defining assumption of a branching process is that a new and independent copy of the process begins each time a new individual is born (Harris, 1963; Mode, 1968, 1971).

To describe the branching process under our assumptions of the main text section 2.1, let us introduce the following notations, which will apply to both the discrete and continuous time processes. First, let *Z*_*i*_(*t*) stand for the random number of individuals in class 𝒞_*i*_ at time *t* that descend from a single newborn (age class zero) mutant progenitor in class 𝒞_*i*_ at *t* = 0, whereby *Z*_*i*_(0) = 1. Second, denote by *L*_*i*_ the random sojourn time of a progenitor of class 𝒞_*i*_ in that class; the progenitor exits class 𝒞_*i*_ either because of death (in which case *L*_*i*_ is the lifespan) and/or because a germline mutation occurred. Third, let

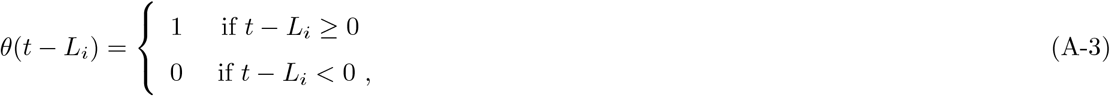

by the indicator random variable describing whether a progenitor of class 𝒞_*i*_ has exited class 𝒞_*i*_ by time *t* (either by death and/or mutation). Namely, if *θ*(*t* − *L*_*i*_) = 0, then the progenitor is still in class 𝒞_*i*_ at *t* (alive without having mutated). Finally, let *n*_*i*_(*t*) = E[*Z*_*i*_(*t*)] denote the expected number of individuals at time *t* in class 𝒞_*i*_ that descend from a single class 𝒞_*i*_ newborn progenitor at *t* = 0, where the expectation is overall stochastic events affecting survival and reproduction (we refer to Crump and Mode, 1968; Mode, 1971 for a construction of the probability spaces for branching processes, here we simply assume they exist).

#### Appendix A.2.1 Discrete time

Let us now focus on the discrete-time process where *t* = 0, 1, 2, … and further introduce the random number *W*_*i*_(*a*) of offspring without new mutations born to the progenitor of class 𝒞_*i*_ during the time interval corresponding to the *a*-th age group where *a* = 1, 2, … so that, recall, we begin counting discrete age classes with 1 (one of the two possible conventions to count discrete age classes, Fig. 3.1 Case, 2000). Because a branching process starts afresh with the birth of any offspring, the random variable *Z*_*i*_(*t*) satisfies the discrete time stochastic renewal equation

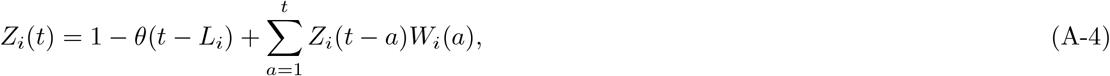

which can be understood as follows. First, if *θ*(*t* − *L*_*i*_) = 0, then the progenitor is still in class 𝒞_*i*_ at *t* and thus contributes one individual to its lineage. Second, each offspring produced without mutation during the time interval corresponding to the *a*-th age class starts a new independent branching process and thus contributes *Z*_*i*_(*t*−*a*) individuals of class 𝒞_*i*_ to the progenitor’s lineage at *t*. Summing over all offspring and all age groups the progenitors can be in, we obtain the second term in eq. (A-4). In other words, the sum 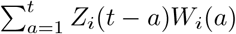 is the total random number of class 𝒞_*i*_ individuals at *t* that descendent from each of the offspring of the progenitor including the surviving selves (i.e., offspring, offspring from offspring, offspring from offspring from offspring, etc). Thus, the right-hand side of eq. (A-4) is the progenitor’s random lineage size at *t*. Bearing differences of notations and context, eq. (A-4) is conceptually equivalent to eq. (5.6) of Mode (1974).

Taking the expectation over realizations of the stochastic process over both sides of eq. (A-4) yields

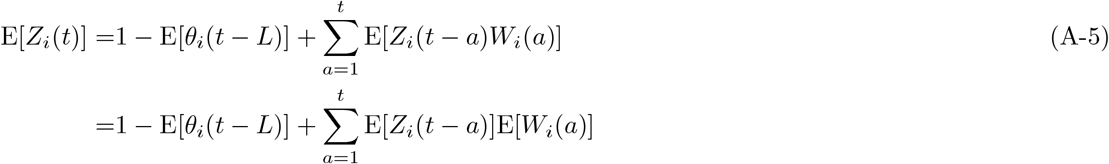

where the first equality follows from the linearity of the expectation operator and the second equality follows from the assumption of the independence of each new branching process. In order to further simplify this expression, we first note that the probability that the progenitor is still of class 𝒞_*i*_ at *t* is 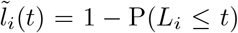, and that by standard probability arguments, P(*L*_*i*_ ≤ *t*) is the expectation of the indicator random variable of the event that the progenitor exits class 𝒞_*i*_ at *t*: E[*θ*(*t* − *L*_*i*_)] = P(*L*_*i*_ ≤ *t*) (Grimmett and Stirzaker, 2001). Second, we assume stochastic independence between survival and reproduction, whereby the expected number of offspring without mutations born to a progenitor of class 𝒞_*i*_ during the time interval corresponding to the *a*-th age class is 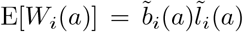, which is the probability 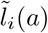 that the progenitor has survived until age class *a* without mutating times the expected number 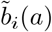 of its offspring without mutations produced when residing in the *a*-th age class. Substituting these quantities into eq. (A-5) and noting that, by definition, *n*_*i*_(*t* − *a*) = E[*Z*_*i*_(*t* − *a*)], eq. (A-5) can be written 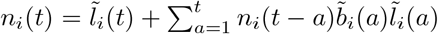, which is eq. (A-1). Bearing differences of notations and context, eq. (A-5) is conceptually equivalent to eq. (5.11) of Mode (1974).

#### Appendix A.2.2 Continuous time

Let us now focus on the continuous time process where *t* ∈ [0, ∞) and for this case, let *N*_*i*_(*t*) denote the random number of offspring without mutations produced until time *t* by a progenitor of class 𝒞_*i*_, where these offspring are produced at the random times *T*_1(*i*)_ ≤ *T*_2(*i*)_ ≤ … ≤ *T*_*Ni*(*t*)_ with *T*_*j*(*i*)_ being the random time until production of the *j*-th offspring. Thus, *N*_*i*_(*t*) is a renewal counting (or point) process (Karlin and Taylor, 1975, p. 31) with rate function 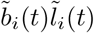, where 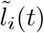 is the probability that a progenitor of class 𝒞_*i*_ is still in that class at time *t* (same as in the discrete-time case), and 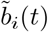 is the birth rate of offspring without mutations at *t*. Because a branching process starts again with the birth of each offspring, the random variable *Z*_*i*_(*t*) satisfies under the continuous time process the stochastic renewal equation

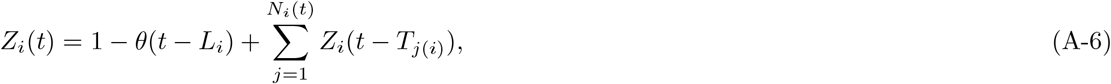

where, as in the discrete case, 1 − *θ*(*t* − *L*_*i*_) counts the surviving progenitor without mutation and the sum counts the total number of class 𝒞_*i*_ individuals at *t* that descend from each of the offspring of the progenitor, where the progenitor’s offspring are produced at random times. Bearing differences of notations and context, eq. (A-6) is conceptually equivalent to eq. (3.1) of Crump and Mode (1968).

Taking the expectation over realizations of the stochastic process over both sides of eq. (A-6) yields

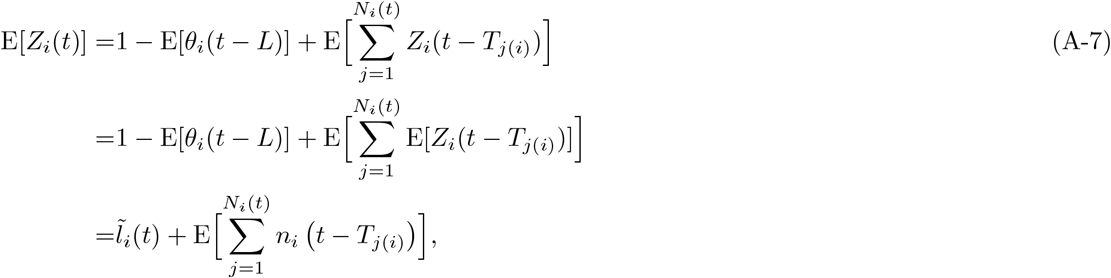

where the second line follows from the independence of each new branching process and the third from the definitions introduced above. Now, owing to Campbell’s theorem relating the rate of a renewal counting process, here 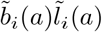, and the expectation of a random sum of a function over the process (e.g., Kingman, 1992, p. 28), we have 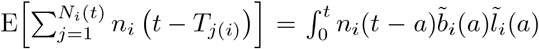 and therefore eq. (A-7) becomes

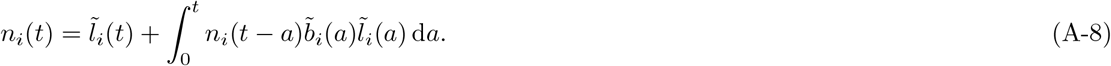

Bearing differences of notations and context, eq. (A-8) is conceptually equivalent to eq. (6.1) of Crump and Mode (1968) once the assumptions of their example 8.2 is endorsed, i.e., stochastic independence between survival and reproduction.

Eq. (A-8) is also functionally equivalent to the standard renewal equation of population dynamics for continous age-structured populations (Charlesworth, 1994, eq. 1.41). As such, and as for the discrete-time case, it then follows from the standard results of population dynamic processes in age-structured populations (Charlesworth, 1994, p. 27) or from the continuous time branching process formulation (Crump and Mode, 1968) that asymptotically, as *t* → ∞, the number *n*_*i*_(*t*) grows geometrically as

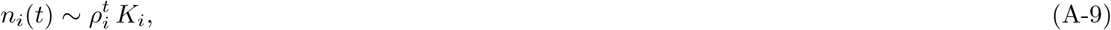

where *K*_*i*_ is some constant depending on the process and *ρ*_*i*_ = exp(*r*_*i*_), where *r*_*i*_ is the mutant growth rate (or Malthusian parameter), which is the unique root of the Euler-Lotka equation

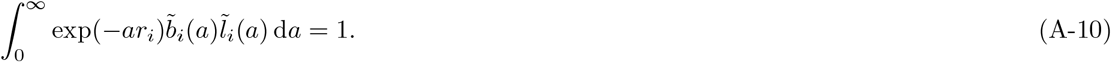

Thereby 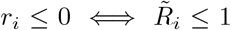 with 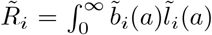 being the basic reproductive number of a class *i* individual (Karlin and Taylor, 1981, p. 424, Mode, 1971, Theorem 7.2 p. 245, Corrolary 6.1 p. 280).

## Appendix B Basic reproductive numbers

In this appendix, we present the explicit expressions for 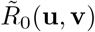 for our two biological scenarios

### Appendix B.1 Coevolution of reproductive effort and germline maintenance

From the model assumptions given in main text section 3.1.1, we have that a juvenile surviving density-dependent competition, has a survival probability of one to the adult stage and that each adult of the least-loaded class survives to the next generation with probability *s*_0_(**u**). In the meantime, the probability that each type of individual has not acquired a germline mutation is exp (−*μ*(**u**)). Therefore, for this model, eq. (3) becomes

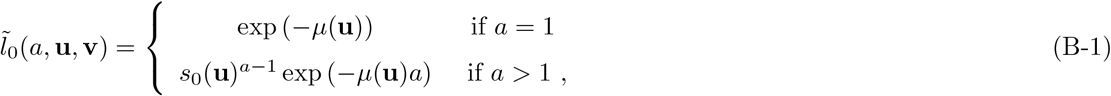

while the effective fecundity of the least-loaded class (eq. 2) is

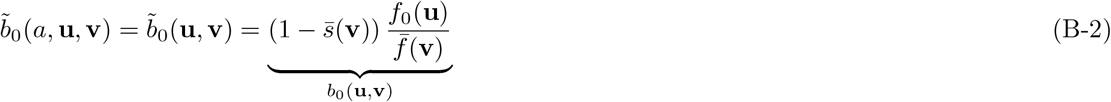

since no mutation occurs specifically during reproduction (in addition of the germline mutation mentioned above). The effective fecundity depends on the mean survival and fecundity in the population, respectively, 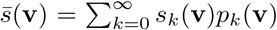 and 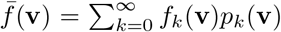. Here, *p*_*i*_(**v**) is the probability that an individual randomly sampled from the resident population carries *i* deleterious mutations (and so **p**(**v**) = {*p*_*i*_(**v**)}_*i*∈N_ for this model). This can be understood by noting that 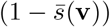 is the fraction of open breeding spots available to a juvenile and the probability that the offspring of a given adult acquires a breeding spot depends on the fecundity of the adult relative to the population average fecundity (as each juvenile is equally likely to acquire a breeding spot).

Since there is no fixed end to lifespan under the above life-cycle assumptions (so *T* → ∞), the basic reproductive number of the least-loaded class, eq. (1), is

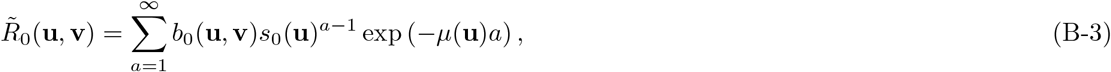

which yields

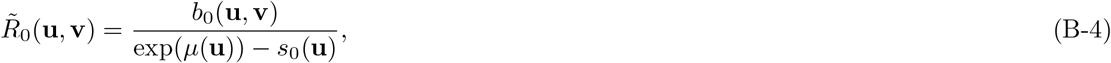

(see also the accompanying Mathematica notebook S.I.). Since *b*_0_(**u, v**) is multiplicatively separable with respect to its arguments, then it follows from eq. (B-4) that the model satisfies the condition of an optimisation principle (e.g., Metz et al., 2008). Namely, 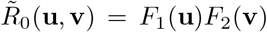 for the functions *F*_1_(**u**) = *f*_0_(**u**)*/*[exp(*μ*(**u**)) − *s*_0_(**u**)] depending only on the mutant and 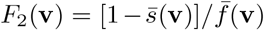 depending only on the resident. It follows that maximising *F*_1_(**u**) is sufficient to ascertain uninvadability (Metz et al., 2008). An optimization principle further entails that uninvadability implies (absolute) convergence stability, since the selection gradient 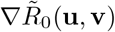 points in the same direction as the gradient ∇*F*_1_(**u**) (Leimar, 2009b, p. 199). Finally, the explicit expressions for 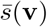 and 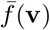, and thus the distribution **p**(**v**) are not needed to carry out the invasion analysis. This allows us to markedly simplify the evolutionary analysis. But we will nevertheless work out the resident distribution **p**(**v**) = {*p*_*k*_(**v**)}_*k*∈N_, where *p*_*k*_(**v**) is the frequency of individuals with *k* deleterious mutations, so as to have a fully worked example that allows for consistency checks and illustrating the concepts.

Since we consider a deterministic resident population process, the frequency *p*_*k*_(**v**) satisfies at equilibrium the equation

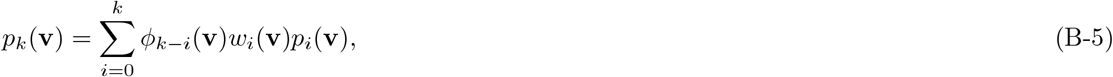

where 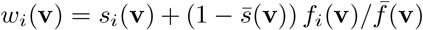 is the individual fitness– survival plus effective fecundity–of an individual with *i* deleterious mutations, and *ϕ*_*k*_(**v**) is the probability that *k* deleterious mutations are produced upon reproduction. Assuming that the mutation distribution is Poisson with mean *μ*(**v**) and *σ*_s_ = *σ*_f_ = *σ*, then eq. (B-5) becomes structurally equivalent to eq. (1) of Haigh (1978) and eq. (5.3) of Bürger (2000, p. 300) (with mean fitness 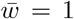 since population size is constant) and as such the equilibrium distribution **p**(**v**) is Poisson with mean *λ*(**v**) = *μ*(**v**)*/σ* (see also the section 1.1.1. in SM). This completely characterises the genetic state of the resident population and implies that

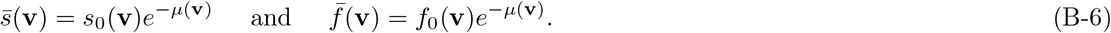

Substituting the explicit expression for the survival and effective fecundity (eq. B-6) into eq. (B-2) and eq. (B-4) shows that in a monomorphic **v** population 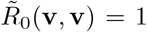, as required for a consistent model formulation. Eq. (B-6) generalises the standard mutation-accumulation model of population genetics to overlapping generations with survival probability depending on the number of deleterious mutations (see e.g. eq. 3.3 Kimura and Maruyama, 1966).

### Appendix B.2 Coevolution of age at maturity and germline maintenance

From the model assumptions given in the main text section 3.2.1 and using eq. (4) (under *T* → ∞ since there is also no definite end to lifepan), the basic reproductive number of the least-loaded class reduces to

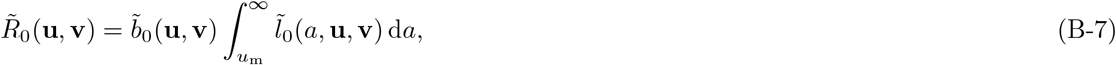

where 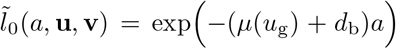. Substituting the expression for eq. (13) into eq. (B-7) and integrating yields

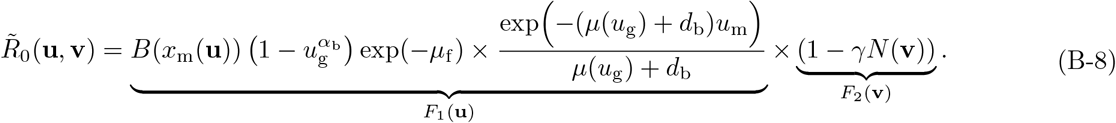

This shows that one can again express the basic reproductive number as a product of the form 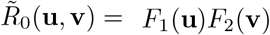 and thus this model has also an optimisation principle. This means that evaluating *N* (**v**) explicitly is not needed to ascertain uninvadability since *N* (**v**) only affects *F*_2_(**v**) (and uninvadability will again imply convergence stability for this model). We will nevertheless work the explicit expression for *N* (**v**) out as it is useful to understand how model parameters affect equilibrium population size. For this, it suffices to note that in a monomorphic resident population at a joint demographic and genetic equilibrium, each individual belonging to the least-loaded class must leave on average one descendant with zero new mutations. Hence 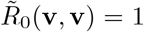 implies that that the population size at the demographic steady state is

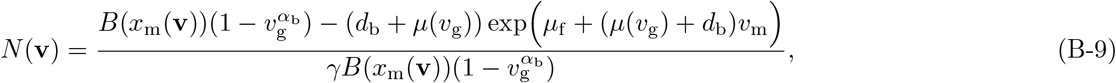

which holds regardless of the effects of deleterious mutations on the vital rates. This is a demographic representation and generalisation of the surprising simple result noted for unstructured semelparous populations of constant size that the nature of epistasis of deleterious mutations has no effect on the genetic load (Kimura and Maruyama, 1966; Gillespie, 2004). Eq. (B-9) also allows to evaluate the equilibrium population size *N* (**u**^∗^) at the evolutionary equilibrium in terms of model parameters by setting *v* = **u**^∗^ and substituting eqs. (19)–(20). The resulting expression is complicated and given in section 2.1.4. of the S.I.

## Appendix C Individual-based simulations

We here describe how we carried out the individual-based (stochastic) simulations used for the two model examples in the main text. The simulation algorithms scrupulously implement the life-cycle assumption of these models with the only difference that the mutation rate at the life-history locus is positive *μ*_*LH*_ *>* 0 (but kept small) in the simulations. This makes the evolutionary process in the simulations irreducible (see also discussion section 2.3.2) and subject to genetic drift along with mutation and natural selection.

### Appendix C.1 Coevolution of reproductive effort and germline maintenance

The simulation algorithm for this scenario (see section 1.3. of the S.I for the Mathematica code) follows a population composed of a finite and fixed number N (=7500 in the simulations) of individuals, where each individual is described by its genetic state (vector of traits consisting of allocation to maintenance, allocation to survival and number of deleterious mutations the individual has). One life-cycle iteration then proceeds as follows. We start by computing the fecundity of each adult individual, which is determined by its trait values (eq. 6). Then, we evaluate the survival probability of each adult individual according to its trait values (the survival of an individual is given by a Bernoulli random variable with mean given by its survival probability eq. 6). After eliminating the dead individuals, we fill the “vacated breeding spots” by randomly sampling offspring from the relative fecundity of all adult individuals before survival, thus effectively implementing a Wright-Fisher process for reproduction (Mode and Gallop, 2008). Once a newborn is chosen to fill the breeding spot, each of its traits mutates independently with probability (*μ*_LH_ = 0.01 in our actual simulations). The effect size of a mutation follows a Normal distribution with zero mean and a standard deviation (=0.1 in our simulations). Finally, we allow deleterious mutations to accumulate at the deleterious mutation locus according to a Poisson distribution with a mean that depends on the life-history locus (as specified by eq. 6). To obtain the results shown in Fig. 2, we initialised the simulation with a monomorphic population, with no deleterious mutations and life-history trait values given by the analytically predicted equilibrium. In Table C-4, we have depicted the time-averaged standard deviations from the main trait for parameter values in Fig. (4). In Fig. (3) we demonstrate the convergence stability of our simulations and we started the simulations away from the equilibrium for four different initial values of the traits.

**Table C-4:**
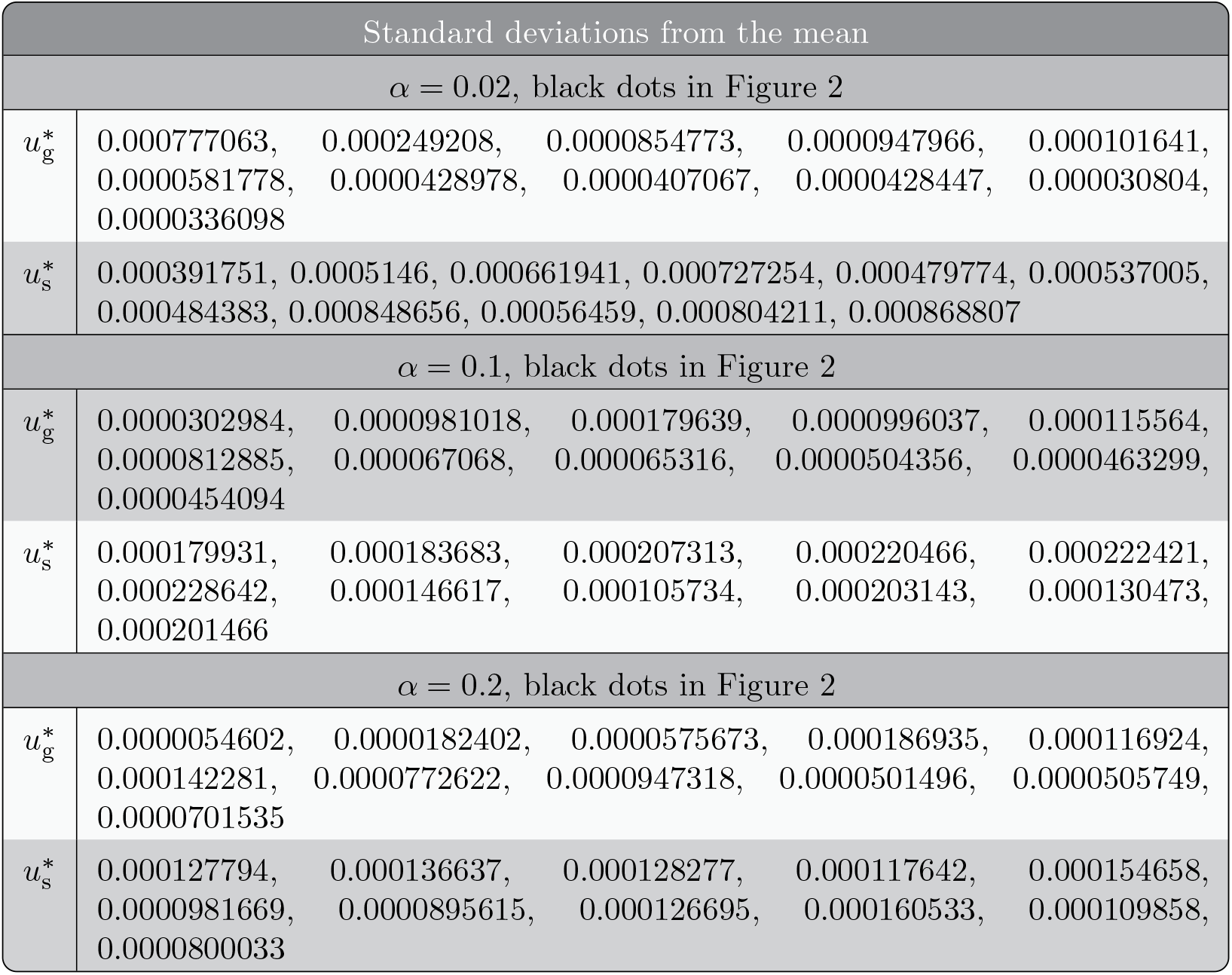
List of standard deviations from the mean (measured over 7500 generations), for the same parameter values as in Figure 2.

### Appendix C.2 Coevolution of age at maturity and germline maintenance

The simulation algorithm for this scenario (see section 2.3. of the S.I for the Mathematica code) follows a population whose size is endogenously determined according to a continuous-time stochastic updating process using the so-called “thinning” algorithm described in Section 3.1 of Ferriere and Tran (2009), which allows to exactly implement our life-cycle assumptions. A thinning algorithm is essentially an algorithm to simulate the points in an inhomogeneous Poisson process (inhomogeneous Poisson processes can be simulated by “thinning” the points from the homogeneous Poisson process), where the points or events take place sequentially (see e.g. Chen, 2016 for a conceptual description). Hence, under this algorithm, each individual is described by a vector specifying its age, allocation to repair, the age at maturity, and the number of deleterious mutations the individual has. The events in the thinning algorithm then follow a Poisson point process whose mean is determined by the vital rates (eq. 12) and where the occurrence of the events depends on the relative weights set by birth, death, and mutation rates of an individual. We defined as a “generation” *N* (**u**^∗^) iterations of the thinning algorithm, where *N* (**u**^∗^) is the analytical prediction of the carrying capacity of the model at the uninvadable trait value **u**^∗^. This is so because during one iteration of the thinning algorithm, a maximum of one event can occur (birth, death, or mutation of an individual) to one randomly chosen individual and so after having iterated the process *N* (**u**^∗^) times, on average the total population has been sampled. Thus, in order to produce a single data point in Fig. (4), we ran the six million (=*N* (**u**^∗^) × *N*_generations_ ≈ 2000 × 3000) iterations of the thinning algorithm. The mutation rate in the life-history locus is set to *μ*_*LH*_ = 0.1 and the effect size of the mutation follows a Normal distribution with zero mean and a standard deviation (=0.07 in our simulations). Simulating the results shown in Fig. (4), we initialised the simulation with a monomorphic population, where individual age is given by *a* = 1*/d*_b_ (recall, that *d*_b_ is the baseline mortality, with no deleterious mutations and life-history trait values given by the analytically predicted equilibrium. To obtain the results shown in Fig. 4, we initialised the simulation with a monomorphic population, with no deleterious mutations and life-history trait values given by the analytically predicted equilibrium. In Table C-5, we have depicted the time-averaged standard deviations from the main trait for parameter values in Fig. (4). In Fig. (5) we demonstrate the convergence stability of our simulations and we started the simulations away from equilibrium for four different initial values of the traits.

**Table C-5:**
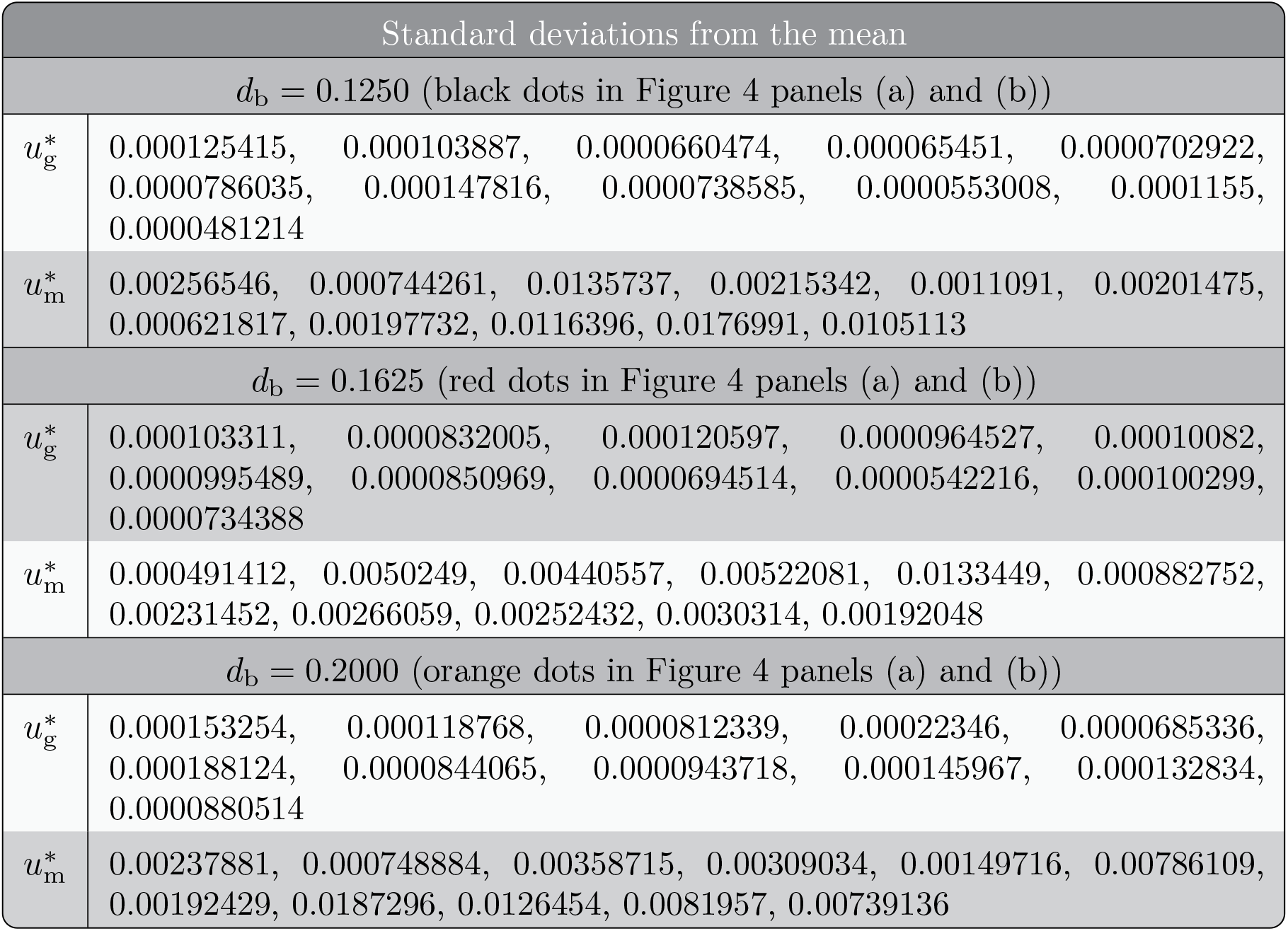
List of standard deviations from the mean (measured over 1500 “generations”), for the same parameter values as in Figure 4.

## Notes

**Conflict of Interest Statement:** All authors declare that they have no conflicts of interest.

### Competing Interest Statement

The authors have declared no competing interest.

### Summary of Updates

Minor revisions in writing.

