## Supplementary Information for "Life history and deleterious mutation rate coevolution"

### “Life-history and mutation rate joint evolution”

*Supplementary Mathematica file*

Piret Avila and Laurent Lehmann
Department of Ecology and Evolution
University of Lausanne

v. 26/07/2023

This notebook (written in Mathematica 12) contains the necessary documentation to derive analytically the results of the two examples and the code for the simulations of the paper “Life-history and mutation rate joint evolution” by Piret Avila and Laurent Lehmann.

**Table of contents**

1 .............**“Co-evolution of reproductive effort and mutation rate”**
1.1. ........Analysis of the model
1.2. ........Second-order condition for uninvadability and convergence stability
1.3. ........Individual-based simulation code
1.4. ........Simulation results
1.4b........Simulation results (with standard deviations)
1.5. ........Graphical illustrations (Fig. 2 of the main text)

2 .............**“Co-evolution of age-at-maturity and mutation rate”**
2.1. ........Analysis of the model
2.2. ........Second-order condition for uninvadability and convergence stability
2.3. ........Individual-based simulation code
2.4. ........Simulation results
2.4b.......Simulation results (with standard deviations)
2.5. ........Graphical illustrations (Fig. 3 and Fig. 4 of the main text)

In[ ]:=
```
Clear["Global`*"]
```

### 1. ”Co-evolution of reproductive effort and mutation rate”

```
In[ ]:= Clear["Global`*"]
```

### 1.1. Analysis of the model

**This color means that it is a key equation in the main text.**

**Important parameters and their notations in this Mathematica file (note that symbols slightly differ from the main text):**

us, ur - allocation to survival and repair (mutant) (u_s, u_g  in the main text)
vs, vr - allocation to survival and repair (resident) (v_s, v_g  in the main text)
$\mu$ - mutation rate during survival
a - age
f0, s0 - fecundity and survival of the least - loaded class
fp, sp - average fecundity and survival
b0, f0 - effective fecundity and fecundity of the least loaded class
fp - average fecundity
lt0, bt0 - survival with zero new mutations, effective fecundity with zero new mutations of the least-loaded class

#### 1.1.1. Basic reproductive number, vital rates and equilibrium distribution of individuals with different number of deleterious mutations

```
In[ ]:= Clear["Global`*"]
```

##### 1.1.1.1. Basic reproductive number

```
(*vital rates with no mutations of the least loaded class,
eqs. B-1 and B-2 in Appendix *)
```

```
In[ ]:= lt0[a_, us_, ur_] :=  (s0[us, ur])^(a-1) * e^(-a*μ[ur])
bt0[a, us, ur, vs, vr] := b0[us, ur, vs, vr]
```

```
(*Ro of the least least loaded, derivation of eq. B-4 of Appendix*)
```

$$\text{In[ ]:= R0[us\_, ur\_, vs\_, vr\_]} = \sum_{a=1}^{\infty} \text{bt0[a, us, ur, vs, vr] * lt0[a, us, ur] // Simplify}$$

$$\text{Out[ ]=} \frac{\text{b0[us, ur, vs, vr]}}{e^{\mu[\text{ur}]} - \text{s0[us, ur]}}$$

#### 1.1.1.2. Equilibrium distribution of individuals with different number of deleterious mutations

In[ ]:=
```
(*Here we aim to show that the distribution of individuals
 with diffrent number of deleterious mutations at equilibrium
 follows a Poisson distribution with mean given by mu/ss,
where ss is effect of a seleterious mutation on fecundity and survival *)
```

In[ ]:=
```
f[j_] := fo (1 - ss)^j (* fecundity of indidivudal with j mutations,
where fo → fecundity of indidivual with 0 mutations*)
s[j_] := so (1 - ss)^j (* survival of an indidivudal with j mutations,
where so → survival of an indidivual with 0 mutations*)
w[j_] := s[j] + (1 - sa) f[j]/fa
(* indidivual fitness of an indidivual with j mutations,
here  sa is average survival and fa is average fecundity *)
ϕ[j_] := PDF[PoissonDistribution[μ], j]
 (*probability that j mutations are produced upon reproduction*)


(* Here we check that if ϕ[j] is Poisson distributed with mean  μ,
then it follows that p[j] is  is Poisson distributed with mean  μ/ss and sa==
 ⅇ^-μ so,fa==ⅇ^-μ fo. We first do this by calculating average survival
   and fecundity assuming that p(j) is Poisson distributed and
   then show that Poisson distributed p(j) satisfies eq. B-5 *)


(* Lets first assume that p[j] is  is Poisson distributed
  with mean  μ/ss and then see that it satisfies eq. B-5 *)
```

In[ ]:=
```
p[j_] := PDF[PoissonDistribution[μ/ss], j]

(* Lets first assume that p[j] is  is Poisson distributed with mean
  μ/ss and then see that it satisfies eq. B-5 of the main text *)


(* Mean vital rates when  p[j] is  is Poisson distributed,
gives Eq. B-6 of the main text *)
```

In[ ]:=

$$\left\{sa == \sum_{j=0}^{\infty} s[j]\times p[j],\ fa == \sum_{j=0}^{\infty} f[j]\times p[j]\right\} \text{ // FullSimplify}$$

Out[ ]= $\{sa == e^{-\mu}\, so,\ fa == e^{-\mu}\, fo\}$

```
(* We now check that Poisson distribution p[j] satisfies Eq. B-
 5 of the main text *)
```

In[ ]:=
```
Assuming[k ≥ 0, Simplify[p[k]]] ==
  Evaluate[Sum[w[j] ϕ[k - j] p[j], {j, 0, k}] /. {fa → ⅇ^-μ fo, sa → ⅇ^-μ so} // FullSimplify]
```

Out[ ]= True

##### 1.1.1.3. Definition of vital rates and checking that R0 = 1 at equilibrium

```
(*effective fecundity of the least loaded class, eq. B-2*)
```

In[ ]:=
```
b0[us, ur, vs, vr] := (1 - sp[vs, vr]) * f0[us, ur] / fp[vs, vr]
```

```
(* fo→f0,
so→s0 survival and fecundity and mutation rate of the leas loaded class,
eq. 7*)
```

In[ ]:=
```
f0[us_, ur_] := fb * ((1 - us) * (1 - ur))^αf
s0[us_, ur_] := sb * (us * (1 - ur))^αs
(* mutation rate during survival, last equation of eq. 6*)
μ[ur_] := μb * (1 - ur)^αμ
```

```
(* average population survival and fecundity (fa→ fp, sa→sp),
eq. B-6, here we just define them, proof is in the previous section *)
```

In[ ]:=
```
sp[vs_, vr_] := s0[vs, vr] * ⅇ^(-μ[vr])
fp[vs_, vr_] := f0[vs, vr] * ⅇ^(-μ[vr])
```

In[ ]:=
```
(*Checking if R0=1 in resident population*)
```

In[ ]:=
```
FullSimplify[R0[us, ur, vs, vr] /. {us → vs, ur → vr}]
```

Out[ ]= R0[vs, vr, vs, vr]

#### 1.1.2. Selection gradients

In[ ]:=
```
(*Selection gradient on repair*)

Sur[vs, vr] = FullSimplify[
   PowerExpand[Evaluate[D[R0[us, ur, vs, vr], ur] /. {us → vs, ur → vr}]]];

(*Derivation of eq. 8 *)
```

In[ ]:=
```
FullSimplify[PowerExpand[1/((1 - vr) (1 - sp[vs, vr])) (μ[vr] α_μ - (α_s * sp[vs, vr] + α_f * (1 - sp[vs, vr])))] ==
  PowerExpand[Sur[vs, vr]]]
```

Out[ ]= True

In[ ]:= `(*Selection gradient on survival*)`

In[ ]:=
```
Sus[vs, vr] = FullSimplify[
  PowerExpand[Evaluate[D[R0[us, ur, vs, vr], us] /. {us → vs, ur → vr}]]]
```

Out[ ]= $\frac{\alpha_f}{-1+vs}+\frac{sb\,(1-vr)^{\alpha_s}\,vs^{-1+\alpha_s}\,\alpha_s}{e^{(1-vr)^{\alpha_\mu}\,\mu b}-sb\,(1-vr)^{\alpha_s}\,vs^{\alpha_s}}$

`(*Derivation of eq. 9 *)`

In[ ]:=
```
FullSimplify[
  PowerExpand[1/(1 - sp[vs, vr]) ((α_s * sp[vs, vr])/vs - (α_f * (1 - sp[vs, vr]))/(1 - vs))] ==
  PowerExpand[Sus[vs, vr]]]
```

Out[ ]= True

##### 1.1.3. Singular strategies

In[ ]:= `FullSimplify[Sur[vs, vr] /. {α_f → α, α_s → α }];`

In[ ]:= `Solve[% == 0, vr] // FullSimplify`

Solve : Inverse functions are being used by Solve, so some solutions may not be found; use Reduce for complete solution information.

Out[ ]= $\left\{\left\{vr\to 1-\left(\frac{\alpha}{\mu b\,\alpha_\mu}\right)^{\frac{1}{\alpha_\mu}}\right\}\right\}$

```
(* Eq. (10), if μb>α/α_μ ,
otherwise vr=0 (since vr can not be smaller than 0) *)
```

In[ ]:= `FullSimplify[Sus[vs, vr] /. {α_f → α, α_s → α }] /. {vr → 1 - (α/(μb α_μ))^(1/α_μ)} // FullSimplify;`

In[•]:=
```
Solve[% == 0, vs] // FullSimplify
```

Solve : Inverse functions are being used by Solve, so some solutions may not be found; use Reduce for complete solution information.

Out[•]= $\left\{\left\{vs \to \left(e^{-\mu b \left(\left(\frac{\alpha}{\mu b\,\alpha_\mu}\right)^{\frac{1}{\alpha_\mu}}\right)^{\alpha_\mu}}\ sb \left(\left(\frac{\alpha}{\mu b\,\alpha_\mu}\right)^{\frac{1}{\alpha_\mu}}\right)^{\alpha}\right)^{\frac{1}{1-\alpha}}\right\}\right\}$

In[•]:=
```
(* Second equation of eq. (20), when  vr>0 *)
```

In[•]:=
```
FullSimplify[
```
$\text{PowerExpand}\left[\left(\frac{e^{\left(\frac{\alpha}{\alpha_\mu}\right)}\left(\frac{\alpha}{\mu b\,\alpha_\mu}\right)^{-\frac{\alpha}{\alpha_\mu}}}{sb}\right)^{\frac{1}{\alpha-1}}\right] == \text{PowerExpand}\left[\left(e^{-\mu b \left(\left(\frac{\alpha}{\mu b\,\alpha_\mu}\right)^{\frac{1}{\alpha_\mu}}\right)^{\alpha_\mu}}\ sb \left(\left(\frac{\alpha}{\mu b\,\alpha_\mu}\right)^{\frac{1}{\alpha_\mu}}\right)^{\alpha}\right)^{\frac{1}{1-\alpha}}\right]]$

Out[•]= True

This holds only if $\mu b > \alpha / \alpha_\mu$ and $vr > 0$

In[•]:=

In[•]:=
```
FullSimplify[Sus[vs, vr] /. {αf → α, αs → α }] /. {vr → 0} // FullSimplify;
```

In[•]:=
```
Solve[% == 0, vs] // FullSimplify
```

Solve : Inverse functions are being used by Solve, so some solutions may not be found; use Reduce for complete solution information.

Out[•]= $\left\{\left\{vs \to \left(\frac{e^{\mu b}}{sb}\right)^{\frac{1}{-1+\alpha}}\right\}\right\}$

In[•]:=
```
(* Second equation of eq. (20), when  vr=0 *)
```

This holds if $\mu b \leq \alpha / \alpha_\mu$ and $vr = 0$

In[•]:=

In[•]:=
```
(*Allocation between survival and reproduction according to classical life-
 history prediction (as vr→0,μb→0) *)
```

In[•]:=
```
FullSimplify[Sus[vs, vr] /. {αf → α, αs → α }] /. {vr → 0, μb → 0} // FullSimplify;
```

In[ ]:= `Solve[% == 0, vs] // FullSimplify`

Solve : Inverse functions are being used by Solve, so some solutions may not be found; use Reduce for complete solution information.

Out[ ]= $\left\{\left\{vs \to \left(\frac{1}{sb}\right)^{\frac{1}{-1+\alpha}}\right\}\right\}$

#### 1.1.4. Additional (numerical) analysis when $\alpha_f \neq \alpha_s$

##### 1.1.4.1. Numerical results $\alpha_f \neq \alpha_s$ and as a function of $\mu b$

In[ ]:= $Sur[vs\_, vr\_] = \frac{\alpha_f - \alpha_s + \frac{e^{(1-vr)^{\alpha_\mu}\mu b}\left(\alpha_s - (1-vr)^{\alpha_\mu}\mu b\,\alpha_\mu\right)}{e^{(1-vr)^{\alpha_\mu}\mu b} - sb\,(1-vr)^{\alpha_s}\,vs^{\alpha_s}}}{-1+vr};$

In[ ]:= $Sus[vs\_, vr\_] = \frac{\alpha_f}{-1+vs} + \frac{sb\,(1-vr)^{\alpha_s}\,vs^{-1+\alpha_s}\,\alpha_s}{e^{(1-vr)^{\alpha_\mu}\mu b} - sb\,(1-vr)^{\alpha_s}\,vs^{\alpha_s}};$

In[ ]:= $Data = \{\alpha_f \to 1/4, \alpha_s \to 1/2, \alpha_\mu \to 2, fb \to 5, sb \to 1/2\};$

In[ ]:= `SurData[vs_, vr_, μb_] = FullSimplify[Sur[vs, vr] /. Data]`

Out[ ]= $\frac{-\frac{1}{4} + \frac{e^{(-1+vr)^2\mu b}\left(1-4\,(-1+vr)^2\mu b\right)}{2\,e^{(-1+vr)^2\mu b} - \sqrt{1-vr}\,\sqrt{vs}}}{-1+vr}$

In[ ]:= `SusData[vs_, vr_, μb_] = FullSimplify[Sus[vs, vr] /. Data]`

Out[ ]= $\frac{1}{4}\left(\frac{1}{-1+vs} + \frac{2-2\,vr}{2\,e^{(-1+vr)^2\mu b}\sqrt{1-vr}\,\sqrt{vs} + (-1+vr)\,vs}\right)$

In[ ]:= `FindRoot[{SurData[vs, vr, 0.2], SusData[vs, vr, 0.2]}, {{vs, 0.5}, {vr, 0.5}}]`

Out[ ]= {vs → 0.409445, vr → 0.113496}

In[ ]:=
```
vSin[μb_] := vSin[μb] = {vs, vr} /.
    FindRoot[{SurData[vs, vr, μb], SusData[vs, vr, μb]}, {{vs, 0.5}, {vr, 0.5}}]
```

In[ ]:=

```
Plot[
 {vSin[μb][[1]], Clip[vSin[μb][[2]], {0, 1}], μb * (1 - Clip[vSin[μb][[2]], {0, 1}])^2},
 {μb, 0.02, 2}, PlotStyle → {Red, Green, Black}]
```

Out[ ]=

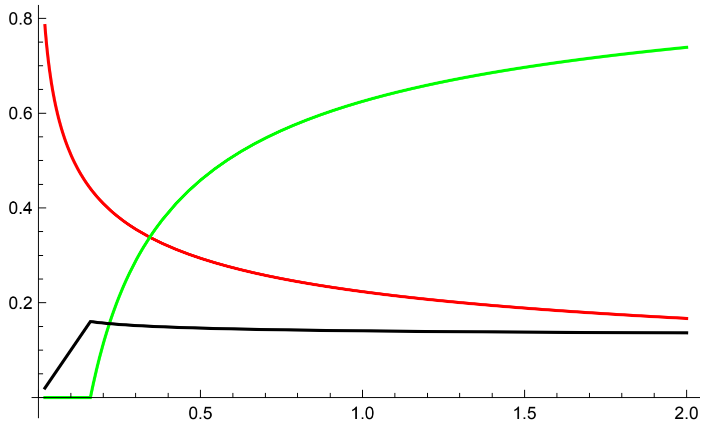


In[ ]:=

```
Data = {αf → 0.9, αs → 0.001, αμ → 2, fb → 5, sb → 0.8};
```

In[ ]:=

```
SurData[vs_, vr_, μb_] = FullSimplify[Sur[vs, vr] /. Data]
```

Out[ ]=

$$\frac{0.899 - \frac{2.\, e^{(-1+vr)^2 \mu b} (-0.0005+\mu b+(-2.+vr)\, vr\, \mu b)}{e^{(-1+vr)^2 \mu b}-0.8\,(1-vr)^{0.001}\, vs^{0.001}}}{-1+vr}$$

In[ ]:=

```
SusData[vs_, vr_, μb_] = FullSimplify[Sus[vs, vr] /. Data]
```

Out[ ]=

$$\frac{0.9}{-1.+vs} + \frac{0.0008\,(1-vr)^{0.001}}{\left(e^{(-1+vr)^2 \mu b} - 0.8\,(1-vr)^{0.001}\, vs^{0.001}\right) vs^{0.999}}$$

In[ ]:=

```
FindRoot[{SurData[vs, vr, 0.2], SusData[vs, vr, 0.2]}, {{vs, 0.5}, {vr, 0.5}}]
```

Out[ ]=

```
{vs → 0.00248903, vr → 0.166363}
```

In[ ]:=

```
vSin[μb_] := vSin[μb] = {vs, vr} /.
   FindRoot[{SurData[vs, vr, μb], SusData[vs, vr, μb]}, {{vs, 0.5}, {vr, 0.5}}]
```

In[ ]:=

```
Plot[
 {vSin[μb][[1]], Clip[vSin[μb][[2]], {0, 1}], μb * (1 - Clip[vSin[μb][[2]], {0, 1}])^2},
 {μb, 0.02, 2}, PlotStyle → {Red, Green, Black}]
```

Out[ ]=

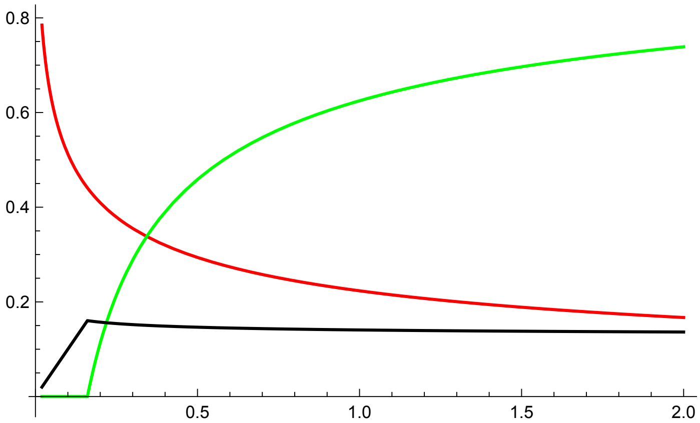


Here the green line is the singular allocation to repair ( $u_r^*$ ), the red line is singular allocation to survival ( $u_s^*$ ), and the black line is the singular mutation rate $\mu$ ( $u_r^*$ ). We see that allocation to repair is an increasing function of $\mu_b$ and survival is a decreasing function of $\mu_b$. Hence, our results are robust in terms of the assumption $\alpha_f \neq \alpha_s$.

##### 1.1.4.2. Some conclusion of graphical analysis when $\alpha_f \neq \alpha_s$

```
(* In this section we plot selection gradients, as a function
 of traits. The singular strategies are determined by the points,
whre the line crosses the x axis; We also plot cost and
 benefit terms to see how they depend on parameter values. *)
```

```
(*Selection gradient on survival, variying vr  *)
```

$$\text{Plot}\Big[\Big\{\left(\frac{\alpha_s * \text{sp}[\text{vs}, \text{vr}]}{\text{vs}} - \frac{\alpha_f * (1 - \text{sp}[\text{vs}, \text{vr}])}{(1 - \text{vs})}\right) \text{/.}$$

$\{\text{vr} \to 0.1, \alpha_f \to 0.2, \alpha_s \to 0.5, \alpha_\mu \to 2, \text{fb} \to 5, \text{sb} \to 0.5, \mu\text{b} \to 2\},$

$$\left(\frac{\alpha_s * \text{sp}[\text{vs}, \text{vr}]}{\text{vs}} - \frac{\alpha_f * (1 - \text{sp}[\text{vs}, \text{vr}])}{(1 - \text{vs})}\right) \text{/.}$$

$\{\text{vr} \to 0.5, \alpha_f \to 0.2, \alpha_s \to 0.5, \alpha_\mu \to 2, \text{fb} \to 5, \text{sb} \to 0.5, \mu\text{b} \to 2\},$

$$\left(\frac{\alpha_s * \text{sp}[\text{vs}, \text{vr}]}{\text{vs}} - \frac{\alpha_f * (1 - \text{sp}[\text{vs}, \text{vr}])}{(1 - \text{vs})}\right) \text{/.}$$

$\{\text{vr} \to 0.9, \alpha_f \to 0.2, \alpha_s \to 0.5, \alpha_\mu \to 2, \text{fb} \to 5, \text{sb} \to 0.5, \mu\text{b} \to 2\}\Big\},$

```
{vs, 0, 0.5}, AxesLabel → {"vs", "Selection gradient on survival"}]
```

Out[ ]=

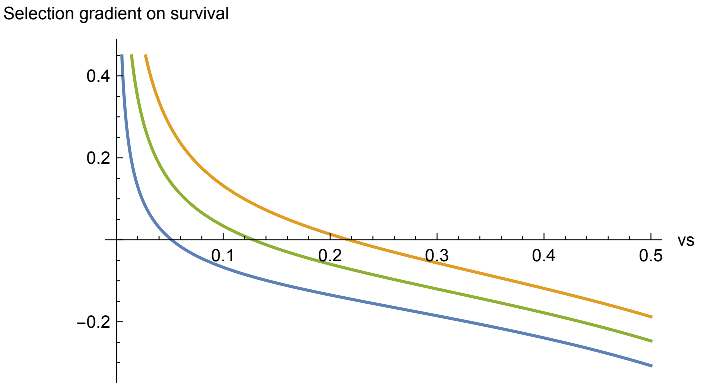


(*We see that survival is higher, with higher repair,
even when $\alpha_f \neq \alpha_s$, we can try out different combinations *)

```
(*Cost of survival, variying repair  *)

Plot[{(αf * (1 - sp[vs, vr]))/(1 - vs) /. {vr → 0.1, αf → 0.2,
    αs → 0.5, αμ → 2, fb → 5, sb → 0.5, μb → 2}, (αf * (1 - sp[vs, vr]))/(1 - vs) /.
   {vr → 0.5, αf → 0.2, αs → 0.5, αμ → 2, fb → 5, sb → 0.5, μb → 2},
  (αf * (1 - sp[vs, vr]))/(1 - vs) /.
   {vr → 0.9, αf → 0.2, αs → 0.5, αμ → 2, fb → 5, sb → 0.5, μb → 2}},
 {vs, 0, 0.5}, AxesLabel → {"vs", "Cost of survival"}]
```

Out[∘]=

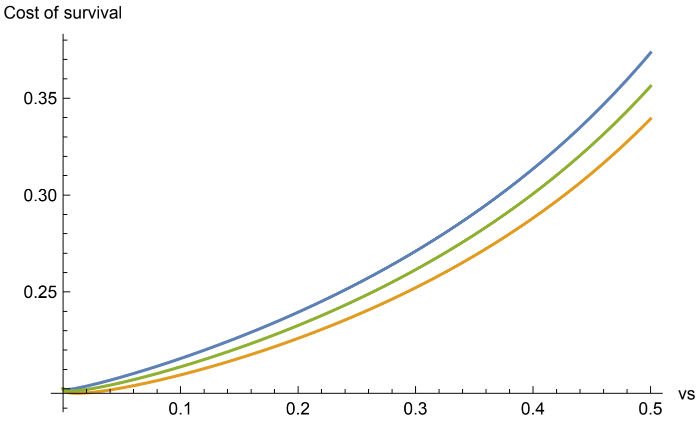


```
(*Cost decreases with repair*)
```

```
(*Benefit of survival, variying αs  *)

Plot[
 {(αs * sp[vs, vr] / vs) /. {vr → 0.1, αf → 0.2, αs → 0.5, αμ → 2, fb → 5, sb → 0.5, μb → 2},
  (αs * sp[vs, vr] / vs) /. {vr → 0.5, αf → 0.2, αs → 0.5, αμ → 2, fb → 5, sb → 0.5, μb → 2},
  (αs * sp[vs, vr] / vs) /.
   {vr → 0.9, αf → 0.2, αs → 0.5, αμ → 2, fb → 5, sb → 0.5, μb → 2}},
 {vs, 0, 0.5}, AxesLabel → {"vs", "Benefit of survival"}]
```

Out[∘]=

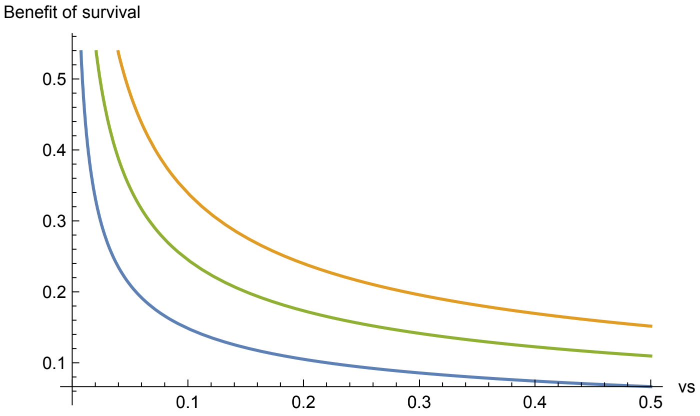


```
(*Benefit increases with repair*)


(*Selection gradient on survival, variying vr  *)
```

In[ ]:=

```
Plot[{(αs * sp[vs, vr]/vs - αf * (1 - sp[vs, vr])/(1 - vs)) /.
    {vr → 0.5, αf → 0.5, αs → 0.01, αμ → 2, fb → 5, sb → 0.5, μb → 2},
   (αs * sp[vs, vr]/vs - αf * (1 - sp[vs, vr])/(1 - vs)) /.
    {vr → 0.5, αf → 0.5, αs → 0.02, αμ → 2, fb → 5, sb → 0.5, μb → 2},
   (αs * sp[vs, vr]/vs - αf * (1 - sp[vs, vr])/(1 - vs)) /.
    {vr → 0.5, αf → 0.5, αs → 0.6, αμ → 2, fb → 5, sb → 0.5, μb → 2},
   (αs * sp[vs, vr]/vs - αf * (1 - sp[vs, vr])/(1 - vs)) /.
    {vr → 0.5, αf → 0.5, αs → 0.9, αμ → 2, fb → 5, sb → 0.5, μb → 2}},
  {vs, 0, 0.5}, AxesLabel → {"vs", "Selection gradient on survival"}]
```

Out[ ]=

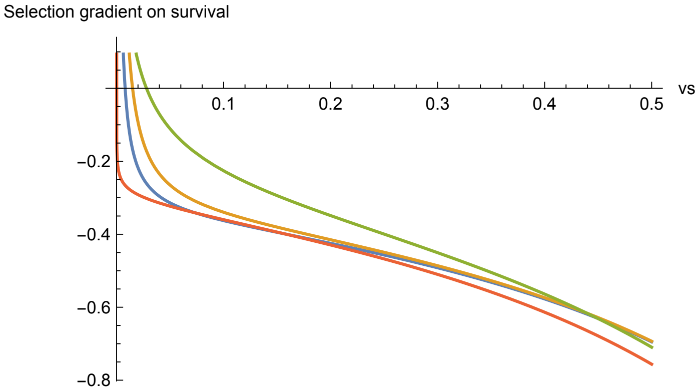


```
(* We see that for higher αs, investment into survival increases
 initially and then after a certain value it decreases again *)

(*Selection gradient on survival, variying αf  *)
```

In[ ]:=

```
Plot[{(αs * sp[vs, vr]/vs - αf * (1 - sp[vs, vr])/(1 - vs)) /.
    {vr → 0.5, αf → 0.1, αs → 0.5, αμ → 2, fb → 5, sb → 0.5, μb → 2},
   (αs * sp[vs, vr]/vs - αf * (1 - sp[vs, vr])/(1 - vs)) /.
    {vr → 0.5, αf → 0.5, αs → 0.5, αμ → 2, fb → 5, sb → 0.5, μb → 2},
   (αs * sp[vs, vr]/vs - αf * (1 - sp[vs, vr])/(1 - vs)) /.
    {vr → 0.5, αf → 0.9, αs → 0.5, αμ → 2, fb → 5, sb → 0.5, μb → 2}},
  {vs, 0, 0.5}, AxesLabel → {"vs", "Selection gradient on survival"}]
```

Out[ ]=

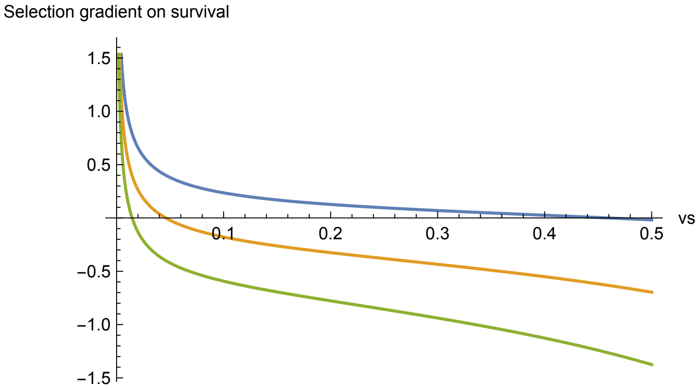


```
(* We see that for higher αf, investment into survival goes down *)
```

#### 1.2. Second-order condition for uninvadability and convergence stability

##### 1.2.1. Uninvadability condition (compute the Hessian)

In[ ]:=

$$\text{vrSol} = 1 - \left(\frac{\alpha}{\mu b\, \alpha_\mu}\right)^{\frac{1}{\alpha_\mu}};$$

$$\text{vsSol} = \left(e^{-\mu b \left(\left(\frac{\alpha}{\mu b\, \alpha_\mu}\right)^{\frac{1}{\alpha_\mu}}\right)^{\alpha_\mu}}\ \text{sb} \left(\left(\frac{\alpha}{\mu b\, \alpha_\mu}\right)^{\frac{1}{\alpha_\mu}}\right)^{\alpha}\right)^{\frac{1}{1-\alpha}};$$

In[ ]:=

```
Hu = FullSimplify[
  FullSimplify[Assuming[μb > 0 && 0 < ur < 1 && sb > 0 && α > 0 && αμ > 0 && αs > 0 && αf > 0 ,
     Refine[D[R0[us, ur, vs, vr], {{us, ur}, 2}] // FullSimplify]] /.
    {us → vs, ur → vr, αf → α, αs → α }] /. {vs → vsSol, vr → vrSol}]
```

Out[ ]=

$$\Bigg\{\Bigg\{R0^{(2,0,0,0)}\Bigg[\left(e^{-\mu b\left(\left(\frac{\alpha}{\mu b\,\alpha_\mu}\right)^{\frac{1}{\alpha_\mu}}\right)^{\alpha_\mu}} sb\left(\left(\left(\frac{\alpha}{\mu b\,\alpha_\mu}\right)^{\frac{1}{\alpha_\mu}}\right)^{\alpha}\right)\right)^{\frac{1}{1-\alpha}},$$

$$1-\left(\frac{\alpha}{\mu b\,\alpha_\mu}\right)^{\frac{1}{\alpha_\mu}},\ \left(e^{-\mu b\left(\left(\frac{\alpha}{\mu b\,\alpha_\mu}\right)^{\frac{1}{\alpha_\mu}}\right)^{\alpha_\mu}} sb\left(\left(\left(\frac{\alpha}{\mu b\,\alpha_\mu}\right)^{\frac{1}{\alpha_\mu}}\right)^{\alpha}\right)\right)^{\frac{1}{1-\alpha}},\ 1-\left(\frac{\alpha}{\mu b\,\alpha_\mu}\right)^{\frac{1}{\alpha_\mu}}\Bigg],$$

$$R0^{(1,1,0,0)}\Bigg[\left(e^{-\mu b\left(\left(\frac{\alpha}{\mu b\,\alpha_\mu}\right)^{\frac{1}{\alpha_\mu}}\right)^{\alpha_\mu}} sb\left(\left(\left(\frac{\alpha}{\mu b\,\alpha_\mu}\right)^{\frac{1}{\alpha_\mu}}\right)^{\alpha}\right)\right)^{\frac{1}{1-\alpha}},\ 1-\left(\frac{\alpha}{\mu b\,\alpha_\mu}\right)^{\frac{1}{\alpha_\mu}},$$

$$\left(e^{-\mu b\left(\left(\frac{\alpha}{\mu b\,\alpha_\mu}\right)^{\frac{1}{\alpha_\mu}}\right)^{\alpha_\mu}} sb\left(\left(\left(\frac{\alpha}{\mu b\,\alpha_\mu}\right)^{\frac{1}{\alpha_\mu}}\right)^{\alpha}\right)\right)^{\frac{1}{1-\alpha}},\ 1-\left(\frac{\alpha}{\mu b\,\alpha_\mu}\right)^{\frac{1}{\alpha_\mu}}\Bigg]\Bigg\},$$

$$\Bigg\{R0^{(1,1,0,0)}\Bigg[\left(e^{-\mu b\left(\left(\frac{\alpha}{\mu b\,\alpha_\mu}\right)^{\frac{1}{\alpha_\mu}}\right)^{\alpha_\mu}} sb\left(\left(\left(\frac{\alpha}{\mu b\,\alpha_\mu}\right)^{\frac{1}{\alpha_\mu}}\right)^{\alpha}\right)\right)^{\frac{1}{1-\alpha}},\ 1-\left(\frac{\alpha}{\mu b\,\alpha_\mu}\right)^{\frac{1}{\alpha_\mu}},$$

$$\left(e^{-\mu b\left(\left(\frac{\alpha}{\mu b\,\alpha_\mu}\right)^{\frac{1}{\alpha_\mu}}\right)^{\alpha_\mu}} sb\left(\left(\left(\frac{\alpha}{\mu b\,\alpha_\mu}\right)^{\frac{1}{\alpha_\mu}}\right)^{\alpha}\right)\right)^{\frac{1}{1-\alpha}},\ 1-\left(\frac{\alpha}{\mu b\,\alpha_\mu}\right)^{\frac{1}{\alpha_\mu}}\Bigg],$$

$$R0^{(0,2,0,0)}\Bigg[\left(e^{-\mu b\left(\left(\frac{\alpha}{\mu b\,\alpha_\mu}\right)^{\frac{1}{\alpha_\mu}}\right)^{\alpha_\mu}} sb\left(\left(\left(\frac{\alpha}{\mu b\,\alpha_\mu}\right)^{\frac{1}{\alpha_\mu}}\right)^{\alpha}\right)\right)^{\frac{1}{1-\alpha}},\ 1-\left(\frac{\alpha}{\mu b\,\alpha_\mu}\right)^{\frac{1}{\alpha_\mu}},$$

$$\left(e^{-\mu b\left(\left(\frac{\alpha}{\mu b\,\alpha_\mu}\right)^{\frac{1}{\alpha_\mu}}\right)^{\alpha_\mu}} sb\left(\left(\left(\frac{\alpha}{\mu b\,\alpha_\mu}\right)^{\frac{1}{\alpha_\mu}}\right)^{\alpha}\right)\right)^{\frac{1}{1-\alpha}},\ 1-\left(\frac{\alpha}{\mu b\,\alpha_\mu}\right)^{\frac{1}{\alpha_\mu}}\Bigg]\Bigg\}\Bigg\}$$

#### 1.2.2. Initialize the Hessian for checking uninvadability

In[ ]:=

```
(* Hessian of the evolutionary system,
we copied Hu from previous section as the computation takes a long time *)
```

In[ ]:=

$$\text{HessianEx1}[\mu b\_] := \Bigg\{\Bigg\{\left(\alpha\left(1-\left(e^{-\mu b\left(\left(\frac{\alpha}{\mu b\,\alpha_\mu}\right)^{\frac{1}{\alpha_\mu}}\right)^{\alpha_\mu}} sb\left(\left(\left(\frac{\alpha}{\mu b\,\alpha_\mu}\right)^{\frac{1}{\alpha_\mu}}\right)^{\alpha}\right)\right)^{\frac{1}{1-\alpha}}\right)\right)^{-2+\alpha}$$

$$\Bigg(e^{2\mu b\left(\left(\frac{\alpha}{\mu b\,\alpha_{\mu}}\right)^{\frac{1}{\alpha_{\mu}}}\right)^{\alpha_{\mu}}}(-1+\alpha)\left(e^{-\mu b\left(\left(\frac{\alpha}{\mu b\,\alpha_{\mu}}\right)^{\frac{1}{\alpha_{\mu}}}\right)^{\alpha_{\mu}}}\,\text{sb}\left(\left(\frac{\alpha}{\mu b\,\alpha_{\mu}}\right)^{\frac{1}{\alpha_{\mu}}}\right)^{\alpha}\right)^{-\frac{2}{-1+\alpha}}+$$

$$e^{\mu b\left(\left(\frac{\alpha}{\mu b\,\alpha_{\mu}}\right)^{\frac{1}{\alpha_{\mu}}}\right)^{\alpha_{\mu}}}\,\text{sb}\Bigg(-1+\alpha-2\,(-1+2\,\alpha)\left(e^{-\mu b\left(\left(\frac{\alpha}{\mu b\,\alpha_{\mu}}\right)^{\frac{1}{\alpha_{\mu}}}\right)^{\alpha_{\mu}}}\,\text{sb}\left(\left(\frac{\alpha}{\mu b\,\alpha_{\mu}}\right)^{\frac{1}{\alpha_{\mu}}}\right)^{\alpha}\right)^{\frac{1}{1-\alpha}}+$$

$$(1+\alpha)\left(e^{-\mu b\left(\left(\frac{\alpha}{\mu b\,\alpha_{\mu}}\right)^{\frac{1}{\alpha_{\mu}}}\right)^{\alpha_{\mu}}}\,\text{sb}\left(\left(\frac{\alpha}{\mu b\,\alpha_{\mu}}\right)^{\frac{1}{\alpha_{\mu}}}\right)^{\alpha}\right)^{-\frac{2}{-1+\alpha}}\Bigg)$$

$$\left(\left(e^{-\mu b\left(\left(\frac{\alpha}{\mu b\,\alpha_{\mu}}\right)^{\frac{1}{\alpha_{\mu}}}\right)^{\alpha_{\mu}}}\,\text{sb}\left(\left(\frac{\alpha}{\mu b\,\alpha_{\mu}}\right)^{\frac{1}{\alpha_{\mu}}}\right)^{\alpha}\right)^{\frac{1}{1-\alpha}}\left(\frac{\alpha}{\mu b\,\alpha_{\mu}}\right)^{\frac{1}{\alpha_{\mu}}}\right)^{\alpha}-$$

$$\text{sb}^{2}\left(-1-\alpha+2\left(e^{-\mu b\left(\left(\frac{\alpha}{\mu b\,\alpha_{\mu}}\right)^{\frac{1}{\alpha_{\mu}}}\right)^{\alpha_{\mu}}}\,\text{sb}\left(\left(\frac{\alpha}{\mu b\,\alpha_{\mu}}\right)^{\frac{1}{\alpha_{\mu}}}\right)^{\alpha}\right)^{\frac{1}{1-\alpha}}\right)$$

$$\left(\left(e^{-\mu b\left(\left(\frac{\alpha}{\mu b\,\alpha_{\mu}}\right)^{\frac{1}{\alpha_{\mu}}}\right)^{\alpha_{\mu}}}\,\text{sb}\left(\left(\frac{\alpha}{\mu b\,\alpha_{\mu}}\right)^{\frac{1}{\alpha_{\mu}}}\right)^{\alpha}\right)^{\frac{1}{1-\alpha}}\left(\frac{\alpha}{\mu b\,\alpha_{\mu}}\right)^{\frac{1}{\alpha_{\mu}}}\right)^{2\,\alpha}\Bigg)$$

$$\left(e^{-\mu b\left(\left(\frac{\alpha}{\mu b\,\alpha_{\mu}}\right)^{\frac{1}{\alpha_{\mu}}}\right)^{\alpha_{\mu}}}\,\text{sb}\left(\left(\frac{\alpha}{\mu b\,\alpha_{\mu}}\right)^{\frac{1}{\alpha_{\mu}}}\right)^{\alpha}\right)^{\frac{2}{-1+\alpha}}\left(\left(\frac{\alpha}{\mu b\,\alpha_{\mu}}\right)^{\frac{1}{\alpha_{\mu}}}\right)^{\alpha}$$

$$\left(-\left(\left(-1+\left(e^{-\mu b\left(\left(\frac{\alpha}{\mu b\,\alpha_{\mu}}\right)^{\frac{1}{\alpha_{\mu}}}\right)^{\alpha_{\mu}}}\,\text{sb}\left(\left(\frac{\alpha}{\mu b\,\alpha_{\mu}}\right)^{\frac{1}{\alpha_{\mu}}}\right)^{\alpha}\right)^{\frac{1}{1-\alpha}}\right)\left(\frac{\alpha}{\mu b\,\alpha_{\mu}}\right)^{\frac{1}{\alpha_{\mu}}}\right)^{-\alpha}\right)\Bigg/$$

$$\left(e^{\mu b\left(\left(\frac{\alpha}{\mu b\,\alpha_{\mu}}\right)^{\frac{1}{\alpha_{\mu}}}\right)^{\alpha_{\mu}}}-\text{sb}\left(\left(e^{-\mu b\left(\left(\frac{\alpha}{\mu b\,\alpha_{\mu}}\right)^{\frac{1}{\alpha_{\mu}}}\right)^{\alpha_{\mu}}}\,\text{sb}\left(\left(\frac{\alpha}{\mu b\,\alpha_{\mu}}\right)^{\frac{1}{\alpha_{\mu}}}\right)^{\alpha}\right)^{\frac{1}{1-\alpha}}\left(\frac{\alpha}{\mu b\,\alpha_{\mu}}\right)^{\frac{1}{\alpha_{\mu}}}\right)^{\alpha}\right)^{2},$$

$$-\Bigg(\Bigg(\text{sb}^{2}\,\alpha\left(1-\left(e^{-\mu b\left(\left(\frac{\alpha}{\mu b\,\alpha_{\mu}}\right)^{\frac{1}{\alpha_{\mu}}}\right)^{\alpha_{\mu}}}\,\text{sb}\left(\left(\frac{\alpha}{\mu b\,\alpha_{\mu}}\right)^{\frac{1}{\alpha_{\mu}}}\right)^{\alpha}\right)^{\frac{1}{1-\alpha}}\right)^{\alpha}\left(e^{-\mu b\left(\left(\frac{\alpha}{\mu b\,\alpha_{\mu}}\right)^{\frac{1}{\alpha_{\mu}}}\right)^{\alpha_{\mu}}}\,\text{sb}\left(\left(\frac{\alpha}{\mu b\,\alpha_{\mu}}\right)^{\frac{1}{\alpha_{\mu}}}\right)^{\alpha}\right)^{-1+\frac{1}{-1+\alpha}}$$

$$\left(\left(\frac{\alpha}{\mu b\,\alpha_{\mu}}\right)^{\frac{1}{\alpha_{\mu}}}\right)^{2\,\alpha}\left(-\left(\left(-1+\left(e^{-\mu b\left(\left(\frac{\alpha}{\mu b\,\alpha_{\mu}}\right)^{\frac{1}{\alpha_{\mu}}}\right)^{\alpha_{\mu}}}\,\text{sb}\left(\left(\frac{\alpha}{\mu b\,\alpha_{\mu}}\right)^{\frac{1}{\alpha_{\mu}}}\right)^{\alpha}\right)^{\frac{1}{1-\alpha}}\right)\left(\frac{\alpha}{\mu b\,\alpha_{\mu}}\right)^{\frac{1}{\alpha_{\mu}}}\right)\right)^{-1-\alpha}$$

$$\left(\left(\left(e^{-\mu b\left(\left(\frac{\alpha}{\mu b\,\alpha_\mu}\right)^{\frac{1}{\alpha_\mu}}\right)^{\alpha_\mu}}\,sb\left(\left(\frac{\alpha}{\mu b\,\alpha_\mu}\right)^{\frac{1}{\alpha_\mu}}\right)^{\alpha}\right)^{\frac{\alpha}{1-\alpha}}\left(\left(\frac{\alpha}{\mu b\,\alpha_\mu}\right)^{\frac{1}{\alpha_\mu}}\right)^{\alpha}\left(-\alpha+\mu b\left(\left(\frac{\alpha}{\mu b\,\alpha_\mu}\right)^{\frac{1}{\alpha_\mu}}\right)^{\alpha_\mu}\alpha_\mu\right)+\right.\right.$$

$$\left(-2+\left(e^{-\mu b\left(\left(\frac{\alpha}{\mu b\,\alpha_\mu}\right)^{\frac{1}{\alpha_\mu}}\right)^{\alpha_\mu}}\,sb\left(\left(\frac{\alpha}{\mu b\,\alpha_\mu}\right)^{\frac{1}{\alpha_\mu}}\right)^{\alpha}\right)^{\frac{1}{1-\alpha}}\right)$$

$$\left(-\alpha\left(\left(e^{-\mu b\left(\left(\frac{\alpha}{\mu b\,\alpha_\mu}\right)^{\frac{1}{\alpha_\mu}}\right)^{\alpha_\mu}}\,sb\left(\left(\frac{\alpha}{\mu b\,\alpha_\mu}\right)^{\frac{1}{\alpha_\mu}}\right)^{\alpha}\right)^{\frac{1}{1-\alpha}}\left(\frac{\alpha}{\mu b\,\alpha_\mu}\right)^{\frac{1}{\alpha_\mu}}\right)^{\alpha}+\right.$$

$$\left.\left.\left.\mu b\left(\left(e^{-\mu b\left(\left(\frac{\alpha}{\mu b\,\alpha_\mu}\right)^{\frac{1}{\alpha_\mu}}\right)^{\alpha_\mu}}\,sb\left(\left(\frac{\alpha}{\mu b\,\alpha_\mu}\right)^{\frac{1}{\alpha_\mu}}\right)^{\alpha}\right)^{\frac{1}{1-\alpha}}\right)^{\alpha}\left(\left(\frac{\alpha}{\mu b\,\alpha_\mu}\right)^{\frac{1}{\alpha_\mu}}\right)^{\alpha+\alpha_\mu}\alpha_\mu\right)\right)\right)\Big/$$

$$\left.\left(e^{\mu b\left(\left(\frac{\alpha}{\mu b\,\alpha_\mu}\right)^{\frac{1}{\alpha_\mu}}\right)^{\alpha_\mu}}-sb\left(\left(e^{-\mu b\left(\left(\frac{\alpha}{\mu b\,\alpha_\mu}\right)^{\frac{1}{\alpha_\mu}}\right)^{\alpha_\mu}}\,sb\left(\left(\frac{\alpha}{\mu b\,\alpha_\mu}\right)^{\frac{1}{\alpha_\mu}}\right)^{\alpha}\right)^{\frac{1}{1-\alpha}}\left(\frac{\alpha}{\mu b\,\alpha_\mu}\right)^{\frac{1}{\alpha_\mu}}\right)^{\alpha}\right)^{2}\right)\right\},$$

$$\left\{-\left(\left(\left(sb^{2}\,\alpha\left(1-\left(\left(e^{-\mu b\left(\left(\frac{\alpha}{\mu b\,\alpha_\mu}\right)^{\frac{1}{\alpha_\mu}}\right)^{\alpha_\mu}}\,sb\left(\left(\frac{\alpha}{\mu b\,\alpha_\mu}\right)^{\frac{1}{\alpha_\mu}}\right)^{\alpha}\right)^{\frac{1}{1-\alpha}}\right)^{\alpha}\right)\left(e^{-\mu b\left(\left(\frac{\alpha}{\mu b\,\alpha_\mu}\right)^{\frac{1}{\alpha_\mu}}\right)^{\alpha_\mu}}\,sb\left(\left(\frac{\alpha}{\mu b\,\alpha_\mu}\right)^{\frac{1}{\alpha_\mu}}\right)^{\alpha}\right)^{-1+\frac{1}{-1+\alpha}}\right.\right.\right.$$

$$\left(\left(\frac{\alpha}{\mu b\,\alpha_\mu}\right)^{\frac{1}{\alpha_\mu}}\right)^{2\alpha}\left(-\left(\left(-1+\left(e^{-\mu b\left(\left(\frac{\alpha}{\mu b\,\alpha_\mu}\right)^{\frac{1}{\alpha_\mu}}\right)^{\alpha_\mu}}\,sb\left(\left(\frac{\alpha}{\mu b\,\alpha_\mu}\right)^{\frac{1}{\alpha_\mu}}\right)^{\alpha}\right)^{\frac{1}{1-\alpha}}\right)\left(\frac{\alpha}{\mu b\,\alpha_\mu}\right)^{\frac{1}{\alpha_\mu}}\right)\right)^{-1-\alpha}$$

$$\left(\left(\left(e^{-\mu b\left(\left(\frac{\alpha}{\mu b\,\alpha_\mu}\right)^{\frac{1}{\alpha_\mu}}\right)^{\alpha_\mu}}\,sb\left(\left(\frac{\alpha}{\mu b\,\alpha_\mu}\right)^{\frac{1}{\alpha_\mu}}\right)^{\alpha}\right)^{\frac{\alpha}{1-\alpha}}\left(\left(\frac{\alpha}{\mu b\,\alpha_\mu}\right)^{\frac{1}{\alpha_\mu}}\right)^{\alpha}\left(-\alpha+\mu b\left(\left(\frac{\alpha}{\mu b\,\alpha_\mu}\right)^{\frac{1}{\alpha_\mu}}\right)^{\alpha_\mu}\alpha_\mu\right)+\right.\right.$$

$$\left(-2+\left(e^{-\mu b\left(\left(\frac{\alpha}{\mu b\,\alpha_\mu}\right)^{\frac{1}{\alpha_\mu}}\right)^{\alpha_\mu}}\,sb\left(\left(\frac{\alpha}{\mu b\,\alpha_\mu}\right)^{\frac{1}{\alpha_\mu}}\right)^{\alpha}\right)^{\frac{1}{1-\alpha}}\right)$$

$$\left(-\alpha\left(\left(e^{-\mu b\left(\left(\frac{\alpha}{\mu b\,\alpha_\mu}\right)^{\frac{1}{\alpha_\mu}}\right)^{\alpha_\mu}}\,sb\left(\left(\frac{\alpha}{\mu b\,\alpha_\mu}\right)^{\frac{1}{\alpha_\mu}}\right)^{\alpha}\right)^{\frac{1}{1-\alpha}}\left(\frac{\alpha}{\mu b\,\alpha_\mu}\right)^{\frac{1}{\alpha_\mu}}\right)^{\alpha}+\right.$$

$$\left.\left.\left.\mu b\left(\left(e^{-\mu b\left(\left(\frac{\alpha}{\mu b\,\alpha_\mu}\right)^{\frac{1}{\alpha_\mu}}\right)^{\alpha_\mu}}\,sb\left(\left(\frac{\alpha}{\mu b\,\alpha_\mu}\right)^{\frac{1}{\alpha_\mu}}\right)^{\alpha}\right)^{\frac{1}{1-\alpha}}\right)^{\alpha}\left(\left(\frac{\alpha}{\mu b\,\alpha_\mu}\right)^{\frac{1}{\alpha_\mu}}\right)^{\alpha+\alpha_\mu}\alpha_\mu\right)\right)\right)\Big/$$

$$\Bigg(e^{\mu b\left(\left(\frac{\alpha}{\mu b\,\alpha_\mu}\right)^{\frac{1}{\alpha_\mu}}\right)^{\alpha_\mu}} - sb\Bigg(\Bigg(e^{-\mu b\left(\left(\frac{\alpha}{\mu b\,\alpha_\mu}\right)^{\frac{1}{\alpha_\mu}}\right)^{\alpha_\mu}}\, sb\left(\left(\frac{\alpha}{\mu b\,\alpha_\mu}\right)^{\frac{1}{\alpha_\mu}}\right)^{\alpha}\Bigg)^{\frac{1}{1-\alpha}}\left(\frac{\alpha}{\mu b\,\alpha_\mu}\right)^{\frac{1}{\alpha_\mu}}\Bigg)^{\alpha}\Bigg)^{2}\Bigg),$$

$$\Bigg(e^{\mu b\left(\left(\frac{\alpha}{\mu b\,\alpha_\mu}\right)^{\frac{1}{\alpha_\mu}}\right)^{\alpha_\mu}}\Bigg(1 - \Bigg(e^{-\mu b\left(\left(\frac{\alpha}{\mu b\,\alpha_\mu}\right)^{\frac{1}{\alpha_\mu}}\right)^{\alpha_\mu}}\, sb\left(\left(\frac{\alpha}{\mu b\,\alpha_\mu}\right)^{\frac{1}{\alpha_\mu}}\right)^{\alpha}\Bigg)^{\frac{1}{1-\alpha}}\Bigg)^{\alpha}\left(\left(\frac{\alpha}{\mu b\,\alpha_\mu}\right)^{\frac{1}{\alpha_\mu}}\right)^{-2+\alpha}$$

$$\Bigg(-\Bigg(\Bigg(-1+\Bigg(e^{-\mu b\left(\left(\frac{\alpha}{\mu b\,\alpha_\mu}\right)^{\frac{1}{\alpha_\mu}}\right)^{\alpha_\mu}}\, sb\left(\left(\frac{\alpha}{\mu b\,\alpha_\mu}\right)^{\frac{1}{\alpha_\mu}}\right)^{\alpha}\Bigg)^{\frac{1}{1-\alpha}}\Bigg)\left(\frac{\alpha}{\mu b\,\alpha_\mu}\right)^{\frac{1}{\alpha_\mu}}\Bigg)\Bigg)^{-\alpha}$$

$$\Bigg(sb\,\alpha\,(1+\alpha)\Bigg(\Bigg(e^{-\mu b\left(\left(\frac{\alpha}{\mu b\,\alpha_\mu}\right)^{\frac{1}{\alpha_\mu}}\right)^{\alpha_\mu}}\, sb\left(\left(\frac{\alpha}{\mu b\,\alpha_\mu}\right)^{\frac{1}{\alpha_\mu}}\right)^{\alpha}\Bigg)^{\frac{1}{1-\alpha}}\left(\frac{\alpha}{\mu b\,\alpha_\mu}\right)^{\frac{1}{\alpha_\mu}}\Bigg)^{\alpha} +$$

$$sb\,\mu b\Bigg(\Bigg(e^{-\mu b\left(\left(\frac{\alpha}{\mu b\,\alpha_\mu}\right)^{\frac{1}{\alpha_\mu}}\right)^{\alpha_\mu}}\, sb\left(\left(\frac{\alpha}{\mu b\,\alpha_\mu}\right)^{\frac{1}{\alpha_\mu}}\right)^{\alpha}\Bigg)^{\frac{1}{1-\alpha}}\Bigg)^{\alpha}\left(\left(\frac{\alpha}{\mu b\,\alpha_\mu}\right)^{\frac{1}{\alpha_\mu}}\right)^{\alpha+\alpha_\mu}$$

$$\alpha_\mu\left(-1-2\,\alpha+\left(1+\mu b\left(\left(\frac{\alpha}{\mu b\,\alpha_\mu}\right)^{\frac{1}{\alpha_\mu}}\right)^{\alpha_\mu}\right)\alpha_\mu\right) + e^{\mu b\left(\left(\frac{\alpha}{\mu b\,\alpha_\mu}\right)^{\frac{1}{\alpha_\mu}}\right)^{\alpha_\mu}}$$

$$\Bigg((-1+\alpha)\,\alpha+\mu b\left(\left(\frac{\alpha}{\mu b\,\alpha_\mu}\right)^{\frac{1}{\alpha_\mu}}\right)^{\alpha_\mu}\alpha_\mu\left(1-2\,\alpha+\left(-1+\mu b\left(\left(\frac{\alpha}{\mu b\,\alpha_\mu}\right)^{\frac{1}{\alpha_\mu}}\right)^{\alpha_\mu}\right)\alpha_\mu\right)\Bigg)\Bigg)\Bigg)\Big/$$

$$\Bigg(e^{\mu b\left(\left(\frac{\alpha}{\mu b\,\alpha_\mu}\right)^{\frac{1}{\alpha_\mu}}\right)^{\alpha_\mu}} - sb\Bigg(\Bigg(e^{-\mu b\left(\left(\frac{\alpha}{\mu b\,\alpha_\mu}\right)^{\frac{1}{\alpha_\mu}}\right)^{\alpha_\mu}}\, sb\left(\left(\frac{\alpha}{\mu b\,\alpha_\mu}\right)^{\frac{1}{\alpha_\mu}}\right)^{\alpha}\Bigg)^{\frac{1}{1-\alpha}}\left(\frac{\alpha}{\mu b\,\alpha_\mu}\right)^{\frac{1}{\alpha_\mu}}\Bigg)^{\alpha}\Bigg)^{2}\}\}$$

##### 1.2.3. Initialize the Jacobian for checking convergence stability

In[ ]:=
```
(*Recall selection gradients*)
```

In[ ]:=
```
μ[vr_] := μb * (1 - vr)^αμ
```

In[ ]:=
```
sp[vs_, vr_] := sb * (vs * (1 - vr))^αs * e^-μ[vr]
```

In[ ]:=
```
Sur[vs_, vr_] :=
  1/((1 - vr) (1 - sp[vs, vr])) (μ[vr] αμ - (αs * sp[vs, vr] + αf * (1 - sp[vs, vr]))) /.
   {αf → α, αs → α } // FullSimplify
```

```
In[•]:= Sus[vs_, vr_] :=
```

$$\frac{1}{(1 - sp[vs, vr])}\left(\frac{\alpha_s * sp[vs, vr]}{vs} - \frac{\alpha_f * (1 - sp[vs, vr])}{(1 - vs)}\right) /. \{\alpha_f \to \alpha, \alpha_s \to \alpha\} //$$

```
FullSimplify
```

In[•]:= $vrSol = 1 - \left(\frac{\alpha}{\mu b\, \alpha_\mu}\right)^{\frac{1}{\alpha_\mu}};$

$$vsSol = \left(e^{-\mu b \left(\left(\frac{\alpha}{\mu b\, \alpha_\mu}\right)^{\frac{1}{\alpha_\mu}}\right)^{\alpha_\mu}} sb\left(\left(\frac{\alpha}{\mu b\, \alpha_\mu}\right)^{\frac{1}{\alpha_\mu}}\right)^{\alpha}\right)^{\frac{1}{1-\alpha}};$$

In[•]:=

$JacobEx1[\mu b\_] = FullSimplify[D[\{Sur[vs, vr], Sus[vs, vr]\}, \{\{vr, vs\}\}] /. \{\alpha_s \to \alpha, \alpha_f \to \alpha, \alpha\mu \to \alpha_\mu\}] /. \{vs \to vsSol, vr \to vrSol\}$

### 1.3. Individual-based simulation code

**Comments about the simulation code:**

The simulations then follow a population composed of a finite and fixed number (=7500 in our simulations) of individuals , where each individual is described by a vector of traits consisting of allocation to repair, allocation to survival and number of deleterious mutations the individual has (see function “PopIteration” in section 1.3.1). Starting with a monomorphic population (function “PopInitial” in section 1.3.3.), we then track the stochastic change in “population state” vector (consisting of the vector of states of each individual) for a fixed number (=7500) of generations of life-cycle iterations (function “PopMeanDyno” in section 1.3.3. , which calls the function “PopDynoFullStat” in section 1.3.1.).

Function “PopIteration”: updates the population by accounting for all the life-cycle steps from one generation to the next: (i) reproduction, (ii) survival, (iii) mutation in life-history locus and deleterious mutations in the locus, where deleterious mutations can accumulate. These three steps are explained in further detail in Appendix B of the paper and comments are also given in the function “PopIteration”. In section “Model Specification” (section 1.3.2.) we define the parameters used to compute “PopIteration” and specify fecundity . Note that in the simulations code we use slightly different notations and section “Model Specification” lists all these notations.

Function “PopDynoFullStat”: updates the population from one generation to the next by calling the function “PopInitial” and storing relevant statistics in “data” vector. The “data” vector contains the following information: generation time, time average of “allocation to maintenance”, variance of “allocation to maintenance”, time average of “allocation to survival”, variance of “allocation to survival”, time average “number of novel mutations”, variance of “number of novel mutations”, and “population state” vector.

In section “Code for running the Program”, we use the “Do” loop to runs the simulation program with different values of the baseline mutation rate and we specify the file where we store the output data. The output of simulations is copied into section 1.4. On average, running the “Do” loop once for the chosen parameter values took about 3-4 hours. In order to reproduce the graph we ran the Do loop three times for a different value of the scaling parameter $\alpha$ of vital rates.

```
In[ ]:= Clear["Global`*"]
```

##### 1.3.1. Population mapping

```
(* Individual={x,y,n} where x is investment into maintenance,
y investment into survival,and n is the number of
 deleterious mutations. Population is an array of individuals *)

PopIteration[Pop_] :=
Block[{Fertility, PopAfterSurviv, PopBeforDeletMut, NewGenerationPheno,
   MutationPhenoVec, MutationDeleteriousVec, MutationMatrix},

  (* Arrays of fertility *)

  Fertility = Table[f[Pop⟦i, 1⟧, Pop⟦i, 2⟧, Pop⟦i, 3⟧], {i, Nind}];

  (* Array of individuals after survival *)

  PopAfterSurviv = DeleteCases[
    Map[If[Random[BernoulliDistribution[s[#⟦1⟧, #⟦2⟧, #⟦3⟧]]] == 0, {}, #] &, Pop],
    _{}];

  (* Array of individuals after survival and reproduction: joints the
     phenotypes of the surviving individuals and those of the newborns *)

  NewGenerationPheno = PopAfterSurviv~Join~
    RandomChoice[Fertility → Pop, Nind - Length[PopAfterSurviv]];

  (* Calculate mutation effect sizes and
   number of deleterious mutation for each individual *)

  MutationMatrix =
   Table[If[RandomChoice[{1 - Pmutation, Pmutation} → {0, 1}] == 0, {0, 0, 0},
     RandomVariate[BinormalDistribution[{mutationStepx, mutationStepy}, 0]]~
      Join~{0}], {i, Nind}];

   (* Construct new generation from fitness and mutations *)

  PopBeforDeletMut = MapAt[Clip[#, {0, 1}] &,
    Evaluate[NewGenerationPheno + MutationMatrix], {{All, 1}, {All, 2}}];

  (* Array of phenotypes after deleterious mutations *)

  Return[ Table[{PopBeforDeletMut⟦i, 1⟧, PopBeforDeletMut⟦i, 2⟧,
     DeletMut[PopBeforDeletMut⟦i, 1⟧, PopBeforDeletMut⟦i, 3⟧]}, {i, Nind}]]]
```

```
In[ ]:= (*MutationDeleteriousVec=
        ParallelTable[If[μRate[PopPhenotypes[[i,1]]]>0,RandomVariate[
          PoissonDistribution[μRate[PopPhenotypes[[i,1]]]]],0],{i, Nind}];)

     MutationMatrix=ParallelTable[
     Join[MutationPhenoVec[[i]],{MutationDeleteriousVec[[i]]}], {i, Nind}];*)

(*Here the data vector which is the argument of PopDynoFullStat
  stores all the relevant statistics about the simulation: data=
 (time, mean "allocation to maintenance",
    variance of "allocation to maintenance", mean "allocation to survival",
    variance of "allocation to survival", mean "number of novel mutations",
    variance of "number of novel mutations", population). Note
  that here variance is computed as a sample variance. *)

PopDynoFullStat[data_] := Block[{PopNew, Time, MeanUr, MeanUs, MeanNu},
  PopNew = PopIteration[data[[9]]];
  Time = data[[1]] + 1;
  MeanUr = Mean[PopNew[[All, 1]]];
  MeanUs = Mean[PopNew[[All, 2]]];
  MeanNu = Mean[PopNew[[All, 3]]];
  {Time,
```

$$\frac{\text{MeanUr}}{\text{Time}} + \frac{(\text{Time}-1)}{\text{Time}}\,\text{data}_{[[2]]},$$
$$\frac{1}{\text{Time}}\left(\text{MeanUr}^2 - 2\,\text{MeanUr}\,\text{data}_{[[2]]} + \text{data}_{[[2]]}{}^2\right) + \frac{(\text{Time}-1)}{\text{Time}}\,\text{data}_{[[3]]},$$
$$\frac{\text{MeanUs}}{\text{Time}} + \frac{(\text{Time}-1)}{\text{Time}}\,\text{data}_{[[4]]},$$
$$\frac{1}{\text{Time}}\left(\text{MeanUs}^2 - 2\,\text{MeanUs}\,\text{data}_{[[4]]} + \text{data}_{[[4]]}{}^2\right) + \frac{(\text{Time}-1)}{\text{Time}}\,\text{data}_{[[5]]},$$
$$\frac{\text{MeanNu}}{\text{Time}} + \frac{(\text{Time}-1)}{\text{Time}}\,\text{data}_{[[6]]},\ \frac{1}{\text{Time}}\left(\text{MeanNu}^2 - 2\,\text{MeanNu}\,\text{data}_{[[6]]} + \text{data}_{[[6]]}{}^2\right) +$$
$$\frac{(\text{Time}-1)}{\text{Time}}\,\text{data}_{[[7]]},\ \frac{\text{Variance}[\text{PopNew}[[\text{All}, 3]]]}{\text{Time}} + \frac{(\text{Time}-1)}{\text{Time}}\,\text{data}_{[[8]]},\ \text{PopNew}\}]$$

#### 1.3.2. Model specification

Here we specify parameter values and vital rates as functions of traits

x - investment into repair
y - investment into survival
nu - mean number of deleterious mutations

```
(* Fertility function, survival, mutation accumulation functions *)


s=Compile[{{x, _Real},{y,_Real},{nu,_Integer}}, so (y(1-x))^ϕ (1-ss)^nu]; (* Survival func
f=Compile[{{x, _Real},{y,_Real},{nu,_Integer}}, bo ((1-y)(1-x))^ν (1-sf)^nu]; (* Ferility
DeletMut[x_,nu_]:=Block[{μinter}, μinter=μb(1-x)^α; Return[nu+If[μinter>0,RandomVariate[I

(* Mutation parameters for evolving phenotypes *)
Pmutation = 0.01;     (* Probability of mutation in the life-history locus *)
mutationStepx = 0.1 (* Standard deviation in mutation step size for trait for repair *)
mutationStepy = 0.1 (*  Standard deviation in mutation step size for trait for survival

(* Parameter values *)
α=2; (* repair efficiency *)
ν=0.1; (* inverse efficiency of fecundity *)
ϕ=ν; (* inverse efficiency of survival *)
ss=sf; (*ss, sf reductions in survival and fecundity from carrying an additional  delet
sf=0.2; (*Baseline fecundity*)
bo=5; (*Baseline fecundity*)
so=0.5; (*Baseline survival*)
```

In[ ]:=
```
(* Analytical predictions for a uninvdable
 strategy {x, y, nu} from invasion analysis *)
```

In[ ]:=

$$\text{Sing} = \left\{1 - \alpha^{-1/\alpha}\,\mu b^{-1/\alpha}\,\nu^{\frac{1}{\alpha}},\ \left(\frac{e^{\frac{\nu}{\alpha}}\left(\frac{\nu}{\alpha\,\mu b}\right)^{-\frac{\nu}{\alpha}}}{\text{so}}\right)^{\frac{1}{-1+\nu}},\ \left(\frac{\nu/\alpha}{\text{sf}}\right)\right\};$$

#### 1.3.3. Code for running the simulations at equilibrium (data for graphs 2)

In[ ]:=
```
Res = {}; (*Initialize the array where we collect the results*)
```

```
In[•]:= Do[
      Nind = 7500; (* Population size *)
      Ngenerations = 7500;  (* Number of generations *)

      (*Data to check if simulations work*)
      Nind = 10; (* We try small number of indidivuals
  to check if simulations work, comment this line out*)
      Ngenerations = 10;  (* We try small number of generartions
  to check if simulations work, comment this line out *)


      (* Initial condition for the population *)
      initialX = Clip[Sing, {0, 1}][[1]]; (*Initial allocation to repair*)
      initialY = Clip[Sing, {0, 1}][[2]];  (*Initial allocation to survival*)
      initialMut = Sing[[3]];
 (*Initial mean number of deleterious mutations *)
 PopInitial = Table[{initialX, initialY,
    RandomVariate[PoissonDistribution[Sing[[3]]]]}, {j, Nind}];

     If[μb == 0.02, startTime = AbsoluteTime[]];
 (*Start counting simulation time*)

           PopMeanDyno = Nest[PopDynoFullStat,
   {1, initialX, 0, initialY, 0, initialMut, 0, 0, PopInitial}, Ngenerations];

 currentTime = AbsoluteTime[] - startTime;

 AppendTo[Res,
  {μb, PopMeanDyno[[2]] // N, PopMeanDyno[[4]] // N, PopMeanDyno[[6]] // N, currentTime}];
 (*Append current results to the array that collects the results *)

 Print["Current simulation for param:  μb = ", μb, "  Time (s) = ",
  currentTime], {μb, {0.02, 0.06, 0.1, 0.15, 0.2, 0.25, 0.3, 0.35, 0.4, 0.45, 0.5}}
 (*Define values of baseline mortality
  rate for which we want to simulate results*)
]
```

```
Current simulation for param:  μb = 0.02  Time (s) = 0.006256
Current simulation for param:  μb = 0.06  Time (s) = 0.012910
Current simulation for param:  μb = 0.1  Time (s) = 0.022555
Current simulation for param:  μb = 0.15  Time (s) = 0.029660
Current simulation for param:  μb = 0.2  Time (s) = 0.036043
Current simulation for param:  μb = 0.25  Time (s) = 0.042379
Current simulation for param:  μb = 0.3  Time (s) = 0.048235
Current simulation for param:  μb = 0.35  Time (s) = 0.054205
Current simulation for param:  μb = 0.4  Time (s) = 0.059712
Current simulation for param:  μb = 0.45  Time (s) = 0.065504
Current simulation for param:  μb = 0.5  Time (s) = 0.071389
```

In[ ]:=
```
(*This allows to track which simulation is
 currently computing and how long it took in seconds *)
```

In[ ]:=
```
Print[Res]
```

```
{{0.02, 0., 0.460789, 0.0863636, 0.006256},
 {0.06, 0.0865136, 0.433106, 0.0772727, 0.012910},
 {0.1, 0.336343, 0.338482, 0.259091, 0.022555},
 {0.15, 0.42265, 0.411991, 0.0681818, 0.029660},
 {0.2, 0.5, 0.405459, 0.368182, 0.036043},
 {0.25, 0.550738, 0.401138, 0.0954545, 0.042379},
 {0.3, 0.562707, 0.438946, 0.586364, 0.048235},
 {0.35, 0.622471, 0.392673, 0.440909, 0.054205},
 {0.4, 0.646447, 0.390142, 0.113636, 0.059712},
 {0.45, 0.676185, 0.381846, 0.177273, 0.065504},
 {0.5, 0.683772, 0.385336, 0.45, 0.071389}}
```

In[ ]:=
```
(*The Result in the following format: array of vectors of results res(μb)=
 {μb,ur,us, nu,simulTime(sec)} for diffrent values of the baseline
  mutation rate μb and where the first element of the results vector
  indicated the current value of the baseline mutation rate μb,
these Results can be copied to section 1.4 "Simulation results" *)
```

#### 1.3.4. Code for running the simulations outside of equilibrium (data for graphs 3)

```
(* Mutation parameters for evolving phenotypes *)
Pmutation = 0.01;  (* Probability of mutation in the life-history locus *)
mutationStepx = 0.05;
 (* Standard deviation in mutation step size for trait for repair *);
mutationStepy = 0.05;
(*  Standard deviation in mutation step size for trait for survival *);

(* Parameter values *)
α = 2; (* repair efficiency *)
ν = 0.1; (* inverse efficiency of fecundity *)
ϕ = ν; (* inverse efficiency of survival *)
ss = sf; (*ss, sf reductions in survival and fecundity
 from carrying an additional  deleterious mutation*)
sf = 0.2; (*Baseline fecundity*)
bo = 5; (*Baseline fecundity*)
so = 0.5; (*Baseline survival*)
```

In[ ]:=
```
Nind = 6000; (* Population size *)
Ngenerations = 6000; (* Number of generations *)
initialMut = 0.000001;
(*Initial mean number of deleterious mutations *)
μb = 0.2;

(* Initial condition for the population *)
initialX1 = 0.1; (*Initial allocation to repair*)
initialY1 = 0.1;  (*Initial allocation to survival*)

initialX2 = 0.1; (*Initial allocation to repair*)
initialY2 = 0.7;  (*Initial allocation to survival*)

initialX3 = 0.7; (*Initial allocation to repair*)
initialY3 = 0.1;  (*Initial allocation to survival*)

initialX4 = 0.7; (*Initial allocation to repair*)
initialY4 = 0.7;  (*Initial allocation to survival*)
```

```
In[ ]:= PopInitial1 = Table[{initialX1, initialY1,
            RandomVariate[PoissonDistribution[initialMut ]]}, {j, Nind}];
        PopInitial2 = Table[{initialX2, initialY2,
            RandomVariate[PoissonDistribution[initialMut ]]}, {j, Nind}];
        PopInitial3 = Table[{initialX3, initialY3,
            RandomVariate[PoissonDistribution[initialMut ]]}, {j, Nind}];
        PopInitial4 = Table[{initialX4, initialY4,
            RandomVariate[PoissonDistribution[initialMut ]]}, {j, Nind}];

In[ ]:=

In[ ]:= PhenoVecTime1 = NestList[PopIteration, PopInitial1 , Ngenerations];

In[ ]:= PhenoVecTime2 = NestList[PopIteration, PopInitial2 , Ngenerations];

In[ ]:= PhenoVecTime3 = NestList[PopIteration, PopInitial3, Ngenerations];

In[ ]:= PhenoVecTime4 = NestList[PopIteration, PopInitial4 , Ngenerations];


In[ ]:= (*dataFile="/Users/pavila/Dropbox/Apps/convergenceResult.csv";*)

In[ ]:= (*dataStream = OpenWrite[dataFile , PageWidth → Infinity];*)

In[ ]:= urTemp1 =
         Table[Mean[Flatten[PhenoVecTime1⟦i⟧⟦All, 1⟧]], {i, Length[PhenoVecTime1]}]

In[ ]:= usTemp1 =
         Table[Mean[Flatten[PhenoVecTime1⟦i⟧⟦All, 2⟧]], {i, Length[PhenoVecTime1]}]

In[ ]:= urTemp2 =
         Table[Mean[Flatten[PhenoVecTime2⟦i⟧⟦All, 1⟧]], {i, Length[PhenoVecTime2]}]

In[ ]:= usTemp2 =
         Table[Mean[Flatten[PhenoVecTime2⟦i⟧⟦All, 2⟧]], {i, Length[PhenoVecTime2]}]

In[ ]:= urTemp3 =
         Table[Mean[Flatten[PhenoVecTime3⟦i⟧⟦All, 1⟧]], {i, Length[PhenoVecTime3]}]

In[ ]:= usTemp3 =
         Table[Mean[Flatten[PhenoVecTime3⟦i⟧⟦All, 2⟧]], {i, Length[PhenoVecTime3]}]

In[ ]:= urTemp4 =
         Table[Mean[Flatten[PhenoVecTime4⟦i⟧⟦All, 1⟧]], {i, Length[PhenoVecTime4]}]

In[ ]:= usTemp4 =
         Table[Mean[Flatten[PhenoVecTime4⟦i⟧⟦All, 2⟧]], {i, Length[PhenoVecTime4]}]

Out[ ]= {0.7, 0.699954, 0.69998, 0.700042, 0.699877, 0.699774, 0.699871, 0.700125,
         0.700255, 0.700249, 0.699967, 0.699657, 0.699879, 0.699737, 0.699884,
         0.699614, 0.699746, 0.699343, 0.698998, 0.699105, 0.699395, 0.699026,
         0.699079, 0.699102, 0.698817, 0.69933, 0.699349, 0.699319, 0.699457, 0.699642,
         0.699108, 0.698828, 0.699052, 0.698919, 0.699467, 0.699112, 0.698988,
         0.699032, 0.699027, 0.699476, 0.699118, 0.699522, 0.699423, 0.698816,
         0.699088, 0.698884, 0.698263, 0.698233, 0.69775, 0.697409, 0.696293, 0.695493,
         0.695494, 0.69522, 0.695101, 0.695103, 0.694866, 0.694017, 0.692773, 0.693297,
         0.692671, 0.692796, 0.692184, 0.692262, 0.691396, 0.690619, 0.689798,
```

0.689271, 0.689723, 0.690028, 0.690228, 0.690713, 0.69029, 0.690929,
0.690121, 0.690443, 0.689205, 0.688341, 0.687502, 0.687269, 0.687672,
0.686858, 0.687564, 0.686394, 0.685603, 0.68571, 0.6865, 0.686659, 0.686564,
0.686023, 0.686966, 0.68627, 0.685607, 0.684377, 0.684239, 0.68387, 0.682849,
0.682869, 0.682723, 0.68267, 0.681379, 0.68036, 0.680727, 0.680648, 0.680368,
0.680446, 0.679845, 0.677985, 0.677949, 0.67708, 0.676294, 0.675989,
0.676379, 0.676272, 0.676519, 0.676249, 0.674233, 0.673841, 0.672553,
0.673565, 0.672434, 0.672188, 0.672154, 0.671974, 0.671027, 0.670994,
0.669956, 0.669837, 0.670127, 0.670607, 0.669685, 0.669368, 0.670553,
0.669444, 0.667843, 0.668033, 0.668081, 0.666386, 0.664647, 0.663449,
0.663312, 0.662178, 0.66238, 0.662725, 0.661825, 0.661969, 0.661197, 0.660192,
0.65811, 0.656577, 0.656893, 0.656301, 0.655873, 0.656562, 0.65603, 0.654667,
0.653114, 0.65112, 0.650898, 0.64917, 0.64868, 0.648674, 0.650006, 0.650436,
0.649865, 0.65048, 0.649144, 0.647975, 0.646889, 0.645431, 0.646433, 0.64556,
0.643487, 0.641866, 0.640349, 0.640769, 0.638859, 0.637356, 0.635603,
0.633769, 0.631844, 0.631667, 0.629808, 0.630085, 0.632534, 0.631893,
0.629254, 0.628418, 0.626103, 0.626683, 0.625046, 0.621652, 0.6197, 0.617841,
0.614677, 0.61285, 0.612775, 0.612479, 0.609504, 0.609333, 0.607812, 0.606452,
0.604586, 0.603927, 0.603573, 0.602593, 0.600412, 0.599071, 0.598998, 0.59864,
0.599645, 0.598189, 0.598083, 0.596593, 0.59475, 0.594603, 0.593305, 0.593649,
0.593446, 0.591738, 0.592616, 0.593951, 0.592311, 0.591538, 0.59031, 0.588713,
0.586013, 0.585665, 0.58393, 0.583537, 0.585674, 0.58707, 0.586712, 0.586199,
0.585155, 0.585434, 0.585639, 0.585173, 0.584108, 0.582628, 0.582875,
0.582686, 0.583307, 0.581844, 0.582335, 0.582235, 0.582282, 0.581335,
0.582553, 0.583716, 0.583903, 0.583672, 0.583358, 0.582922, 0.584369,
0.582443, 0.579592, 0.579459, 0.579462, 0.579349, 0.576107, 0.574256,
0.574806, 0.572414, 0.572177, 0.570707, 0.567716, 0.568282, 0.567591,
0.567996, 0.567957, 0.567078, 0.566081, 0.566033, 0.56573, 0.566469, 0.565322,
0.565257, 0.563305, 0.562375, 0.561564, 0.560766, 0.558797, 0.558429,
0.556191, 0.554725, 0.554055, 0.555114, 0.55403, 0.553919, 0.553911, 0.552495,
0.553169, 0.553971, 0.552109, 0.552497, 0.551291, 0.549656, 0.550767,
0.549363, 0.549739, 0.550869, 0.551804, 0.550806, 0.550405, 0.549942,
0.549577, 0.547832, 0.546311, 0.545092, 0.544438, 0.543586, 0.542849,
0.542167, 0.543319, 0.54415, 0.544197, 0.543749, 0.543152, 0.543451, 0.542454,
0.541662, 0.540455, 0.53878, 0.538391, 0.537003, 0.537233, 0.53687, 0.535722,
0.535906, 0.535785, 0.534394, 0.534217, 0.532864, 0.52988, 0.529575, 0.528812,
0.529515, 0.527933, 0.529114, 0.528163, 0.529281, 0.527453, 0.525816, 0.5251,
0.526255, 0.525749, 0.523842, 0.523198, 0.523985, 0.523564, 0.522185,
0.520879, 0.51838, 0.518128, 0.517338, 0.516479, 0.516373, 0.515729, 0.517726,
0.517277, 0.515874, 0.514426, 0.513833, 0.513911, 0.51489, 0.514535, 0.515548,
0.514507, 0.514348, 0.514348, 0.515378, 0.514697, 0.512317, 0.509745,
0.508086, 0.508119, 0.507302, 0.508078, 0.506446, 0.504654, 0.503903, 0.50343,
0.503195, 0.501223, 0.501456, 0.500631, 0.499496, 0.497513, 0.496444,
0.496482, 0.495707, 0.494461, 0.495957, 0.495165, 0.496138, 0.496731,
0.495191, 0.495451, 0.497094, 0.497258, 0.496542, 0.49602, 0.494738, 0.493994,
0.493185, 0.492004, 0.491462, 0.491708, 0.491105, 0.489555, 0.488942,
0.488294, 0.488582, 0.489391, 0.49076, 0.49139, 0.492365, 0.490235, 0.491978,

0.491822, 0.492969, 0.491298, 0.491374, 0.48978, 0.487836, 0.488622, 0.488122,
0.486724, 0.486416, 0.485476, 0.485918, 0.486434, 0.486991, 0.484587,
0.483706, 0.48269, 0.484136, 0.484281, 0.482761, 0.482286, 0.482263, 0.482236,
0.481118, 0.481507, 0.480905, 0.48092, 0.477419, 0.476903, 0.476699, 0.475789,
0.47565, 0.475624, 0.476283, 0.474472, 0.47387, 0.473585, 0.473614, 0.47182,
0.471931, 0.473248, 0.472498, 0.472308, 0.470879, 0.470687, 0.472126,
0.472431, 0.47166, 0.472459, 0.475586, 0.473883, 0.47245, 0.473943, 0.472917,
0.472462, 0.472715, 0.471717, 0.4706, 0.472155, 0.472011, 0.471964, 0.4707,
0.470931, 0.469625, 0.470878, 0.471933, 0.470446, 0.468252, 0.466377,
0.467524, 0.466992, 0.466352, 0.464993, 0.465108, 0.465864, 0.467347,
0.466362, 0.465915, 0.466649, 0.466776, 0.467092, 0.465815, 0.466681, 0.46824,
0.466504, 0.466912, 0.467815, 0.465643, 0.464092, 0.463176, 0.464928,
0.464979, 0.463449, 0.462877, 0.463183, 0.462358, 0.463077, 0.463512,
0.463621, 0.464706, 0.464098, 0.46166, 0.460854, 0.460637, 0.462018, 0.461467,
0.46084, 0.460374, 0.459382, 0.460785, 0.461521, 0.46004, 0.459573, 0.460743,
0.460671, 0.461144, 0.459216, 0.461651, 0.462186, 0.462297, 0.462303,
0.461722, 0.460319, 0.460376, 0.461283, 0.459992, 0.459431, 0.458403,
0.459796, 0.460846, 0.459631, 0.459191, 0.458578, 0.459074, 0.459042,
0.457967, 0.458115, 0.459223, 0.460012, 0.46013, 0.459996, 0.459553, 0.459889,
0.457715, 0.458401, 0.458614, 0.459404, 0.461495, 0.460982, 0.460452,
0.460732, 0.461582, 0.462647, 0.46279, 0.462583, 0.462495, 0.462255, 0.462749,
0.462638, 0.460471, 0.460517, 0.460687, 0.461425, 0.461622, 0.461832,
0.461003, 0.460776, 0.461174, 0.461238, 0.461178, 0.460763, 0.460457,
0.461247, 0.459977, 0.459736, 0.459437, 0.458168, 0.457488, 0.455069,
0.456604, 0.457969, 0.457324, 0.456984, 0.45478, 0.452333, 0.452228, 0.451732,
0.451894, 0.453412, 0.453865, 0.452743, 0.452892, 0.452269, 0.451119,
0.45187, 0.452197, 0.453731, 0.453903, 0.453001, 0.453192, 0.451381,
0.450457, 0.45071, 0.452655, 0.452917, 0.452833, 0.452841, 0.453692,
0.452675, 0.452825, 0.452043, 0.450939, 0.451778, 0.4522, 0.450716, 0.450529,
0.4513, 0.450313, 0.450659, 0.450409, 0.45048, 0.450672, 0.450578, 0.451881,
0.451725, 0.452433, 0.45204, 0.451664, 0.451197, 0.450671, 0.45094, 0.449952,
0.449158, 0.449414, 0.448049, 0.448277, 0.447137, 0.446427, 0.449668,
0.448924, 0.450187, 0.449713, 0.448903, 0.447962, 0.448297, 0.448316,
0.447385, 0.445999, 0.446976, 0.445635, 0.446599, 0.445108, 0.444481,
0.444385, 0.442846, 0.443448, 0.443358, 0.443946, 0.443904, 0.444999,
0.444817, 0.444756, 0.44416, 0.442017, 0.44162, 0.44162, 0.440943, 0.442735,
0.442617, 0.442079, 0.441505, 0.442121, 0.440766, 0.439456, 0.439031,
0.439309, 0.439634, 0.440246, 0.439793, 0.439705, 0.439304, 0.439393,
0.439648, 0.440111, 0.440155, 0.43988, 0.440475, 0.441308, 0.441284, 0.44036,
0.438868, 0.439345, 0.440381, 0.440957, 0.441886, 0.44211, 0.442315, 0.442768,
0.443323, 0.443179, 0.443, 0.444015, 0.444234, 0.444118, 0.444961, 0.445175,
0.446226, 0.444872, 0.444275, 0.443038, 0.443458, 0.442932, 0.442189,
0.441814, 0.442054, 0.440673, 0.44128, 0.441244, 0.439474, 0.440328,
0.439767, 0.438896, 0.439084, 0.438958, 0.439684, 0.438569, 0.437831,
0.439355, 0.440183, 0.440437, 0.439471, 0.439302, 0.439472, 0.440469,
0.442049, 0.442918, 0.441208, 0.440925, 0.440824, 0.439017, 0.438446,
0.437837, 0.438219, 0.437923, 0.437929, 0.437035, 0.436866, 0.436639,

0.436423, 0.436396, 0.435767, 0.435042, 0.433918, 0.43374, 0.433706,
0.434227, 0.435139, 0.434902, 0.433841, 0.43438, 0.434215, 0.435608,
0.435446, 0.434733, 0.434578, 0.436053, 0.436454, 0.436066, 0.438075,
0.438713, 0.438683, 0.437173, 0.438464, 0.43754, 0.436294, 0.436178,
0.437773, 0.438307, 0.440093, 0.438995, 0.439475, 0.439312, 0.440829,
0.440533, 0.440763, 0.439641, 0.438812, 0.438126, 0.438059, 0.437029,
0.437117, 0.437792, 0.438563, 0.438616, 0.43951, 0.440552, 0.439758,
0.440065, 0.440492, 0.440404, 0.440256, 0.442633, 0.443042, 0.44186,
0.441432, 0.441718, 0.441143, 0.441279, 0.441846, 0.440934, 0.440143,
0.44106, 0.441338, 0.441292, 0.44129, 0.441287, 0.441913, 0.441435, 0.441984,
0.441659, 0.44261, 0.44241, 0.444139, 0.442635, 0.441147, 0.440933, 0.438749,
0.440116, 0.43864, 0.437482, 0.437451, 0.434026, 0.434143, 0.433955, 0.43458,
0.434362, 0.434091, 0.433194, 0.432031, 0.432735, 0.433042, 0.433321,
0.434868, 0.436073, 0.435112, 0.43518, 0.435308, 0.435167, 0.435448,
0.435422, 0.435592, 0.434968, 0.433836, 0.434367, 0.433996, 0.433359,
0.433503, 0.434196, 0.435218, 0.435799, 0.435415, 0.435698, 0.433814,
0.4342, 0.433775, 0.4334, 0.43416, 0.433875, 0.434859, 0.434939, 0.434729,
0.435336, 0.435978, 0.43493, 0.436057, 0.436373, 0.435539, 0.437125,
0.435666, 0.436234, 0.435914, 0.436888, 0.436931, 0.43747, 0.438344, 0.43847,
0.436219, 0.435683, 0.43453, 0.434094, 0.432489, 0.430826, 0.429387,
0.429642, 0.429206, 0.429978, 0.427185, 0.427781, 0.426262, 0.424497,
0.423168, 0.423608, 0.424421, 0.423846, 0.424299, 0.425413, 0.425355,
0.427237, 0.426709, 0.426805, 0.428216, 0.426011, 0.42643, 0.426744,
0.426348, 0.425929, 0.426418, 0.425849, 0.425496, 0.424593, 0.423874,
0.422672, 0.42127, 0.420458, 0.419621, 0.420409, 0.420549, 0.420133,
0.421187, 0.422136, 0.42322, 0.423155, 0.422548, 0.423499, 0.424539,
0.424382, 0.424013, 0.4234, 0.423345, 0.425016, 0.426484, 0.427471, 0.428461,
0.428259, 0.428668, 0.427599, 0.427727, 0.428289, 0.429525, 0.430365,
0.429558, 0.431529, 0.431079, 0.43087, 0.430387, 0.429312, 0.429509,
0.429821, 0.426531, 0.425163, 0.423368, 0.424235, 0.422303, 0.422353,
0.424019, 0.423394, 0.42272, 0.422527, 0.422149, 0.422492, 0.422354,
0.423702, 0.425434, 0.426916, 0.425624, 0.425391, 0.424155, 0.423892,
0.423856, 0.423884, 0.423706, 0.423113, 0.424708, 0.424806, 0.424777,
0.426208, 0.427583, 0.427503, 0.424655, 0.424718, 0.42655, 0.425569,
0.425445, 0.424723, 0.422629, 0.420877, 0.42025, 0.420921, 0.421766,
0.422869, 0.422933, 0.423658, 0.423387, 0.424273, 0.422882, 0.424071,
0.425391, 0.424813, 0.425648, 0.425084, 0.42594, 0.428602, 0.429011,
0.427861, 0.428075, 0.429069, 0.430935, 0.430324, 0.431403, 0.431783,
0.431949, 0.433508, 0.433553, 0.432849, 0.434176, 0.434107, 0.43457,
0.43567, 0.434349, 0.435373, 0.436008, 0.436881, 0.436356, 0.436531,
0.437441, 0.438715, 0.437076, 0.437336, 0.436184, 0.43678, 0.437389,
0.436928, 0.437723, 0.438205, 0.438192, 0.438349, 0.438053, 0.439644,
0.438038, 0.438856, 0.437104, 0.438018, 0.437934, 0.437866, 0.436618,
0.435221, 0.433961, 0.434596, 0.434085, 0.433301, 0.433824, 0.434367,
0.433183, 0.432537, 0.431234, 0.431768, 0.431533, 0.431221, 0.430576,
0.429306, 0.428887, 0.428368, 0.427294, 0.428439, 0.428228, 0.428862,
0.428562, 0.427778, 0.428321, 0.426909, 0.426535, 0.424069, 0.423738,

0.42364, 0.424432, 0.425154, 0.423275, 0.421769, 0.4203, 0.420491, 0.419324,
0.418752, 0.419268, 0.418802, 0.418347, 0.416709, 0.416663, 0.418312,
0.419987, 0.420847, 0.420201, 0.419762, 0.420224, 0.419847, 0.418908,
0.420185, 0.419054, 0.418837, 0.421333, 0.419745, 0.420402, 0.420062,
0.418966, 0.416871, 0.416549, 0.416147, 0.414733, 0.414134, 0.412201,
0.413538, 0.414334, 0.41349, 0.413365, 0.412658, 0.4112, 0.410511, 0.407612,
0.406192, 0.407648, 0.408774, 0.407904, 0.407861, 0.409641, 0.409227,
0.411174, 0.410661, 0.412035, 0.410583, 0.409658, 0.410027, 0.409696,
0.409788, 0.410315, 0.410781, 0.409714, 0.408195, 0.409341, 0.411165,
0.410375, 0.410292, 0.410239, 0.409452, 0.410101, 0.41033, 0.411261,
0.410358, 0.411222, 0.411443, 0.409266, 0.409587, 0.411572, 0.41109,
0.412004, 0.412546, 0.411367, 0.411641, 0.409291, 0.409637, 0.409187,
0.409226, 0.408722, 0.407992, 0.409447, 0.40964, 0.409347, 0.410097,
0.410344, 0.411851, 0.410287, 0.41049, 0.41089, 0.41215, 0.411501, 0.411032,
0.410099, 0.411624, 0.412946, 0.412317, 0.411784, 0.410795, 0.410979,
0.409007, 0.409028, 0.408224, 0.408378, 0.407058, 0.407728, 0.407742,
0.407067, 0.407156, 0.407009, 0.407585, 0.408543, 0.407549, 0.407228,
0.405719, 0.40437, 0.403311, 0.404241, 0.406219, 0.408204, 0.408212,
0.407032, 0.407714, 0.409263, 0.41028, 0.410738, 0.411264, 0.408873,
0.407876, 0.407967, 0.40636, 0.406092, 0.407376, 0.410192, 0.412358,
0.413309, 0.414214, 0.414078, 0.412628, 0.413389, 0.413005, 0.412712,
0.410924, 0.410735, 0.412154, 0.411872, 0.410717, 0.41097, 0.409341,
0.409331, 0.409532, 0.408363, 0.408302, 0.408354, 0.40858, 0.408022,
0.407299, 0.406752, 0.407794, 0.408894, 0.409762, 0.409304, 0.409973,
0.410953, 0.411687, 0.411633, 0.411293, 0.412357, 0.410648, 0.411277,
0.410992, 0.410479, 0.410074, 0.4115, 0.4109, 0.412194, 0.412404, 0.412714,
0.41274, 0.410955, 0.411334, 0.412285, 0.411673, 0.409891, 0.410748,
0.411037, 0.411582, 0.411892, 0.41248, 0.412973, 0.412023, 0.411968,
0.41285, 0.411954, 0.411368, 0.410576, 0.411626, 0.412484, 0.41351,
0.414032, 0.414372, 0.415357, 0.414484, 0.41357, 0.412947, 0.412938,
0.414097, 0.41429, 0.413758, 0.413517, 0.412395, 0.411785, 0.413242,
0.413705, 0.412582, 0.412167, 0.41343, 0.414473, 0.412632, 0.412843,
0.411879, 0.413395, 0.411901, 0.412253, 0.410801, 0.411442, 0.411178,
0.410667, 0.410126, 0.409647, 0.410953, 0.4116, 0.412835, 0.412815,
0.413555, 0.412401, 0.412163, 0.411322, 0.409883, 0.408987, 0.409298,
0.40964, 0.409436, 0.40932, 0.409732, 0.40948, 0.409364, 0.408542, 0.409476,
0.407808, 0.40682, 0.408165, 0.408041, 0.408415, 0.407294, 0.4068, 0.407862,
0.407217, 0.407253, 0.406679, 0.406556, 0.404931, 0.406002, 0.407447,
0.407419, 0.407379, 0.407502, 0.407331, 0.40764, 0.407255, 0.406681,
0.40852, 0.408235, 0.40794, 0.406975, 0.406498, 0.40657, 0.407014, 0.406554,
0.406223, 0.405751, 0.405933, 0.404714, 0.404314, 0.404581, 0.404444,
0.404464, 0.403429, 0.40371, 0.403442, 0.406047, 0.405775, 0.408764,
0.408772, 0.409143, 0.407545, 0.408224, 0.407288, 0.407308, 0.405291,
0.405552, 0.406118, 0.406183, 0.407842, 0.408529, 0.40641, 0.408269,
0.407106, 0.406712, 0.406605, 0.40686, 0.406149, 0.405197, 0.403279,
0.404471, 0.40419, 0.404652, 0.401658, 0.401277, 0.400208, 0.399875,
0.400148, 0.399656, 0.400811, 0.401017, 0.401386, 0.402047, 0.40265,

0.403605, 0.403191, 0.402907, 0.404068, 0.404026, 0.403793, 0.404015,
0.403146, 0.401965, 0.399885, 0.399925, 0.399918, 0.399271, 0.400436,
0.401015, 0.400864, 0.399783, 0.400344, 0.401561, 0.402189, 0.401576,
0.399928, 0.399288, 0.398816, 0.400225, 0.400481, 0.399753, 0.399581,
0.40008, 0.399602, 0.400109, 0.399604, 0.40073, 0.401651, 0.401502,
0.401652, 0.400895, 0.400769, 0.401498, 0.401625, 0.402418, 0.402778,
0.403429, 0.403834, 0.403109, 0.402211, 0.401783, 0.401668, 0.40178,
0.401514, 0.401702, 0.402528, 0.402321, 0.40117, 0.400891, 0.400347,
0.399806, 0.400033, 0.39997, 0.399894, 0.399766, 0.400475, 0.401281,
0.400031, 0.399433, 0.39934, 0.39869, 0.397692, 0.396926, 0.396344, 0.39628,
0.396086, 0.396933, 0.396715, 0.397153, 0.397516, 0.397819, 0.396982,
0.396852, 0.398028, 0.397587, 0.398299, 0.397828, 0.398014, 0.398129,
0.397772, 0.398124, 0.396428, 0.396969, 0.396742, 0.3948, 0.395381,
0.396228, 0.396503, 0.396297, 0.39697, 0.396591, 0.396667, 0.397564,
0.39797, 0.399481, 0.400157, 0.400915, 0.400654, 0.400863, 0.401915,
0.401803, 0.403701, 0.404185, 0.404793, 0.405243, 0.405439, 0.405798,
0.404619, 0.407219, 0.406926, 0.406576, 0.406966, 0.406088, 0.40447,
0.403985, 0.403135, 0.40323, 0.403336, 0.403295, 0.401145, 0.400728,
0.401603, 0.400869, 0.400996, 0.399891, 0.399993, 0.398733, 0.399109,
0.40017, 0.400543, 0.399154, 0.397979, 0.40043, 0.400149, 0.400254,
0.399528, 0.400859, 0.401882, 0.402052, 0.40208, 0.402944, 0.402232,
0.401077, 0.401021, 0.399708, 0.399384, 0.399297, 0.399774, 0.400765,
0.400142, 0.399133, 0.400327, 0.401001, 0.401113, 0.400225, 0.401011,
0.402317, 0.402244, 0.401306, 0.401787, 0.402555, 0.402652, 0.403561,
0.402482, 0.401329, 0.400173, 0.398566, 0.399561, 0.399063, 0.397897,
0.397099, 0.395804, 0.396065, 0.394635, 0.393331, 0.393317, 0.394426,
0.393614, 0.393622, 0.392956, 0.392744, 0.393021, 0.392529, 0.392298,
0.392345, 0.391164, 0.391693, 0.391269, 0.392965, 0.39277, 0.393311,
0.393167, 0.392841, 0.393102, 0.393323, 0.393122, 0.393149, 0.393787,
0.392284, 0.391437, 0.392225, 0.3918, 0.392886, 0.393421, 0.392107,
0.390741, 0.391456, 0.391019, 0.392106, 0.392611, 0.393767, 0.394228,
0.395064, 0.39573, 0.397575, 0.397947, 0.395965, 0.396266, 0.396413,
0.396873, 0.394843, 0.392576, 0.391844, 0.390692, 0.3902, 0.38997, 0.391539,
0.391614, 0.391273, 0.392001, 0.392544, 0.391375, 0.391342, 0.391253,
0.390419, 0.390593, 0.390599, 0.391149, 0.390732, 0.391054, 0.392566,
0.391789, 0.391129, 0.391997, 0.392619, 0.392044, 0.393336, 0.394963,
0.394547, 0.394851, 0.396294, 0.395697, 0.393624, 0.394769, 0.394758,
0.39534, 0.395729, 0.395317, 0.396092, 0.396478, 0.396187, 0.397589,
0.397341, 0.397544, 0.397215, 0.396995, 0.397081, 0.396411, 0.395885,
0.397336, 0.396254, 0.396035, 0.396127, 0.396705, 0.396792, 0.395813,
0.395531, 0.396747, 0.396829, 0.397342, 0.399175, 0.399068, 0.398297,
0.398765, 0.398112, 0.397444, 0.396887, 0.395961, 0.396565, 0.396413,
0.397735, 0.398948, 0.399805, 0.399691, 0.400544, 0.400888, 0.40201,
0.402456, 0.401288, 0.401161, 0.400863, 0.401121, 0.401378, 0.402212,
0.401328, 0.401985, 0.401469, 0.400856, 0.400604, 0.401034, 0.401285,
0.400607, 0.400342, 0.400116, 0.399645, 0.398861, 0.399029, 0.398207,
0.398464, 0.398311, 0.398703, 0.397792, 0.396862, 0.397145, 0.398145,

0.398153, 0.398415, 0.397956, 0.397497, 0.398003, 0.398864, 0.399928,
0.400982, 0.40013, 0.401136, 0.401253, 0.401847, 0.401636, 0.401124,
0.401828, 0.401629, 0.401936, 0.402673, 0.402068, 0.401941, 0.402413,
0.402698, 0.403239, 0.403633, 0.402853, 0.403691, 0.403761, 0.404796,
0.403941, 0.404581, 0.405503, 0.406225, 0.406528, 0.406862, 0.407795,
0.408816, 0.409525, 0.409705, 0.40923, 0.408391, 0.409147, 0.40805,
0.40803, 0.408954, 0.410271, 0.409744, 0.408302, 0.408343, 0.406574,
0.406924, 0.406623, 0.405123, 0.405282, 0.406104, 0.406156, 0.407777,
0.407842, 0.408632, 0.406705, 0.405603, 0.405817, 0.405412, 0.405957,
0.405898, 0.406381, 0.406781, 0.407267, 0.407923, 0.407292, 0.407961,
0.408417, 0.408026, 0.406335, 0.406011, 0.406333, 0.405733, 0.405473,
0.405226, 0.404888, 0.404436, 0.404726, 0.404893, 0.405254, 0.404864,
0.404193, 0.403801, 0.403854, 0.404313, 0.404321, 0.405248, 0.405057,
0.404532, 0.403007, 0.403211, 0.403133, 0.402086, 0.400588, 0.400986,
0.402197, 0.400674, 0.40012, 0.39908, 0.398702, 0.398555, 0.399111,
0.398097, 0.397955, 0.397315, 0.398222, 0.398586, 0.4004, 0.398733,
0.399566, 0.399982, 0.399527, 0.398781, 0.398073, 0.400722, 0.400248,
0.399119, 0.399896, 0.399885, 0.39951, 0.400488, 0.399414, 0.398953,
0.398325, 0.398494, 0.399944, 0.400146, 0.397945, 0.399148, 0.399962,
0.400713, 0.400988, 0.401485, 0.40311, 0.403857, 0.404925, 0.404651,
0.406237, 0.405553, 0.405299, 0.406477, 0.406208, 0.406893, 0.407456,
0.40831, 0.407917, 0.409155, 0.409266, 0.410309, 0.409933, 0.410344,
0.411692, 0.413203, 0.414343, 0.41492, 0.415316, 0.414007, 0.414224,
0.413276, 0.414227, 0.413669, 0.413251, 0.41437, 0.412706, 0.411854,
0.410298, 0.410225, 0.410581, 0.408362, 0.40893, 0.40797, 0.406533,
0.406533, 0.405145, 0.405123, 0.404475, 0.404548, 0.403061, 0.404452,
0.403797, 0.405725, 0.402686, 0.402598, 0.401687, 0.402479, 0.40288,
0.401775, 0.400774, 0.400886, 0.401593, 0.401202, 0.400394, 0.399905,
0.401308, 0.400037, 0.400396, 0.399763, 0.398983, 0.398696, 0.398528,
0.400456, 0.400232, 0.398199, 0.398793, 0.400266, 0.401242, 0.39977,
0.398291, 0.397978, 0.398486, 0.399656, 0.401486, 0.401254, 0.40083,
0.400354, 0.400293, 0.399364, 0.399892, 0.40083, 0.401766, 0.402927,
0.402915, 0.40175, 0.402392, 0.403692, 0.402761, 0.400892, 0.400345,
0.400412, 0.400401, 0.400209, 0.399739, 0.400324, 0.401367, 0.399632,
0.397202, 0.396258, 0.395841, 0.396809, 0.397256, 0.399034, 0.398525,
0.398296, 0.398234, 0.397823, 0.397516, 0.398215, 0.397488, 0.397094,
0.398778, 0.399272, 0.400596, 0.400655, 0.401453, 0.40178, 0.401464,
0.402451, 0.401731, 0.40105, 0.399954, 0.401978, 0.40178, 0.40142,
0.402403, 0.402415, 0.404152, 0.403795, 0.403184, 0.40197, 0.402907,
0.401983, 0.401989, 0.402783, 0.403356, 0.403324, 0.402365, 0.404222,
0.405787, 0.407518, 0.408071, 0.409533, 0.409794, 0.411996, 0.410773,
0.409997, 0.411202, 0.412361, 0.411358, 0.412536, 0.412947, 0.412932,
0.412049, 0.410892, 0.411142, 0.410433, 0.409844, 0.407302, 0.407768,
0.407858, 0.408341, 0.407968, 0.409021, 0.40794, 0.406926, 0.407066,
0.406319, 0.404973, 0.403275, 0.403777, 0.405611, 0.406591, 0.407478,
0.407349, 0.408091, 0.409449, 0.40744, 0.408915, 0.410417, 0.409946,
0.40875, 0.407921, 0.407498, 0.407728, 0.408158, 0.409342, 0.409025,

0.408626, 0.408559, 0.40672, 0.407497, 0.405743, 0.406139, 0.407862,
0.405874, 0.405936, 0.405629, 0.405567, 0.405171, 0.405722, 0.407628,
0.407396, 0.405691, 0.404718, 0.404021, 0.40386, 0.402749, 0.404987,
0.404219, 0.40659, 0.406515, 0.407027, 0.40734, 0.407869, 0.407721,
0.407618, 0.408464, 0.406159, 0.4074, 0.408777, 0.409983, 0.408887,
0.410002, 0.412152, 0.410814, 0.410338, 0.411529, 0.410439, 0.410168,
0.410794, 0.410217, 0.410661, 0.412058, 0.412433, 0.412802, 0.41241,
0.413269, 0.414469, 0.414891, 0.415619, 0.415466, 0.41319, 0.411507,
0.410672, 0.411719, 0.41121, 0.410947, 0.408988, 0.409876, 0.409287,
0.409842, 0.410199, 0.410734, 0.410989, 0.409428, 0.409255, 0.407379,
0.408711, 0.407125, 0.407849, 0.408179, 0.407881, 0.407923, 0.408717,
0.409546, 0.408754, 0.408327, 0.407958, 0.4089, 0.408532, 0.407587,
0.407963, 0.407647, 0.409588, 0.410347, 0.411113, 0.409265, 0.408622,
0.408468, 0.406912, 0.4067, 0.405877, 0.406785, 0.407452, 0.406453,
0.406228, 0.407251, 0.406122, 0.402484, 0.402975, 0.402328, 0.40203,
0.402025, 0.401651, 0.403568, 0.401885, 0.401725, 0.401929, 0.401247,
0.402161, 0.401653, 0.401848, 0.401989, 0.4009, 0.400474, 0.39911, 0.399814,
0.400485, 0.400663, 0.401953, 0.402473, 0.401264, 0.401058, 0.399086,
0.401441, 0.40018, 0.39977, 0.397586, 0.397894, 0.396397, 0.394571,
0.394868, 0.394361, 0.395845, 0.395225, 0.395546, 0.394841, 0.395169,
0.394136, 0.39373, 0.393551, 0.391325, 0.392313, 0.391694, 0.39113,
0.390647, 0.390489, 0.389431, 0.388235, 0.388451, 0.386789, 0.385153,
0.386013, 0.384957, 0.386485, 0.388856, 0.389593, 0.388908, 0.38939,
0.389761, 0.389152, 0.39038, 0.391746, 0.390938, 0.392408, 0.391528,
0.392122, 0.39147, 0.391546, 0.392205, 0.392088, 0.391845, 0.391473,
0.392437, 0.392329, 0.393759, 0.393236, 0.393838, 0.394601, 0.39546,
0.396986, 0.396746, 0.3972, 0.397608, 0.397453, 0.397052, 0.396275,
0.396963, 0.397001, 0.395506, 0.394972, 0.394659, 0.39524, 0.39484,
0.395133, 0.394524, 0.393142, 0.394075, 0.396175, 0.396678, 0.396713,
0.397726, 0.396726, 0.397065, 0.398947, 0.396579, 0.396493, 0.395518,
0.396099, 0.395102, 0.395515, 0.395456, 0.395569, 0.396633, 0.397068,
0.395787, 0.396705, 0.396769, 0.395081, 0.397013, 0.398655, 0.398416,
0.397276, 0.39827, 0.397252, 0.39777, 0.398114, 0.398508, 0.396741,
0.395891, 0.396695, 0.394911, 0.394513, 0.39505, 0.394777, 0.395245,
0.395182, 0.395867, 0.396902, 0.396535, 0.396477, 0.395242, 0.396484,
0.397024, 0.398843, 0.398983, 0.399619, 0.399022, 0.396471, 0.396761,
0.396058, 0.397324, 0.397598, 0.396023, 0.399172, 0.398977, 0.399979,
0.401354, 0.401927, 0.402151, 0.402701, 0.404261, 0.40513, 0.405197,
0.405727, 0.40505, 0.407004, 0.407197, 0.406874, 0.409298, 0.410073,
0.408813, 0.409458, 0.408823, 0.40975, 0.410511, 0.410164, 0.411117,
0.409358, 0.411468, 0.4108, 0.41082, 0.412401, 0.411379, 0.411714, 0.413408,
0.413315, 0.413743, 0.413363, 0.414233, 0.413881, 0.412406, 0.4104,
0.41105, 0.411203, 0.412248, 0.411673, 0.411598, 0.409234, 0.411007,
0.410119, 0.410164, 0.409243, 0.409418, 0.409774, 0.408939, 0.407828,
0.406641, 0.406124, 0.404103, 0.404309, 0.404261, 0.403525, 0.402952,
0.401828, 0.402588, 0.403497, 0.403237, 0.40325, 0.40351, 0.403963,
0.404315, 0.404184, 0.401791, 0.401704, 0.400833, 0.401112, 0.401674,

0.400928, 0.400656, 0.399987, 0.399903, 0.399741, 0.400155, 0.401381,
0.401196, 0.400803, 0.401838, 0.403049, 0.404064, 0.40652, 0.407552,
0.409925, 0.411537, 0.412242, 0.410404, 0.410333, 0.410582, 0.40942,
0.40783, 0.407536, 0.407651, 0.406436, 0.405203, 0.404743, 0.404212,
0.40486, 0.405985, 0.407464, 0.4086, 0.409406, 0.409587, 0.410202, 0.410991,
0.409697, 0.407765, 0.407872, 0.408005, 0.40688, 0.405601, 0.405053,
0.406652, 0.407034, 0.407428, 0.40655, 0.407096, 0.408392, 0.407085,
0.407896, 0.407525, 0.407372, 0.408949, 0.409859, 0.410154, 0.409092,
0.408204, 0.408823, 0.410073, 0.41068, 0.412315, 0.412246, 0.411829,
0.411644, 0.41063, 0.411547, 0.411278, 0.410907, 0.412467, 0.411871,
0.411383, 0.411298, 0.410836, 0.411051, 0.410281, 0.410783, 0.409568,
0.409883, 0.411821, 0.412044, 0.411236, 0.411446, 0.411113, 0.409286,
0.409488, 0.410676, 0.41128, 0.410524, 0.410961, 0.409233, 0.408281,
0.40756, 0.407891, 0.407918, 0.407991, 0.408475, 0.407836, 0.407702,
0.407406, 0.406577, 0.405972, 0.406552, 0.406327, 0.40647, 0.40848,
0.408024, 0.407134, 0.406566, 0.404376, 0.404519, 0.404731, 0.403766,
0.403993, 0.40476, 0.405431, 0.405237, 0.404245, 0.404186, 0.402894,
0.404612, 0.403692, 0.403619, 0.403, 0.403974, 0.40364, 0.40494, 0.405538,
0.404373, 0.404937, 0.40385, 0.404006, 0.402514, 0.402487, 0.403199,
0.40243, 0.402542, 0.403107, 0.402615, 0.401566, 0.401906, 0.402448,
0.402551, 0.401465, 0.400993, 0.400674, 0.400049, 0.399292, 0.398085,
0.398563, 0.397608, 0.398189, 0.39774, 0.396953, 0.396156, 0.395065,
0.395339, 0.396155, 0.396064, 0.395093, 0.397161, 0.39779, 0.398015,
0.398302, 0.39872, 0.397409, 0.396176, 0.396591, 0.396838, 0.396273,
0.39786, 0.398133, 0.397674, 0.398541, 0.397396, 0.397444, 0.398703,
0.397984, 0.397034, 0.396835, 0.396585, 0.39764, 0.399617, 0.397287,
0.397277, 0.395559, 0.395169, 0.395778, 0.395047, 0.39435, 0.393161,
0.39443, 0.395212, 0.3949, 0.39708, 0.39714, 0.396617, 0.397422, 0.398021,
0.39882, 0.400248, 0.399616, 0.401795, 0.399772, 0.39754, 0.397859,
0.397045, 0.396253, 0.396928, 0.396917, 0.395739, 0.397927, 0.397696,
0.399337, 0.4, 0.400498, 0.401335, 0.4013, 0.400412, 0.400715, 0.400902,
0.400867, 0.401221, 0.403513, 0.403871, 0.40468, 0.403704, 0.404333,
0.404013, 0.40589, 0.406876, 0.405279, 0.405817, 0.407411, 0.408392,
0.411035, 0.409795, 0.409039, 0.408133, 0.407499, 0.406264, 0.407887,
0.40888, 0.409276, 0.410656, 0.410674, 0.410855, 0.411627, 0.410564,
0.411199, 0.411516, 0.411629, 0.411354, 0.411412, 0.412616, 0.412467,
0.411472, 0.411921, 0.411364, 0.410674, 0.410432, 0.409048, 0.408782,
0.409386, 0.408066, 0.407668, 0.407467, 0.409333, 0.408794, 0.409723,
0.410868, 0.412038, 0.411773, 0.410705, 0.412265, 0.412205, 0.411069,
0.410731, 0.409517, 0.407633, 0.40947, 0.409597, 0.409781, 0.40991,
0.411506, 0.412348, 0.410711, 0.410559, 0.410723, 0.410263, 0.41033,
0.410962, 0.411027, 0.411101, 0.412256, 0.413965, 0.416215, 0.41722,
0.417991, 0.418761, 0.421821, 0.422432, 0.422834, 0.421032, 0.421249,
0.420588, 0.422824, 0.424522, 0.424053, 0.425337, 0.426366, 0.427745,
0.428251, 0.428156, 0.430327, 0.431086, 0.431822, 0.430948, 0.43017,
0.431599, 0.429893, 0.428207, 0.428264, 0.428364, 0.429698, 0.428265,
0.428554, 0.427564, 0.425819, 0.42651, 0.426384, 0.426525, 0.428062,

0.42936, 0.429833, 0.430226, 0.430221, 0.429435, 0.427726, 0.428808,
0.430523, 0.431452, 0.431385, 0.433438, 0.433502, 0.433804, 0.434405,
0.434093, 0.435181, 0.436622, 0.438096, 0.436004, 0.435998, 0.434936,
0.435394, 0.434558, 0.434378, 0.433069, 0.432644, 0.432372, 0.433461,
0.433528, 0.434816, 0.433593, 0.432977, 0.434102, 0.433666, 0.43473,
0.435278, 0.434124, 0.433417, 0.43402, 0.432124, 0.432341, 0.43194,
0.434102, 0.434641, 0.434849, 0.434378, 0.43318, 0.432164, 0.432419,
0.430613, 0.430636, 0.430157, 0.431767, 0.432489, 0.43132, 0.431126,
0.431041, 0.431779, 0.43158, 0.431258, 0.432047, 0.431885, 0.432434,
0.430844, 0.430206, 0.431046, 0.429771, 0.429626, 0.430709, 0.430035,
0.429876, 0.430359, 0.429893, 0.429667, 0.429211, 0.427998, 0.426087,
0.428198, 0.428406, 0.428464, 0.429087, 0.427148, 0.426278, 0.425723,
0.426667, 0.427146, 0.426653, 0.426053, 0.424692, 0.424414, 0.424502,
0.424973, 0.424487, 0.423337, 0.42325, 0.424086, 0.424798, 0.425082,
0.425892, 0.42495, 0.422003, 0.420881, 0.419834, 0.419277, 0.417701,
0.419703, 0.421043, 0.420563, 0.420173, 0.420599, 0.420873, 0.420868,
0.423449, 0.423875, 0.423256, 0.424716, 0.425186, 0.425516, 0.425707,
0.426217, 0.426651, 0.427151, 0.427122, 0.42743, 0.426677, 0.426957,
0.42664, 0.429304, 0.42889, 0.429903, 0.428819, 0.429484, 0.430266,
0.431336, 0.430028, 0.429146, 0.428407, 0.426739, 0.425997, 0.424486,
0.422938, 0.421563, 0.423651, 0.423016, 0.423246, 0.422426, 0.42201,
0.421921, 0.421474, 0.421374, 0.420521, 0.418969, 0.417708, 0.419302,
0.417412, 0.417775, 0.41722, 0.4173, 0.418426, 0.416351, 0.416216, 0.417957,
0.416395, 0.41652, 0.414444, 0.414491, 0.415916, 0.41445, 0.414235, 0.4147,
0.416084, 0.415954, 0.413871, 0.414118, 0.414686, 0.41484, 0.414316,
0.415006, 0.414544, 0.414452, 0.413897, 0.414494, 0.414517, 0.414897,
0.41539, 0.415499, 0.417404, 0.417587, 0.417393, 0.416508, 0.416725,
0.416385, 0.415543, 0.414318, 0.413214, 0.412947, 0.414867, 0.414882,
0.416215, 0.41573, 0.415362, 0.415387, 0.414845, 0.414952, 0.414218,
0.413192, 0.414227, 0.413884, 0.413769, 0.412704, 0.411279, 0.409788,
0.409859, 0.409429, 0.409071, 0.409397, 0.408797, 0.408334, 0.407469,
0.40929, 0.410617, 0.410978, 0.411816, 0.411643, 0.411435, 0.41141,
0.410688, 0.411606, 0.411417, 0.411772, 0.413068, 0.413683, 0.413356,
0.412094, 0.412128, 0.413881, 0.413829, 0.414944, 0.413622, 0.413744,
0.413823, 0.412606, 0.411439, 0.411403, 0.410859, 0.411345, 0.411046,
0.410056, 0.411199, 0.411583, 0.412723, 0.413661, 0.413807, 0.415444,
0.415751, 0.415007, 0.415856, 0.416473, 0.415202, 0.415026, 0.41371,
0.413794, 0.412062, 0.413205, 0.415013, 0.414086, 0.413477, 0.413041,
0.413011, 0.414016, 0.414331, 0.414381, 0.413248, 0.412486, 0.413168,
0.413762, 0.413373, 0.41216, 0.411599, 0.412632, 0.412139, 0.412183,
0.412027, 0.41327, 0.414649, 0.414068, 0.4122, 0.411915, 0.412545,
0.412737, 0.413349, 0.412231, 0.411581, 0.41076, 0.411003, 0.41087, 0.4104,
0.409671, 0.410827, 0.410263, 0.411192, 0.411644, 0.412319, 0.412197,
0.41168, 0.411453, 0.411644, 0.411055, 0.411934, 0.410247, 0.408887,
0.409339, 0.411978, 0.411495, 0.411523, 0.410646, 0.410882, 0.411573,
0.41262, 0.411733, 0.412103, 0.412753, 0.412334, 0.411716, 0.411252,
0.409631, 0.410955, 0.410911, 0.409989, 0.409996, 0.411047, 0.411869,

0.411681, 0.410982, 0.413026, 0.41218, 0.412301, 0.411726, 0.411475,
0.411856, 0.411628, 0.411071, 0.411432, 0.413231, 0.412453, 0.413965,
0.414321, 0.414198, 0.414459, 0.41413, 0.413805, 0.41309, 0.414228,
0.413688, 0.412811, 0.412247, 0.412474, 0.412169, 0.412443, 0.412633,
0.413583, 0.41346, 0.41317, 0.414452, 0.414746, 0.414766, 0.415621,
0.414889, 0.41609, 0.416438, 0.416039, 0.415301, 0.41696, 0.41688,
0.417342, 0.41709, 0.417567, 0.417134, 0.417297, 0.417594, 0.418593,
0.418403, 0.419166, 0.419763, 0.418014, 0.418279, 0.419179, 0.41941,
0.420789, 0.42112, 0.421461, 0.422014, 0.420824, 0.420418, 0.420587,
0.419116, 0.420697, 0.421223, 0.422546, 0.422502, 0.421234, 0.421116,
0.420899, 0.422233, 0.420894, 0.420787, 0.422347, 0.422335, 0.421966,
0.422024, 0.422277, 0.422801, 0.423614, 0.424622, 0.424502, 0.425031,
0.424572, 0.423979, 0.423368, 0.424907, 0.424627, 0.424754, 0.424544,
0.425465, 0.426959, 0.426159, 0.425853, 0.426614, 0.425818, 0.427162,
0.426417, 0.426601, 0.426034, 0.425824, 0.426026, 0.426453, 0.425904,
0.425665, 0.423806, 0.423861, 0.423035, 0.423331, 0.423544, 0.422104,
0.420222, 0.420897, 0.419697, 0.41895, 0.418647, 0.419, 0.42012, 0.42006,
0.419692, 0.419346, 0.4191, 0.419726, 0.420256, 0.420342, 0.420476,
0.420873, 0.420913, 0.420597, 0.422124, 0.421767, 0.421489, 0.422327,
0.422386, 0.424135, 0.423641, 0.423625, 0.423467, 0.423655, 0.42321,
0.422998, 0.423939, 0.423325, 0.422526, 0.42091, 0.420872, 0.419975,
0.420496, 0.421892, 0.421249, 0.420047, 0.418589, 0.418653, 0.419192,
0.42013, 0.420567, 0.419395, 0.420439, 0.420156, 0.418929, 0.417842,
0.420321, 0.42046, 0.419924, 0.420928, 0.420875, 0.420861, 0.421016,
0.42135, 0.421852, 0.422121, 0.421352, 0.419979, 0.419024, 0.420474,
0.419837, 0.421022, 0.420299, 0.419955, 0.418321, 0.418232, 0.419631,
0.420416, 0.421701, 0.421672, 0.422137, 0.423071, 0.424182, 0.424064,
0.423481, 0.423858, 0.425397, 0.425463, 0.424329, 0.423732, 0.423797,
0.424579, 0.42502, 0.425016, 0.424145, 0.424007, 0.423959, 0.425145,
0.426011, 0.42592, 0.425343, 0.425885, 0.426641, 0.42609, 0.426155,
0.424149, 0.424402, 0.423446, 0.42395, 0.423256, 0.42326, 0.424578,
0.424391, 0.425503, 0.425139, 0.423451, 0.423014, 0.422263, 0.422236,
0.422416, 0.42208, 0.422453, 0.421406, 0.420874, 0.419784, 0.420109,
0.420536, 0.420401, 0.420155, 0.41946, 0.419341, 0.419354, 0.41806,
0.417209, 0.416735, 0.41694, 0.41835, 0.417842, 0.418642, 0.420565,
0.420841, 0.421553, 0.422714, 0.423662, 0.424158, 0.423966, 0.423288,
0.424218, 0.424568, 0.42714, 0.42607, 0.425626, 0.425506, 0.426239,
0.426688, 0.425408, 0.424148, 0.423551, 0.422863, 0.423234, 0.422424,
0.422245, 0.421795, 0.420369, 0.420552, 0.421314, 0.421599, 0.422079,
0.420452, 0.419684, 0.420651, 0.42101, 0.421677, 0.422338, 0.422561,
0.42245, 0.421635, 0.42221, 0.422926, 0.423239, 0.423723, 0.422556,
0.423768, 0.424758, 0.424938, 0.424694, 0.425573, 0.425266, 0.426656,
0.425315, 0.425508, 0.42565, 0.425945, 0.427699, 0.427719, 0.426161,
0.426921, 0.426185, 0.426762, 0.427288, 0.427212, 0.426807, 0.426485,
0.425723, 0.425393, 0.425574, 0.425374, 0.426899, 0.427029, 0.427197,
0.42762, 0.427223, 0.427877, 0.428666, 0.427954, 0.428886, 0.428327,
0.428243, 0.428529, 0.42812, 0.428245, 0.428569, 0.42914, 0.428202,

0.427917, 0.427857, 0.427902, 0.428354, 0.428285, 0.429333, 0.430971,
0.431123, 0.431111, 0.430167, 0.42958, 0.430167, 0.428774, 0.427617,
0.427162, 0.428015, 0.427893, 0.429755, 0.429425, 0.429831, 0.430585,
0.431232, 0.432816, 0.432357, 0.432381, 0.43162, 0.431013, 0.43012,
0.430757, 0.430605, 0.429825, 0.430001, 0.430374, 0.429703, 0.430181,
0.429447, 0.429064, 0.429724, 0.429749, 0.429905, 0.430498, 0.429306,
0.4285, 0.42877, 0.428716, 0.428158, 0.428307, 0.428651, 0.430334,
0.431049, 0.43131, 0.431545, 0.431162, 0.430235, 0.430338, 0.431165,
0.431289, 0.430334, 0.430123, 0.431713, 0.431888, 0.433721, 0.433487,
0.433622, 0.433309, 0.433553, 0.433188, 0.43362, 0.433795, 0.433418,
0.432711, 0.432447, 0.432205, 0.431807, 0.43055, 0.430717, 0.430796,
0.430282, 0.429897, 0.429948, 0.42995, 0.43034, 0.429599, 0.430749,
0.431275, 0.431145, 0.431339, 0.429355, 0.429329, 0.430041, 0.430095,
0.431235, 0.432305, 0.431853, 0.431288, 0.430718, 0.431499, 0.432917,
0.433968, 0.433921, 0.433585, 0.433839, 0.433547, 0.432919, 0.431669,
0.431818, 0.431663, 0.431916, 0.431502, 0.431288, 0.432607, 0.43376,
0.434405, 0.433862, 0.433874, 0.433289, 0.435157, 0.435304, 0.435034,
0.434893, 0.435326, 0.436162, 0.435377, 0.434527, 0.435213, 0.435477,
0.436158, 0.436125, 0.433888, 0.432681, 0.432719, 0.431931, 0.431651,
0.43095, 0.429891, 0.430155, 0.429025, 0.429859, 0.429313, 0.429176,
0.428902, 0.428114, 0.428569, 0.429295, 0.429345, 0.430267, 0.430251,
0.429716, 0.42942, 0.430783, 0.43016, 0.430969, 0.430972, 0.432219,
0.433058, 0.43138, 0.431406, 0.432258, 0.431608, 0.432555, 0.432524,
0.430725, 0.432287, 0.431722, 0.430065, 0.430469, 0.430756, 0.430601,
0.429946, 0.429086, 0.428609, 0.427506, 0.428006, 0.425664, 0.425298,
0.426, 0.427698, 0.427012, 0.427516, 0.42747, 0.42672, 0.425662, 0.425682,
0.425399, 0.426584, 0.427895, 0.427737, 0.427649, 0.426803, 0.427259,
0.426924, 0.427394, 0.427805, 0.428011, 0.427629, 0.428331, 0.427613,
0.426373, 0.427002, 0.426861, 0.427125, 0.428262, 0.428966, 0.427999,
0.428437, 0.4274, 0.428204, 0.428491, 0.427884, 0.428523, 0.4296, 0.430606,
0.430139, 0.429045, 0.428163, 0.428128, 0.428932, 0.429182, 0.431792,
0.429204, 0.4279, 0.42884, 0.429308, 0.429016, 0.428376, 0.427063,
0.427004, 0.426811, 0.426939, 0.42873, 0.428515, 0.428191, 0.428027,
0.4289, 0.427416, 0.426585, 0.42591, 0.425947, 0.426101, 0.425288,
0.424577, 0.42365, 0.423238, 0.422831, 0.422103, 0.421979, 0.421965,
0.421485, 0.422225, 0.421096, 0.420203, 0.419714, 0.416972, 0.416537,
0.417113, 0.418046, 0.418329, 0.419339, 0.417622, 0.416652, 0.417262,
0.417485, 0.416172, 0.417849, 0.419092, 0.420135, 0.420221, 0.418366,
0.418831, 0.420201, 0.420348, 0.422169, 0.42339, 0.421907, 0.421063,
0.420822, 0.42179, 0.421827, 0.422422, 0.423185, 0.421284, 0.422092,
0.421217, 0.4211, 0.423402, 0.424336, 0.422369, 0.422391, 0.424953,
0.427029, 0.4288, 0.428265, 0.42946, 0.429254, 0.432119, 0.431801,
0.432132, 0.432959, 0.432529, 0.43258, 0.43184, 0.429364, 0.428557,
0.426613, 0.425907, 0.425237, 0.425775, 0.42435, 0.423888, 0.424016,
0.423314, 0.422879, 0.422806, 0.422764, 0.424783, 0.424862, 0.425341,
0.425866, 0.426313, 0.427288, 0.424809, 0.425559, 0.424583, 0.424194,
0.426207, 0.424103, 0.425426, 0.425426, 0.426721, 0.426058, 0.426983,

0.426757, 0.426988, 0.426757, 0.426554, 0.425705, 0.425755, 0.426108,
0.425597, 0.426702, 0.426283, 0.42621, 0.425673, 0.42566, 0.424824,
0.423533, 0.423015, 0.42275, 0.422528, 0.420936, 0.421476, 0.421189,
0.421252, 0.421533, 0.421797, 0.423088, 0.423184, 0.423025, 0.424677,
0.425424, 0.424623, 0.426012, 0.428121, 0.429123, 0.428029, 0.429095,
0.428397, 0.428585, 0.428581, 0.42845, 0.427569, 0.425605, 0.42575,
0.424889, 0.423834, 0.423356, 0.422322, 0.422309, 0.421133, 0.420377,
0.421153, 0.420526, 0.42012, 0.418598, 0.418879, 0.417923, 0.418749,
0.418131, 0.419046, 0.418339, 0.419337, 0.419401, 0.421459, 0.423317,
0.422927, 0.42197, 0.422251, 0.422876, 0.421993, 0.421986, 0.421036,
0.420431, 0.420517, 0.420156, 0.419204, 0.419737, 0.420912, 0.422018,
0.421692, 0.42161, 0.422959, 0.421626, 0.420356, 0.419854, 0.421094,
0.420408, 0.420064, 0.420794, 0.422108, 0.421965, 0.422052, 0.421098,
0.421083, 0.419951, 0.418854, 0.417381, 0.416293, 0.415088, 0.414168,
0.413684, 0.414435, 0.413299, 0.413268, 0.412022, 0.412128, 0.411805,
0.414055, 0.413215, 0.414487, 0.415137, 0.41498, 0.416873, 0.415946,
0.416374, 0.416356, 0.416762, 0.415652, 0.415215, 0.417237, 0.417594,
0.417362, 0.416019, 0.416565, 0.416259, 0.416168, 0.416449, 0.415682,
0.416028, 0.416674, 0.41689, 0.41901, 0.419275, 0.420786, 0.421376,
0.420933, 0.421276, 0.420543, 0.421582, 0.420259, 0.421037, 0.421247,
0.419993, 0.418218, 0.417962, 0.416763, 0.416312, 0.416108, 0.415537,
0.415438, 0.415928, 0.41681, 0.417882, 0.41772, 0.416887, 0.417119,
0.417682, 0.419768, 0.419542, 0.418273, 0.418086, 0.417408, 0.416689,
0.41684, 0.417688, 0.41845, 0.4195, 0.41933, 0.420903, 0.420708, 0.423045,
0.422215, 0.423355, 0.422808, 0.422505, 0.422567, 0.421509, 0.423147,
0.422276, 0.423217, 0.424072, 0.423056, 0.422373, 0.422435, 0.422251,
0.420751, 0.421181, 0.421156, 0.421582, 0.420499, 0.420972, 0.421641,
0.421452, 0.42245, 0.422927, 0.42246, 0.422985, 0.421919, 0.420936,
0.420135, 0.420778, 0.422456, 0.42215, 0.422136, 0.42207, 0.421529,
0.423107, 0.422322, 0.422517, 0.422622, 0.422114, 0.423033, 0.423865,
0.424108, 0.425009, 0.426093, 0.428218, 0.428399, 0.42899, 0.429583,
0.430678, 0.430875, 0.431591, 0.431123, 0.430918, 0.430341, 0.430529,
0.429364, 0.429062, 0.428099, 0.428205, 0.429184, 0.43059, 0.429181,
0.42939, 0.428952, 0.428524, 0.429531, 0.430283, 0.429799, 0.428269,
0.427569, 0.427677, 0.427546, 0.425215, 0.424981, 0.424995, 0.424171,
0.422987, 0.423081, 0.424556, 0.423675, 0.422499, 0.421702, 0.421551,
0.421183, 0.419892, 0.421325, 0.420372, 0.421525, 0.421816, 0.421889,
0.42098, 0.421993, 0.421128, 0.422599, 0.422522, 0.421787, 0.42368,
0.422023, 0.423384, 0.422553, 0.423339, 0.424524, 0.423099, 0.422917,
0.423632, 0.42293, 0.423964, 0.421808, 0.42123, 0.419817, 0.419565,
0.419248, 0.419148, 0.418127, 0.416689, 0.416525, 0.415769, 0.41617,
0.416168, 0.415486, 0.414689, 0.413361, 0.411947, 0.410319, 0.408288,
0.407708, 0.408983, 0.409904, 0.408395, 0.409292, 0.410106, 0.411314,
0.411064, 0.409352, 0.41006, 0.409222, 0.40903, 0.409113, 0.407869,
0.40757, 0.406322, 0.40618, 0.405176, 0.403975, 0.403509, 0.402268,
0.402045, 0.402782, 0.402625, 0.40132, 0.399991, 0.400104, 0.399337,
0.400894, 0.402329, 0.402859, 0.402499, 0.402488, 0.403019, 0.402867,

0.402145, 0.404173, 0.402943, 0.401978, 0.402826, 0.402159, 0.401478,
0.401207, 0.401926, 0.402653, 0.402591, 0.404333, 0.403969, 0.40648,
0.407855, 0.409688, 0.410371, 0.411244, 0.409999, 0.408351, 0.408074,
0.406957, 0.406529, 0.406896, 0.407732, 0.408478, 0.408961, 0.408532,
0.407158, 0.404993, 0.401782, 0.402089, 0.402418, 0.403088, 0.404545,
0.403609, 0.403862, 0.403768, 0.402902, 0.40463, 0.403603, 0.403447,
0.403923, 0.405678, 0.40744, 0.407013, 0.406298, 0.40553, 0.405431,
0.405371, 0.407114, 0.407989, 0.408519, 0.407083, 0.408436, 0.410285,
0.410182, 0.410682, 0.410897, 0.410768, 0.411228, 0.412057, 0.414056,
0.414166, 0.414147, 0.41621, 0.416101, 0.417044, 0.415798, 0.416106,
0.415727, 0.415394, 0.414843, 0.415489, 0.413871, 0.414752, 0.41489,
0.415892, 0.414118, 0.413595, 0.414914, 0.415475, 0.415155, 0.414149,
0.413901, 0.41382, 0.41471, 0.416254, 0.415055, 0.415715, 0.415199,
0.415324, 0.414647, 0.413236, 0.412164, 0.413213, 0.413049, 0.414056,
0.413777, 0.41246, 0.413412, 0.412662, 0.411827, 0.409702, 0.411354,
0.411974, 0.41286, 0.414636, 0.415092, 0.41624, 0.414937, 0.41502, 0.415439,
0.41688, 0.417149, 0.418337, 0.4199, 0.418723, 0.421187, 0.421701,
0.420087, 0.418795, 0.419976, 0.421635, 0.422286, 0.422548, 0.422881,
0.421265, 0.422307, 0.420164, 0.420752, 0.419708, 0.420262, 0.420165,
0.420699, 0.421009, 0.419978, 0.420028, 0.41925, 0.417943, 0.417452,
0.416384, 0.416284, 0.416091, 0.417394, 0.416773, 0.416856, 0.417638,
0.418691, 0.41929, 0.418393, 0.41822, 0.41853, 0.418016, 0.41837, 0.417205,
0.417241, 0.417632, 0.417336, 0.417257, 0.416419, 0.415393, 0.417829,
0.418249, 0.419946, 0.419529, 0.418208, 0.417954, 0.417985, 0.416792,
0.417227, 0.415025, 0.414687, 0.414188, 0.415169, 0.413998, 0.415665,
0.413887, 0.415081, 0.416054, 0.415679, 0.415494, 0.415847, 0.415384,
0.415204, 0.414794, 0.414288, 0.413701, 0.413796, 0.412602, 0.412558,
0.413952, 0.414702, 0.416267, 0.415832, 0.417641, 0.41656, 0.418075,
0.419172, 0.41984, 0.42012, 0.42053, 0.420091, 0.420071, 0.419721,
0.41749, 0.418301, 0.418883, 0.420985, 0.419567, 0.421542, 0.420204,
0.420119, 0.419854, 0.419438, 0.419471, 0.419875, 0.42047, 0.419281,
0.41962, 0.419927, 0.419598, 0.419107, 0.420394, 0.420344, 0.42116,
0.421115, 0.41881, 0.415913, 0.414619, 0.414539, 0.414423, 0.41349,
0.413497, 0.411675, 0.41286, 0.413726, 0.415798, 0.41611, 0.416817,
0.416877, 0.417704, 0.416944, 0.415941, 0.417363, 0.416549, 0.417009,
0.416294, 0.416553, 0.41603, 0.416647, 0.417355, 0.416773, 0.416928,
0.417133, 0.41768, 0.417049, 0.416214, 0.417274, 0.416786, 0.417674,
0.418195, 0.419345, 0.420362, 0.419639, 0.418095, 0.418213, 0.416459,
0.417583, 0.418113, 0.417981, 0.419274, 0.419687, 0.419031, 0.418798,
0.421944, 0.421388, 0.420855, 0.420896, 0.421375, 0.420952, 0.420271,
0.419657, 0.418992, 0.419121, 0.419796, 0.420611, 0.421998, 0.421147,
0.42157, 0.420971, 0.421575, 0.420066, 0.421429, 0.421703, 0.421873,
0.423546, 0.424205, 0.424356, 0.423946, 0.425894, 0.427183, 0.42749,
0.427762, 0.42781, 0.427029, 0.427159, 0.426924, 0.42699, 0.426564,
0.426938, 0.426852, 0.427266, 0.426458, 0.42537, 0.42541, 0.425498,
0.426062, 0.425347, 0.424733, 0.425178, 0.425281, 0.426654, 0.426059,
0.425836, 0.42665, 0.426957, 0.426511, 0.425267, 0.424144, 0.423308,

0.423829, 0.423578, 0.423284, 0.423546, 0.422618, 0.422697, 0.421417,
0.42124, 0.420923, 0.420188, 0.420697, 0.421135, 0.420744, 0.421096,
0.422124, 0.421745, 0.420399, 0.419956, 0.422254, 0.423879, 0.424164,
0.423536, 0.42435, 0.424519, 0.422171, 0.422891, 0.423897, 0.423779,
0.424281, 0.423304, 0.423128, 0.424743, 0.424161, 0.425135, 0.425818,
0.427499, 0.426482, 0.428532, 0.428581, 0.42958, 0.429811, 0.431764,
0.43125, 0.430106, 0.428363, 0.426586, 0.426407, 0.424901, 0.423073,
0.421579, 0.421976, 0.42223, 0.422238, 0.421826, 0.421858, 0.421059,
0.421486, 0.422306, 0.422187, 0.42254, 0.423273, 0.422948, 0.421328,
0.42034, 0.418969, 0.419871, 0.419616, 0.418533, 0.419019, 0.418456,
0.417911, 0.417088, 0.418206, 0.417475, 0.415427, 0.414003, 0.413706,
0.412547, 0.411188, 0.412595, 0.412574, 0.412186, 0.411188, 0.411229,
0.411455, 0.411959, 0.411353, 0.410028, 0.410034, 0.408987, 0.408418,
0.408248, 0.408491, 0.408793, 0.408962, 0.409142, 0.408162, 0.408721,
0.409452, 0.408837, 0.409057, 0.408983, 0.40952, 0.408192, 0.407132,
0.408382, 0.408028, 0.408321, 0.410186, 0.410137, 0.411553, 0.41123,
0.411748, 0.41091, 0.411769, 0.411451, 0.411662, 0.412126, 0.410422,
0.409867, 0.411511, 0.411261, 0.410673, 0.412506, 0.413206, 0.414517,
0.413635, 0.413381, 0.414741, 0.415843, 0.417741, 0.41713, 0.417944,
0.41717, 0.416546, 0.416955, 0.417829, 0.417434, 0.418584, 0.41771,
0.416769, 0.418549, 0.418311, 0.418164, 0.41826, 0.416944, 0.417508,
0.41814, 0.418626, 0.418135, 0.418598, 0.419425, 0.41895, 0.417264,
0.417136, 0.418028, 0.418638, 0.417827, 0.417763, 0.418385, 0.417247,
0.418318, 0.418612, 0.417579, 0.41784, 0.41785, 0.418117, 0.418388,
0.419114, 0.42039, 0.42144, 0.421407, 0.420816, 0.420725, 0.419974,
0.419395, 0.418132, 0.416758, 0.417035, 0.417317, 0.415635, 0.414678,
0.414361, 0.414336, 0.411566, 0.411723, 0.41257, 0.41215, 0.412335,
0.411229, 0.411182, 0.411382, 0.411368, 0.410214, 0.410276, 0.410024,
0.408496, 0.40859, 0.408182, 0.409334, 0.409272, 0.409099, 0.409227,
0.40774, 0.406831, 0.406971, 0.405923, 0.402547, 0.401984, 0.400783,
0.400623, 0.399484, 0.398985, 0.398834, 0.398198, 0.398241, 0.396058,
0.396238, 0.395829, 0.396573, 0.395146, 0.395381, 0.395666, 0.395536,
0.395853, 0.395057, 0.396396, 0.395865, 0.395605, 0.396426, 0.396187,
0.395005, 0.395288, 0.395956, 0.394152, 0.39498, 0.394676, 0.39455,
0.394228, 0.393402, 0.393634, 0.395174, 0.39596, 0.397702, 0.396881,
0.396034, 0.39482, 0.395916, 0.398007, 0.396734, 0.398803, 0.399089,
0.400191, 0.398088, 0.400086, 0.399003, 0.398035, 0.398516, 0.397117,
0.396171, 0.397233, 0.39579, 0.396353, 0.394726, 0.394077, 0.393677,
0.393177, 0.39311, 0.392568, 0.393013, 0.392905, 0.392117, 0.393081,
0.393353, 0.392396, 0.394934, 0.394373, 0.395247, 0.395207, 0.394545,
0.395473, 0.395976, 0.396117, 0.398052, 0.398422, 0.396523, 0.396394,
0.396401, 0.39503, 0.395216, 0.393306, 0.393462, 0.393383, 0.394728,
0.395552, 0.396661, 0.398917, 0.398113, 0.399706, 0.4003, 0.400988,
0.40249, 0.403338, 0.404764, 0.405144, 0.405211, 0.404647, 0.404817,
0.403677, 0.402667, 0.402221, 0.402754, 0.403837, 0.403933, 0.40278,
0.402476, 0.402931, 0.401918, 0.400201, 0.399329, 0.399056, 0.398853,
0.398972, 0.399684, 0.399674, 0.399072, 0.399899, 0.39899, 0.397192,

0.396383, 0.396025, 0.397307, 0.395571, 0.396447, 0.39667, 0.39787,
0.398008, 0.39804, 0.397975, 0.397935, 0.398426, 0.398687, 0.398412,
0.398348, 0.398968, 0.398618, 0.398645, 0.399954, 0.400001, 0.399466,
0.398651, 0.398237, 0.399359, 0.398144, 0.39873, 0.398314, 0.397804,
0.39854, 0.397707, 0.397703, 0.397449, 0.39933, 0.3993, 0.399786, 0.399524,
0.401205, 0.401344, 0.400439, 0.399752, 0.400103, 0.400774, 0.401734,
0.401207, 0.401217, 0.402494, 0.403307, 0.40255, 0.404142, 0.403915,
0.403465, 0.404025, 0.404509, 0.405659, 0.405464, 0.406617, 0.406718,
0.406519, 0.406875, 0.408377, 0.408953, 0.409367, 0.408317, 0.409251,
0.409301, 0.407165, 0.406924, 0.407695, 0.406851, 0.406605, 0.407689,
0.408208, 0.407282, 0.406891, 0.407195, 0.407634, 0.408052, 0.406912,
0.406792, 0.405866, 0.404734, 0.404769, 0.406476, 0.405832, 0.405017,
0.406146, 0.406052, 0.406966, 0.406394, 0.406983, 0.407974, 0.406773,
0.406793, 0.409154, 0.409553, 0.409316, 0.409524, 0.40908, 0.406617,
0.407037, 0.408413, 0.408831, 0.410463, 0.411669, 0.411122, 0.410269,
0.4089, 0.409321, 0.409334, 0.408385, 0.408876, 0.408951, 0.408389,
0.40957, 0.409487, 0.410127, 0.410582, 0.410076, 0.409155, 0.409947,
0.408538, 0.408947, 0.409601, 0.409668, 0.409649, 0.410202, 0.410261,
0.410218, 0.411933, 0.412186, 0.412814, 0.413154, 0.413178, 0.412495,
0.413232, 0.41373, 0.413189, 0.414625, 0.413671, 0.413365, 0.414401,
0.415502, 0.415376, 0.416818, 0.416898, 0.41881, 0.418644, 0.42001,
0.419633, 0.420347, 0.41963, 0.419072, 0.419557, 0.420742, 0.421192,
0.421313, 0.418485, 0.41893, 0.419718, 0.419877, 0.420635, 0.421682,
0.422909, 0.423519, 0.423672, 0.423511, 0.424497, 0.426352, 0.42812,
0.427361, 0.426512, 0.427368, 0.426049, 0.42571, 0.425045, 0.425865,
0.425552, 0.424478, 0.424871, 0.424526, 0.423751, 0.422922, 0.423206,
0.42293, 0.422587, 0.422637, 0.42386, 0.422848, 0.422639, 0.422701,
0.423977, 0.423066, 0.423149, 0.421328, 0.420918, 0.420429, 0.418665,
0.419298, 0.417595, 0.41644, 0.415589, 0.415361, 0.414933, 0.415418,
0.41471, 0.414381, 0.414705, 0.413219, 0.413397, 0.413537, 0.411581,
0.412029, 0.41191, 0.411818, 0.411055, 0.413201, 0.41226, 0.409597,
0.409652, 0.408331, 0.40811, 0.408593, 0.408514, 0.410438, 0.409508,
0.410147, 0.411364, 0.412443, 0.413402, 0.412835, 0.41228, 0.414116,
0.414685, 0.414998, 0.416078, 0.416046, 0.415121, 0.415041, 0.412946,
0.411733, 0.411654, 0.41207, 0.411651, 0.410583, 0.410571, 0.410299,
0.410251, 0.409991, 0.412123, 0.412058, 0.413148, 0.414231, 0.413212,
0.414081, 0.414446, 0.414848, 0.413683, 0.4129, 0.412717, 0.413692,
0.414727, 0.415871, 0.419403, 0.420142, 0.418985, 0.419832, 0.418743,
0.418323, 0.41871, 0.417894, 0.417862, 0.417907, 0.418921, 0.417722,
0.417143, 0.414943, 0.415065, 0.414841, 0.415109, 0.41465, 0.413847,
0.412749, 0.411544, 0.413591, 0.413092, 0.411824, 0.412605, 0.411516,
0.411632, 0.411763, 0.412783, 0.411351, 0.412435, 0.411321, 0.409228,
0.408341, 0.408597, 0.411105, 0.410264, 0.410719, 0.411701, 0.412336,
0.412496, 0.413774, 0.413797, 0.415504, 0.415337, 0.416155, 0.41584,
0.4149, 0.415392, 0.415685, 0.414855, 0.416421, 0.416963, 0.416372,
0.415174, 0.414543, 0.414163, 0.411931, 0.411337, 0.41088, 0.410162,
0.407783, 0.408863, 0.407405, 0.407852, 0.405774, 0.406399, 0.407627,

0.408187, 0.407147, 0.407325, 0.406601, 0.40708, 0.407541, 0.407662,
0.408849, 0.408998, 0.408618, 0.408435, 0.407863, 0.407137, 0.406623,
0.405515, 0.406354, 0.407155, 0.406483, 0.405672, 0.405173, 0.406526,
0.406524, 0.40698, 0.408329, 0.40843, 0.409128, 0.40865, 0.409472,
0.41045, 0.409824, 0.410149, 0.409264, 0.41059, 0.411467, 0.411903,
0.412661, 0.41263, 0.410707, 0.411357, 0.40965, 0.40983, 0.409544,
0.410983, 0.409944, 0.410509, 0.409917, 0.408882, 0.408022, 0.408508,
0.409738, 0.410817, 0.410636, 0.411625, 0.411546, 0.411637, 0.41258,
0.410483, 0.413121, 0.412593, 0.411494, 0.411059, 0.411857, 0.411495,
0.410143, 0.409409, 0.4065, 0.406794, 0.406243, 0.406581, 0.40704,
0.407478, 0.406835, 0.407087, 0.407108, 0.405366, 0.404694, 0.403585,
0.403683, 0.403071, 0.401716, 0.400813, 0.401317, 0.400912, 0.400182,
0.399261, 0.399382, 0.400118, 0.399411, 0.39879, 0.401191, 0.402171,
0.40283, 0.403098, 0.402552, 0.400652, 0.401842, 0.401455, 0.400259,
0.400919, 0.401114, 0.400733, 0.398618, 0.398372, 0.398086, 0.398799,
0.39981, 0.400224, 0.398809, 0.399134, 0.399523, 0.399731, 0.400015,
0.39952, 0.399535, 0.401515, 0.400904, 0.400045, 0.399472, 0.399635,
0.400672, 0.40184, 0.401092, 0.40294, 0.401791, 0.401906, 0.401288,
0.400916, 0.400713, 0.40057, 0.400962, 0.401282, 0.402889, 0.403942,
0.403498, 0.404957, 0.404714, 0.404382, 0.404437, 0.403869, 0.404038,
0.405329, 0.405201, 0.404463, 0.40345, 0.402182, 0.403514, 0.404995,
0.406053, 0.405107, 0.405634, 0.405114, 0.405354, 0.404917, 0.404467,
0.404936, 0.405078, 0.405372, 0.405993, 0.406286, 0.406566, 0.408155,
0.407245, 0.405749, 0.405313, 0.40596, 0.40777, 0.408456, 0.408332,
0.407543, 0.407106, 0.40678, 0.407484, 0.409439, 0.410627, 0.410003,
0.408292, 0.406718, 0.407155, 0.406429, 0.405034, 0.404502, 0.406726,
0.407164, 0.405764, 0.403812, 0.404324, 0.405429, 0.406284, 0.405732,
0.404694, 0.40552, 0.406306, 0.406073, 0.406374, 0.404618, 0.403579,
0.402377, 0.401855, 0.400834, 0.400412, 0.400804, 0.401876, 0.402516,
0.402568, 0.402668, 0.40258, 0.400999, 0.400581, 0.400743, 0.400435,
0.399563, 0.399919, 0.400515, 0.400361, 0.399869, 0.399479, 0.399746,
0.40014, 0.400482, 0.401083, 0.399942, 0.400466, 0.400751, 0.401361,
0.402104, 0.403265, 0.402541, 0.40153, 0.402363, 0.401963, 0.400907,
0.40213, 0.400917, 0.401853, 0.40199, 0.402842, 0.401161, 0.401122,
0.401947, 0.402112, 0.402217, 0.399645, 0.400035, 0.400144, 0.400108,
0.400402, 0.402264, 0.40303, 0.403132, 0.403997, 0.402252, 0.401837,
0.401766, 0.40147, 0.401207, 0.400725, 0.400622, 0.399889, 0.399293,
0.396582, 0.396712, 0.3955, 0.394943, 0.395136, 0.395449, 0.393975,
0.39384, 0.395933, 0.397637, 0.398647, 0.399343, 0.400144, 0.400228,
0.397785, 0.396434, 0.39821, 0.3985, 0.398397, 0.400337, 0.400802,
0.399989, 0.400519, 0.401408, 0.401065, 0.400064, 0.399362, 0.399455,
0.398926, 0.399564, 0.397924, 0.398091, 0.398374, 0.397381, 0.398602,
0.397266, 0.396616, 0.397328, 0.39708, 0.397789, 0.397053, 0.397377,
0.396305, 0.396744, 0.397663, 0.396497, 0.395762, 0.39606, 0.394663,
0.396729, 0.397305, 0.397008, 0.397672, 0.397283, 0.395167, 0.395666,
0.396021, 0.396753, 0.396781, 0.397019, 0.395657, 0.398089, 0.399518,
0.39972, 0.397827, 0.397609, 0.397548, 0.397836, 0.397561, 0.397596,

0.398298, 0.398893, 0.397244, 0.399751, 0.398651, 0.398505, 0.397685,
0.396872, 0.397369, 0.39793, 0.396737, 0.397028, 0.398191, 0.396743,
0.395655, 0.395695, 0.397774, 0.398337, 0.39853, 0.397873, 0.397178,
0.39793, 0.399043, 0.398275, 0.397649, 0.395924, 0.395819, 0.394761,
0.395027, 0.394334, 0.394899, 0.395016, 0.396769, 0.396679, 0.397321,
0.395989, 0.396541, 0.39607, 0.395714, 0.394228, 0.39532, 0.394465,
0.395703, 0.396496, 0.397738, 0.396408, 0.393893, 0.391494, 0.390419,
0.390442, 0.391031, 0.391322, 0.392266, 0.39295, 0.392986, 0.392925,
0.394628, 0.395601, 0.396993, 0.395486, 0.394964, 0.395039, 0.39848,
0.399815, 0.400595, 0.400109, 0.399274, 0.398912, 0.396913, 0.397994,
0.398379, 0.398246, 0.397073, 0.398365, 0.398035, 0.398132, 0.398725,
0.398069, 0.397738, 0.397916, 0.39775, 0.396521, 0.396194, 0.395215,
0.395393, 0.395001, 0.395131, 0.396293, 0.395025, 0.394716, 0.394696,
0.39668, 0.3961, 0.395441, 0.396485, 0.397771, 0.396714, 0.39803,
0.398766, 0.40089, 0.402459, 0.404151, 0.405444, 0.404646, 0.403739,
0.40406, 0.404747, 0.404981, 0.403613, 0.404187, 0.40457, 0.404923,
0.404432, 0.403613, 0.401469, 0.401579, 0.401718, 0.401613, 0.402568,
0.402932, 0.402233, 0.402631, 0.402742, 0.40433, 0.403034, 0.403846,
0.404356, 0.405516, 0.40739, 0.406673, 0.407817, 0.406909, 0.405875,
0.406698, 0.405606, 0.405394, 0.405918, 0.405187, 0.40558, 0.404806,
0.404103, 0.405795, 0.407364, 0.408485, 0.408498, 0.406391, 0.407125,
0.405179, 0.404403, 0.403601, 0.401678, 0.402872, 0.402675, 0.402416,
0.403218, 0.403326, 0.40342, 0.403702, 0.40268, 0.402854, 0.40299,
0.403635, 0.404568, 0.405864, 0.403793, 0.404229, 0.402937, 0.403684,
0.401849, 0.402049, 0.402306, 0.402536, 0.403571, 0.403896, 0.402586,
0.403954, 0.404477, 0.405882, 0.405583, 0.40439, 0.40321, 0.402228,
0.400349, 0.399408, 0.399784, 0.401596, 0.400598, 0.400049, 0.399512,
0.398446, 0.399845, 0.398715, 0.395896, 0.394915, 0.39654, 0.396424,
0.397057, 0.395435, 0.395608, 0.396315, 0.396242, 0.394561, 0.39576,
0.395613, 0.396148, 0.39867, 0.399819, 0.402158, 0.402153, 0.402018,
0.401047, 0.402271, 0.402853, 0.403954, 0.403669, 0.403963, 0.404207,
0.405975, 0.405312, 0.406489, 0.408085, 0.409309, 0.408978, 0.409093,
0.410075, 0.409431, 0.409585, 0.409293, 0.408922, 0.407432, 0.407487,
0.406525, 0.406945, 0.407655, 0.407972, 0.407103, 0.407152, 0.406743,
0.407643, 0.407663, 0.408068, 0.406637, 0.405236, 0.406397, 0.405576,
0.404647, 0.405171, 0.405348, 0.403363, 0.400666, 0.398411, 0.399039,
0.398186, 0.396764, 0.396411, 0.39644, 0.395112, 0.394816, 0.395448,
0.395051, 0.396089, 0.396242, 0.398004, 0.399056, 0.400446, 0.402114,
0.400716, 0.40025, 0.400618, 0.400958, 0.403265, 0.402374, 0.40314,
0.402789, 0.401043, 0.402132, 0.402681, 0.403138, 0.40279, 0.402091,
0.402448, 0.4017, 0.401912, 0.402052, 0.400701, 0.401106, 0.400967,
0.39991, 0.39944, 0.399403, 0.399449, 0.39942, 0.398072, 0.399981,
0.400299, 0.400242, 0.399852, 0.399106, 0.398062, 0.401136, 0.399904,
0.399444, 0.40057, 0.400036, 0.401451, 0.401074, 0.400568, 0.399881,
0.39829, 0.39949, 0.398527, 0.399341, 0.3984, 0.39811, 0.399442, 0.398846,
0.399602, 0.40132, 0.400693, 0.401238, 0.401326, 0.40226, 0.401719,
0.399763, 0.399035, 0.397886, 0.396699, 0.395909, 0.396908, 0.397786,

```
0.398737, 0.397705, 0.397156, 0.396222, 0.395688, 0.394733, 0.395543,
0.396304, 0.397219, 0.39793, 0.398096, 0.395529, 0.394113, 0.391499,
0.392666, 0.39104, 0.391345, 0.394344, 0.393719, 0.391253, 0.393426,
0.391103, 0.39104, 0.390643, 0.391418, 0.390531, 0.388816, 0.388369,
0.389513, 0.387808, 0.388627, 0.389227, 0.388516, 0.386569, 0.386342,
0.387529, 0.387836, 0.387104, 0.386815, 0.386909, 0.386449, 0.38871,
0.388762, 0.388096, 0.389642, 0.38889, 0.389425, 0.387942, 0.389448,
0.388412, 0.389701, 0.391842, 0.39288, 0.392844, 0.39172, 0.393084,
0.393979, 0.394851, 0.39422, 0.395198, 0.396084, 0.39637, 0.396546,
0.398052, 0.397534, 0.397868, 0.398007, 0.396525, 0.396182, 0.396158,
0.398203, 0.39904, 0.400772, 0.401206, 0.400272, 0.400112, 0.40239,
0.404987, 0.403981, 0.404191, 0.405126, 0.403569, 0.404431, 0.403608,
0.403571, 0.403191, 0.403445, 0.405009, 0.405262, 0.405255, 0.405703,
0.4045, 0.403987, 0.401509, 0.400154, 0.400831, 0.401953, 0.400702,
0.399259, 0.400172, 0.402617, 0.402804, 0.399889, 0.399929, 0.400052,
0.401274, 0.404125, 0.404305, 0.404042, 0.402443, 0.402346, 0.40395,
0.404395, 0.405328, 0.40542, 0.404294, 0.406079, 0.406164, 0.40628,
0.406701, 0.407184, 0.407071, 0.405898, 0.403757, 0.405099, 0.405931,
0.404906, 0.405507, 0.402773, 0.402824, 0.402494, 0.400046, 0.399416,
0.400479, 0.400832, 0.401926, 0.401992, 0.400066, 0.401034, 0.401521,
0.402859, 0.401151, 0.401925, 0.40138, 0.402275, 0.400761, 0.400641,
0.39946, 0.400116, 0.399105, 0.400246, 0.401648, 0.401391, 0.402694,
0.403242, 0.400515, 0.398454, 0.398289, 0.399761, 0.399951, 0.399222,
0.398596, 0.397239, 0.397389, 0.398614, 0.398546, 0.395639, 0.393672,
0.393709, 0.393713, 0.392581, 0.390432, 0.391334, 0.391468, 0.393169,
0.393024, 0.392205, 0.39226, 0.392279, 0.390432, 0.387923, 0.390149,
0.39187, 0.390947, 0.391201, 0.390081, 0.390669, 0.392318, 0.390807,
0.390567, 0.392373, 0.392597, 0.39281, 0.392936, 0.393739, 0.394828,
0.392963, 0.392136, 0.392633, 0.393219, 0.394501, 0.393578, 0.393699,
0.393088, 0.392386, 0.393076, 0.391583, 0.392374, 0.392733, 0.39394,
0.393581, 0.394399, 0.39503, 0.395389, 0.395673, 0.395234, 0.393482,
0.394046, 0.393378, 0.394744, 0.396283, 0.395735, 0.394438, 0.394315,
0.395977, 0.396141, 0.395885, 0.397473, 0.398348, 0.397236, 0.396418,
0.397699, 0.399328, 0.39818, 0.399727, 0.399808, 0.399624, 0.39984,
0.398854, 0.398642, 0.399413, 0.39879, 0.399928, 0.400769, 0.401074,
0.400642, 0.398775, 0.398258, 0.397885, 0.399098, 0.398104, 0.3965,
0.398071, 0.398872, 0.397934, 0.397528, 0.398191, 0.398495, 0.39974,
0.399191, 0.401414, 0.401481, 0.399701, 0.398342, 0.397475, 0.399755}
```

In[ ]:=
```
threadJoin = Quiet[Re@##] /. Re → List &;
(*Defines a useful function for elementwise join of two arrays*)

uTemp1 = threadJoin[urTemp1, usTemp1, Range[Length[usTemp1]]];
uTemp2 = threadJoin[urTemp2, usTemp2, Range[Length[usTemp2]]];
uTemp3 = threadJoin[urTemp3, usTemp3, Range[Length[usTemp3]]];
uTemp4 = threadJoin[urTemp4, usTemp4, Range[Length[usTemp4]]];
```

In[ ]:=
```
(*PutAppend[uTemp1,dataStream];*)
```

In[ ]:=
```
ListPlot[{uTemp1⟦All, {1, 2}⟧,
  uTemp2⟦All, {1, 2}⟧, uTemp3⟦All, {1, 2}⟧, uTemp4⟦All, {1, 2}⟧},
 PlotRange → {{0.05, 0.9}, {0.1, 0.9}}, Joined → True, ColorFunction → "Rainbow"]
```

Out[ ]=

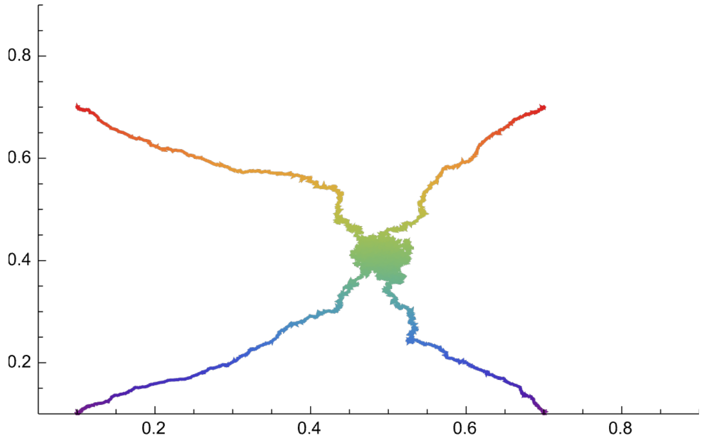


#### 1.4. Simulation results

In[ ]:=
```
(*The Result in the following format: array of vectors of results res(μb)=
 {μb, ur, us, nu, simulTime(sec)} for diffrent values of the baseline
  mutation rate μb and where the first element of the results vector
  indicated the current value of the baseline mutation rate μb *)
```

##### 1.4.1. Results ($\alpha = 0.02$)

```
In[ ]:= dataSimul1 =
  {{0.02`, 0.2849779022839121`, 0.5048810222998984`,
    0.05160171965606885`, 996.003999`9.449806075634681},
   {0.06`, 0.5813207826219774`, 0.46733173432893177`, 0.05386858628274335`,
    2010.807486`9.754915486882348}, {0.1`, 0.6809779393186428`,
    0.46149113019361176`, 0.05236212757448487`, 3110.338816`9.944352693746099},
   {0.15`, 0.7509533324887845`, 0.48794275483200183`, 0.0482923415316935`,
    4085.873184`10.062829876511023}, {0.2`, 0.772107170209215`,
    0.4713514115641462`, 0.053857868426314556`, 5102.01964`10.15928711957045},
   {0.25`, 0.7978991016745477`, 0.49066865962920053`, 0.053423875224954856`,
    6146.217929`10.240152948649039}, {0.3`, 0.8127989844307917`,
    0.45034109170055986`, 0.05542943411317732`, 7278.832768`10.31360673502611},
   {0.35`, 0.8289972735780193`, 0.4812894605178182`, 0.05395924815036964`,
    8357.00564`10.373595688761156}, {0.4`, 0.8349619263545627`,
    0.44326846546091947`, 0.05709438112377531`, 9333.262758`10.42157848612635},
   {0.45`, 0.8435376161366338`, 0.4775049009577848`,
    0.05796404719056182`, 10312.032287`10.464889257620605},
   {0.5`, 0.8583137380942792`, 0.49079552563434703`,
    0.05363495300939796`, 11289.000341`10.50420047965397}};

In[ ]:=

In[ ]:= urSimul1 = dataSimul1⟦All, {1, 2}⟧;

In[ ]:= urSimul1Inter = Interpolation[urSimul1];

In[ ]:= usSimul1 = dataSimul1⟦All, {1, 3}⟧;

In[ ]:= usSimul1Inter = Interpolation[urSimul1];

In[ ]:= laSimul1 = dataSimul1⟦All, {1, 4}⟧;

In[ ]:= laSimul1Inter = Interpolation[urSimul1];

(*Computes mutation rate from the data*)

threadJoin = Quiet[Re@##] /. Re → List &;
(*Defines a useful function for elementwise join of two arrays*)
muTemp1 = Flatten[urSimul1⟦All, {1}⟧ * (1 - urSimul1⟦All, {2}⟧)^2 ]
  (* μ=μb*(1-ur) *);
muSimul1 = threadJoin[Flatten[urSimul1⟦All, {1}⟧], muTemp1];
```

#### 1.4.2. Results ($\alpha$= 0.1)

```
In[•]:= dataSimul2 =
  {{0.02`, 0.018158426518732035`, 0.44652978429218126`, 0.096603679264147`,
    965.36382`9.436236011752426}, {0.06`, 0.08073552458182442`,
    0.43205071326590333`, 0.2532807438512292`, 3266.504107`9.965628202063844},
   {0.1`, 0.304345676473806`, 0.4201322276931529`, 0.24336032793441253`,
    4536.083242`10.108226009224202}, {0.15`, 0.4182326757536784`,
    0.4186310653787863`, 0.25599492101579574`, 5505.31057`10.192326817263822},
   {0.2`, 0.5082921437801758`, 0.4204231928000513`, 0.2434567886422709`,
    6475.362863`10.262809103698386}, {0.25`, 0.5566675237777752`,
    0.4033993197465713`, 0.2483994001199761`, 7445.713198`10.323451297077934},
   {0.3`, 0.5883672555043401`, 0.40251524139341166`, 0.256688662267545`,
    8415.85455`10.376643214521625}, {0.35`, 0.619348725041838`,
    0.4008517789150927`, 0.256720855828833`, 9386.945195`10.42406927577234},
   {0.4`, 0.6429326854394969`, 0.38371846212355637`,
    0.2586430313937221`, 10355.570909`10.466719040307641},
   {0.45`, 0.6623101144214092`, 0.38663491667117367`,
    0.2595608478304328`, 11341.969242`10.506233458695311},
   {0.5`, 0.6820635524755411`, 0.39074459889486124`,
    0.2548461507698449`, 12328.281137`10.542447523071974}};

In[•]:=

In[•]:= urSimul2 = dataSimul2[[All, {1, 2}]];

In[•]:= urSimul2Inter = Interpolation[urSimul2];

In[•]:= usSimul2 = dataSimul2[[All, {1, 3}]];

In[•]:= usSimul2Inter = Interpolation[urSimul2];

In[•]:= laSimul2 = dataSimul2[[All, {1, 4}]];

In[•]:= laSimul2Inter = Interpolation[urSimul2];

In[•]:= (*Computes mutation rate from the data*)
threadJoin = Quiet[Re@##] /. Re → List &;
muTemp2 = Flatten[urSimul2[[All, {1}]] * (1 - urSimul2[[All, {2}]])^2]
  (* μ=μb*(1-ur) *);
muSimul2 = threadJoin[Flatten[urSimul2[[All, {1}]]], muTemp2];
```

##### 1.4.3. Results ($\alpha = 0.2$)

```
In[ ]:= dataSimul3 =
  {{0.02`, 0.011603402121799855`, 0.42232840139154815`, 0.09820875824834975`,
    865.40578`9.388764784998525}, {0.06`, 0.018399001209311455`,
    0.4054102634386388`, 0.29003511297740436`, 1732.604231`9.690244364102481},
   {0.1`, 0.029321353217799953`, 0.3733725024705082`, 0.47142839432113504`,
    6167.69421`10.241667827053513}, {0.15`, 0.17919754067519786`,
    0.3483281945217647`, 0.5074791441711632`, 7061.067312`10.300415345068759},
   {0.2`, 0.29510970552615606`, 0.3396631588606742`, 0.49896740651869465`,
    7939.052011`10.351313640637022}, {0.25`, 0.367643017121403`,
    0.32752097783030903`, 0.5016513097380487`, 8826.40903`10.397329042978754},
   {0.3`, 0.4220098601039076`, 0.3270279481897332`, 0.5037566486702658`,
    9709.25819`10.438731043558834}, {0.35`, 0.46067916619893634`,
    0.3301424961232047`, 0.5121071785642851`, 10594.625843`10.476630617384718},
   {0.4`, 0.4972929460778841`, 0.3130897258788491`, 0.509563687262544`,
    11477.585936`10.51139554663439}, {0.45`, 0.5215893266194344`,
    0.3019888129238775`, 0.5180496700659856`, 12347.731149`10.543132158379818},
   {0.5`, 0.5508190531584227`, 0.3113914312327404`,
    0.5072319936012782`, 13225.346023`10.572952037070008}};

In[ ]:=

In[ ]:=

In[ ]:= urSimul3 = dataSimul3[[All, {1, 2}]];

In[ ]:= urSimul3Inter = Interpolation[urSimul3];

In[ ]:= usSimul3 = dataSimul3[[All, {1, 3}]];

In[ ]:= usSimul3Inter = Interpolation[urSimul3];

In[ ]:= laSimul3 = dataSimul3[[All, {1, 4}]];

In[ ]:= laSimul3Inter = Interpolation[urSimul3];

In[ ]:= (*Computes mutation rate from the data*)
threadJoin = Quiet[Re@##] /. Re → List &;
muTemp3 = Flatten[urSimul3[[All, {1}]] * (1 - urSimul3[[All, {2}]])^2 ]
  (* μ=μb*(1-ur) *);
muSimul3 = threadJoin[Flatten[urSimul3[[All, {1}]]], muTemp3];
```

### 1.4b. Simulation results (with standard deviations)

In[ ]:=
```
(*The Result in the following format: array of vectors of results res(μb) =
 {μb, mean(ur), std(ur), mean(us), std(us), mean(nu), std(nu),
   simulTime(sec)} for diffrent values of the baseline mutation
  rate μb and where the first element of the results vector
  indicated the current value of the baseline mutation rate μb *)
```

#### 1.4.1. Results ($\alpha = 0.02$)

In[ ]:=
```
dataSimul1 =
  {{0.02`, 0.2653341225987212`, 0.0007770630022012713`,
    0.4830919423889296`, 0.0003917507790959075`, 0.05662908945473924`,
    0.00003475628730752835`, 1064.134145`9.478541372170307},
   {0.06`, 0.5746822148271555`, 0.0002492079154426532`,
    0.49246265307039266`, 0.0005146003458797949`, 0.05955092209927547`,
    0.00003828204202577005`, 2148.302833`9.783640494653248},
   {0.1`, 0.6580550771321518`, 0.0000854772808039606`,
    0.4663722378229636`, 0.0006619412179353601`, 0.06470628805048208`,
    0.00003318201130458127`, 3129.211023`9.946979844924126},
   {0.15`, 0.722394258993336`, 0.00009479663231630993`,
    0.49536572884522634`, 0.0007272537114948919`, 0.0649294760698573`,
    0.00003986708715735241`, 4107.176083`10.065088315837324},
   {0.2`, 0.7603028986334707`, 0.0001016406904867358`,
    0.48212381432904444`, 0.0004797737421529596`, 0.06567975825445535`,
    0.00004486176766267906`, 5086.534462`10.15796698474707},
   {0.25`, 0.7796772648964672`, 0.00005817776457049837`,
    0.47945827075633946`, 0.0005370050710096884`, 0.06889390747900295`,
    0.00004230238562697686`, 6065.176356`10.234388426781686},
   {0.3`, 0.799202802099558`, 0.00004289783018846616`,
    0.47331198465970686`, 0.0004843832173849361`, 0.06921823756832395`,
    0.00004240854183754868`, 7043.545815`10.299336337327455},
   {0.35`, 0.8112326135143108`, 0.00004070671336973548`,
    0.46909746682573095`, 0.0008486560612485168`, 0.0716911878416212`,
    0.00004346745485923293`, 8023.748773`10.355922315775551},
   {0.4`, 0.8227664775709478`, 0.0000428447441325412`,
    0.47246826831487704`, 0.0005645903625260015`, 0.07245270408389995`,
    0.00004650651374134637`, 9002.405579`10.405903568500774},
   {0.45`, 0.832491363761243`, 0.00003080404498575562`,
    0.47329896466338467`, 0.0008042114020419718`, 0.07309948006932407`,
    0.00004703322234795706`, 9981.702709`10.45074962436734},
   {0.5`, 0.8381325235776502`, 0.00003360976946557548`,
    0.4695824958753394`, 0.0008688074811255905`, 0.0758725325512149`,
    0.00005648642576634448`, 10962.499876`10.491454594957641}};
```

In[ ]:=
```
urSimul1 = dataSimul1[[All, {1, 2}]];
```

```
In[ ]:= urSimul1Inter = Interpolation[urSimul1];

In[ ]:= urErr1 = ScientificForm[dataSimul1[[All, 3]]]
Out[ ]//ScientificForm=
        {7.77063 × 10^-4, 2.49208 × 10^-4, 8.54773 × 10^-5,
         9.47966 × 10^-5, 1.01641 × 10^-4, 5.81778 × 10^-5, 4.28978 × 10^-5,
         4.07067 × 10^-5, 4.28447 × 10^-5, 3.0804 × 10^-5, 3.36098 × 10^-5}

In[ ]:= usSimul1 = dataSimul1[[All, {1, 4}]];

In[ ]:= usErr1 = ScientificForm[dataSimul1[[All, 5]]]
Out[ ]//ScientificForm=
        {3.91751 × 10^-4, 5.146 × 10^-4, 6.61941 × 10^-4,
         7.27254 × 10^-4, 4.79774 × 10^-4, 5.37005 × 10^-4, 4.84383 × 10^-4,
         8.48656 × 10^-4, 5.6459 × 10^-4, 8.04211 × 10^-4, 8.68807 × 10^-4}

In[ ]:= urWithErr1 =
          Transpose[{dataSimul1[[All, 1]], dataSimul1[[All, 2]], dataSimul1[[All, 3]]}];

In[ ]:= usWithErr1 =
          Transpose[{dataSimul1[[All, 1]], dataSimul1[[All, 4]], dataSimul1[[All, 5]]}];

In[ ]:= usSimul1Inter = Interpolation[urSimul1];

In[ ]:= laSimul1 = dataSimul1[[All, {1, 6}]];


In[ ]:= laErr1 = ScientificForm[dataSimul1[[All, 7]]]
Out[ ]//ScientificForm=
        {3.47563 × 10^-5, 3.8282 × 10^-5, 3.3182 × 10^-5,
         3.98671 × 10^-5, 4.48618 × 10^-5, 4.23024 × 10^-5, 4.24085 × 10^-5,
         4.34675 × 10^-5, 4.65065 × 10^-5, 4.70332 × 10^-5, 5.64864 × 10^-5}

In[ ]:= laWithErr1 =
          Transpose[{dataSimul1[[All, 1]], dataSimul1[[All, 6]], dataSimul1[[All, 7]]}];

In[ ]:= laSimul1Inter = Interpolation[urSimul1];

        (*Computes mutation rate from the data*)

        threadJoin = Quiet[Re@##] /. Re → List &;
        (*Defines a useful function for elementwise join of two arrays*)
        muTemp1 = Flatten[urSimul1[[All, {1}]] * (1 - urSimul1[[All, {2}]])^2 ]
         (* μ=μb*(1-ur) *);
        muSimul1 = threadJoin[Flatten[urSimul1[[All, {1}]]], muTemp1];
```

##### 1.4.2. Results ($\alpha$= 0.1)

In[ ]:=
```
dataSimul2 =
  {{0.02`, 0.035023861396192305`, 0.000030298398560145486`,
    0.4490901500407161`, 0.00017993112127637407`, 0.09349032573434611`,
    0.00004733976903201299`, 1053.164199`9.474041080883085},
   {0.06`, 0.09898169304054275`, 0.00009810183623506608`,
    0.4414121021734595`, 0.00018368325701149802`, 0.24630852775185386`,
    0.00011587586677166762`, 2137.534589`9.78145814458755},
   {0.1`, 0.2793887567795987`, 0.00017963920506485658`,
    0.41767903569972614`, 0.00020731264859722334`, 0.2666235257521204`,
    0.00019600410727122804`, 3127.030166`9.946677064334096},
   {0.15`, 0.41096438023567694`, 0.00009960367510427825`,
    0.41325868020750794`, 0.00022046621208444517`, 0.267632120161756`,
    0.00016866470093965575`, 4111.261607`10.065520106097708},
   {0.2`, 0.4870413976112483`, 0.0001155641032800947`,
    0.4168117397402329`, 0.00022242118991339509`, 0.27167012398346746`,
    0.00022787012933846024`, 5095.709685`10.158749670712975},
   {0.25`, 0.5398076993485078`, 0.00008128848526691243`,
    0.4082984624107034`, 0.00022864189776327 73`, 0.2746920321734873`,
    0.0002054340122711202`, 6079.141344`10.235387234627682},
   {0.3`, 0.5805414170299812`, 0.000067068031250471 21`,
    0.4044627631519853`, 0.00014661745384869407`, 0.27419734257654593`,
    0.0002128597349900427`, 7061.825331`10.30046196490473},
   {0.35`, 0.6108548312650953`, 0.0000653159548201411`,
    0.3956143158871077`, 0.00010573378986410787`, 0.2752995600586573`,
    0.00021403736869802029`, 8047.278505`10.357194025404718},
   {0.4`, 0.6354806879015831`, 0.00005043561088532316`,
    0.3998019465441064`, 0.00020314252137866877`, 0.27679066791094586`,
    0.00020977655606353789`, 9031.316332`10.407296047692148},
   {0.45`, 0.6560546132835193`, 0.000046329917358847505`,
    0.38384340015739177`, 0.00013047347606998469`, 0.27760632804514657`,
    0.0002244171815077906`, 10016.201251`10.452248035532527},
   {0.5`, 0.6721674114073624`, 0.000045409371526099 04`,
    0.3966672374111581`, 0.0002014659359904439`, 0.2806113140470149`,
    0.00022459278192695894`, 11002.189628`10.493024119447323}};
```

In[ ]:=
```
urSimul2 = dataSimul2[[All, {1, 2}]];
```

In[ ]:=
```
urSimul2Inter = Interpolation[urSimul2];
```

In[ ]:=
```
urErr2 = ScientificForm[dataSimul2[[All, 3]]]
```

Out[ ]//ScientificForm=

$\{3.02984\times10^{-5}, 9.81018\times10^{-5}, 1.79639\times10^{-4}, 9.96037\times10^{-5}, 1.15564\times10^{-4}, 8.12885\times10^{-5}, 6.7068\times10^{-5}, 6.5316\times10^{-5}, 5.04356\times10^{-5}, 4.63299\times10^{-5}, 4.54094\times10^{-5}\}$

In[ ]:=
```
usSimul2 = dataSimul2[[All, {1, 4}]];
```

```
In[ ]:= usSimul2Inter = Interpolation[urSimul2];

In[ ]:= usErr2 = ScientificForm[dataSimul2[[All, 5]]]
Out[ ]//ScientificForm=
        {1.79931 × 10^-4, 1.83683 × 10^-4, 2.07313 × 10^-4,
         2.20466 × 10^-4, 2.22421 × 10^-4, 2.28642 × 10^-4, 1.46617 × 10^-4,
         1.05734 × 10^-4, 2.03143 × 10^-4, 1.30473 × 10^-4, 2.01466 × 10^-4}

In[ ]:= urWithErr2 =
          Transpose[{dataSimul2[[All, 1]], dataSimul2[[All, 2]], dataSimul2[[All, 3]]}];

In[ ]:= usWithErr2 =
          Transpose[{dataSimul2[[All, 1]], dataSimul2[[All, 4]], dataSimul2[[All, 5]]}];

In[ ]:= laSimul2 = dataSimul2[[All, {1, 6}]];

In[ ]:= laErr2 = ScientificForm[dataSimul2[[All, 7]]]
Out[ ]//ScientificForm=
        {4.73398 × 10^-5, 1.15876 × 10^-4, 1.96004 × 10^-4,
         1.68665 × 10^-4, 2.2787 × 10^-4, 2.05434 × 10^-4, 2.1286 × 10^-4,
         2.14037 × 10^-4, 2.09777 × 10^-4, 2.24417 × 10^-4, 2.24593 × 10^-4}

In[ ]:= laWithErr2 =
          Transpose[{dataSimul2[[All, 1]], dataSimul2[[All, 6]], dataSimul2[[All, 7]]}];

In[ ]:= laSimul2Inter = Interpolation[urSimul2];

In[ ]:= (*Computes mutation rate from the data*)
        threadJoin = Quiet[Re@##] /. Re → List &;
        muTemp2 = Flatten[urSimul2[[All, {1}]] * (1 - urSimul2[[All, {2}]])^2 ]
          (* μ=μb*(1-ur) *);
        muSimul2 = threadJoin[Flatten[urSimul2[[All, {1}]]], muTemp2];
```

#### 1.4.3. Results ($\alpha = 0.2$)

```
In[∘]:= dataSimul3 =
  {{0.02`, 0.01850159210346847`, 5.460209009046523`*^-6,
    0.4162429684425838`, 0.00012779377569777412`, 0.09714317202150798`,
    0.00009756732763289809`, 1042.778051`9.469736874805172},
   {0.06`, 0.025523184392661043`, 0.000018240234775526184`,
    0.39587195320552976`, 0.00013663748873111753`, 0.2865639070346181`,
    0.0001277350806026791`, 2114.690441`9.776791795653832},
   {0.1`, 0.04521825095917123`, 0.0000575673356796422`,
    0.37813398612278365`, 0.00012827683919801154`, 0.45868872594765014`,
    0.00022596490846476958`, 3100.370195`9.942958546700208},
   {0.15`, 0.16329054017033567`, 0.00018693472859462163`,
    0.35752183148272265`, 0.0001176421175685025`, 0.5323922321468199`,
    0.0005275825008858013`, 4092.485806`10.063532175315673},
   {0.2`, 0.27989282452084635`, 0.00011692430290430947`,
    0.34877866649033845`, 0.00015465752709529077`, 0.526387734968678`,
    0.00046000844671725805`, 5088.352082000000000001`10.158122147626022},
   {0.25`, 0.35363972615746725`, 0.0001422808729151391 8`,
    0.3350506467197765`, 0.0000981669468908208`, 0.5319256099186745`,
    0.000611235270686678`, 6083.346906`10.235687576532259},
   {0.3`, 0.4092253948480109`, 0.000077262248693253 28`,
    0.33008024022180427`, 0.0000895615447537377`, 0.5321908012265028`,
    0.00045527476032020315`, 7076.662101`10.301373452747615},
   {0.35`, 0.45479101338314315`, 0.00009473179606817591`,
    0.3229888085362869`, 0.00012669519359186486`, 0.5291196373816804`,
    0.0005670553379583534`, 8072.532844`10.358554814164295},
   {0.4`, 0.49064368942921993`, 0.00005014961389256292`,
    0.3170319269366864`, 0.0001605326962097691`, 0.5290798560191959`,
    0.0003945592580779918 7`, 9065.286054`10.408926506253332},
   {0.45`, 0.5164552921596441`, 0.000050574893448722646`,
    0.314024498167354`, 0.00010985768840953844`, 0.5369244100786534`,
    0.00043370221242543944`, 10058.937044`10.454097083528444},
   {0.5`, 0.5415978028457531`, 0.00007015345552123 85`,
    0.3138082657067113`, 0.00008000329253589138`, 0.5373403723947897`,
    0.000632025343858712`, 11054.024414`10.495065412928792}};

In[∘]:= urSimul3 = dataSimul3⟦All, {1, 2}⟧;

In[∘]:= urSimul3Inter = Interpolation[urSimul3];

In[∘]:= usSimul3 = dataSimul3⟦All, {1, 4}⟧;

In[∘]:= usSimul3Inter = Interpolation[urSimul3];

In[∘]:= urWithErr3 =
  Transpose[{dataSimul3⟦All, 1⟧, dataSimul3⟦All, 2⟧, dataSimul3⟦All, 3⟧}];

In[∘]:= usWithErr3 =
  Transpose[{dataSimul3⟦All, 1⟧, dataSimul3⟦All, 4⟧, dataSimul3⟦All, 5⟧}];
```

```
In[ ]:= laSimul3 = dataSimul3[[All, {1, 6}]];

In[ ]:= laWithErr3 =
          Transpose[{dataSimul3[[All, 1]], dataSimul3[[All, 6]], dataSimul3[[All, 7]]}];

In[ ]:= laSimul3Inter = Interpolation[urSimul3];

In[ ]:= (*Computes mutation rate from the data*)
        threadJoin = Quiet[Re@##] /. Re → List &;
        muTemp3 = Flatten[urSimul3[[All, {1}]] * (1 - urSimul3[[All, {2}]])^2]
          (* μ=μb*(1-ur) *);
        muSimul3 = threadJoin[Flatten[urSimul3[[All, {1}]]], muTemp3];

In[ ]:= urErr3 = dataSimul3[[All, 3]]
Out[ ]= {5.46021 × 10^-6, 0.0000182402, 0.0000575673, 0.000186935, 0.000116924, 0.000142281,
         0.0000772622, 0.0000947318, 0.0000501496, 0.0000505749, 0.0000701535}

In[ ]:= urErr3 = ScientificForm[dataSimul3[[All, 3]]]
Out[ ]//ScientificForm=
        {5.46021 × 10^-6, 1.82402 × 10^-5, 5.75673 × 10^-5,
         1.86935 × 10^-4, 1.16924 × 10^-4, 1.42281 × 10^-4, 7.72622 × 10^-5,
         9.47318 × 10^-5, 5.01496 × 10^-5, 5.05749 × 10^-5, 7.01535 × 10^-5}

In[ ]:= usErr3 = ScientificForm[dataSimul3[[All, 5]]]
Out[ ]//ScientificForm=
        {1.27794 × 10^-4, 1.36637 × 10^-4, 1.28277 × 10^-4,
         1.17642 × 10^-4, 1.54658 × 10^-4, 9.81669 × 10^-5, 8.95615 × 10^-5,
         1.26695 × 10^-4, 1.60533 × 10^-4, 1.09858 × 10^-4, 8.00033 × 10^-5}

In[ ]:= laErr3 = ScientificForm[dataSimul3[[All, 7]]]
Out[ ]//ScientificForm=
        {9.75673 × 10^-5, 1.27735 × 10^-4, 2.25965 × 10^-4,
         5.27583 × 10^-4, 4.60008 × 10^-4, 6.11235 × 10^-4, 4.55275 × 10^-4,
         5.67055 × 10^-4, 3.94559 × 10^-4, 4.33702 × 10^-4, 6.32025 × 10^-4}
```

### 1.5. Graphical illustrations (Fig. 2 of the main text)

```
In[ ]:= (*Theoretical predictions*)
```

In[ ]:=
```
urSin[μb_, α_ , αμ_] := 1 - (α/(αμ μb))^(1/αμ)
usSin[μb_, α_ , αμ_, sb_] :=
 Piecewise[{{(e^μb/sb)^(1/(-1+α)), μb < α/αμ}, {((e^(α/αμ) * (α/(αμ μb))^(-α/αμ))/sb)^(1/(α-1)), μb > α/αμ}}]



(*Uninvadable mutation rate*)
mut[μb_, α_ , αμ_] = μb * (1 - Clip[urSin[μb, α, αμ], {0, 1}])^2 // FullSimplify;

(*Average number of novel mutations*)
laSin[α_ , αμ_, sf_] = mut[μb, α , αμ]/sf;

(*Allocation to survival according to classical life-history results*)

usSinClassic[sb_, α_ ] := (1/sb)^(1/(-1+α))
```

In[ ]:=

##### 1.5.1. Fig 2 in the main text

In[∘]:=

```
Show[Plot[{Clip[urSin[x, 0.02, 2], {0, 1}],
    Clip[urSin[x, 0.1, 2], {0, 1}], Clip[urSin[x, 0.2, 2], {0, 1}]}, {x, 0, 0.515},
  PlotStyle → {{Black, Thickness[0.012]}, {Red, Thickness[0.012]},
    {Orange, Thickness[0.012]}}, PlotRange → {{0, 0.525}, {-0.075, 1}},
  PlotLegends → Placed[{Style["α=0.02", 16, FontFamily → "Times New Roman"],
     Style["α=0.1", 16, FontFamily → "Times New Roman"],
     Style["α=0.2", 16, FontFamily → "Times New Roman"]}, {0.75, 0.25}]
 ],
 ListPlot[urSimul1, PlotMarkers → {Graphics[{Black, Disk[]}], 0.055}],
 ListPlot[urSimul2, PlotMarkers → {Graphics[{Red, Disk[]}], 0.055}],
 ListPlot[urSimul3, PlotMarkers → {Graphics[{Orange, Disk[]}], 0.055}],
 Frame → True,
 FrameLabel → {Style["Baseline mutation rate, μb"],
   Style["Uninvadable investment into repair, ug*"]},
 BaseStyle → {FontFamily → "Times New Roman", 14},
 LabelStyle → {FontFamily → "Times New Roman", 14}, AspectRatio → 0.85]
```

Out[∘]=

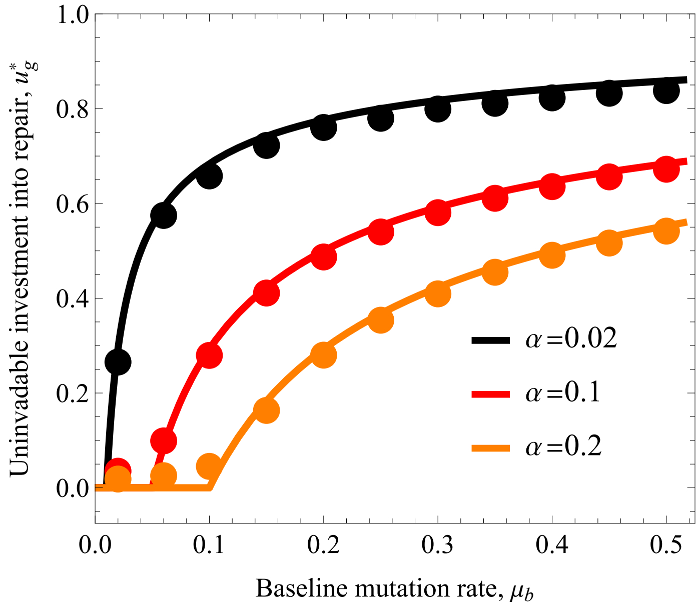

In[ ]:=

```
Show[Plot[{Clip[usSin[x, 0.02, 2, 0.5], {0, 1}],
    Clip[usSin[x, 0.1, 2, 0.5], {0, 1}], Clip[usSin[x, 0.2, 2, 0.5], {0, 1}]},
   {x, 0, 0.515}, PlotStyle → {{Black, Thickness[0.012]},
     {Red, Thickness[0.012]}, {Orange, Thickness[0.012]}},
   PlotRange → {{0, 0.525}, {-0.075, 1.0}}],
  ListPlot[usSimul1, PlotMarkers → {Graphics[{Black, Disk[]}], 0.055}],
  ListPlot[usSimul2, PlotMarkers → {Graphics[{Red, Disk[]}], 0.055}],
  ListPlot[usSimul3, PlotMarkers → {Graphics[{Orange, Disk[]}], 0.055}],
  Frame → True,
  FrameLabel → {Style["Baseline mutation rate, μb"],
    Style["Uninvadable investment into survival, u*s"]},
  BaseStyle → {FontFamily → "Times New Roman", 14},
  LabelStyle → {FontFamily → "Times New Roman", 14}, AspectRatio → 0.85]
```

Out[ ]=

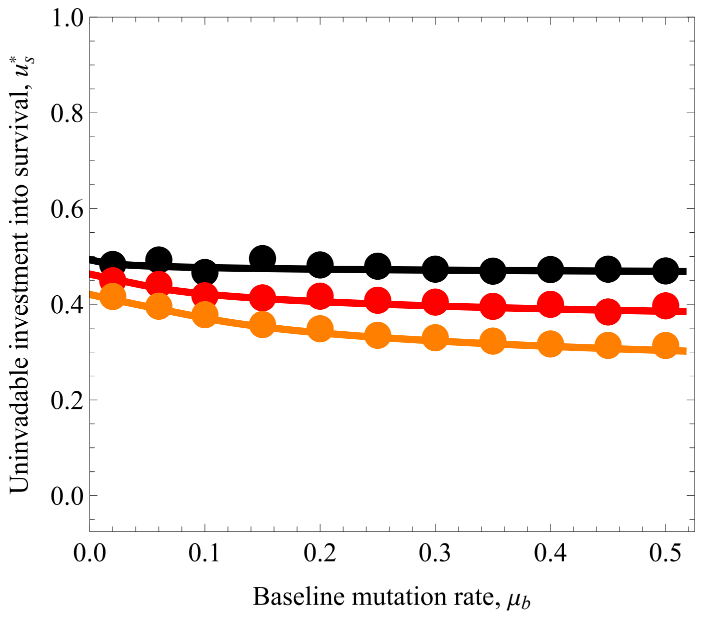

In[ ]:=

```
Show[Plot[{laSin[ 0.02, 2, 0.2], laSin[0.1, 2, 0.2], laSin[ 0.2, 2, 0.2]},
  {μb, 0, 0.515}, PlotStyle → {{Black, Thickness[0.012]}, {Red, Thickness[0.012]},
    {Orange, Thickness[0.012]}}, PlotRange → {{0, 0.525}, {-0.075, 1}}],
 ListPlot[laSimul1, PlotMarkers → {Graphics[{Black, Disk[]}], 0.055}],
 ListPlot[laSimul2, PlotMarkers → {Graphics[{Red, Disk[]}], 0.055}],
 ListPlot[laSimul3, PlotMarkers → {Graphics[{Orange, Disk[]}], 0.055}],
 Frame → True,
 FrameLabel → {Style["Baseline mutation rate, μb"],
   Style["The mean number of novel mutations, λ(u*)"]},
 BaseStyle → {FontFamily → "Times New Roman", 14},
 LabelStyle → {FontFamily → "Times New Roman", 14}, AspectRatio → 0.85]
```

Out[ ]=

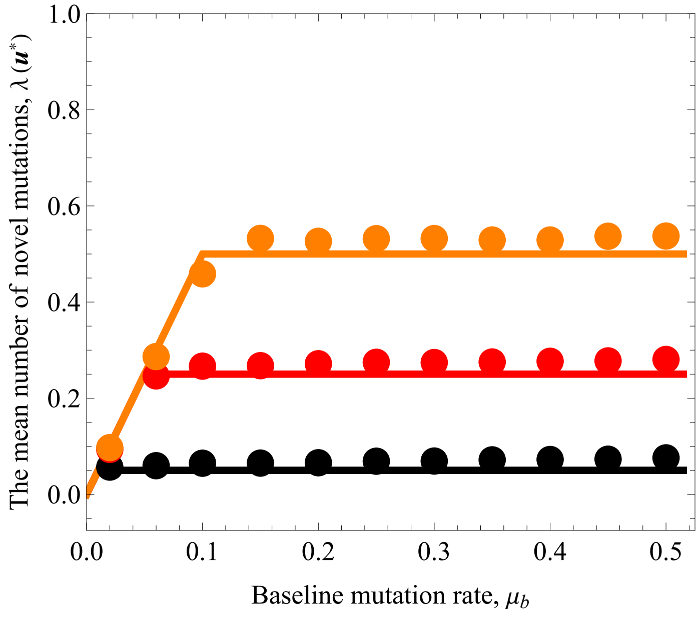

In[ ]:=

```
Show[Plot[{mut[x, 0.02, 2], mut[x, 0.1, 2], mut[x, 0.2, 2]}, {x, 0, 0.515},
   PlotStyle → {{Black, Thickness[0.012]}, {Red, Thickness[0.012]},
     {Orange, Thickness[0.012]}}, PlotRange → {{0, 0.525}, {0, 0.2}}
  ],
  ListPlot[muSimul1, PlotMarkers → {Graphics[{Black, Disk[]}], 0.055}],
  ListPlot[muSimul2, PlotMarkers → {Graphics[{Red, Disk[]}], 0.055}],
  ListPlot[muSimul3, PlotMarkers → {Graphics[{Orange, Disk[]}], 0.055}],
  Frame → True,
  FrameLabel → {Style["Baseline mutation rate, μb"],
    Style["Uninvadable mutation rate, μ(u*)"]},
  BaseStyle → {FontFamily → "Times New Roman", 14},
  LabelStyle → {FontFamily → "Times New Roman", 14}, AspectRatio → 0.85]
```

Out[ ]=

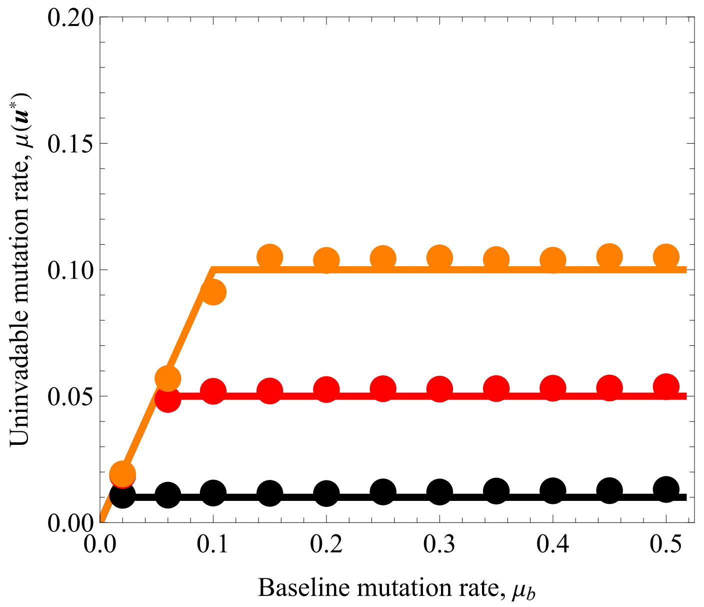

#### 1.5.1b. Average trait values and mean number of novel mutations with standard deviations

In[ ]:=

```
Needs["ErrorBarPlots`"]
Show[Plot[{Clip[urSin[x, 0.02, 2], {0, 1}],
    Clip[urSin[x, 0.1, 2], {0, 1}], Clip[urSin[x, 0.2, 2], {0, 1}]}, {x, 0, 0.515},
  PlotStyle → {{Black, Thickness[0.003]}, {Red, Thickness[0.003]},
    {Orange, Thickness[0.003]}}, PlotRange → {{0, 0.525}, {-0.075, 1}},
  PlotLegends → Placed[{Style["α=0.02", 16, FontFamily → "Times New Roman"],
     Style["α=0.1", 16, FontFamily → "Times New Roman"],
     Style["α=0.2", 16, FontFamily → "Times New Roman"]}, {0.75, 0.25}]
 ],
 (*ListPlot[urSimul1, PlotMarkers →{Graphics[{Black,Disk[]}],0.001}],
 ListPlot[urSimul2, PlotMarkers →{Graphics[{Red,Disk[]}],0.001}],
 ListPlot[urSimul3, PlotMarkers →{Graphics[{Orange,Disk[]}],0.001}],*)
 ErrorListPlot[urWithErr1, PlotStyle → Black,
  PlotMarkers → {Graphics[{Black, Disk[]}], 0.001}],
 ErrorListPlot[urWithErr2, PlotStyle → Red,
  PlotMarkers → {Graphics[{Black, Disk[]}], 0.001}],
 ErrorListPlot[urWithErr3, PlotStyle → Orange,
  PlotMarkers → {Graphics[{Black, Disk[]}], 0.001}], Frame → True,
 FrameLabel → {Style["Baseline mutation rate, μb"],
   Style["Uninvadable investment into repair, u*g"]},
 BaseStyle → {FontFamily → "Times New Roman", 14},
 LabelStyle → {FontFamily → "Times New Roman", 14}, AspectRatio → 0.85]
```

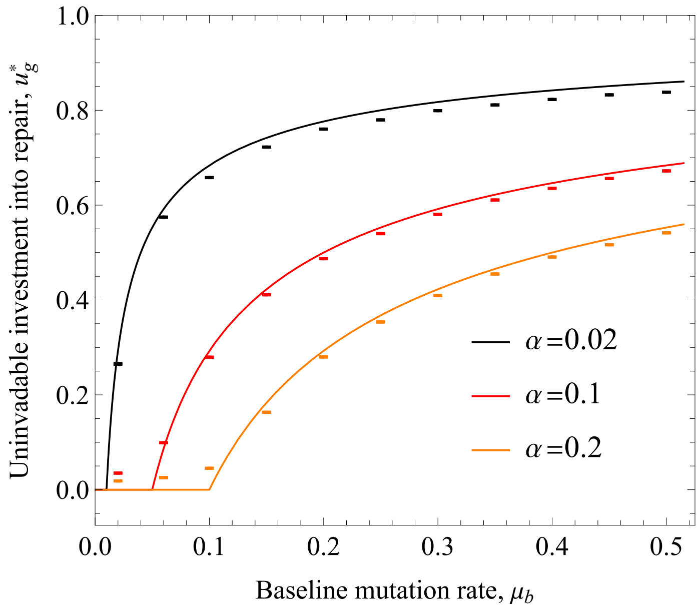

In[ ]:=
```
Needs["ErrorBarPlots`"]
Show[Plot[{Clip[usSin[x, 0.02, 2, 0.5], {0, 1}],
   Clip[usSin[x, 0.1, 2, 0.5], {0, 1}], Clip[usSin[x, 0.2, 2, 0.5], {0, 1}]},
  {x, 0, 0.515}, PlotStyle → {{Black, Thickness[0.003]},
    {Red, Thickness[0.003]}, {Orange, Thickness[0.003]}},
  PlotRange → {{0, 0.525}, {-0.075, 1.0}}],
 (*ListPlot[usSimul1, PlotMarkers →{Graphics[{Black,Disk[]}],0.055}],
 ListPlot[usSimul2, PlotMarkers →{Graphics[{Red,Disk[]}],0.055}],
 ListPlot[usSimul3, PlotMarkers →{Graphics[{Orange,Disk[]}],0.055}],*)
 ErrorListPlot[usWithErr1, PlotStyle → Black,
  PlotMarkers → {Graphics[{Black, Disk[]}], 0.001}],
 ErrorListPlot[usWithErr2, PlotStyle → Red,
  PlotMarkers → {Graphics[{Black, Disk[]}], 0.001}],
 ErrorListPlot[usWithErr3, PlotStyle → Orange,
  PlotMarkers → {Graphics[{Black, Disk[]}], 0.001}],
 Frame → True,
 FrameLabel → {Style["Baseline mutation rate, μb"],
   Style["Uninvadable investment into survival, u*s"]},
 BaseStyle → {FontFamily → "Times New Roman", 14},
 LabelStyle → {FontFamily → "Times New Roman", 14}, AspectRatio → 0.85]
```

Out[ ]=

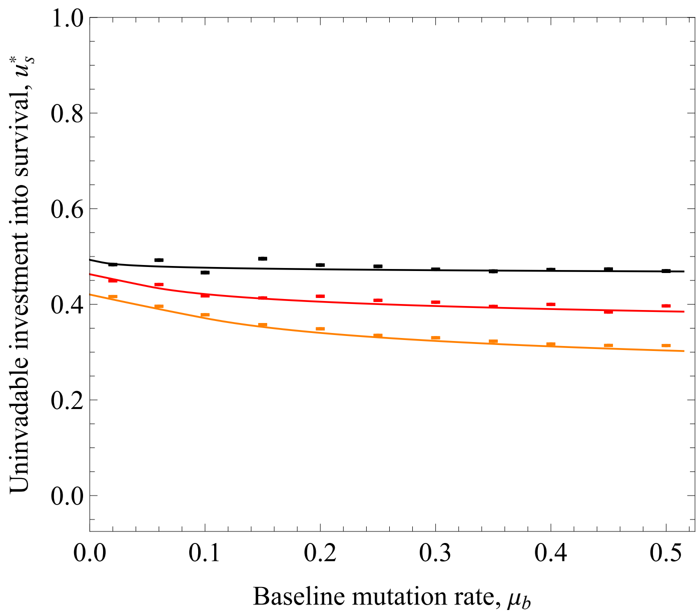

In[ ]:=
```
Needs["ErrorBarPlots`"]
Show[Plot[{laSin[ 0.02, 2, 0.2], laSin[0.1, 2, 0.2], laSin[ 0.2, 2, 0.2]},
  {μb, 0, 0.515}, PlotStyle → {{Black, Thickness[0.003]}, {Red, Thickness[0.003]},
    {Orange, Thickness[0.003]}}, PlotRange → {{0, 0.525}, {-0.075, 1}}],
 (*ListPlot[laSimul1, PlotMarkers →{Graphics[{Black,Disk[]}],0.055}],
 ListPlot[laSimul2, PlotMarkers →{Graphics[{Red,Disk[]}],0.055}],
 ListPlot[laSimul3, PlotMarkers →{Graphics[{Orange,Disk[]}],0.055}], *)
 ErrorListPlot[laWithErr1, PlotStyle → Black,
  PlotMarkers → {Graphics[{Black, Disk[]}], 0.001}],
 ErrorListPlot[laWithErr2, PlotStyle → Red,
  PlotMarkers → {Graphics[{Black, Disk[]}], 0.001}],
 ErrorListPlot[laWithErr3, PlotStyle → Orange,
  PlotMarkers → {Graphics[{Black, Disk[]}], 0.001}],
 Frame → True,
 FrameLabel → {Style["Baseline mutation rate, μb"],
   Style["The mean number of novel mutations, λ(u*)"]},
 BaseStyle → {FontFamily → "Times New Roman", 14},
 LabelStyle → {FontFamily → "Times New Roman", 14}, AspectRatio → 0.85]
```

Out[ ]=

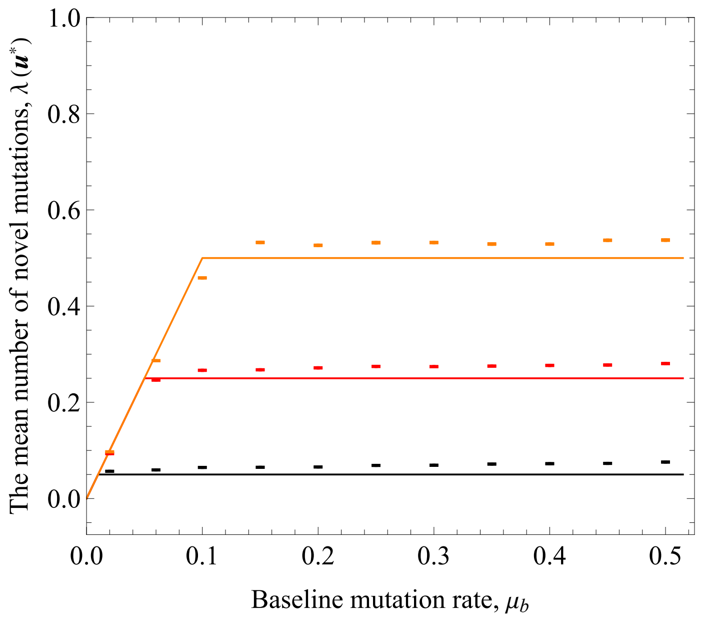


#### 1.5.2. Evaluate uninvadability at the parameter values plotted

In[ ]:=
```
(*Check uninvdability,
note that the Hessian should be initialized here from section 1.2.2. *)
```

In[ ]:=
```
(* α→0.02,α→0.1,α→0.2*)
```

In[ ]:=
```
Do[
 Print[NegativeDefiniteMatrixQ[HessianEx1[μb] /. {sb → 0.5, α → 0.02, αμ → 2}]],
 {μb, 0.1, 2, 0.1}]
```

```
True
True
True
True
True
True
True
True
True
True
True
True
True
True
True
True
True
True
True
True
```

In[ ]:=

```
Do[Print[NegativeDefiniteMatrixQ[HessianEx1[μb] /. {sb → 0.5, α → 0.1, αμ → 2}]],
  {μb, 0.1, 2, 0.1}]
```

```
True
True
True
True
True
True
True
True
True
True
True
True
True
True
True
True
True
True
True
True
```

In[ ]:=
```
Do[Print[NegativeDefiniteMatrixQ[HessianEx1[μb] /. {sb → 0.5, α → 0.2, αμ → 2}]],
 {μb, 0.1, 2, 0.1}]
```

```
True
True
True
True
True
True
True
True
True
True
True
True
True
True
True
True
True
True
True
True
```

In[ ]:=

```
(*If TRUE→ Hessian in negative definite for these parameter values →
  Singular strategy is uninvdable*)
```

##### 1.5.3. Evaluate convergence stability  at the parameter values plotted

In[ ]:=

```
(*Check convergence stability,
note that the Jacobian should be initialized here from section 1.2.2. *)
```

In[ ]:=

```
Do[Print[NegativeDefiniteMatrixQ[JacobEx1[μb] /. {sb → 0.5, α → 0.02, α_μ → 2}]],
  {μb, 0.1, 2, 0.1}]
```

```
True
True
True
True
True
True
True
True
True
True
True
True
True
True
True
True
True
True
True
True
```

In[ ]:=
```
Do[Print[NegativeDefiniteMatrixQ[JacobEx1[μb] /. {sb → 0.5, α → 0.1, αμ → 2}]],
  {μb, 0.1, 2, 0.1}]
```

True

True

True

True

True

True

True

True

True

True

True

True

True

True

True

True

True

True

True

True

In[ ]:=
```
Do[Print[NegativeDefiniteMatrixQ[JacobEx1[μb] /. {sb → 0.5, α → 0.2, αμ → 2}]],
  {μb, 0.1, 2, 0.1}]
```

```
True
True
True
True
True
True
True
True
True
True
True
True
True
True
True
True
True
True
True
True
```

```
In[ ]:= (*If TRUE→ Jacobian of selection gradients is negative definite for
        these parameter values → Singular strategy is converegnce stable*)
```

#### 1.5.4 . Figure 3 (convergence stability) in the main text

```
In[ ]:= Clear["Global`*"]
```

In[ ]:= $\mu[vr\_] := \mu b * (1 - vr)^{\alpha\mu}$

In[ ]:= $sp[vs\_, vr\_] := sb * (vs * (1 - vr))^{\alpha_s} * e^{-\mu[vr]}$

In[ ]:= Sur[vs_, vr_] :=

$$\frac{1}{(1 - vr)\ (1 - sp[vs, vr])} (\mu[vr]\ \alpha\mu - (\alpha_s * sp[vs, vr] + \alpha_f * (1 - sp[vs, vr])))\ /.\ \{\alpha_f \to \alpha, \alpha_s \to \alpha\}\ //\ FullSimplify$$

In[ ]:= Sus[vs_, vr_] :=

$$\frac{1}{(1 - sp[vs, vr])} \left(\frac{\alpha_s * sp[vs, vr]}{vs} - \frac{\alpha_f * (1 - sp[vs, vr])}{(1 - vs)}\right)\ /.\ \{\alpha_f \to \alpha, \alpha_s \to \alpha\}\ //\ FullSimplify$$

In[ ]:= $Data = \{\alpha \to 0.1, \mu b \to 0.2, \alpha\mu \to 2, sb \to 0.5\};$

In[ ]:= $urSin[\mu b\_, \alpha\_, \alpha\mu\_] := 1 - \left(\frac{\alpha}{\alpha\mu\ \mu b}\right)^{\frac{1}{\alpha\mu}}$

In[ ]:=
```
usSin[μb_, α_ , αμ_, sb_] :=
```

$$\text{Piecewise}\left[\left\{\left\{\left(\frac{e^{\mu b}}{sb}\right)^{\frac{1}{-1+\alpha}},\ \mu b < \frac{\alpha}{\alpha\mu}\right\},\ \left\{\left(\frac{e^{\left(\frac{\alpha}{\alpha\mu}\right)} * \left(\frac{\alpha}{\alpha\mu\ \mu b}\right)^{-\frac{\alpha}{\alpha\mu}}}{sb}\right)^{\frac{1}{(\alpha-1)}},\ \mu b > \frac{\alpha}{\alpha\mu}\right\}\right\}\right]$$

In[ ]:=
```
urESS = urSin[μb, α, αμ] /. Data (* ESS allocation to repair*)
usESS = usSin[μb, α, αμ, sb] /. Data  (*ESS allocation to survival*)
```

Out[ ]= 0.5

Out[ ]= 0.405459

In[ ]:=
```
p1 = VectorPlot[{Sur[vs, vr] /. Data, Sus[vs, vr] /. Data},
  {vr, 0.05, 0.95}, {vs, 0.05, 0.95}]
```

Out[ ]=

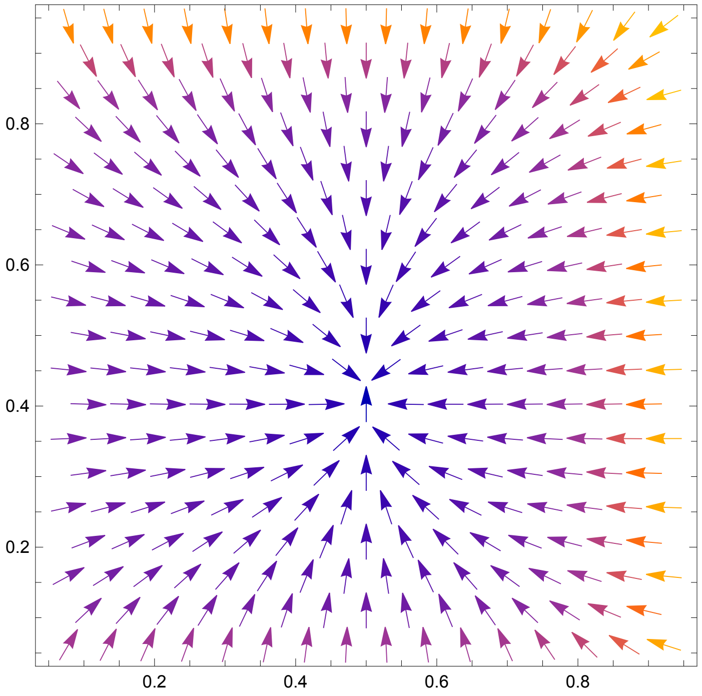


Get Data 1

Plot Data 1

Get Data 2

Plot Data 2

Plot Data 3

Get Data 4

Plot Data 4

Get Data 4

Plot Data 4

#### Plot it all together

In[ ]:= pAnalysis = 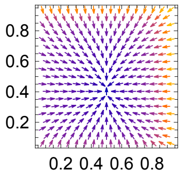

;

```
In[ ]:= ESSPoint =
  Graphics[{EdgeForm[Black], Gray, Thickness[.01], Disk[{urESS, usESS}, 0.015]}];
```

```
In[ ]:= Show[{pAnalysis, ESSPoint }]
```

Out[ ]=

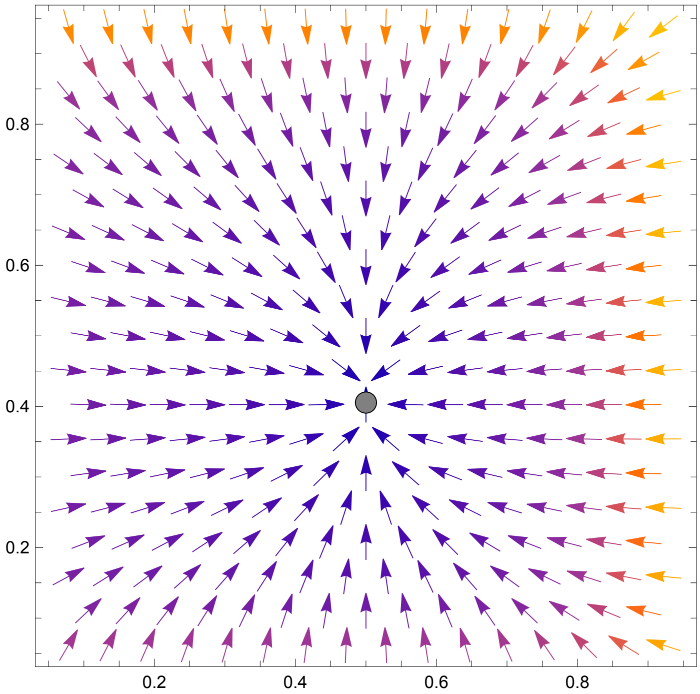

In[ ]:=
```
Show[{pAnalysis, pData1, pData2, pData3, pData4, ESSPoint },
  FrameLabel → {Style["Investment into maintenance, vg"],
    Style["Investment into survival, vs"]},
  BaseStyle → {FontFamily → "Times New Roman", 14},
  LabelStyle → {FontFamily → "Times New Roman", 14}]
```

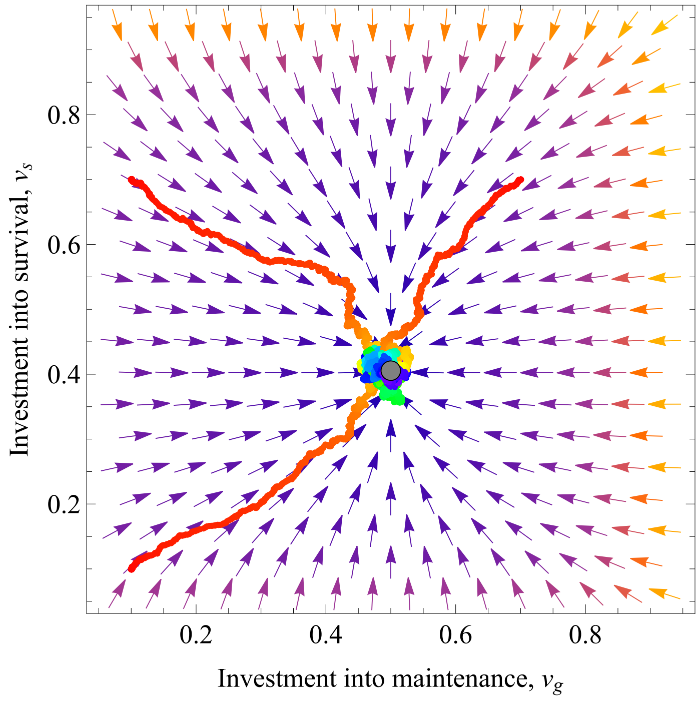


In[ ]:=
```
bar = BarLegend[{{"Rainbow", "Reverse"}, {0, 3000}}]
```

Out[ ]=

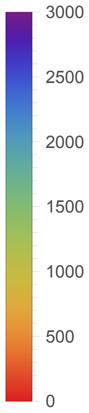

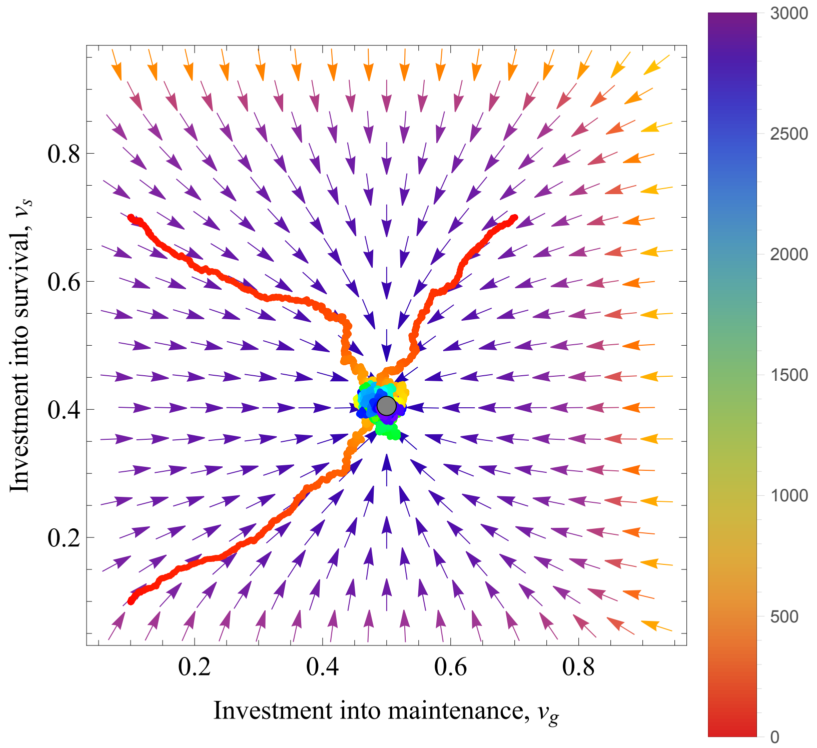
0.8
0.6
0.4
0.2
Investment into survival, $v_s$
0.2
0.4
0.6
0.8
Investment into maintenance, $v_g$
3000
2500
2000
1500
1000
500
0

#### 1.5.5. Additional plots

In[ ]:=

```
Show[Plot[{Clip[urSin[x, 0.1, 1], {0, 1}],
    Clip[urSin[x, 0.1, 2], {0, 1}], Clip[urSin[x, 0.1, 4], {0, 1}]}, {x, 0, 0.515},
  PlotStyle → {{Black, Thickness[0.012]}, {Red, Thickness[0.012]},
    {Orange, Thickness[0.012]}}, PlotRange → {{0, 0.525}, {-0.075, 1}},
  PlotLegends → Placed[{Style["αμ=1", 16, FontFamily → "Times New Roman"],
     Style["αμ=2", 16, FontFamily → "Times New Roman"],
     Style["αμ=4", 16, FontFamily → "Times New Roman"]}, {0.75, 0.25}]
 ],
 FrameLabel → {Style["Baseline mutation rate, μb"],
   Style["Uninvadable investment into repair, u*g"]},
 BaseStyle → {FontFamily → "Times New Roman", 14},
 LabelStyle → {FontFamily → "Times New Roman", 14}, AspectRatio → 0.85]
```

Out[ ]=

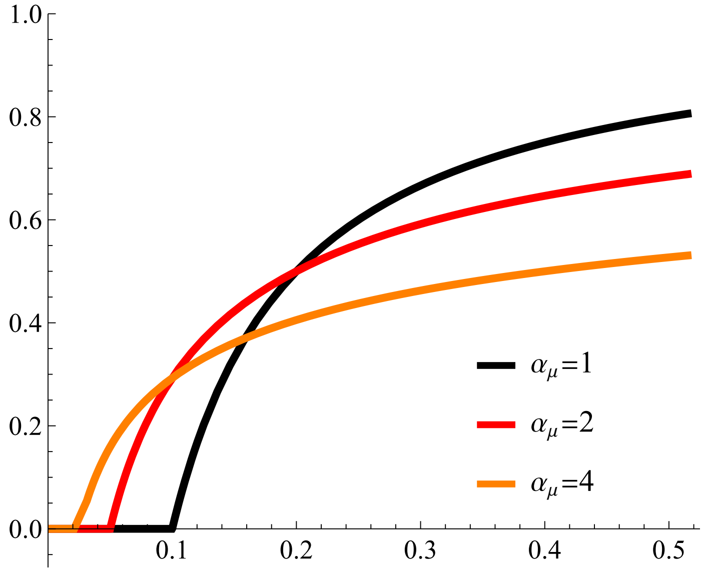

In[ ]:=

```
Show[Plot[{Clip[usSin[x, 0.1, 1, 0.5], {0, 1}],
    Clip[usSin[x, 0.1, 2, 0.5], {0, 1}], Clip[usSin[x, 0.1, 4, 0.5], {0, 1}]},
   {x, 0, 0.515}, PlotStyle → {{Black, Thickness[0.012]},
     {Red, Thickness[0.012]}, {Orange, Thickness[0.012]}},
   PlotRange → {{0, 0.525}, {-0.075, 1.0}}],
  Frame → True,
  FrameLabel → {Style["Baseline mutation rate, μb"],
    Style["Uninvadable investment into survival, u*s"]},
  BaseStyle → {FontFamily → "Times New Roman", 14},
  LabelStyle → {FontFamily → "Times New Roman", 14}, AspectRatio → 0.85]
```

Out[ ]=

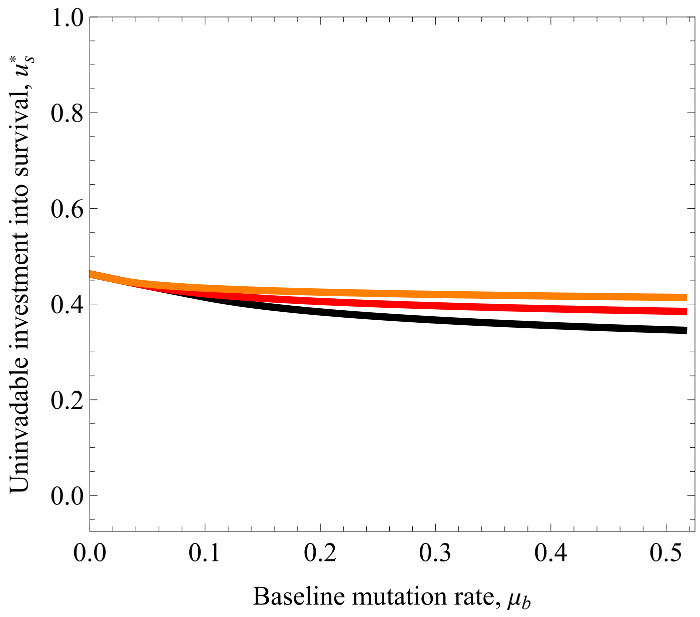

```
Show[Plot[{Clip[usSin[x, 0.02, 2, 0.5], {0, 1}],
   Clip[usSin[x, 0.1, 2, 0.5], {0, 1}], Clip[usSin[x, 0.2, 2, 0.5], {0, 1}]},
  {x, 0, 0.515}, PlotStyle → {{Black, Thickness[0.012]},
    {Red, Thickness[0.012]}, {Orange, Thickness[0.012]}},
  GridLines → {{}, {{usSinClassic[0.5, 0.02 ], Black},
     {usSinClassic[0.5, 0.1 ], Red}, {usSinClassic[0.5, 0.2 ], Orange}}},
  GridLinesStyle → Directive[Dashed, Thick],
  PlotRange → {{0, 0.525}, {-0.075, 1.0}}],
 ListPlot[usSimul1, PlotMarkers → {Graphics[{Black, Disk[]}], 0.055}],
 ListPlot[usSimul2, PlotMarkers → {Graphics[{Red, Disk[]}], 0.055}],
 ListPlot[usSimul3, PlotMarkers → {Graphics[{Orange, Disk[]}], 0.055}],
 Frame → True,
 FrameLabel → {Style["Baseline mutation rate,
      \!\(\*SubscriptBox[StyleBox[\"μ\",FontSlant->\"Italic\"],
      \"b\"]\)"], Style["Uninvadable investment into survival,
      \!\(\*SubsuperscriptBox[StyleBox[\"u\",FontSlant->\"Italic\"],
      \"s\", \"*\"]\)"]},
 BaseStyle → {FontFamily → "Times New Roman",
   14},
 LabelStyle → {FontFamily → "Times New Roman", 14}, AspectRatio → 0.85]
```

Below we also plot of allocation to survival with Classic life - history lines (dashed) . However, the lines are a bit too close to each other, so we suppressed the classic lines in the main text and zooming is also not ideal as other allocation strategy is  not zoomed in.

In[ ]:=

```
Show[Plot[{Clip[usSin[x, 0.02, 2, 0.5], {0, 1}],
    Clip[usSin[x, 0.1, 2, 0.5], {0, 1}], Clip[usSin[x, 0.2, 2, 0.5], {0, 1}]},
  {x, 0, 0.515}, PlotStyle → {{Black, Thickness[0.012]},
    {Red, Thickness[0.012]}, {Orange, Thickness[0.012]}},
  GridLines → {{}, {{usSinClassic[0.5, 0.02 ], Black},
     {usSinClassic[0.5, 0.1 ], Red}, {usSinClassic[0.5, 0.2 ], Orange}}},
  GridLinesStyle → Directive[Dashed, Thick],
  PlotRange → {{0, 0.525}, {-0.075, 1.0}}],
 ListPlot[usSimul1, PlotMarkers → {Graphics[{Black, Disk[]}], 0.055}],
 ListPlot[usSimul2, PlotMarkers → {Graphics[{Red, Disk[]}], 0.055}],
 ListPlot[usSimul3, PlotMarkers → {Graphics[{Orange, Disk[]}], 0.055}],
 Frame → True,
 FrameLabel → {Style["Baseline mutation rate, μb"],
   Style["Uninvadable investment into survival, u*s"]},
 BaseStyle → {FontFamily → "Times New Roman", 14},
 LabelStyle → {FontFamily → "Times New Roman", 14}, AspectRatio → 0.85]
```

Out[ ]=

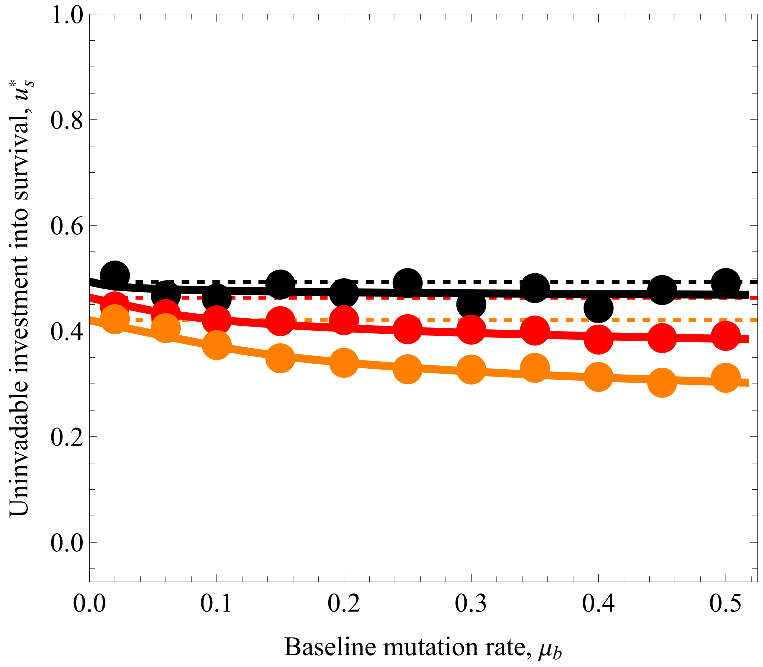

In[ ]:=

```
Show[Plot[{Clip[usSin[x, 0.02, 2, 0.5], {0, 1}],
    Clip[usSin[x, 0.1, 2, 0.5], {0, 1}], Clip[usSin[x, 0.2, 2, 0.5], {0, 1}]},
   {x, 0, 0.515}, PlotStyle → {{Black, Thickness[0.012]},
     {Red, Thickness[0.012]}, {Orange, Thickness[0.012]}},
   GridLines → {{}, {{usSinClassic[0.5, 0.02 ], Black},
      {usSinClassic[0.5, 0.1 ], Red}, {usSinClassic[0.5, 0.2 ], Orange}}},
   GridLinesStyle → Directive[Dashed, Thick],
   PlotRange → {{0, 0.525}, {0.2, 0.6}}],
  ListPlot[usSimul1, PlotMarkers → {Graphics[{Black, Disk[]}], 0.055}],
  ListPlot[usSimul2, PlotMarkers → {Graphics[{Red, Disk[]}], 0.055}],
  ListPlot[usSimul3, PlotMarkers → {Graphics[{Orange, Disk[]}], 0.055}],
  Frame → True,
  FrameLabel → {Style["Baseline mutation rate, μb"],
    Style["Uninvadable investment into survival, u*s"]},
  BaseStyle → {FontFamily → "Times New Roman", 14},
  LabelStyle → {FontFamily → "Times New Roman", 14}, AspectRatio → 0.85]
```

Out[ ]=

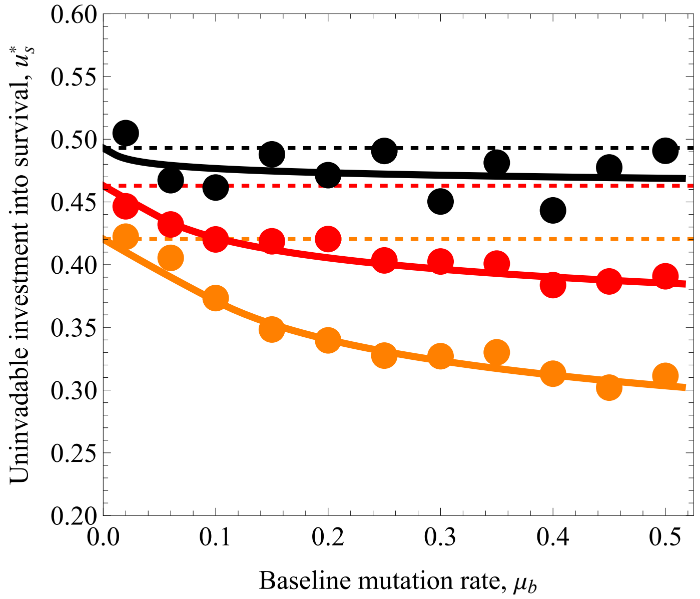


In[ ]:= (*Repair allocation is highest when α is small, $\alpha_\mu$ is small*)

In[ ]:=
```
ContourPlot[urSin[μb, α , α_μ] /. {μb → 2},
  {α, 0.25, 1}, {α_μ, 1, 4}, PlotLegends → Automatic]
```

Out[ ]=

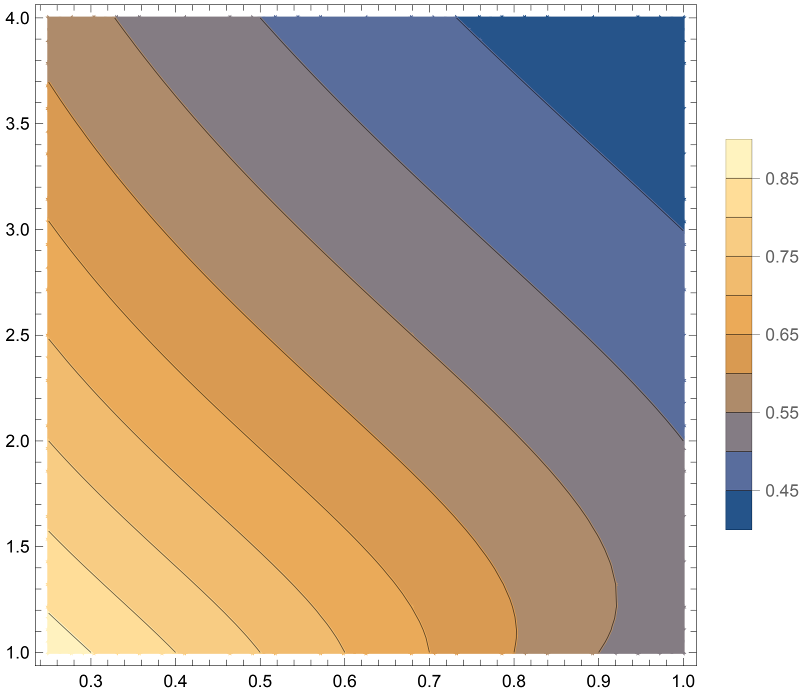


In[ ]:=
```
(*Survival allocation is highest when α is small, α_μ is large*)
```

In[ ]:=
```
ContourPlot[usSin[μb, α , α_μ, sb] /. {μb → 2, sb → 0.5},
  {α, 0.25, 1}, {α_μ, 1, 4}, PlotLegends → Automatic]
```

Out[ ]=

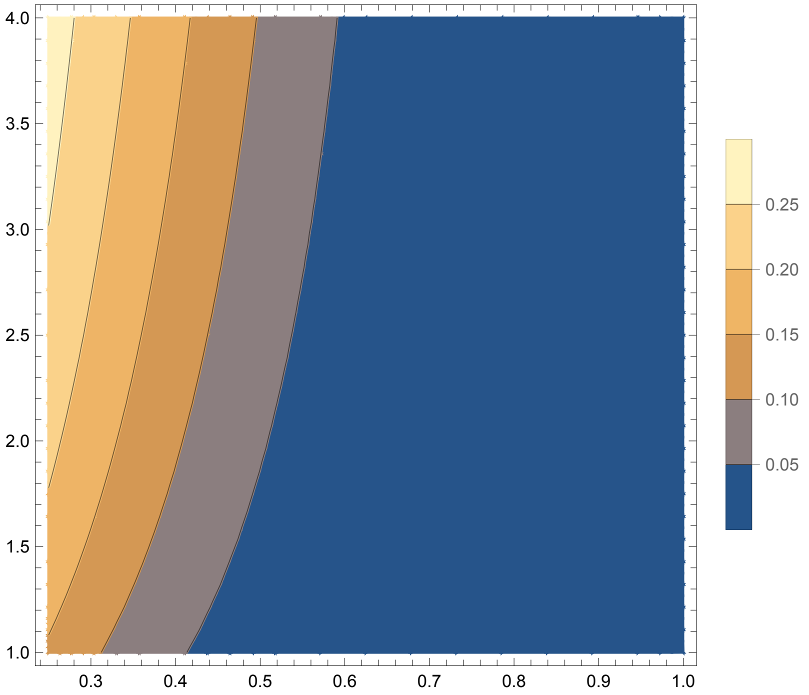


### 2. ”Co-evolution of age-at-maturation and

mutation rate”

In[ ]:=
```
Clear["Global`*"]
```

### 2.1. Analysis of the model

**Important parameters and their notations in this Mathematica file:**

ut, vt - mutant and resident age at maturity (u_m, v_m, in main text)
ur, vr - mutant and resident allocation to repair (u_g, v_g in the main text)
x[t], xm[t] - body size and body size at maturity
mut[ur], mf - mutation rate during ageing, mutation rate during reproduction
B[ut,ut] or Bsyn - energy available to allocation

d - baseline death rate (external mortality)
αb - allocation to growth or fecundity exponent
α - allocation to repair exponent

b0, l0 - effective fecundity and survival with no new mutations of the least loaded class
NGen[ut, ur] - population size as a function of traits

In[ ]:=
```
Clear["Global`*"]
```

#### 2.1.1. Basic reproductive value and equilibrium population size

In[ ]:=
```
(*mutation rate during lifetime, last line of eq. (12) *)
```

In[ ]:=
```
mut[ur_] = μb (1 - ur)^α;
```

In[ ]:=
```
(*Growth equation, eq. (14)*)
```

In[ ]:=
```
FullSimplify[
 DSolve[{x'[t] == β * (1 - ur^αb) * a * (x[t])^c , x[0] == xo}, x[t], t] , Reals]
```

Solve : Inverse functions are being used by Solve, so some solutions may not be found; use Reduce for complete solution information.

Solve : Inverse functions are being used by Solve, so some solutions may not be found; use Reduce for complete solution information.

Out[ ]=
$$\left\{\left\{x[t] \to \left(xo^{1-c} + a\,(-1+c)\,t\,\left(-1+ur^{\alpha b}\right)\beta\right)^{\frac{1}{1-c}}\right\}\right\}$$

In[ ]:= (*Body size at maturity, eq. (15)*)

$xm[ut\_, ur\_] = \left(xo^{1-c} + a\,(-1 + c)\; ut\,\left(-1 + ur^{\alpha b}\right)\beta\right)^{\frac{1}{1-c}};$

In[ ]:= (* Bsyn Productivity as a function of body size *)

$B[ut\_, ur\_] := a * (xm[ut, ur])^{c}$

In[ ]:=

(*Effective fecundity with no new mutations, eq. (13)*)

In[ ]:= $b0[ut\_, ur\_] = \left(1 - ur^{\alpha b}\right) * B[ut, ur]\;(1 - \gamma * N)\; e^{-mf}$ // FullSimplify;

(*survival with no new mutations, see below eq. B-7*)

In[ ]:= FullSimplify[DSolve[{l′[t] == - (d + mut[ur]) * l[t], l[0] == 1}, l[t], t] , Reals]

Out[ ]= $\left\{\left\{l[t] \to e^{-t\,\left(d + (1-ur)^{\alpha}\,\mu_b\right)}\right\}\right\}$

(*Def. survival with no new mutations, below eq. B-7 in the main text*)

In[ ]:= $l0[t\_] = e^{-t\,\left(d + (1-ur)^{\alpha}\,\mu_b\right)};$

(* R0, eq. A-8 of the main text, note that
b0= $\left(1-ur^{\alpha b}\right)$*BSyn* (1-$\gamma$ *N)$e^{-mf}$ where Bsyn=B[ut,ur] *)

In[ ]:= $Ro[ut\_, ur\_] = FullSimplify\left[\left(1 - ur^{\alpha b}\right) * BSyn * (1 - \gamma * N)\; e^{-mf} \int_{ut}^{\infty} l0[t]\, dt\, , \{(d + (1 - ur)^{\alpha}\,\mu_b) > 0\}\right];$

(* Proof of the simplification eq. A-8 *)

In[ ]:= $FullSimplify\left[PowerExpand\left[BSyn * \left(1 - ur^{\alpha b}\right) * e^{-mf}\, \frac{e^{-ut\,(d+mut[ur])}}{d + mut[ur]} * (1 - N\gamma)\right] == PowerExpand[Ro[ut, ur]]\right]$

Out[ ]= True

In[ ]:= (*Derivation of equilibrium population size in the resident population*)

In[ ]:=

Solve[Ro[vt, vr] == 1, N] // FullSimplify

Out[ ]= $\left\{\left\{N \to \frac{1 + \frac{e^{mf + d\,vt + (1-vr)^{\alpha}\,vt\,\mu_b}\,\left(d + (1-vr)^{\alpha}\,\mu_b\right)}{BSyn\,\left(-1 + vr^{\alpha b}\right)}}{\gamma}\right\}\right\}$

In[ ]:= $NGen[vt\_, vr\_] := \frac{1 + \frac{e^{mf + d\,vt + (1-vr)^{\alpha}\,vt\,\mu_b}\,\left(d + (1-vr)^{\alpha}\,\mu_b\right)}{BSyn\,\left(-1 + vr^{\alpha b}\right)}}{\gamma}$

(* Proof of eq. B-9 , where BSyn=a*(xm[vt,vr])$^{c}$, see above *)

In[ ]:=
```
FullSimplify[PowerExpand[(BSyn (1 - vr^αb) - (d + mut[vr]) ⅇ^(mf+(mut[vr]+d)*vt)) /
    (γ * BSyn (1 - vr^αb))] == PowerExpand[NGen[vt, vr]]]
```

Out[ ]= True

In[ ]:=
```
(*Equilibrium  population size in terms of traits *)
```

In[ ]:=
```
NEquil[vt_, vr_] = NGen[vt, vr] /. {BSyn → B[vt, vr]}
```

Out[ ]=
$$\frac{1+\frac{e^{mf+d\,vt+(1-vr)^{\alpha}\,vt\,\mu_b}\left(\left(xo^{1-c}+a\,(-1+c)\,\left(-1+vr^{\alpha b}\right)\,vt\,\beta\right)^{\frac{1}{1-c}}\right)^{-c}\,(d+(1-vr)^{\alpha}\,\mu_b)}{a\,\left(-1+vr^{\alpha b}\right)}}{\gamma}$$

In[ ]:=
```
(* Equilibrium population size in classical life-history theory*)
```

In[ ]:=
```
NClassic[ut_] = FullSimplify[NEquil[vt, vr] /. {vr → 0, μb → 0, αb → 2, α → 2, mf → 0}]
```

Out[ ]=
$$\frac{1-\frac{d\,e^{d\,vt}\left(\left(xo^{1-c}-a\,(-1+c)\,vt\,\beta\right)^{\frac{1}{1-c}}\right)^{-c}}{a}}{\gamma}$$

#### 2.1.2. Selection gradients

In[ ]:=
```
(*Redefine Ro, where Bsyn is expressed in terms of traits*)
Ro[ut_, ur_] = Ro[ut, ur] /. {BSyn → B[ut, ur]};

(* Selection gradient on repair, derivation of eq. (16) *)
```

In[ ]:=
```
Su[vt_, vr_] =
  FullSimplify[Assuming[μb > 0 && 0 < ur < 1 && a > 0 && α > 0 && αb > 0 && xo > 0 &&
      c > 0 && ts > 0 && γ > 0 && d > 0 && β > 0,
     Refine[D[Ro[ut, ur], ur] // FullSimplify]] /. {ut → vt, ur → vr}]
```

Out[ ]=
$$\frac{1}{(d+(1-vr)^{\alpha}\,\mu_b)^2}\,a\,e^{-mf-vt\,(d+(1-vr)^{\alpha}\,\mu_b)}$$
$$\left(\left(xo^{1-c}+a\,(-1+c)\,\left(-1+vr^{\alpha b}\right)\,vt\,\beta\right)^{\frac{1}{1-c}}\right)^{c}\,(-1+N\,\gamma)\,\Bigg((1-vr)^{-1+\alpha}\,\left(-1+vr^{\alpha b}\right)\,\alpha\,\mu_b+$$
$$vr^{-1+\alpha b}\,\alpha b\,(d+(1-vr)^{\alpha}\,\mu_b)-\frac{a\,c\,vr^{-1+\alpha b}\,\left(-1+vr^{\alpha b}\right)\,vt\,xo^{c}\,\alpha b\,\beta\,(d+(1-vr)^{\alpha}\,\mu_b)}{xo+a\,(-1+c)\,\left(-1+vr^{\alpha b}\right)\,vt\,xo^{c}\,\beta}+$$
$$(1-vr)^{-1+\alpha}\,\left(-1+vr^{\alpha b}\right)\,vt\,\alpha\,\mu_b\,(d+(1-vr)^{\alpha}\,\mu_b)\Bigg)$$

```
(* Selection gradient on the age at maturity, derivation of eq. (17) *)
```

In[ ]:=
```
St[vt_, vr_] =
  FullSimplify[Assuming[μb > 0 && 0 < ur < 1 && a > 0 && α > 0 && αb > 0 && xo > 0 &&
      c > 0 && ut > 0 && γ > 0 && d > 0 && β > 0,
     Refine[D[Ro[ut, ur], ut] // FullSimplify]] /. {ut → vt, ur → vr}]
```

Out[ ]=
$$-\left(\left(a\, e^{-mf-d\,vt-(1-vr)^{\alpha}\,vt\,\mu_b}\left(-1+vr^{\alpha b}\right)\left(\left(xo^{1-c}+a\,(-1+c)\left(-1+vr^{\alpha b}\right)vt\,\beta\right)^{\frac{1}{1-c}}\right)^{c}\right.\right.$$
$$(-1+N\,\gamma)\left(d\,xo+a\left(-1+vr^{\alpha b}\right)(c+(-1+c)\,d\,vt)\,xo^{c}\,\beta+\right.$$
$$\left.\left.(1-vr)^{\alpha}\left(xo+a\,(-1+c)\left(-1+vr^{\alpha b}\right)vt\,xo^{c}\,\beta\right)\mu_b\right)\right)\Big/$$
$$\left(\left(xo+a\,(-1+c)\left(-1+vr^{\alpha b}\right)vt\,xo^{c}\,\beta\right)\left(d+(1-vr)^{\alpha}\,\mu_b\right)\right)$$

**(*Selection gradient on repair, proof of eq. 16*)**

In[ ]:=
$$\text{FullSimplify}\Big[\text{PowerExpand}\Big[\frac{b0[vt,vr]\,e^{-(d+mut[vr])\,vt}}{(d+mut[vr])}*\left(\frac{\alpha\,mut[vr]}{(1-vr)}*\left(vt+\frac{1}{(d+mut[vr])}\right)-\alpha b*vr^{\alpha b-1}\left(\beta\frac{B[vt,vr]\,c\,vt}{xm[vt,vr]}+\frac{1}{\left(1-vr^{\alpha b}\right)}\right)\right)==Su[vt,vr]\Big]\Big]$$

Out[ ]= True

In[ ]:=
$$\text{FullSimplify}\Big[\text{PowerExpand}\Big[\frac{b0[vt,vr]\,e^{-(d+mut[vr])\,vt}}{(d+mut[vr])}*\left(\frac{\alpha\,mut[vr]}{(1-vr)}*\left(vt+\frac{1}{(d+mut[vr])}\right)-\frac{\alpha b*vr^{\alpha b-1}}{\left(1-vr^{\alpha b}\right)xm[vt,vr]}\left(\beta\left(1-vr^{\alpha b}\right)B[vt,vr]\,c\,vt+xm[vt,vr]\right)\right)==Su[vt,vr]\Big]\Big]$$

Out[ ]= True

NB! Note that at (ut = vt, ur = vr),

we have that $R0=\frac{b0[vt,vr]*e^{-(d+mut[vr])\,vt}}{(d+mut[vr])}=1$,

this can be obtained when substituting eq. 13 into eq. A - 8.

**(* Selection gradient on the switching time, proof of eq. 17*)**

In[ ]:=
$$\text{FullSimplify}\Big[\text{PowerExpand}\Big[\frac{b0[vt,vr]*e^{-(d+mut[vr])\,vt}}{(d+mut[vr])}*\left(c*\frac{\beta*\left(1-vr^{\alpha b}\right)*B[vt,vr]}{xm[vt,vr]}-(d+mut[vr])\right)==St[vt,vr]\Big]\Big]$$

Out[ ]= True

##### 2.1.3. Singular strategies

In[ ]:=
```
(* Switching time as a function of repair*)
Solve[St[ut, ur] == 0, ut] // FullSimplify
```

Solve : Inverse functions are being used by Solve, so some solutions may not be found; use Reduce for complete solution information.

Out[ ]=
$$\left\{\left\{ut \to \frac{\frac{xo^{1-c}}{a\,\beta - a\,ur^{\alpha b}\,\beta} - \frac{c}{d + (1-ur)^{\alpha}\,\mu_b}}{-1+c}\right\}\right\}$$

```
(* Proof of eq. 18, Age at maturity as a function of repair*)
```

In[ ]:=
```
FullSimplify[
 PowerExpand[1/(1 - c) (c/(d + mut[ur]) - xo/(a β (1 - ur^αb) xo^c)) == ((xo^(1-c))/(a β - a ur^αb β) - c/(d + (1 - ur)^α μb))/(-1 + c)]]
```

$$\text{FullSimplify}\left[\text{PowerExpand}\left[\frac{1}{1-c}\left(\frac{c}{(d+\text{mut}[ur])} - \frac{xo}{a\,\beta\,\left(1-ur^{\alpha b}\right)\,xo^{c}}\right) == \frac{\frac{xo^{1-c}}{a\,\beta - a\,ur^{\alpha b}\,\beta} - \frac{c}{d+(1-ur)^{\alpha}\,\mu_b}}{-1+c}\right]\right]$$

Out[ ]= True

In[ ]:=

$$\text{vtClassic} = \text{FullSimplify}\left[\frac{\frac{xo^{1-c}}{a\,\beta - a\,ur^{\alpha b}\,\beta} - \frac{c}{d+(1-ur)^{\alpha}\,\mu_b}}{-1+c} \;/.\; \{\mu_b \to 0, \alpha b \to 1\} \;/.\; \{ur \to 0\}\right];$$

In[ ]:=
```
(* Proof of age at maturity in classic life-history theory*)
```

In[ ]:=

$$\text{FullSimplify}\left[\text{PowerExpand}\left[\frac{1}{1-c}\left(\frac{c}{d} - \frac{xo}{a * \beta * xo^{c}}\right) == \text{vtClassic}\right]\right]$$

Out[ ]= True

In[ ]:=
```
(* Solution for αb→2, α→2 *)
```

In[ ]:=
```
sol = Solve[FullSimplify[St[vt, vr] == 0 /. {αb → 2, α → 2}] &&
    FullSimplify[Su[vt, vr] == 0 /. {αb → 2, α → 2} ], {vt, vr}] // FullSimplify
```

Solve : Inverse functions are being used by Solve, so some solutions may not be found; use Reduce for complete solution information.

Out[ ]=
$$\left\{\left\{vt \to \frac{xo^{-c}\left(2\sqrt{d}\,xo\,\mu_b + \left(d\,xo - a\,c\,xo^c\,\beta\right)\left(\sqrt{d} + \sqrt{d+4\,\mu_b}\right)\right)}{2\,a\,(-1+c)\,d\,\beta\,\sqrt{d+4\,\mu_b}},\; vr \to \frac{d + 2\,\mu_b - \sqrt{d}\,\sqrt{d+4\,\mu_b}}{2\,\mu_b}\right\},\right.$$
$$\left\{vt \to \frac{xo^{-c}\left(-2\sqrt{d}\,xo\,\mu_b + \left(d\,xo - a\,c\,xo^c\,\beta\right)\left(-\sqrt{d} + \sqrt{d+4\,\mu_b}\right)\right)}{2\,a\,(-1+c)\,d\,\beta\,\sqrt{d+4\,\mu_b}},\; vr \to \frac{d + 2\,\mu_b + \sqrt{d}\,\sqrt{d+4\,\mu_b}}{2\,\mu_b}\right\},$$
$$\left\{vt \to -\frac{xo^{-c}\left(a\,xo^c\,\beta + xo\,\mu_b\right)^2}{a\,\beta\left(a\,d\,xo^c\,\beta + \mu_b\left(d\,xo + 2\left(a\,xo^c\,\beta + \sqrt{a\,xo^c\,\beta\left(-d\,xo + a\,xo^c\,\beta\right) - d\,xo^2\,\mu_b}\right)\right)\right)},\; vr \to \frac{xo\,\mu_b - \sqrt{a\,xo^c\,\beta\left(-d\,xo + a\,xo^c\,\beta\right) - d\,xo^2\,\mu_b}}{a\,xo^c\,\beta + xo\,\mu_b}\right\},$$
$$\left.\left\{vt \to -\frac{xo^{-c}\left(a\,xo^c\,\beta + xo\,\mu_b\right)^2}{a\,\beta\left(a\,d\,xo^c\,\beta + \mu_b\left(d\,xo + 2\,a\,xo^c\,\beta - 2\sqrt{a\,xo^c\,\beta\left(-d\,xo + a\,xo^c\,\beta\right) - d\,xo^2\,\mu_b}\right)\right)},\; vr \to \frac{xo\,\mu_b + \sqrt{a\,xo^c\,\beta\left(-d\,xo + a\,xo^c\,\beta\right) - d\,xo^2\,\mu_b}}{a\,xo^c\,\beta + xo\,\mu_b}\right\}\right\}$$

In[ ]:=
```
ReplaceAll[sol, {xo → 0.1, a → 1, μb → 1, c → 0.75, d → 0.1, β → 1}]
```

Out[ ]=
```
{{vt → 12.5294, vr → 0.729844}, {vt → 15.2213, vr → 1.37016},
 {vt → -0.600269, vr → -0.251367}, {vt → -9.91795, vr → 0.971237}}
```

In[ ]:=
```
ReplaceAll[sol, {xo → 0.1, a → 1, μb → 10, c → 0.75, d → 0.1, β → 1}]
```

Out[ ]=
```
{{vt → 3.33542, vr → 0.904875}, {vt → 24.4152, vr → 1.10512},
 {vt → -1.20172, vr → 0.729419}, {vt → -9.10369, vr → 0.968622}}
```

In[ ]:=
```
ReplaceAll[sol, {xo → 0.1, a → 1, μb → 0.1, c → 0.75, d → 0.1, β → 1}]
```

Out[ ]=
```
{{vt → 19.0746, vr → 0.381966}, {vt → 8.67603, vr → 2.61803},
 {vt → -2.23308, vr → -0.864972}, {vt → -9.99186, vr → 0.971453}}
```

In[ ]:=
```
ReplaceAll[sol, {xo → 0.1, a → 1, μb → 0.01, c → 0.75, d → 0.1, β → 1}]
```

Out[ ]=
```
{{vt → 25.412, vr → 0.0839202}, {vt → 2.33864, vr → 11.9161},
 {vt → -7.22401, vr → -0.96029}, {vt → -9.99919, vr → 0.971474}}
```

```
(*vr -first solution*)
```

In[•]:= Manipulate$\left[\text{Plot}\left[\frac{d + 2\,\mu - \sqrt{d}\ \sqrt{d + 4\,\mu}}{2\,\mu}, \{d, 0.00001, 2\}\right], \{\mu, 0.0001, 2\}\right]$

Out[•]=

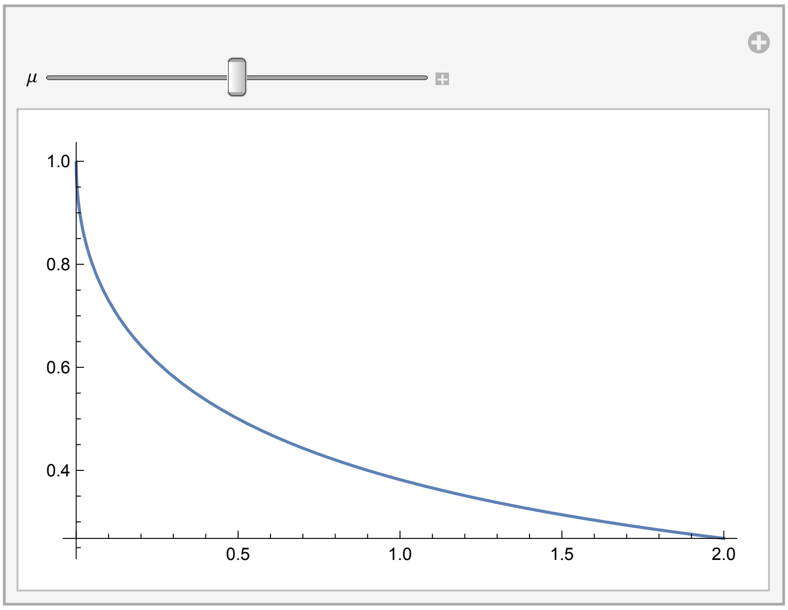


(*vr second solution*)

In[•]:= Manipulate$\left[\text{Plot}\left[\frac{d + 2\,\mu + \sqrt{d}\ \sqrt{d + 4\,\mu}}{2\,\mu}, \{d, 0.00001, 2\}\right], \{\mu, 0.0001, 10\}\right]$

Out[•]=

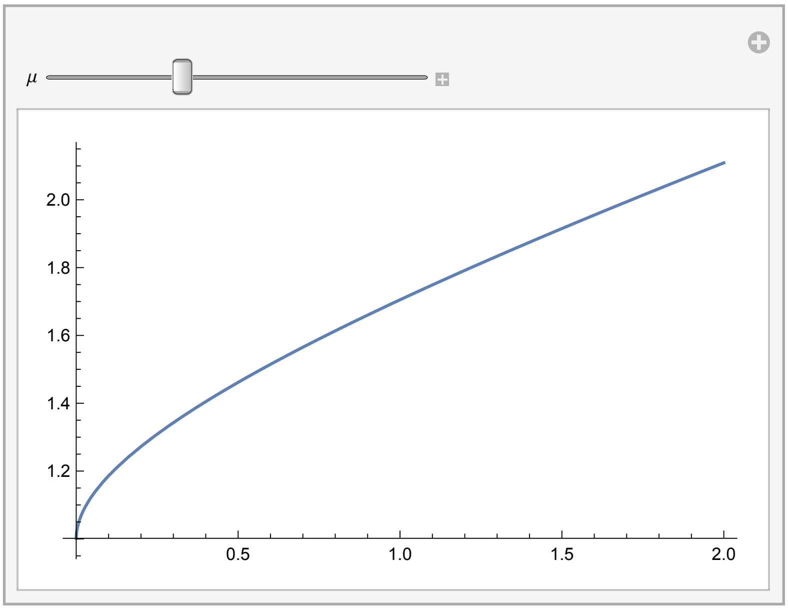


In[•]:= $\frac{d + 2\,\mu + \sqrt{d}\ \sqrt{d + 4\,\mu}}{2\,\mu} < 1$ // FullSimplify

Out[•]= $d\,\mu + \sqrt{d}\ \mu\ \sqrt{d + 4\,\mu} < 0$

(*this can not hold biologically, because μ and d are poth positive*)

(*vr third solution*)

In[∘]:=

```
(*Note thate we can always scale initial size to one*)
```

In[∘]:= $\frac{xo\,\mu_b - \sqrt{a\,xo^c\,\beta\,\left(-d\,xo + a\,xo^c\,\beta\right) - d\,xo^2\,\mu_b}}{a\,xo^c\,\beta + xo\,\mu_b} > 0$ /. xo → 1 // FullSimplify

Out[∘]= $(a\,\beta + \mu_b)\,\left(-\mu_b + \sqrt{a\,\beta\,(-d + a\,\beta) - d\,\mu_b}\right) < 0$

In[∘]:= (* Lets look at the solution when external mortality can be neglected*)

In[∘]:= $\left(-\mu_b + \sqrt{a\,\beta\,(-d + a\,\beta) - d\,\mu_b}\right) < 0$ /. d → 0 // FullSimplify

Out[∘]= $\sqrt{a^2\,\beta^2} < \mu_b$

```
(*Here, aβ can be interpretated as growth rate as size at birth
 is scaled to one, this solution implies that allocation to repair
 is only positive if mutation rate is larger than the growth rate,
this is not biologically not feasible *)
```

```
(*vt the fourth solution*)
```

In[∘]:= $-\frac{xo^{-c}\,\left(a\,xo^c\,\beta + xo\,\mu_b\right)^2}{a\,\beta\,\left(a\,d\,xo^c\,\beta + \mu_b\,\left(d\,xo + 2\,a\,xo^c\,\beta - 2\,\sqrt{a\,xo^c\,\beta\,\left(-d\,xo + a\,xo^c\,\beta\right) - d\,xo^2\,\mu_b}\right)\right)} > 0$ /.
xo → 1 // FullSimplify

Out[∘]= $a\,\beta\,(a\,\beta + \mu_b)^2\,\left(a\,d\,\beta + \mu_b\,\left(d + 2\,a\,\beta - 2\,\sqrt{a\,\beta\,(-d + a\,\beta) - d\,\mu_b}\right)\right) < 0$

In[∘]:= $\left(a\,d\,\beta + \mu_b\,\left(d + 2\,a\,\beta - 2\,\sqrt{a\,\beta\,(-d + a\,\beta) - d\,\mu_b}\right)\right) < 0$ /. {$\mu_b$ → 0} // FullSimplify

Out[∘]= $a\,d\,\beta < 0$

```
(*This is not biologically feasible*)
```

In[∘]:= (*The first solution produces biologically feasible results.  *)

In[∘]:=

```
(* Singular strategies, eq. 32,
copied from the first solution of "sol" above  *)
```

In[∘]:= vrSol = $\frac{d + 2\,\mu_b - \sqrt{d^2 + 4\,d * \mu_b}}{2\,\mu_b}$;

vtSol = $\left(xo^{-c}\,\left(2\,\sqrt{d}\,xo\,\mu_b + \left(d\,xo - a\,c\,xo^c\,\beta\right)\,\left(\sqrt{d} + \sqrt{d + 4\,\mu_b}\right)\right)\right) /$
$\left(2\,a\,(-1 + c)\,d\,\beta\,\sqrt{d + 4\,\mu_b}\right)$;

In[ ]:=

```
(*Simplification of vtSol, eq. 32 second line *)
```

In[ ]:=

$$\text{FullSimplify}\Big[\text{PowerExpand}\Big[\frac{1}{(1-c)}\left(c*\left(\frac{\sqrt{d}+\sqrt{(d+4\mu_b)}}{2\,d\,\sqrt{(d+4\mu_b)}}\right)-\frac{xo}{\beta*a\,xo^c}*\frac{\left(d+2\mu_b+\sqrt{d\,(d+4\mu_b)}\right)}{2\sqrt{d\,(d+4\mu_b)}}\right)==\left(xo^{-c}\left(2\sqrt{d}\,xo\,\mu_b+\left(d\,xo-a\,c\,xo^c\,\beta\right)\left(\sqrt{d}+\sqrt{d+4\mu_b}\right)\right)\right)\Big/\left(2\,a\,(-1+c)\,d\,\beta\sqrt{d+4\mu_b}\right)\Big]\Big]$$

Out[ ]= True

#### 2.1.4. Singular mutation rate and population size

In[ ]:=

```
(*Uninvadable mutation rate*)
```

$$\text{mutSin}=\mu_b\,(1-\text{vrSol})^2\ \text{// FullSimplify}$$

Out[ ]=

$$\frac{\left(d-\sqrt{d\,(d+4\mu_b)}\right)^2}{4\mu_b}$$

In[ ]:= 

```
(*Proof of 33 *)
```

In[ ]:=

$$\text{FullSimplify}\left[\text{PowerExpand}\left[\frac{\left(d-\sqrt{d\,(d+4\mu_b)}\right)^2}{4\mu_b}==\text{mutSin}\right],\ \{d>0,\ \mu_b>0\}\right]$$

Out[ ]= True

In[ ]:=

```
(*Uninvdable population size, the equiation is too large,
so not given in the Manuscript *)
```

In[ ]:=
```
NSing = FullSimplify[NEquil[vt, vr] /. {vt → vtSol, vr → vrSol, αb → 2, α → 2}]
```

Out[ ]=

$$\frac{1}{\gamma}\Bigg(1+\Bigg(d\, e^{\frac{xo^{-c}\,(8\,\sqrt{d}\,xo\,\mu_b^2+(d\,xo-a\,c\,xo^c\,\beta)\,(\sqrt{d}+\sqrt{d+4\,\mu_b})\,(d-\sqrt{d\,(d+4\,\mu_b)})-2\,\mu_b\,(-3\,d^{3/2}\,xo-2\,d\,xo\,\sqrt{d+4\,\mu_b}-2\,a\,(-c+(-1+c)\,mf)\,xo^c\,\beta\,\sqrt{d+4\,\mu_b}+\sqrt{d}\,(2\,a\,c\,xo^c\,\beta+xo\,\sqrt{d\,(d}{4\,a\,(-1+c)\,\beta\,\mu_b\,\sqrt{d+4\,\mu_b}}}$$

$$\mu_b\,\left(d+4\,\mu_b-\sqrt{d\,(d+4\,\mu_b)}\right)$$

$$\Bigg(\Bigg(xo^{1-c}+\Bigg(xo^{-c}\,\left(2\,\sqrt{d}\,xo\,\mu_b+(d\,xo-a\,c\,xo^c\,\beta)\,\left(\sqrt{d}+\sqrt{d+4\,\mu_b}\right)\right)$$

$$\left(-1+\frac{1}{4\,\mu_b^2}\,\left(d+2\,\mu_b-\sqrt{d\,(d+4\,\mu_b)}\right)^2\right)\Bigg)\Big/\left(2\,d\,\sqrt{d+4\,\mu_b}\right)\Bigg)^{\frac{1}{1-c}}\Bigg)^{-c}\Bigg)\Big/$$

$$\left(a\,\left(d\,\left(d-\sqrt{d\,(d+4\,\mu_b)}\right)-2\,\mu_b\,\left(-2\,d+\sqrt{d\,(d+4\,\mu_b)}\right)\right)\right)\Bigg)$$

In[ ]:=
```
(*Uninvdable popualation size. Saving the solution here,
because re-calculation takes several minutes*)
```

In[ ]:= **NUninvdable =**

$$\text{NUninvdable} = \frac{1}{\gamma}\left(1+\left(d\, e^{\frac{xo^{-c}\left(8\sqrt{d}\,xo\,\mu_b^2+(d\,xo-a\,c\,xo^c\,\beta)\left(\sqrt{d}+\sqrt{d+4\mu_b}\right)\left(d-\sqrt{d(d+4\mu_b)}\right)-2\mu_b\left(-3d^{3/2}\,xo-2\,d\,xo\sqrt{d+4\mu_b}-2a(-c+(-1+c)\,mf)\,xo^c\,\beta\sqrt{d+4\mu_b}+\sqrt{d}\left(2\,a\,c\,xo^c\,\beta+xo\right.\right.\right.}{4a(-1+c)\beta\mu_b\sqrt{d+4\mu_b}}}\right.\right.$$

$$\mu_b\left(d+4\mu_b-\sqrt{d(d+4\mu_b)}\right)\left(\left(xo^{1-c}+\left(xo^{-c}\left(2\sqrt{d}\,xo\,\mu_b+(d\,xo-a\,c\,xo^c\,\beta)\left(\sqrt{d}+\sqrt{d+4\mu_b}\right)\right)\left(-1+\frac{1}{4\mu_b^2}\left(d+2\mu_b-\sqrt{d(d+4\mu_b)}\right)^2\right)\right)\Big/\left(2d\sqrt{d+4\mu_b}\right)\right)^{\frac{1}{1-c}}\right)^{-c}\Big/$$

$$\left.\left.\left(a\left(d\left(d-\sqrt{d(d+4\mu_b)}\right)-2\mu_b\left(-2d+\sqrt{d(d+4\mu_b)}\right)\right)\right)\right)\ \text{/. } \{\mu_b\to\mu\}$$

Out[ ]=

$$\frac{1}{\gamma}\left(1+\left(d\, e^{\frac{xo^{-c}\left(8\sqrt{d}\,xo\,\mu^2+(d\,xo-a\,c\,xo^c\,\beta)\left(\sqrt{d}+\sqrt{d+4\mu}\right)\left(d-\sqrt{d(d+4\mu)}\right)-2\mu\left(-3d^{3/2}\,xo-2\,d\,xo\sqrt{d+4\mu}-2a(-c+(-1+c)\,mf)\,xo^c\,\beta\sqrt{d+4\mu}+\sqrt{d}\left(2\,a\,c\,xo^c\,\beta+xo\sqrt{d(d+4\mu)}\right.\right.\right.}{4a(-1+c)\beta\mu\sqrt{d+4\mu}}}\right.\right.$$

$$\mu\left(d+4\mu-\sqrt{d(d+4\mu)}\right)\left(\left(xo^{1-c}+\frac{xo^{-c}\left(2\sqrt{d}\,xo\,\mu+(d\,xo-a\,c\,xo^c\,\beta)\left(\sqrt{d}+\sqrt{d+4\mu}\right)\right)\left(-1+\frac{\left(d+2\mu-\sqrt{d(d+4\mu)}\right)^2}{4\mu^2}\right)}{2d\sqrt{d+4\mu}}\right)^{\frac{1}{1-c}}\right.$$

$$\left.\left.\left(a\left(d\left(d-\sqrt{d(d+4\mu)}\right)-2\mu\left(-2d+\sqrt{d(d+4\mu)}\right)\right)\right)\right)$$

In[ ]:= **NClassicUninvadable = FullSimplify[NClassic[vt] /. vt → vtClassic]**

Out[ ]=

$$\frac{1-\frac{d\, e^{-\frac{d\,xo^{1-c}-a\,c\,\beta}{a\beta-a\,c\,\beta}}\left(\left(\frac{a\,c\,\beta}{d}\right)^{\frac{1}{1-c}}\right)^{-c}}{a}}{\gamma}$$

### 2.2. Second-order condition for uninvadability and convergence stability

#### 2.2.1. Uninvadability condition

```
(* Hessian *)
```

In[ ]:=
```
Hu[vt_, vr_] =
  FullSimplify[Assuming[μb > 0 && 0 < ur < 1 && a > 0 && α > 0 && αb > 0 && xo > 0 &&
     c > 0 && ts > 0 && γ > 0 && d > 0 && β > 0,
    Refine[D[Ro[ut, ur], {{ut, ur}, 2}] // FullSimplify]] /. {ut → vt, ur → vr}]
```

Out[ ]=

$$\Bigg\{\Bigg\{a\, e^{-mf-vt\,(d+(1-vr)^{\alpha}\mu_b)}\,(-1+vr^{\alpha b})\,\left(\left(xo^{1-c}+a\,(-1+c)\,(-1+vr^{\alpha b})\,vt\,\beta\right)^{\frac{1}{1-c}}\right)^{c}$$

$$(-1+N\,\gamma)\,\Bigg(d+\frac{2\,a\,c\,(-1+vr^{\alpha b})\,xo^{c}\,\beta}{xo+a\,(-1+c)\,(-1+vr^{\alpha b})\,vt\,xo^{c}\,\beta}+$$

$$(1-vr)^{\alpha}\,\mu_b+\left(a^2\,(-1+c)\,c\,(-1+vr^{\alpha b})^2\,xo^{2\,c}\,\beta^2\right)\Big/$$

$$\left(\left(xo+a\,(-1+c)\,(-1+vr^{\alpha b})\,vt\,xo^{c}\,\beta\right)^2\,(d+(1-vr)^{\alpha}\,\mu_b)\right)+\Big(a^2\,c^2\,(-1+vr^{\alpha b})^2$$

$$xo^{2\,c}\,\beta^2\Big)\Big/\left(\left(xo+a\,(-1+c)\,(-1+vr^{\alpha b})\,vt\,xo^{c}\,\beta\right)^2\,(d+(1-vr)^{\alpha}\,\mu_b)\right)\Bigg),$$

$$\Bigg(a\, e^{-mf-vt\,(d+(1-vr)^{\alpha}\mu_b)}\,\left(\left(xo^{1-c}+a\,(-1+c)\,(-1+vr^{\alpha b})\,vt\,\beta\right)^{\frac{1}{1-c}}\right)^{c}\,(-1+N\,\gamma)$$

$$\Big(vr^{-1+\alpha b}\,\alpha b\,(d+(1-vr)^{\alpha}\,\mu_b)\,\Big(-xo^2\,(d+(1-vr)^{\alpha}\,\mu_b)-$$

$$a\,(-1+vr^{\alpha b})\,xo^{1+c}\,\beta\,(2\,c+(-2+c)\,d\,vt+(-2+c)\,(1-vr)^{\alpha}\,vt\,\mu_b)+$$

$$a^2\,(-1+vr^{\alpha b})^2\,vt\,xo^{2\,c}\,\beta^2\,(c+(-1+c)\,d\,vt+(-1+c)\,(1-vr)^{\alpha}\,vt\,\mu_b)\Big)-$$

$$(1-vr)^{-1+\alpha}\,(-1+vr^{\alpha b})\,\alpha\,(xo+a\,(-1+c)\,(-1+vr^{\alpha b})\,vt\,xo^{c}\,\beta)\,\mu_b$$

$$\Big(d^2\,vt\,xo+a\,(-1+vr^{\alpha b})\,(c+c\,d\,vt+(-1+c)\,d^2\,vt^2)\,xo^{c}\,\beta+$$

$$(1-vr)^{\alpha}\,vt\,\mu_b\,\Big(2\,d\,xo+a\,(-1+vr^{\alpha b})\,(c+2\,(-1+c)\,d\,vt)\,xo^{c}\,\beta+$$

$$(1-vr)^{\alpha}\,(xo+a\,(-1+c)\,(-1+vr^{\alpha b})\,vt\,xo^{c}\,\beta)\,\mu_b\Big)\Big)\Big)\Bigg)\Big/$$

$$\left(\left(xo+a\,(-1+c)\,(-1+vr^{\alpha b})\,vt\,xo^{c}\,\beta\right)^2\,(d+(1-vr)^{\alpha}\,\mu_b)^2\right)\Bigg\},$$

$$\Bigg\{\Bigg(a\, e^{-mf-vt\,(d+(1-vr)^{\alpha}\mu_b)}\,\left(\left(xo^{1-c}+a\,(-1+c)\,(-1+vr^{\alpha b})\,vt\,\beta\right)^{\frac{1}{1-c}}\right)^{c}$$

$$(-1+N\,\gamma)\,\Big(vr^{-1+\alpha b}\,\alpha b\,(d+(1-vr)^{\alpha}\,\mu_b)\,\Big(-xo^2\,(d+(1-vr)^{\alpha}\,\mu_b)-$$

$$a\,(-1+vr^{\alpha b})\,xo^{1+c}\,\beta\,(2\,c+(-2+c)\,d\,vt+(-2+c)\,(1-vr)^{\alpha}\,vt\,\mu_b)+$$

$$a^2\,(-1+vr^{\alpha b})^2\,vt\,xo^{2\,c}\,\beta^2\,(c+(-1+c)\,d\,vt+(-1+c)\,(1-vr)^{\alpha}\,vt\,\mu_b)\Big)-$$

$$(1-vr)^{-1+\alpha}\,(-1+vr^{\alpha b})\,\alpha\,(xo+a\,(-1+c)\,(-1+vr^{\alpha b})\,vt\,xo^{c}\,\beta)\,\mu_b$$

$$\Big(d^2\,vt\,xo+a\,(-1+vr^{\alpha b})\,(c+c\,d\,vt+(-1+c)\,d^2\,vt^2)\,xo^{c}\,\beta+$$

$$(1-vr)^{\alpha}\,vt\,\mu_b\,\Big(2\,d\,xo+a\,(-1+vr^{\alpha b})\,(c+2\,(-1+c)\,d\,vt)\,xo^{c}\,\beta+$$

$$(1-vr)^{\alpha}\,(xo+a\,(-1+c)\,(-1+vr^{\alpha b})\,vt\,xo^{c}\,\beta)\,\mu_b\Big)\Big)\Big)\Bigg)\Big/$$

$$\left(\left(xo+a\,(-1+c)\,(-1+vr^{\alpha b})\,vt\,xo^{c}\,\beta\right)^2\,(d+(1-vr)^{\alpha}\,\mu_b)^2\right),$$

$$\frac{1}{(d+(1-vr)^{\alpha}\mu_b)^3}$$

$$a$$

$$e^{-mf-vt\,(d+(1-vr)^{\alpha}\mu_b)}$$

$$\left(\left(xo^{1-c}+a\,(-1+c)\,(-1+vr^{\alpha b})\,vt\,\beta\right)^{\frac{1}{1-c}}\right)^{c}$$

$$(-1+N\,\gamma)$$

$$\Big(2\,(1-vr)^{2\,(-1+\alpha)}\,(-1+vr^{\alpha b})\,\alpha^2\,\mu_b^2 -$$

$$(1-vr)^{-2+\alpha}\,(-1+vr^{\alpha b})\,(-1+\alpha)\,\alpha\,\mu_b\,(d+(1-vr)^{\alpha}\mu_b) +$$

$$2\,(1-vr)^{-1+\alpha}\,vr^{-1+\alpha b}\,\alpha\,\alpha b\,\mu_b\,(d+(1-vr)^{\alpha}\mu_b) -$$

$$\big(2\,a\,c\,(1-vr)^{-1+\alpha}\,vr^{-1+\alpha b}\,(-1+vr^{\alpha b})\,vt\,xo^{c}\,\alpha\,\alpha b\,\beta\,\mu_b\,(d+(1-vr)^{\alpha}\mu_b)\big)\,\Big/$$

$$\big(xo+a\,(-1+c)\,(-1+vr^{\alpha b})\,vt\,xo^{c}\,\beta\big)+2\,(1-vr)^{2\,(-1+\alpha)}\,(-1+vr^{\alpha b})$$

$$vt\,\alpha^2\,\mu_b^2\,(d+(1-vr)^{\alpha}\mu_b)+vr^{-2+\alpha b}\,(-1+\alpha b)\,\alpha b\,(d+(1-vr)^{\alpha}\mu_b)^2 +$$

$$\big(a^2\,(-1+c)\,c\,vr^{-2+2\,\alpha b}\,(-1+vr^{\alpha b})\,vt^2\,xo^{2\,c}\,\alpha b^2\,\beta^2\,(d+(1-vr)^{\alpha}\mu_b)^2\big)\,\Big/$$

$$\big(xo+a\,(-1+c)\,(-1+vr^{\alpha b})\,vt\,xo^{c}\,\beta\big)^2 +$$

$$\big(a^2\,c^2\,vr^{-2+2\,\alpha b}\,(-1+vr^{\alpha b})\,vt^2\,xo^{2\,c}\,\alpha b^2\,\beta^2\,(d+(1-vr)^{\alpha}\mu_b)^2\big)\,\Big/$$

$$\big(xo+a\,(-1+c)\,(-1+vr^{\alpha b})\,vt\,xo^{c}\,\beta\big)^2 -$$

$$\big(a\,c\,vr^{-2+\alpha b}\,(-1+vr^{\alpha b})\,vt\,xo^{c}\,(-1+\alpha b)\,\alpha b\,\beta\,(d+(1-vr)^{\alpha}\mu_b)^2\big)\,\Big/$$

$$\big(xo+a\,(-1+c)\,(-1+vr^{\alpha b})\,vt\,xo^{c}\,\beta\big) -$$

$$\big(2\,a\,c\,vr^{-2+2\,\alpha b}\,vt\,xo^{c}\,\alpha b^2\,\beta\,(d+(1-vr)^{\alpha}\mu_b)^2\big)\,\Big/$$

$$\big(xo+a\,(-1+c)\,(-1+vr^{\alpha b})\,vt\,xo^{c}\,\beta\big) -$$

$$(1-vr)^{-2+\alpha}\,(-1+vr^{\alpha b})\,vt\,(-1+\alpha)\,\alpha\,\mu_b\,(d+(1-vr)^{\alpha}\mu_b)^2 +$$

$$2\,(1-vr)^{-1+\alpha}\,vr^{-1+\alpha b}\,vt\,\alpha\,\alpha b\,\mu_b\,(d+(1-vr)^{\alpha}\mu_b)^2 -$$

$$\big(2\,a\,c\,(1-vr)^{-1+\alpha}\,vr^{-1+\alpha b}\,(-1+vr^{\alpha b})\,vt^2\,xo^{c}\,\alpha\,\alpha b\,\beta\,\mu_b\,(d+(1-vr)^{\alpha}\mu_b)^2\big)\,\Big/$$

$$\big(xo+a\,(-1+c)\,(-1+vr^{\alpha b})\,vt\,xo^{c}\,\beta\big) +$$

$$(1-vr)^{2\,(-1+\alpha)}\,(-1+vr^{\alpha b})\,vt^2\,\alpha^2\,\mu_b^2\,(d+(1-vr)^{\alpha}\mu_b)^2\Big)\}\}$$

In[ ]:= (*Recall the singular solution*)

$$\text{vrSol} = \frac{d + 2\,\mu_b - \sqrt{d^2 + 4\,d * \mu_b}}{2\,\mu_b};$$

$$\text{vtSol} = \left(xo^{-c}\left(2\sqrt{d}\;xo\,\mu_b + \left(d\,xo - a\,c\,xo^{c}\,\beta\right)\left(\sqrt{d} + \sqrt{d + 4\,\mu_b}\right)\right)\right) \Big/ \left(2\,a\,(-1 + c)\,d\,\beta\,\sqrt{d + 4\,\mu_b}\right);$$

$$\text{NUninvdable} = \frac{1}{\gamma}\left(1 - \left(2^{\alpha b}\,\mathbb{e}^{-\frac{-2\,d^{3/2}\,xo^{1-c}\,\mu_b - 2\,a\,(-1+c)\,d\,mf\,\beta\,\sqrt{d+4\,\mu_b} + d\,xo^{-c}\left(-d\,xo + a\,c\,xo^{c}\,\beta\right)\left(\sqrt{d}+\sqrt{d+4\,\mu_b}\right) + 2^{-\alpha}\,xo^{-c}\,\mu_b\left(\frac{-d+\sqrt{d\,(d+4\,\mu_b)}}{\mu_b}\right)^{\alpha}\left(-2\sqrt{d}\;xo\,\mu_b - \left(d\,xo - a\,c\,xo^{c}\,\beta\right)\left(\sqrt{d} + \sqrt{d+4\,[illegible]}\right.\right.}{2\,a\,(-1+c)\,d\,\beta\,\sqrt{d+4\,\mu_b}}}\left(d + 2^{-\alpha}\,\mu_b\left(\frac{-d + \sqrt{d\,(d + 4\,\mu_b)}}{\mu_b}\right)^{\alpha}\right)\left(\left(xo^{1-c} + \frac{1}{2\,d\,\sqrt{d + 4\,\mu_b}}\,xo^{-c}\left(2\sqrt{d}\;xo\,\mu_b + \left(d\,xo - a\,c\,xo^{c}\,\beta\right)\left(\sqrt{d} + \sqrt{d + 4\,\mu_b}\right)\right)\left(-1 + 2^{-\alpha b}\left(\frac{d + 2\,\mu_b - \sqrt{d\,(d + 4\,\mu_b)}}{\mu_b}\right)^{\alpha b}\right)\right)^{\frac{1}{1-c}}\right)^{-c}\right) \Big/ \left(a\left(2^{\alpha b} - \left(\frac{d + 2\,\mu_b - \sqrt{d\,(d + 4\,\mu_b)}}{\mu_b}\right)^{\alpha b}\right)\right)\right);$$

In[ ]:= HuEx2FullParam = Hu[vt, vr] /. {vt → vtSol, vr → vrSol, N → NUninvdable}

In[ ]:= HuEx2FullParam = HuEx2FullParam /. {$\mu_b$ → $\mu$}

Out[ ]=

$$\left\{\left\{-\left(\left(2^{\alpha b}\,\mathbb{e}^{-mf-\frac{-2\,d^{3/2}\,xo^{1-c}\,\mu - 2\,a\,(-1+c)\,d\,mf\,\beta\,\sqrt{d+4\,\mu} + d\,xo^{-c}\left(-d\,xo + a\,c\,xo^{c}\,\beta\right)\left(\sqrt{d}+\sqrt{d+4\,\mu}\right) + 2^{-\alpha}\,xo^{-c}\,\mu\left(\frac{-d+\sqrt{d\,(d+4\,\mu)}}{\mu}\right)^{\alpha}\left(-2\sqrt{d}\;xo\,\mu - \left(d\,xo - a\,c\,xo^{c}\,\beta\right)\left(\sqrt{d} + \sqrt{d+4\,\mu}\right)\right)}{2\,a\,(-1+c)\,d\,\beta\,\sqrt{d+4\,\mu}}}\left(d + 2^{-\alpha}\,\mu\left(\frac{-d + \sqrt{d\,(d + 4\,\mu)}}{\mu}\right)^{\alpha}\right)\left(-1 + 2^{-\alpha b}\left(\frac{d + 2\,\mu - \sqrt{d^2 + 4\,d\,\mu}}{\mu}\right)^{\alpha b}\right)\right.\right.\right.$$

$$\left(\left(xo^{1-c} + \frac{1}{2\,d\,\sqrt{d + 4\,\mu}}\,xo^{-c}\left(2\sqrt{d}\;xo\,\mu + \left(d\,xo - a\,c\,xo^{c}\,\beta\right)\left(\sqrt{d} + \sqrt{d + 4\,\mu}\right)\right)\right.\right.$$

$$\left(-1+2^{-\alpha b}\left(\frac{d+2\mu-\sqrt{d\,(d+4\mu)}}{\mu}\right)^{\alpha b}\right)\Bigg)^{\frac{1}{1-c}}\Bigg)^{-c}$$

$$\Bigg(\Bigg(xo^{1-c}+\frac{1}{2\,d\,\sqrt{d+4\mu}}\,xo^{-c}\,\left(2\,\sqrt{d}\,xo\,\mu+\left(d\,xo-a\,c\,xo^{c}\,\beta\right)\,\left(\sqrt{d}+\sqrt{d+4\mu}\right)\right)$$

$$\left(-1+2^{-\alpha b}\left(\frac{d+2\mu-\sqrt{d^{2}+4\,d\,\mu}}{\mu}\right)^{\alpha b}\right)\Bigg)^{\frac{1}{1-c}}\Bigg)^{c}$$

$$\left(d+\mu\left(1-\frac{d+2\mu-\sqrt{d^{2}+4\,d\,\mu}}{2\mu}\right)^{\alpha}+\Bigg(a^{2}\,(-1+c)\,c\,xo^{2c}\,\beta^{2}\right.$$

$$\left(-1+2^{-\alpha b}\left(\frac{d+2\mu-\sqrt{d^{2}+4\,d\,\mu}}{\mu}\right)^{\alpha b}\right)^{2}\Bigg)\Bigg/\Bigg(\left(d+\mu\left(1-\frac{d+2\mu-\sqrt{d^{2}+4\,d\,\mu}}{2\mu}\right)^{\alpha}\right)$$

$$\Bigg(xo+\frac{1}{2\,d\,\sqrt{d+4\mu}}\,\left(2\,\sqrt{d}\,xo\,\mu+\left(d\,xo-a\,c\,xo^{c}\,\beta\right)\,\left(\sqrt{d}+\sqrt{d+4\mu}\right)\right)$$

$$\left(-1+2^{-\alpha b}\left(\frac{d+2\mu-\sqrt{d^{2}+4\,d\,\mu}}{\mu}\right)^{\alpha b}\right)\Bigg)^{2}\Bigg)+$$

$$\left(a^{2}\,c^{2}\,xo^{2c}\,\beta^{2}\left(-1+2^{-\alpha b}\left(\frac{d+2\mu-\sqrt{d^{2}+4\,d\,\mu}}{\mu}\right)^{\alpha b}\right)^{2}\right)\Bigg/$$

$$\Bigg(\left(d+\mu\left(1-\frac{d+2\mu-\sqrt{d^{2}+4\,d\,\mu}}{2\mu}\right)^{\alpha}\right)$$

$$\Bigg(xo+\frac{1}{2\,d\,\sqrt{d+4\mu}}\,\left(2\,\sqrt{d}\,xo\,\mu+\left(d\,xo-a\,c\,xo^{c}\,\beta\right)\,\left(\sqrt{d}+\sqrt{d+4\mu}\right)\right)$$

$$\left(-1+2^{-\alpha b}\left(\frac{d+2\mu-\sqrt{d^{2}+4\,d\,\mu}}{\mu}\right)^{\alpha b}\right)\Bigg)^{2}\Bigg)+$$

$$\left(2\,a\,c\,xo^{c}\,\beta\left(-1+2^{-\alpha b}\left(\frac{d+2\mu-\sqrt{d^{2}+4\,d\,\mu}}{\mu}\right)^{\alpha b}\right)\right)\Bigg/$$

$$\Bigg(xo+\Bigg(\left(2\,\sqrt{d}\,xo\,\mu+\left(d\,xo-a\,c\,xo^{c}\,\beta\right)\,\left(\sqrt{d}+\sqrt{d+4\mu}\right)\right)$$

$$\left(-1+2^{-\alpha b}\left(\frac{d+2\mu-\sqrt{d^{2}+4\,d\,\mu}}{\mu}\right)^{\alpha b}\right)\Bigg)\Bigg/\left(2\,d\,\sqrt{d+4\mu}\right)\Bigg)\Bigg)\Bigg)\Bigg/$$

$$\left(2^{\alpha b}-\left(\frac{d+2\,\mu-\sqrt{d\,(d+4\,\mu)}}{\mu}\right)^{\alpha b}\right)\Bigg),\ -\Bigg(\Bigg(\Bigg(2^{\alpha b}$$

$$e^{-mf-\frac{-2\,d^{3/2}\,xo^{1-c}\,\mu-2\,a\,(-1+c)\,d\,mf\,\beta\,\sqrt{d+4\,\mu}+d\,xo^{-c}\,(-d\,xo+a\,c\,xo^{c}\,\beta)\,\left(\sqrt{d}+\sqrt{d+4\,\mu}\right)+2^{-\alpha}\,xo^{-c}\,\mu\,\left(\frac{-d+\sqrt{d\,(d+4\,\mu)}}{\mu}\right)^{\alpha}\left(-2\,\sqrt{d}\,xo\,\mu-(d\,xo-a\,c\,xo^{c}\,\beta)\,\left(\sqrt{d}+\sqrt{d+4\,\mu}\right)\right)}{2\,a\,(-1+c)\,d\,\beta\,\sqrt{d+4\,\mu}}}$$

$$\left(d+2^{-\alpha}\,\mu\,\left(\frac{-d+\sqrt{d\,(d+4\,\mu)}}{\mu}\right)^{\alpha}\right)$$

$$\Bigg(\Bigg(xo^{1-c}+\frac{1}{2\,d\,\sqrt{d+4\,\mu}}\,xo^{-c}\,\left(2\,\sqrt{d}\,xo\,\mu+(d\,xo-a\,c\,xo^{c}\,\beta)\,\left(\sqrt{d}+\sqrt{d+4\,\mu}\right)\right)$$

$$\left(-1+2^{-\alpha b}\,\left(\frac{d+2\,\mu-\sqrt{d\,(d+4\,\mu)}}{\mu}\right)^{\alpha b}\right)\Bigg)^{\frac{1}{1-c}}\Bigg)^{-c}$$

$$\Bigg(\Bigg(xo^{1-c}+\frac{1}{2\,d\,\sqrt{d+4\,\mu}}\,xo^{-c}\,\left(2\,\sqrt{d}\,xo\,\mu+(d\,xo-a\,c\,xo^{c}\,\beta)\,\left(\sqrt{d}+\sqrt{d+4\,\mu}\right)\right)$$

$$\left(-1+2^{-\alpha b}\,\left(\frac{d+2\,\mu-\sqrt{d^{2}+4\,d\,\mu}}{\mu}\right)^{\alpha b}\right)\Bigg)^{\frac{1}{1-c}}\Bigg)^{c}$$

$$\Bigg(2^{1-\alpha b}\,\alpha b\,\left(\frac{d+2\,\mu-\sqrt{d^{2}+4\,d\,\mu}}{\mu}\right)^{-1+\alpha b}\,\left(d+\mu\,\left(1-\frac{d+2\,\mu-\sqrt{d^{2}+4\,d\,\mu}}{2\,\mu}\right)^{\alpha}\right)$$

$$\Bigg(-xo^{2}\,\left(d+\mu\,\left(1-\frac{d+2\,\mu-\sqrt{d^{2}+4\,d\,\mu}}{2\,\mu}\right)^{\alpha}\right)+\Bigg(a\,xo^{c}\,\beta\,\Big(2\,\sqrt{d}\,xo\,\mu+\Big(d\,xo-$$

$$a\,c\,xo^{c}\,\beta\Big)\,\left(\sqrt{d}+\sqrt{d+4\,\mu}\right)\Big)\,\left(-1+2^{-\alpha b}\,\left(\frac{d+2\,\mu-\sqrt{d^{2}+4\,d\,\mu}}{\mu}\right)^{\alpha b}\right)^{2}$$

$$\Bigg(c+\left(xo^{-c}\,\left(2\,\sqrt{d}\,xo\,\mu+(d\,xo-a\,c\,xo^{c}\,\beta)\,\left(\sqrt{d}+\sqrt{d+4\,\mu}\right)\right)\right)\Big/$$

$$\left(2\,a\,\beta\,\sqrt{d+4\,\mu}\right)+\frac{1}{2\,a\,d\,\beta\,\sqrt{d+4\,\mu}}\,xo^{-c}\,\mu\,\Big(2\,\sqrt{d}\,xo\,\mu+$$

$$(d\,xo-a\,c\,xo^{c}\,\beta)\,\left(\sqrt{d}+\sqrt{d+4\,\mu}\right)\Big)\,\left(1-\frac{d+2\,\mu-\sqrt{d^{2}+4\,d\,\mu}}{2\,\mu}\right)^{\alpha}\Bigg)\Bigg)\Big/$$

$$\left(2\,(-1+c)\,d\,\sqrt{d+4\,\mu}\right)-a\,xo^{1+c}\,\beta\,\left(-1+2^{-\alpha b}\,\left(\frac{d+2\,\mu-\sqrt{d^{2}+4\,d\,\mu}}{\mu}\right)^{\alpha b}\right)$$

$$\Bigg(2\,c+\left((-2+c)\,xo^{-c}\,\left(2\,\sqrt{d}\,xo\,\mu+(d\,xo-a\,c\,xo^{c}\,\beta)\,\left(\sqrt{d}+\sqrt{d+4\,\mu}\right)\right)\right)\Big/$$

$$(2\, a\, (-1 + c)\, \beta\, \sqrt{d + 4\,\mu}\,) +$$

$$\Big((-2 + c)\, xo^{-c}\, \mu\, \big(2\, \sqrt{d}\, xo\, \mu + (d\, xo - a\, c\, xo^{c}\, \beta)\, (\sqrt{d} + \sqrt{d + 4\,\mu}\,)\big)$$

$$\left(1 - \frac{d + 2\,\mu - \sqrt{d^{2} + 4\, d\, \mu}}{2\,\mu}\right)^{\alpha}\Big) \Big/ \; (2\, a\, (-1 + c)\, d\, \beta\, \sqrt{d + 4\,\mu}\,)\Big)\Big) -$$

$$\alpha\, \mu\, \left(1 - \frac{d + 2\,\mu - \sqrt{d^{2} + 4\, d\, \mu}}{2\,\mu}\right)^{-1+\alpha} \left(-1 + 2^{-\alpha b} \left(\frac{d + 2\,\mu - \sqrt{d^{2} + 4\, d\, \mu}}{\mu}\right)^{\alpha b}\right)$$

$$\Big(xo + \frac{1}{2\, d\, \sqrt{d + 4\,\mu}}\, \big(2\, \sqrt{d}\, xo\, \mu + (d\, xo - a\, c\, xo^{c}\, \beta)\, (\sqrt{d} + \sqrt{d + 4\,\mu}\,)\big)$$

$$\left(-1 + 2^{-\alpha b} \left(\frac{d + 2\,\mu - \sqrt{d^{2} + 4\, d\, \mu}}{\mu}\right)^{\alpha b}\right)\Big)\Big)$$

$$\Big(\big(d\, xo^{1-c}\, \big(2\, \sqrt{d}\, xo\, \mu + (d\, xo - a\, c\, xo^{c}\, \beta)\, (\sqrt{d} + \sqrt{d + 4\,\mu}\,)\big)\big) \Big/$$

$$(2\, a\, (-1 + c)\, \beta\, \sqrt{d + 4\,\mu}\,) + a\, xo^{c}\, \beta \left(-1 + 2^{-\alpha b} \left(\frac{d + 2\,\mu - \sqrt{d^{2} + 4\, d\, \mu}}{\mu}\right)^{\alpha b}\right)$$

$$\Big(c + \big(c\, xo^{-c}\, \big(2\, \sqrt{d}\, xo\, \mu + (d\, xo - a\, c\, xo^{c}\, \beta)\, (\sqrt{d} + \sqrt{d + 4\,\mu}\,)\big)\big) \Big/$$

$$(2\, a\, (-1 + c)\, \beta\, \sqrt{d + 4\,\mu}\,) +$$

$$\Big(xo^{-2\, c}\, \big(2\, \sqrt{d}\, xo\, \mu + (d\, xo - a\, c\, xo^{c}\, \beta)\, (\sqrt{d} + \sqrt{d + 4\,\mu}\,)\big)^{2}\Big) \Big/$$

$$(4\, a^{2}\, (-1 + c)\, \beta^{2}\, (d + 4\,\mu))\Big)\Big) + \Big(xo^{-c}\, \mu\, \big(2\, \sqrt{d}\, xo\, \mu +$$

$$(d\, xo - a\, c\, xo^{c}\, \beta)\, (\sqrt{d} + \sqrt{d + 4\,\mu}\,)\big) \left(1 - \frac{d + 2\,\mu - \sqrt{d^{2} + 4\, d\, \mu}}{2\,\mu}\right)^{\alpha}$$

$$\left(2\, d\, xo + a\, xo^{c}\, \beta \left(-1 + 2^{-\alpha b} \left(\frac{d + 2\,\mu - \sqrt{d^{2} + 4\, d\, \mu}}{\mu}\right)^{\alpha b}\right)\right.$$

$$\Big(c + \big(xo^{-c}\, \big(2\, \sqrt{d}\, xo\, \mu + (d\, xo - a\, c\, xo^{c}\, \beta)\, (\sqrt{d} + \sqrt{d + 4\,\mu}\,)\big)\big) \Big/$$

$$(a\, \beta\, \sqrt{d + 4\,\mu}\,)\Big)\Big) + \mu \left(1 - \frac{d + 2\,\mu - \sqrt{d^{2} + 4\, d\, \mu}}{2\,\mu}\right)^{\alpha}$$

$$\Big(xo + \frac{1}{2\, d\, \sqrt{d + 4\,\mu}}\, \big(2\, \sqrt{d}\, xo\, \mu + (d\, xo - a\, c\, xo^{c}\, \beta)\, (\sqrt{d} + \sqrt{d + 4\,\mu}\,)\big)$$

$$\left(-1 + 2^{-\alpha b} \left(\frac{d + 2\,\mu - \sqrt{d^{2} + 4\, d\, \mu}}{\mu}\right)^{\alpha b}\right)\Big)\Big)\Big)\Big) \Big/$$

$$(2 a (-1+c) d \beta \sqrt{d+4\mu}))))\Big/ \left(\left(2^{\alpha b} - \left(\frac{d+2\mu-\sqrt{d(d+4\mu)}}{\mu}\right)^{\alpha b}\right)\right.$$

$$\left(d+\mu\left(1-\frac{d+2\mu-\sqrt{d^2+4d\mu}}{2\mu}\right)^{\alpha}\right)^2$$

$$\left(xo+\frac{1}{2d\sqrt{d+4\mu}}\,(2\sqrt{d}\,xo\,\mu+(d\,xo-a\,c\,xo^c\,\beta)(\sqrt{d}+\sqrt{d+4\mu}))\right.$$

$$\left.\left.\left.\left(-1+2^{-\alpha b}\left(\frac{d+2\mu-\sqrt{d^2+4d\mu}}{\mu}\right)^{\alpha b}\right)\right)^2\right)\right)\Big\}, \Big\{-\left(\left(2^{\alpha b}\right.\right.$$

$$e^{-mf-\frac{-2 d^{3/2} xo^{1-c}\mu-2a(-1+c)d\,mf\,\beta\sqrt{d+4\mu}+d\,xo^{-c}(-d\,xo+a\,c\,xo^c\beta)(\sqrt{d}+\sqrt{d+4\mu})+2^{-\alpha}xo^{-c}\mu\left(\frac{-d+\sqrt{d(d+4\mu)}}{\mu}\right)^{\alpha}(-2\sqrt{d}\,xo\,\mu-(d\,xo-a\,c\,xo^c\beta)(\sqrt{d}+\sqrt{d+4\mu}))}{2a(-1+c)d\beta\sqrt{d+4\mu}}}$$

$$\left(d+2^{-\alpha}\mu\left(\frac{-d+\sqrt{d(d+4\mu)}}{\mu}\right)^{\alpha}\right)$$

$$\left(\left(xo^{1-c}+\frac{1}{2d\sqrt{d+4\mu}}\,xo^{-c}\,(2\sqrt{d}\,xo\,\mu+(d\,xo-a\,c\,xo^c\,\beta)(\sqrt{d}+\sqrt{d+4\mu}))\right.\right.$$

$$\left.\left.\left(-1+2^{-\alpha b}\left(\frac{d+2\mu-\sqrt{d(d+4\mu)}}{\mu}\right)^{\alpha b}\right)\right)^{\frac{1}{1-c}}\right)^{-c}$$

$$\left(\left(xo^{1-c}+\frac{1}{2d\sqrt{d+4\mu}}\,xo^{-c}\,(2\sqrt{d}\,xo\,\mu+(d\,xo-a\,c\,xo^c\,\beta)(\sqrt{d}+\sqrt{d+4\mu}))\right.\right.$$

$$\left.\left.\left(-1+2^{-\alpha b}\left(\frac{d+2\mu-\sqrt{d^2+4d\mu}}{\mu}\right)^{\alpha b}\right)\right)^{\frac{1}{1-c}}\right)^{c}$$

$$\left(2^{1-\alpha b}\,\alpha b\left(\frac{d+2\mu-\sqrt{d^2+4d\mu}}{\mu}\right)^{-1+\alpha b}\left(d+\mu\left(1-\frac{d+2\mu-\sqrt{d^2+4d\mu}}{2\mu}\right)^{\alpha}\right)\right.$$

$$\left(-xo^2\left(d+\mu\left(1-\frac{d+2\mu-\sqrt{d^2+4d\mu}}{2\mu}\right)^{\alpha}\right)+\left(a\,xo^c\,\beta\,(2\sqrt{d}\,xo\,\mu+(d\,xo-\right.\right.$$

$$\left.a\,c\,xo^c\,\beta)(\sqrt{d}+\sqrt{d+4\mu}))\left(-1+2^{-\alpha b}\left(\frac{d+2\mu-\sqrt{d^2+4d\mu}}{\mu}\right)^{\alpha b}\right)^2\right.$$

$$\left(c+(xo^{-c}\,(2\sqrt{d}\,xo\,\mu+(d\,xo-a\,c\,xo^c\,\beta)(\sqrt{d}+\sqrt{d+4\mu})))\right)\Big/$$

$$(2a\beta\sqrt{d+4\mu})+\frac{1}{2ad\beta\sqrt{d+4\mu}}\,xo^{-c}\,\mu\,(2\sqrt{d}\,xo\,\mu+$$

$$(d\,xo - a\,c\,xo^{c}\,\beta)\;(\sqrt{d} + \sqrt{d+4\mu}))\;\left(1 - \frac{d+2\mu-\sqrt{d^2+4\,d\,\mu}}{2\mu}\right)^{\alpha}))\;\Big/$$

$$(2\;(-1+c)\;d\;\sqrt{d+4\mu}) - a\,xo^{1+c}\,\beta\,\left(-1+2^{-\alpha b}\left(\frac{d+2\mu-\sqrt{d^2+4\,d\,\mu}}{\mu}\right)^{\alpha b}\right)$$

$$(2\,c + ((-2+c)\;xo^{-c}\;(2\,\sqrt{d}\;xo\,\mu + (d\,xo - a\,c\,xo^{c}\,\beta)\;(\sqrt{d} + \sqrt{d+4\mu})))\;/$$

$$(2\,a\;(-1+c)\;\beta\,\sqrt{d+4\mu}) +$$

$$((-2+c)\;xo^{-c}\,\mu\,(2\,\sqrt{d}\;xo\,\mu + (d\,xo - a\,c\,xo^{c}\,\beta)\;(\sqrt{d} + \sqrt{d+4\mu}))$$

$$\left(1 - \frac{d+2\mu-\sqrt{d^2+4\,d\,\mu}}{2\mu}\right)^{\alpha})\Big/\;(2\,a\;(-1+c)\;d\,\beta\,\sqrt{d+4\mu}))) -$$

$$\alpha\,\mu\,\left(1 - \frac{d+2\mu-\sqrt{d^2+4\,d\,\mu}}{2\mu}\right)^{-1+\alpha}\left(-1+2^{-\alpha b}\left(\frac{d+2\mu-\sqrt{d^2+4\,d\,\mu}}{\mu}\right)^{\alpha b}\right)$$

$$(xo + \frac{1}{2\,d\,\sqrt{d+4\mu}}\;(2\,\sqrt{d}\;xo\,\mu + (d\,xo - a\,c\,xo^{c}\,\beta)\;(\sqrt{d} + \sqrt{d+4\mu}))$$

$$\left(-1+2^{-\alpha b}\left(\frac{d+2\mu-\sqrt{d^2+4\,d\,\mu}}{\mu}\right)^{\alpha b}\right))$$

$$((d\,xo^{1-c}\;(2\,\sqrt{d}\;xo\,\mu + (d\,xo - a\,c\,xo^{c}\,\beta)\;(\sqrt{d} + \sqrt{d+4\mu})))\;/$$

$$(2\,a\;(-1+c)\;\beta\,\sqrt{d+4\mu}) + a\,xo^{c}\,\beta\,\left(-1+2^{-\alpha b}\left(\frac{d+2\mu-\sqrt{d^2+4\,d\,\mu}}{\mu}\right)^{\alpha b}\right)$$

$$(c + (c\,xo^{-c}\;(2\,\sqrt{d}\;xo\,\mu + (d\,xo - a\,c\,xo^{c}\,\beta)\;(\sqrt{d} + \sqrt{d+4\mu})))\;/$$

$$(2\,a\;(-1+c)\;\beta\,\sqrt{d+4\mu}) +$$

$$(xo^{-2\,c}\;(2\,\sqrt{d}\;xo\,\mu + (d\,xo - a\,c\,xo^{c}\,\beta)\;(\sqrt{d} + \sqrt{d+4\mu}))^{2})\;/$$

$$(4\,a^2\;(-1+c)\;\beta^2\;(d+4\mu))) + (xo^{-c}\,\mu\,(2\,\sqrt{d}\;xo\,\mu +$$

$$(d\,xo - a\,c\,xo^{c}\,\beta)\;(\sqrt{d} + \sqrt{d+4\mu}))\;\left(1 - \frac{d+2\mu-\sqrt{d^2+4\,d\,\mu}}{2\mu}\right)^{\alpha}$$

$$(2\,d\,xo + a\,xo^{c}\,\beta\,\left(-1+2^{-\alpha b}\left(\frac{d+2\mu-\sqrt{d^2+4\,d\,\mu}}{\mu}\right)^{\alpha b}\right)$$

$$(c + (xo^{-c}\;(2\,\sqrt{d}\;xo\,\mu + (d\,xo - a\,c\,xo^{c}\,\beta)\;(\sqrt{d} + \sqrt{d+4\mu})))\;/$$

$$\left(a\,\beta\,\sqrt{d+4\,\mu}\,\right)\Big)+\mu\left(1-\frac{d+2\,\mu-\sqrt{d^2+4\,d\,\mu}}{2\,\mu}\right)^{\alpha}$$

$$\Bigg(xo+\frac{1}{2\,d\,\sqrt{d+4\,\mu}}\,\left(2\,\sqrt{d}\,\,xo\,\mu+\left(d\,xo-a\,c\,xo^{c}\,\beta\right)\,\left(\sqrt{d}\,+\sqrt{d+4\,\mu}\,\right)\right)$$

$$\left(-1+2^{-\alpha b}\,\left(\frac{d+2\,\mu-\sqrt{d^2+4\,d\,\mu}}{\mu}\right)^{\alpha b}\right)\Bigg)\Bigg)\Bigg)\Bigg)\Bigg/$$

$$\left(2\,a\,\left(-1+c\right)\,d\,\beta\,\sqrt{d+4\,\mu}\,\right)\Bigg)\Bigg)\Bigg)\Bigg/\;\Bigg(\Bigg(2^{\alpha b}-\left(\frac{d+2\,\mu-\sqrt{d\,\left(d+4\,\mu\right)}}{\mu}\right)^{\alpha b}\Bigg)$$

$$\left(d+\mu\,\left(1-\frac{d+2\,\mu-\sqrt{d^2+4\,d\,\mu}}{2\,\mu}\right)^{\alpha}\right)^{2}$$

$$\Bigg(xo+\frac{1}{2\,d\,\sqrt{d+4\,\mu}}\,\left(2\,\sqrt{d}\,\,xo\,\mu+\left(d\,xo-a\,c\,xo^{c}\,\beta\right)\,\left(\sqrt{d}\,+\sqrt{d+4\,\mu}\,\right)\right)$$

$$\left(-1+2^{-\alpha b}\,\left(\frac{d+2\,\mu-\sqrt{d^2+4\,d\,\mu}}{\mu}\right)^{\alpha b}\right)\Bigg)^{2}\Bigg)\Bigg)\,,\;-\Bigg(\Bigg(2^{\alpha b}$$

$$e^{-mf-\frac{-2\,d^{3/2}\,xo^{1-c}\,\mu-2\,a\,(-1+c)\,d\,mf\,\beta\,\sqrt{d+4\,\mu}\,+d\,xo^{-c}\,\left(-d\,xo+a\,c\,xo^{c}\,\beta\right)\,\left(\sqrt{d}\,+\sqrt{d+4\,\mu}\,\right)+2^{-\alpha}\,xo^{-c}\,\mu\,\left(\frac{-d+\sqrt{d\,(d+4\,\mu)}}{\mu}\right)^{\alpha}\,\left(-2\,\sqrt{d}\,\,xo\,\mu-\left(d\,xo-a\,c\,xo^{c}\,\beta\right)\,\left(\sqrt{d}\,+\sqrt{d+4\,\mu}\,\right)\right)}{2\,a\,(-1+c)\,d\,\beta\,\sqrt{d+4\,\mu}}}$$

$$\left(d+2^{-\alpha}\,\mu\,\left(\frac{-d+\sqrt{d\,\left(d+4\,\mu\right)}}{\mu}\right)^{\alpha}\right)$$

$$\Bigg(\Bigg(xo^{1-c}+\frac{1}{2\,d\,\sqrt{d+4\,\mu}}\,xo^{-c}\,\left(2\,\sqrt{d}\,\,xo\,\mu+\left(d\,xo-a\,c\,xo^{c}\,\beta\right)\,\left(\sqrt{d}\,+\sqrt{d+4\,\mu}\,\right)\right)$$

$$\left(-1+2^{-\alpha b}\,\left(\frac{d+2\,\mu-\sqrt{d\,\left(d+4\,\mu\right)}}{\mu}\right)^{\alpha b}\right)\Bigg)^{\frac{1}{1-c}}\Bigg)^{-c}$$

$$\Bigg(\Bigg(xo^{1-c}+\frac{1}{2\,d\,\sqrt{d+4\,\mu}}\,xo^{-c}\,\left(2\,\sqrt{d}\,\,xo\,\mu+\left(d\,xo-a\,c\,xo^{c}\,\beta\right)\,\left(\sqrt{d}\,+\sqrt{d+4\,\mu}\,\right)\right)$$

$$\left(-1+2^{-\alpha b}\,\left(\frac{d+2\,\mu-\sqrt{d^2+4\,d\,\mu}}{\mu}\right)^{\alpha b}\right)\Bigg)^{\frac{1}{1-c}}\Bigg)^{c}$$

$$\Bigg(2\,\alpha^2\,\mu^2\,\left(1-\frac{d+2\,\mu-\sqrt{d^2+4\,d\,\mu}}{2\,\mu}\right)^{2\,(-1+\alpha)}\,\left(-1+2^{-\alpha b}\,\left(\frac{d+2\,\mu-\sqrt{d^2+4\,d\,\mu}}{\mu}\right)^{\alpha b}\right)+$$

$$2^{2-\alpha b}\,\alpha\,\alpha b\,\mu\left(\frac{d+2\,\mu-\sqrt{d^2+4\,d\,\mu}}{\mu}\right)^{-1+\alpha b}\left(1-\frac{d+2\,\mu-\sqrt{d^2+4\,d\,\mu}}{2\,\mu}\right)^{-1+\alpha}$$

$$\left(d+\mu\left(1-\frac{d+2\,\mu-\sqrt{d^2+4\,d\,\mu}}{2\,\mu}\right)^{\alpha}\right)-(-1+\alpha)\,\alpha\,\mu\left(1-\frac{d+2\,\mu-\sqrt{d^2+4\,d\,\mu}}{2\,\mu}\right)^{-2+\alpha}$$

$$\left(-1+2^{-\alpha b}\left(\frac{d+2\,\mu-\sqrt{d^2+4\,d\,\mu}}{\mu}\right)^{\alpha b}\right)\left(d+\mu\left(1-\frac{d+2\,\mu-\sqrt{d^2+4\,d\,\mu}}{2\,\mu}\right)^{\alpha}\right)+$$

$$\Bigg(xo^{-c}\,\alpha^2\,\mu^2\left(2\,\sqrt{d}\,\,xo\,\mu+\left(d\,xo-a\,c\,xo^{c}\,\beta\right)\left(\sqrt{d}+\sqrt{d+4\,\mu}\right)\right)$$

$$\left(1-\frac{d+2\,\mu-\sqrt{d^2+4\,d\,\mu}}{2\,\mu}\right)^{2\,(-1+\alpha)}\left(-1+2^{-\alpha b}\left(\frac{d+2\,\mu-\sqrt{d^2+4\,d\,\mu}}{\mu}\right)^{\alpha b}\right)$$

$$\left(d+\mu\left(1-\frac{d+2\,\mu-\sqrt{d^2+4\,d\,\mu}}{2\,\mu}\right)^{\alpha}\right)\Bigg)\Bigg/\left(a\,(-1+c)\,d\,\beta\,\sqrt{d+4\,\mu}\right)+$$

$$2^{2-\alpha b}\,(-1+\alpha b)\,\alpha b\left(\frac{d+2\,\mu-\sqrt{d^2+4\,d\,\mu}}{\mu}\right)^{-2+\alpha b}\left(d+\mu\left(1-\frac{d+2\,\mu-\sqrt{d^2+4\,d\,\mu}}{2\,\mu}\right)^{\alpha}\right)^2+$$

$$\Bigg(2^{1-\alpha b}\,xo^{-c}\,\alpha\,\alpha b\,\mu\left(\frac{d+2\,\mu-\sqrt{d^2+4\,d\,\mu}}{\mu}\right)^{-1+\alpha b}$$

$$\left(2\,\sqrt{d}\,\,xo\,\mu+\left(d\,xo-a\,c\,xo^{c}\,\beta\right)\left(\sqrt{d}+\sqrt{d+4\,\mu}\right)\right)\left(1-\frac{d+2\,\mu-\sqrt{d^2+4\,d\,\mu}}{2\,\mu}\right)^{-1+\alpha}$$

$$\left(d+\mu\left(1-\frac{d+2\,\mu-\sqrt{d^2+4\,d\,\mu}}{2\,\mu}\right)^{\alpha}\right)^2\Bigg)\Bigg/\left(a\,(-1+c)\,d\,\beta\,\sqrt{d+4\,\mu}\right)-$$

$$\Bigg(xo^{-c}\,(-1+\alpha)\,\alpha\,\mu\left(2\,\sqrt{d}\,\,xo\,\mu+\left(d\,xo-a\,c\,xo^{c}\,\beta\right)\left(\sqrt{d}+\sqrt{d+4\,\mu}\right)\right)$$

$$\left(1-\frac{d+2\,\mu-\sqrt{d^2+4\,d\,\mu}}{2\,\mu}\right)^{-2+\alpha}\left(-1+2^{-\alpha b}\left(\frac{d+2\,\mu-\sqrt{d^2+4\,d\,\mu}}{\mu}\right)^{\alpha b}\right)$$

$$\left(d+\mu\left(1-\frac{d+2\,\mu-\sqrt{d^2+4\,d\,\mu}}{2\,\mu}\right)^{\alpha}\right)^2\Bigg)\Bigg/\left(2\,a\,(-1+c)\,d\,\beta\,\sqrt{d+4\,\mu}\right)+$$

$$\Bigg(xo^{-2\,c}\,\alpha^2\,\mu^2\left(2\,\sqrt{d}\,\,xo\,\mu+\left(d\,xo-a\,c\,xo^{c}\,\beta\right)\left(\sqrt{d}+\sqrt{d+4\,\mu}\right)\right)^2$$

$$\left(1-\frac{d+2\,\mu-\sqrt{d^2+4\,d\,\mu}}{2\,\mu}\right)^{2\,(-1+\alpha)}\left(-1+2^{-\alpha b}\left(\frac{d+2\,\mu-\sqrt{d^2+4\,d\,\mu}}{\mu}\right)^{\alpha b}\right)$$

$$\left(d+\mu\left(1-\frac{d+2\mu-\sqrt{d^2+4d\mu}}{2\mu}\right)^{\alpha}\right)^2\Bigg/$$

$$\left(4a^2(-1+c)^2 d^2\beta^2(d+4\mu)\right)+\Bigg(2^{-2\alpha b}c\,\alpha b^2\left(\frac{d+2\mu-\sqrt{d^2+4d\mu}}{\mu}\right)^{-2+2\alpha b}$$

$$\left(2\sqrt{d}\,xo\,\mu+\left(d\,xo-a\,c\,xo^c\beta\right)\left(\sqrt{d}+\sqrt{d+4\mu}\right)\right)^2$$

$$\left(-1+2^{-\alpha b}\left(\frac{d+2\mu-\sqrt{d^2+4d\mu}}{\mu}\right)^{\alpha b}\right)\left(d+\mu\left(1-\frac{d+2\mu-\sqrt{d^2+4d\mu}}{2\mu}\right)^{\alpha}\right)^2\Bigg)\Bigg/$$

$$\Bigg((-1+c)\,d^2\,(d+4\mu)\Bigg(xo+\frac{1}{2d\sqrt{d+4\mu}}\left(2\sqrt{d}\,xo\,\mu+\left(d\,xo-a\,c\,xo^c\beta\right)\right.$$

$$\left.\left(\sqrt{d}+\sqrt{d+4\mu}\right)\right)\left(-1+2^{-\alpha b}\left(\frac{d+2\mu-\sqrt{d^2+4d\mu}}{\mu}\right)^{\alpha b}\right)\Bigg)^2\Bigg)+$$

$$\Bigg(2^{-2\alpha b}c^2\alpha b^2\left(\frac{d+2\mu-\sqrt{d^2+4d\mu}}{\mu}\right)^{-2+2\alpha b}\left(2\sqrt{d}\,xo\,\mu+\left(d\,xo-a\,c\,xo^c\beta\right)\right.$$

$$\left.\left(\sqrt{d}+\sqrt{d+4\mu}\right)\right)^2\left(-1+2^{-\alpha b}\left(\frac{d+2\mu-\sqrt{d^2+4d\mu}}{\mu}\right)^{\alpha b}\right)$$

$$\left(d+\mu\left(1-\frac{d+2\mu-\sqrt{d^2+4d\mu}}{2\mu}\right)^{\alpha}\right)^2\Bigg)\Bigg/\Bigg((-1+c)^2 d^2(d+4\mu)$$

$$\Bigg(xo+\frac{1}{2d\sqrt{d+4\mu}}\left(2\sqrt{d}\,xo\,\mu+\left(d\,xo-a\,c\,xo^c\beta\right)\left(\sqrt{d}+\sqrt{d+4\mu}\right)\right)$$

$$\left(-1+2^{-\alpha b}\left(\frac{d+2\mu-\sqrt{d^2+4d\mu}}{\mu}\right)^{\alpha b}\right)\Bigg)^2\Bigg)-$$

$$\Bigg(2^{1-\alpha b}c\,\alpha\,\alpha b\,\mu\left(\frac{d+2\mu-\sqrt{d^2+4d\mu}}{\mu}\right)^{-1+\alpha b}\left(2\sqrt{d}\,xo\,\mu+\left(d\,xo-a\,c\,xo^c\beta\right)\right.$$

$$\left.\left(\sqrt{d}+\sqrt{d+4\mu}\right)\right)\left(1-\frac{d+2\mu-\sqrt{d^2+4d\mu}}{2\mu}\right)^{-1+\alpha}$$

$$\left(-1+2^{-\alpha b}\left(\frac{d+2\mu-\sqrt{d^2+4d\mu}}{\mu}\right)^{\alpha b}\right)\left(d+\mu\left(1-\frac{d+2\mu-\sqrt{d^2+4d\mu}}{2\mu}\right)^{\alpha}\right)\Bigg)\Bigg/$$

$$\Bigg((-1+c)\,d\sqrt{d+4\mu}\Bigg(xo+\frac{1}{2d\sqrt{d+4\mu}}\left(2\sqrt{d}\,xo\,\mu+\left(d\,xo-a\,c\,xo^c\beta\right)\right.$$

$$\left(\sqrt{d}+\sqrt{d+4\mu}\right)\right)\left(-1+2^{-\alpha b}\left(\frac{d+2\mu-\sqrt{d^{2}+4d\mu}}{\mu}\right)^{\alpha b}\right)\right)\right)\right)-$$

$$\left(2^{2-2\alpha b}\,c\,\alpha b^{2}\left(\frac{d+2\mu-\sqrt{d^{2}+4d\mu}}{\mu}\right)^{-2+2\alpha b}\left(2\sqrt{d}\,xo\,\mu+\left(d\,xo-a\,c\,xo^{c}\,\beta\right)\right.\right.$$

$$\left.\left.\left(\sqrt{d}+\sqrt{d+4\mu}\right)\right)\left(d+\mu\left(1-\frac{d+2\mu-\sqrt{d^{2}+4d\mu}}{2\mu}\right)^{\alpha}\right)^{2}\right)\Big/$$

$$\left((-1+c)\,d\,\sqrt{d+4\mu}\left(xo+\frac{1}{2d\sqrt{d+4\mu}}\left(2\sqrt{d}\,xo\,\mu+\left(d\,xo-a\,c\,xo^{c}\,\beta\right)\right.\right.\right.$$

$$\left(\sqrt{d}+\sqrt{d+4\mu}\right)\right)\left(-1+2^{-\alpha b}\left(\frac{d+2\mu-\sqrt{d^{2}+4d\mu}}{\mu}\right)^{\alpha b}\right)\right)\right)\right)-$$

$$\left(2^{1-\alpha b}\,c\,(-1+\alpha b)\,\alpha b\left(\frac{d+2\mu-\sqrt{d^{2}+4d\mu}}{\mu}\right)^{-2+\alpha b}\left(2\sqrt{d}\,xo\,\mu+\right.\right.$$

$$\left(d\,xo-a\,c\,xo^{c}\,\beta\right)\left(\sqrt{d}+\sqrt{d+4\mu}\right)\right)$$

$$\left(-1+2^{-\alpha b}\left(\frac{d+2\mu-\sqrt{d^{2}+4d\mu}}{\mu}\right)^{\alpha b}\right)\left(d+\mu\left(1-\frac{d+2\mu-\sqrt{d^{2}+4d\mu}}{2\mu}\right)^{\alpha}\right)^{2}\right)\Big/$$

$$\left((-1+c)\,d\,\sqrt{d+4\mu}\left(xo+\frac{1}{2d\sqrt{d+4\mu}}\left(2\sqrt{d}\,xo\,\mu+\left(d\,xo-a\,c\,xo^{c}\,\beta\right)\right.\right.\right.$$

$$\left(\sqrt{d}+\sqrt{d+4\mu}\right)\right)\left(-1+2^{-\alpha b}\left(\frac{d+2\mu-\sqrt{d^{2}+4d\mu}}{\mu}\right)^{\alpha b}\right)\right)\right)\right)-$$

$$\left(2^{-\alpha b}\,c\,xo^{-c}\,\alpha\,\alpha b\,\mu\left(\frac{d+2\mu-\sqrt{d^{2}+4d\mu}}{\mu}\right)^{-1+\alpha b}\left(2\sqrt{d}\,xo\,\mu+\right.\right.$$

$$\left.\left(d\,xo-a\,c\,xo^{c}\,\beta\right)\left(\sqrt{d}+\sqrt{d+4\mu}\right)\right)^{2}\left(1-\frac{d+2\mu-\sqrt{d^{2}+4d\mu}}{2\mu}\right)^{-1+\alpha}$$

$$\left(-1+2^{-\alpha b}\left(\frac{d+2\mu-\sqrt{d^{2}+4d\mu}}{\mu}\right)^{\alpha b}\right)\left(d+\mu\left(1-\frac{d+2\mu-\sqrt{d^{2}+4d\mu}}{2\mu}\right)^{\alpha}\right)^{2}\right)\Big/$$

$$\left(a\,(-1+c)^{2}\,d^{2}\,\beta\,(d+4\mu)\left(xo+\frac{1}{2d\sqrt{d+4\mu}}\left(2\sqrt{d}\,xo\,\mu+\left(d\,xo-a\,c\,xo^{c}\,\beta\right)\right.\right.\right.$$

$$\left(\sqrt{d}+\sqrt{d+4\mu}\right)\right)\left(-1+2^{-\alpha b}\left(\frac{d+2\mu-\sqrt{d^{2}+4d\mu}}{\mu}\right)^{\alpha b}\right)\right)\right)\right)\right)\Big/$$

$$\left(\left(2^{\alpha b}-\left(\frac{d+2\,\mu-\sqrt{d\,(d+4\,\mu)}}{\mu}\right)^{\alpha b}\right)\left(d+\mu\left(1-\frac{d+2\,\mu-\sqrt{d^2+4\,d\,\mu}}{2\,\mu}\right)^{\alpha}\right)^3\right)\Bigg\}\Bigg\}$$

#### 2.2.2. Initialize the Hessian (previously computed)

In[ ]:= HessianEx2FullParamSave = $\Bigg\{\Bigg\{-\Bigg(\Bigg(2^{\alpha b}$

$$e^{-mf-\frac{-2\,d^{3/2}\,xo^{1-c}\,\mu-2\,a\,(-1+c)\,d\,mf\,\beta\,\sqrt{d+4\,\mu}+d\,xo^{-c}\,(-d\,xo+a\,c\,xo^{c}\,\beta)\left(\sqrt{d}+\sqrt{d+4\,\mu}\right)+2^{-\alpha}\,xo^{-c}\,\mu\left(\frac{-d+\sqrt{d\,(d+4\,\mu)}}{\mu}\right)^{\alpha}\left(-2\,\sqrt{d}\,xo\,\mu-(d\,xo-a\,c\,xo^{c}\,\beta)\left(\sqrt{d}+\sqrt{d+4}\right.\right.}{2\,a\,(-1+c)\,d\,\beta\,\sqrt{d+4\,\mu}}}$$

$$\left(d+2^{-\alpha}\,\mu\left(\frac{-d+\sqrt{d\,(d+4\,\mu)}}{\mu}\right)^{\alpha}\right)$$

$$\left(-1+2^{-\alpha b}\left(\frac{d+2\,\mu-\sqrt{d^2+4\,d\,\mu}}{\mu}\right)^{\alpha b}\right)$$

$$\left(\left(xo^{1-c}+\frac{1}{2\,d\,\sqrt{d+4\,\mu}}\,xo^{-c}\left(2\,\sqrt{d}\,xo\,\mu+(d\,xo-a\,c\,xo^{c}\,\beta)\left(\sqrt{d}+\sqrt{d+4\,\mu}\right)\right)\right.\right.$$

$$\left.\left.\left(-1+2^{-\alpha b}\left(\frac{d+2\,\mu-\sqrt{d\,(d+4\,\mu)}}{\mu}\right)^{\alpha b}\right)\right)^{\frac{1}{1-c}}\right)^{-c}$$

$$\left(\left(xo^{1-c}+\frac{1}{2\,d\,\sqrt{d+4\,\mu}}\,xo^{-c}\left(2\,\sqrt{d}\,xo\,\mu+(d\,xo-a\,c\,xo^{c}\,\beta)\left(\sqrt{d}+\sqrt{d+4\,\mu}\right)\right)\right.\right.$$

$$\left.\left.\left(-1+2^{-\alpha b}\left(\frac{d+2\,\mu-\sqrt{d^2+4\,d\,\mu}}{\mu}\right)^{\alpha b}\right)\right)^{\frac{1}{1-c}}\right)^{c}$$

$$\left(d+\mu\left(1-\frac{d+2\,\mu-\sqrt{d^2+4\,d\,\mu}}{2\,\mu}\right)^{\alpha}\right)+\left(a^2\,(-1+c)\,c\,xo^{2\,c}\,\beta^2\left(-1+2^{-\alpha b}\right.\right.$$

$$\left.\left.\left(\frac{d+2\,\mu-\sqrt{d^2+4\,d\,\mu}}{\mu}\right)^{\alpha b}\right)^2\right)\Bigg/\left(\left(d+\mu\left(1-\frac{d+2\,\mu-\sqrt{d^2+4\,d\,\mu}}{2\,\mu}\right)^{\alpha}\right)\right.$$

$$\left(xo+\frac{1}{2\,d\,\sqrt{d+4\,\mu}}\left(2\,\sqrt{d}\,xo\,\mu+(d\,xo-a\,c\,xo^{c}\,\beta)\left(\sqrt{d}+\sqrt{d+4\,\mu}\right)\right)\right.$$

$$\left.\left.\left(-1+2^{-\alpha b}\left(\frac{d+2\,\mu-\sqrt{d^2+4\,d\,\mu}}{\mu}\right)^{\alpha b}\right)\right)^2\right)+$$

$$\left(a^2\, c^2\, xo^{2\,c}\, \beta^2 \left(-1 + 2^{-\alpha b} \left(\frac{d + 2\,\mu - \sqrt{d^2 + 4\,d\,\mu}}{\mu}\right)^{\alpha b}\right)^2\right) \Big/$$

$$\Bigg( \Bigg( d + \mu \left(1 - \frac{d + 2\,\mu - \sqrt{d^2 + 4\,d\,\mu}}{2\,\mu}\right)^{\alpha} \Bigg)$$

$$\Bigg( xo + \frac{1}{2\,d\,\sqrt{d + 4\,\mu}} \left(2\,\sqrt{d}\,\,xo\,\mu + (d\,xo - a\,c\,xo^c\,\beta)\,\left(\sqrt{d} + \sqrt{d + 4\,\mu}\right)\right)$$

$$\left(-1 + 2^{-\alpha b} \left(\frac{d + 2\,\mu - \sqrt{d^2 + 4\,d\,\mu}}{\mu}\right)^{\alpha b}\right)\Bigg)^2 \Bigg) +$$

$$\left(2\,a\,c\,xo^c\,\beta \left(-1 + 2^{-\alpha b} \left(\frac{d + 2\,\mu - \sqrt{d^2 + 4\,d\,\mu}}{\mu}\right)^{\alpha b}\right)\right) \Big/$$

$$\Bigg( xo + \Bigg( \left(2\,\sqrt{d}\,\,xo\,\mu + (d\,xo - a\,c\,xo^c\,\beta)\,\left(\sqrt{d} + \sqrt{d + 4\,\mu}\right)\right)$$

$$\left(-1 + 2^{-\alpha b} \left(\frac{d + 2\,\mu - \sqrt{d^2 + 4\,d\,\mu}}{\mu}\right)^{\alpha b}\right)\Bigg) \Big/ \left(2\,d\,\sqrt{d + 4\,\mu}\right)\Bigg)\Bigg) \Big/$$

$$\left(2^{\alpha b} - \left(\frac{d + 2\,\mu - \sqrt{d\,(d + 4\,\mu)}}{\mu}\right)^{\alpha b}\right)\Bigg),\; - \Bigg(\Bigg(2^{\alpha b}$$

$$e^{-mf - \frac{-2\,d^{3/2}\,xo^{1-c}\,\mu - 2\,a\,(-1+c)\,d\,mf\,\beta\,\sqrt{d+4\,\mu} + d\,xo^{-c}\,(-d\,xo + a\,c\,xo^c\,\beta)\,\left(\sqrt{d} + \sqrt{d+4\,\mu}\right) + 2^{-\alpha}\,xo^{-c}\,\mu\,\left(\frac{-d + \sqrt{d\,(d+4\,\mu)}}{\mu}\right)^{\alpha}\,\left(-2\,\sqrt{d}\,\,xo\,\mu - (d\,xo - a\,c\,xo^c\,\beta)\,\left(\sqrt{d} + \sqrt{d+4}\right.\right.}{2\,a\,(-1+c)\,d\,\beta\,\sqrt{d+4\,\mu}}}$$

$$\left(d + 2^{-\alpha}\,\mu \left(\frac{-d + \sqrt{d\,(d + 4\,\mu)}}{\mu}\right)^{\alpha}\right)$$

$$\Bigg(\Bigg(xo^{1-c} + \frac{1}{2\,d\,\sqrt{d + 4\,\mu}}\,xo^{-c} \left(2\,\sqrt{d}\,\,xo\,\mu + (d\,xo - a\,c\,xo^c\,\beta)\,\left(\sqrt{d} + \sqrt{d + 4\,\mu}\right)\right)$$

$$\left(-1 + 2^{-\alpha b} \left(\frac{d + 2\,\mu - \sqrt{d\,(d + 4\,\mu)}}{\mu}\right)^{\alpha b}\right)\Bigg)^{\frac{1}{1-c}}\Bigg)^{-c}$$

$$\Bigg(\Bigg(xo^{1-c} + \frac{1}{2\,d\,\sqrt{d + 4\,\mu}}\,xo^{-c} \left(2\,\sqrt{d}\,\,xo\,\mu + (d\,xo - a\,c\,xo^c\,\beta)\,\left(\sqrt{d} + \sqrt{d + 4\,\mu}\right)\right)$$

$$\left(-1 + 2^{-\alpha b} \left(\frac{d + 2\,\mu - \sqrt{d^2 + 4\,d\,\mu}}{\mu}\right)^{\alpha b}\right)\Bigg)^{\frac{1}{1-c}}\Bigg)^{c}$$

$$\Bigg(2^{1-\alpha b}\,\alpha b\left(\frac{d+2\,\mu-\sqrt{d^2+4\,d\,\mu}}{\mu}\right)^{-1+\alpha b}\left(d+\mu\left(1-\frac{d+2\,\mu-\sqrt{d^2+4\,d\,\mu}}{2\,\mu}\right)^{\alpha}\right)\Bigg)$$

$$\Bigg(-xo^2\left(d+\mu\left(1-\frac{d+2\,\mu-\sqrt{d^2+4\,d\,\mu}}{2\,\mu}\right)^{\alpha}\right)+$$

$$\Bigg(a\,xo^c\,\beta\left(2\,\sqrt{d}\,xo\,\mu+\left(d\,xo-a\,c\,xo^c\,\beta\right)\left(\sqrt{d}+\sqrt{d+4\,\mu}\right)\right)$$

$$\left(-1+2^{-\alpha b}\left(\frac{d+2\,\mu-\sqrt{d^2+4\,d\,\mu}}{\mu}\right)^{\alpha b}\right)^2\Bigg(c+\Bigg(xo^{-c}\Bigg(2\,\sqrt{d}\,xo\,\mu+$$

$$\left(d\,xo-a\,c\,xo^c\,\beta\right)\left(\sqrt{d}+\sqrt{d+4\,\mu}\right)\Bigg)\Bigg)\Big/\left(2\,a\,\beta\,\sqrt{d+4\,\mu}\right)+$$

$$\frac{1}{2\,a\,d\,\beta\,\sqrt{d+4\,\mu}}\,xo^{-c}\,\mu\,\Bigg(2\,\sqrt{d}\,xo\,\mu+\Bigg(d\,xo-a\,c\,xo^c\,\beta\Bigg)$$

$$\left(\sqrt{d}+\sqrt{d+4\,\mu}\right)\Bigg)\left(1-\frac{d+2\,\mu-\sqrt{d^2+4\,d\,\mu}}{2\,\mu}\right)^{\alpha}\Bigg)\Bigg)\Big/\Bigg(2\,(-1+c)$$

$$d\,\sqrt{d+4\,\mu}\Bigg)-a\,xo^{1+c}\,\beta\left(-1+2^{-\alpha b}\left(\frac{d+2\,\mu-\sqrt{d^2+4\,d\,\mu}}{\mu}\right)^{\alpha b}\right)$$

$$\Bigg(2\,c+\left(\left(-2+c\right)\,xo^{-c}\left(2\,\sqrt{d}\,xo\,\mu+\left(d\,xo-a\,c\,xo^c\,\beta\right)\left(\sqrt{d}+\sqrt{d+4\,\mu}\right)\right)\right)\Big/$$

$$\left(2\,a\,(-1+c)\,\beta\,\sqrt{d+4\,\mu}\right)+$$

$$\Bigg(\left(-2+c\right)\,xo^{-c}\,\mu\left(2\,\sqrt{d}\,xo\,\mu+\left(d\,xo-a\,c\,xo^c\,\beta\right)\left(\sqrt{d}+\sqrt{d+4\,\mu}\right)\right)$$

$$\left(1-\frac{d+2\,\mu-\sqrt{d^2+4\,d\,\mu}}{2\,\mu}\right)^{\alpha}\Bigg)\Big/\left(2\,a\,(-1+c)\,d\,\beta\,\sqrt{d+4\,\mu}\right)\Bigg)\Bigg)-$$

$$\alpha\,\mu\left(1-\frac{d+2\,\mu-\sqrt{d^2+4\,d\,\mu}}{2\,\mu}\right)^{-1+\alpha}\left(-1+2^{-\alpha b}\left(\frac{d+2\,\mu-\sqrt{d^2+4\,d\,\mu}}{\mu}\right)^{\alpha b}\right)$$

$$\Bigg(xo+\frac{1}{2\,d\,\sqrt{d+4\,\mu}}\left(2\,\sqrt{d}\,xo\,\mu+\left(d\,xo-a\,c\,xo^c\,\beta\right)\left(\sqrt{d}+\sqrt{d+4\,\mu}\right)\right)$$

$$\left(-1+2^{-\alpha b}\left(\frac{d+2\,\mu-\sqrt{d^2+4\,d\,\mu}}{\mu}\right)^{\alpha b}\right)\Bigg)\Bigg)$$

$$\Bigg(\left(d\,xo^{1-c}\left(2\,\sqrt{d}\,xo\,\mu+\left(d\,xo-a\,c\,xo^c\,\beta\right)\left(\sqrt{d}+\sqrt{d+4\,\mu}\right)\right)\right)\Big/$$

$$\Big(2\,a\,(-1+c)\,\beta\,\sqrt{d+4\,\mu}\Big)+a\,xo^{c}\,\beta\,\Bigg(-1+2^{-\alpha b}\,\bigg(\frac{d+2\,\mu-\sqrt{d^{2}+4\,d\,\mu}}{\mu}\bigg)^{\alpha b}\Bigg)$$

$$\Big(c+\Big(c\,xo^{-c}\,\Big(2\,\sqrt{d}\,xo\,\mu+\big(d\,xo-a\,c\,xo^{c}\,\beta\big)\,\big(\sqrt{d}+\sqrt{d+4\,\mu}\big)\Big)\Big)\Big/$$

$$\Big(2\,a\,(-1+c)\,\beta\,\sqrt{d+4\,\mu}\Big)+$$

$$\Big(xo^{-2\,c}\,\Big(2\,\sqrt{d}\,xo\,\mu+\big(d\,xo-a\,c\,xo^{c}\,\beta\big)\,\big(\sqrt{d}+\sqrt{d+4\,\mu}\big)\Big)^{2}\Big)\Big/$$

$$\Big(4\,a^{2}\,(-1+c)\,\beta^{2}\,(d+4\,\mu)\Big)\Big)+\Bigg(xo^{-c}\,\mu\,\Big(2\,\sqrt{d}\,xo\,\mu+$$

$$\big(d\,xo-a\,c\,xo^{c}\,\beta\big)\,\big(\sqrt{d}+\sqrt{d+4\,\mu}\big)\Big)\,\Bigg(1-\frac{d+2\,\mu-\sqrt{d^{2}+4\,d\,\mu}}{2\,\mu}\Bigg)^{\alpha}$$

$$\Bigg(2\,d\,xo+a\,xo^{c}\,\beta\,\Bigg(-1+2^{-\alpha b}\,\bigg(\frac{d+2\,\mu-\sqrt{d^{2}+4\,d\,\mu}}{\mu}\bigg)^{\alpha b}\Bigg)$$

$$\Big(c+\Big(xo^{-c}\,\Big(2\,\sqrt{d}\,xo\,\mu+\big(d\,xo-a\,c\,xo^{c}\,\beta\big)\,\big(\sqrt{d}+\sqrt{d+4\,\mu}\big)\Big)\Big)\Big/$$

$$\Big(a\,\beta\,\sqrt{d+4\,\mu}\Big)\Big)+\mu\,\Bigg(1-\frac{d+2\,\mu-\sqrt{d^{2}+4\,d\,\mu}}{2\,\mu}\Bigg)^{\alpha}\,\Bigg(xo+$$

$$\frac{1}{2\,d\,\sqrt{d+4\,\mu}}\,\Big(2\,\sqrt{d}\,xo\,\mu+\big(d\,xo-a\,c\,xo^{c}\,\beta\big)\,\big(\sqrt{d}+\sqrt{d+4\,\mu}\big)\Big)$$

$$\Bigg(-1+2^{-\alpha b}\,\bigg(\frac{d+2\,\mu-\sqrt{d^{2}+4\,d\,\mu}}{\mu}\bigg)^{\alpha b}\Bigg)\Bigg)\Bigg)\Bigg)\Bigg/\Big(2$$

$$a\,(-1+c)\,d\,\beta\,\sqrt{d+4\,\mu}\Big)\Bigg)\Bigg)\Bigg)\Bigg/$$

$$\Bigg(\Bigg(2^{\alpha b}-\bigg(\frac{d+2\,\mu-\sqrt{d\,(d+4\,\mu)}}{\mu}\bigg)^{\alpha b}\Bigg)\,\Bigg(d+\mu\,\Bigg(1-\frac{d+2\,\mu-\sqrt{d^{2}+4\,d\,\mu}}{2\,\mu}\Bigg)^{\alpha}\Bigg)^{2}$$

$$\Bigg(xo+\frac{1}{2\,d\,\sqrt{d+4\,\mu}}\,\Big(2\,\sqrt{d}\,xo\,\mu+\big(d\,xo-a\,c\,xo^{c}\,\beta\big)\,\big(\sqrt{d}+\sqrt{d+4\,\mu}\big)\Big)$$

$$\Bigg(-1+2^{-\alpha b}\,\bigg(\frac{d+2\,\mu-\sqrt{d^{2}+4\,d\,\mu}}{\mu}\bigg)^{\alpha b}\Bigg)\Bigg)^{2}\Bigg)\Bigg\},\ \Bigg\{-\Bigg(\Bigg(2^{\alpha b}$$

$$e^{-mf-\frac{-2\,d^{3/2}\,xo^{1-c}\,\mu-2\,a\,(-1+c)\,d\,mf\,\beta\,\sqrt{d+4\,\mu}+d\,xo^{-c}\,(-d\,xo+a\,c\,xo^{c}\,\beta)\,\left(\sqrt{d}+\sqrt{d+4\,\mu}\right)+2^{-\alpha}\,xo^{-c}\,\mu\,\left(\frac{-d+\sqrt{d\,(d+4\,\mu)}}{\mu}\right)^{\alpha}\,\left(-2\,\sqrt{d}\,xo\,\mu-(d\,xo-a\,c\,xo^{c}\,\beta)\,\left(\sqrt{d}+\sqrt{d+4\text{[illegible]}\right.\right.}{2\,a\,(-1+c)\,d\,\beta\,\sqrt{d+4\,\mu}}}$$

$$\Bigg(d+2^{-\alpha}\,\mu\,\bigg(\frac{-d+\sqrt{d\,(d+4\,\mu)}}{\mu}\bigg)^{\alpha}\Bigg)$$

$$\Bigg(\Bigg(xo^{1-c}+\frac{1}{2\,d\,\sqrt{d+4\,\mu}}\,xo^{-c}\,\Big(2\,\sqrt{d}\,xo\,\mu+\big(d\,xo-a\,c\,xo^{c}\,\beta\big)\,\big(\sqrt{d}+\sqrt{d+4\,\mu}\big)\Big)$$

$$\left(-1+2^{-\alpha b}\left(\frac{d+2\,\mu-\sqrt{d\,(d+4\,\mu)}}{\mu}\right)^{\alpha b}\right)\Bigg)^{\frac{1}{1-c}}\Bigg)^{-c}$$

$$\Bigg(\Bigg(xo^{1-c}+\frac{1}{2\,d\,\sqrt{d+4\,\mu}}\,xo^{-c}\,\Big(2\,\sqrt{d}\,xo\,\mu+\big(d\,xo-a\,c\,xo^{c}\,\beta\big)\,\big(\sqrt{d}+\sqrt{d+4\,\mu}\big)\Big)$$

$$\left(-1+2^{-\alpha b}\left(\frac{d+2\,\mu-\sqrt{d^{2}+4\,d\,\mu}}{\mu}\right)^{\alpha b}\right)\Bigg)^{\frac{1}{1-c}}\Bigg)^{c}$$

$$\left(2^{1-\alpha b}\,\alpha b\left(\frac{d+2\,\mu-\sqrt{d^{2}+4\,d\,\mu}}{\mu}\right)^{-1+\alpha b}\left(d+\mu\left(1-\frac{d+2\,\mu-\sqrt{d^{2}+4\,d\,\mu}}{2\,\mu}\right)^{\alpha}\right)\right.$$

$$\left(-xo^{2}\left(d+\mu\left(1-\frac{d+2\,\mu-\sqrt{d^{2}+4\,d\,\mu}}{2\,\mu}\right)^{\alpha}\right)+\right.$$

$$\Bigg(a\,xo^{c}\,\beta\,\Big(2\,\sqrt{d}\,xo\,\mu+\big(d\,xo-a\,c\,xo^{c}\,\beta\big)\,\big(\sqrt{d}+\sqrt{d+4\,\mu}\big)\Big)$$

$$\left(-1+2^{-\alpha b}\left(\frac{d+2\,\mu-\sqrt{d^{2}+4\,d\,\mu}}{\mu}\right)^{\alpha b}\right)^{2}\Bigg(c+\Big(xo^{-c}\,\Big(2\,\sqrt{d}\,xo\,\mu+$$

$$\big(d\,xo-a\,c\,xo^{c}\,\beta\big)\,\big(\sqrt{d}+\sqrt{d+4\,\mu}\big)\Big)\Big)\Big/\Big(2\,a\,\beta\,\sqrt{d+4\,\mu}\Big)+$$

$$\frac{1}{2\,a\,d\,\beta\,\sqrt{d+4\,\mu}}\,xo^{-c}\,\mu\,\Big(2\,\sqrt{d}\,xo\,\mu+\big(d\,xo-a\,c\,xo^{c}\,\beta\big)$$

$$\big(\sqrt{d}+\sqrt{d+4\,\mu}\big)\Big)\left(1-\frac{d+2\,\mu-\sqrt{d^{2}+4\,d\,\mu}}{2\,\mu}\right)^{\alpha}\Bigg)\Bigg)\Bigg/\Big(2\,(-1+c)$$

$$d\,\sqrt{d+4\,\mu}\Big)-a\,xo^{1+c}\,\beta\left(-1+2^{-\alpha b}\left(\frac{d+2\,\mu-\sqrt{d^{2}+4\,d\,\mu}}{\mu}\right)^{\alpha b}\right)$$

$$\Bigg(2\,c+\Big(\big(-2+c\big)\,xo^{-c}\,\Big(2\,\sqrt{d}\,xo\,\mu+\big(d\,xo-a\,c\,xo^{c}\,\beta\big)\,\big(\sqrt{d}+\sqrt{d+4\,\mu}\big)\Big)\Big)\Big/$$

$$\Big(2\,a\,(-1+c)\,\beta\,\sqrt{d+4\,\mu}\Big)+$$

$$\Big(\big(-2+c\big)\,xo^{-c}\,\mu\,\Big(2\,\sqrt{d}\,xo\,\mu+\big(d\,xo-a\,c\,xo^{c}\,\beta\big)\,\big(\sqrt{d}+\sqrt{d+4\,\mu}\big)\Big)$$

$$\left(1-\frac{d+2\mu-\sqrt{d^2+4d\mu}}{2\mu}\right)^{\alpha}\Bigg)\Bigg/\left(2a(-1+c)d\beta\sqrt{d+4\mu}\right)\Bigg)\Bigg)-$$

$$\alpha\mu\left(1-\frac{d+2\mu-\sqrt{d^2+4d\mu}}{2\mu}\right)^{-1+\alpha}\left(-1+2^{-\alpha b}\left(\frac{d+2\mu-\sqrt{d^2+4d\mu}}{\mu}\right)^{\alpha b}\right)$$

$$\Bigg(xo+\frac{1}{2d\sqrt{d+4\mu}}\left(2\sqrt{d}\,xo\,\mu+\left(d\,xo-a\,c\,xo^c\beta\right)\left(\sqrt{d}+\sqrt{d+4\mu}\right)\right)$$

$$\left(-1+2^{-\alpha b}\left(\frac{d+2\mu-\sqrt{d^2+4d\mu}}{\mu}\right)^{\alpha b}\right)\Bigg)$$

$$\Bigg(\Bigg(d\,xo^{1-c}\left(2\sqrt{d}\,xo\,\mu+\left(d\,xo-a\,c\,xo^c\beta\right)\left(\sqrt{d}+\sqrt{d+4\mu}\right)\right)\Bigg)\Bigg/$$

$$\left(2a(-1+c)\beta\sqrt{d+4\mu}\right)+a\,xo^c\beta\left(-1+2^{-\alpha b}\left(\frac{d+2\mu-\sqrt{d^2+4d\mu}}{\mu}\right)^{\alpha b}\right)$$

$$\Bigg(c+\Bigg(c\,xo^{-c}\left(2\sqrt{d}\,xo\,\mu+\left(d\,xo-a\,c\,xo^c\beta\right)\left(\sqrt{d}+\sqrt{d+4\mu}\right)\right)\Bigg)\Bigg/$$

$$\left(2a(-1+c)\beta\sqrt{d+4\mu}\right)+$$

$$\left(xo^{-2c}\left(2\sqrt{d}\,xo\,\mu+\left(d\,xo-a\,c\,xo^c\beta\right)\left(\sqrt{d}+\sqrt{d+4\mu}\right)\right)^2\right)\Bigg/$$

$$\left(4a^2(-1+c)\beta^2(d+4\mu)\right)\Bigg)+\Bigg(xo^{-c}\mu\Big(2\sqrt{d}\,xo\,\mu+$$

$$\left(d\,xo-a\,c\,xo^c\beta\right)\left(\sqrt{d}+\sqrt{d+4\mu}\right)\Big)\left(1-\frac{d+2\mu-\sqrt{d^2+4d\mu}}{2\mu}\right)^{\alpha}$$

$$\left(2d\,xo+a\,xo^c\beta\left(-1+2^{-\alpha b}\left(\frac{d+2\mu-\sqrt{d^2+4d\mu}}{\mu}\right)^{\alpha b}\right)\right.$$

$$\Bigg(c+\Bigg(xo^{-c}\left(2\sqrt{d}\,xo\,\mu+\left(d\,xo-a\,c\,xo^c\beta\right)\left(\sqrt{d}+\sqrt{d+4\mu}\right)\right)\Bigg)\Bigg/$$

$$\left(a\beta\sqrt{d+4\mu}\right)\Bigg)+\mu\left(1-\frac{d+2\mu-\sqrt{d^2+4d\mu}}{2\mu}\right)^{\alpha}\Bigg(xo+$$

$$\frac{1}{2d\sqrt{d+4\mu}}\left(2\sqrt{d}\,xo\,\mu+\left(d\,xo-a\,c\,xo^c\beta\right)\left(\sqrt{d}+\sqrt{d+4\mu}\right)\right)$$

$$\left(-1+2^{-\alpha b}\left(\frac{d+2\mu-\sqrt{d^2+4d\mu}}{\mu}\right)^{\alpha b}\right)\Bigg)\Bigg)\Bigg)\Bigg)\Bigg/\Big(2$$

$$a(-1+c)d\beta\sqrt{d+4\mu}\Big)\Bigg)\Bigg)\Bigg)\Bigg/$$

$$\Bigg(\Bigg(2^{\alpha b} - \left(\frac{d + 2\,\mu - \sqrt{d\,(d + 4\,\mu)}}{\mu}\right)^{\alpha b}\Bigg)\;\left(d + \mu\left(1 - \frac{d + 2\,\mu - \sqrt{d^2 + 4\,d\,\mu}}{2\,\mu}\right)^{\alpha}\right)^2$$

$$\Bigg(xo + \frac{1}{2\,d\,\sqrt{d + 4\,\mu}}\;\Big(2\,\sqrt{d}\;xo\,\mu + \big(d\,xo - a\,c\,xo^{c}\,\beta\big)\,\big(\sqrt{d} + \sqrt{d + 4\,\mu}\big)\Big)$$

$$\Bigg(-1 + 2^{-\alpha b}\left(\frac{d + 2\,\mu - \sqrt{d^2 + 4\,d\,\mu}}{\mu}\right)^{\alpha b}\Bigg)\Bigg)\Bigg)^{2}\Bigg)\Bigg),\; -\Bigg(\Bigg(\Bigg(2^{\alpha b}$$

$$e^{-mf - \frac{-2\,d^{3/2}\,xo^{1-c}\,\mu - 2\,a\,(-1+c)\,d\,mf\,\beta\,\sqrt{d+4\,\mu} + d\,xo^{-c}\,(-d\,xo + a\,c\,xo^{c}\,\beta)\,\left(\sqrt{d} + \sqrt{d+4\,\mu}\right) + 2^{-\alpha}\,xo^{-c}\,\mu\,\left(\frac{-d + \sqrt{d\,(d+4\,\mu)}}{\mu}\right)^{\alpha}\,\left(-2\,\sqrt{d}\;xo\,\mu - (d\,xo - a\,c\,xo^{c}\,\beta)\,\Big(\sqrt{d} + \sqrt{d+4}\right.}{2\,a\,(-1+c)\,d\,\beta\,\sqrt{d+4\,\mu}}}$$

$$\left(d + 2^{-\alpha}\,\mu\left(\frac{-d + \sqrt{d\,(d + 4\,\mu)}}{\mu}\right)^{\alpha}\right)$$

$$\Bigg(\Bigg(xo^{1-c} + \frac{1}{2\,d\,\sqrt{d + 4\,\mu}}\;xo^{-c}\,\Big(2\,\sqrt{d}\;xo\,\mu + \big(d\,xo - a\,c\,xo^{c}\,\beta\big)\,\big(\sqrt{d} + \sqrt{d + 4\,\mu}\big)\Big)$$

$$\Bigg(-1 + 2^{-\alpha b}\left(\frac{d + 2\,\mu - \sqrt{d\,(d + 4\,\mu)}}{\mu}\right)^{\alpha b}\Bigg)\Bigg)^{\frac{1}{1-c}}\Bigg)^{-c}$$

$$\Bigg(\Bigg(xo^{1-c} + \frac{1}{2\,d\,\sqrt{d + 4\,\mu}}\;xo^{-c}\,\Big(2\,\sqrt{d}\;xo\,\mu + \big(d\,xo - a\,c\,xo^{c}\,\beta\big)\,\big(\sqrt{d} + \sqrt{d + 4\,\mu}\big)\Big)$$

$$\Bigg(-1 + 2^{-\alpha b}\left(\frac{d + 2\,\mu - \sqrt{d^2 + 4\,d\,\mu}}{\mu}\right)^{\alpha b}\Bigg)\Bigg)^{\frac{1}{1-c}}\Bigg)^{c}$$

$$\Bigg(2\,\alpha^2\,\mu^2\left(1 - \frac{d + 2\,\mu - \sqrt{d^2 + 4\,d\,\mu}}{2\,\mu}\right)^{2\,(-1+\alpha)}\left(-1 + 2^{-\alpha b}\left(\frac{d + 2\,\mu - \sqrt{d^2 + 4\,d\,\mu}}{\mu}\right)^{\alpha b}\right) +$$

$$2^{2-\alpha b}\,\alpha\,\alpha b\,\mu\left(\frac{d + 2\,\mu - \sqrt{d^2 + 4\,d\,\mu}}{\mu}\right)^{-1+\alpha b}\left(1 - \frac{d + 2\,\mu - \sqrt{d^2 + 4\,d\,\mu}}{2\,\mu}\right)^{-1+\alpha}$$

$$\left(d + \mu\left(1 - \frac{d + 2\,\mu - \sqrt{d^2 + 4\,d\,\mu}}{2\,\mu}\right)^{\alpha}\right) - (-1 + \alpha)\,\alpha\,\mu\left(1 - \frac{d + 2\,\mu - \sqrt{d^2 + 4\,d\,\mu}}{2\,\mu}\right)^{-2+\alpha}$$

$$\left(-1 + 2^{-\alpha b}\left(\frac{d + 2\,\mu - \sqrt{d^2 + 4\,d\,\mu}}{\mu}\right)^{\alpha b}\right)\left(d + \mu\left(1 - \frac{d + 2\,\mu - \sqrt{d^2 + 4\,d\,\mu}}{2\,\mu}\right)^{\alpha}\right) +$$

$$\Bigg(xo^{-c}\,\alpha^2\,\mu^2\,\Big(2\,\sqrt{d}\;xo\,\mu + \big(d\,xo - a\,c\,xo^{c}\,\beta\big)\,\big(\sqrt{d} + \sqrt{d + 4\,\mu}\big)\Big)$$

$$\Bigg(1-\frac{d+2\mu-\sqrt{d^2+4\,d\,\mu}}{2\,\mu}\Bigg)^{2\,(-1+\alpha)}\Bigg(-1+2^{-\alpha b}\Bigg(\frac{d+2\mu-\sqrt{d^2+4\,d\,\mu}}{\mu}\Bigg)^{\alpha b}\Bigg)$$

$$\Bigg(d+\mu\Bigg(1-\frac{d+2\mu-\sqrt{d^2+4\,d\,\mu}}{2\,\mu}\Bigg)^{\alpha}\Bigg)\Bigg)\Bigg/\Big(a\,(-1+c)\;d\,\beta\,\sqrt{d+4\,\mu}\Big)+2^{2-\alpha b}$$

$$(-1+\alpha b)\;\alpha b\,\Bigg(\frac{d+2\mu-\sqrt{d^2+4\,d\,\mu}}{\mu}\Bigg)^{-2+\alpha b}\Bigg(d+\mu\Bigg(1-\frac{d+2\mu-\sqrt{d^2+4\,d\,\mu}}{2\,\mu}\Bigg)^{\alpha}\Bigg)^{2}+$$

$$\Bigg(2^{1-\alpha b}\,xo^{-c}\,\alpha\,\alpha b\,\mu\,\Bigg(\frac{d+2\mu-\sqrt{d^2+4\,d\,\mu}}{\mu}\Bigg)^{-1+\alpha b}\Big(2\,\sqrt{d}\;xo\,\mu+$$

$$\Big(d\,xo-a\,c\,xo^{c}\,\beta\Big)\Big(\sqrt{d}+\sqrt{d+4\,\mu}\Big)\Big)\Bigg(1-\frac{d+2\mu-\sqrt{d^2+4\,d\,\mu}}{2\,\mu}\Bigg)^{-1+\alpha}$$

$$\Bigg(d+\mu\Bigg(1-\frac{d+2\mu-\sqrt{d^2+4\,d\,\mu}}{2\,\mu}\Bigg)^{\alpha}\Bigg)^{2}\Bigg)\Bigg/\Big(a\,(-1+c)\;d\,\beta\,\sqrt{d+4\,\mu}\Big)-$$

$$\Bigg(xo^{-c}\,(-1+\alpha)\;\alpha\,\mu\,\Big(2\,\sqrt{d}\;xo\,\mu+\Big(d\,xo-a\,c\,xo^{c}\,\beta\Big)\Big(\sqrt{d}+\sqrt{d+4\,\mu}\Big)\Big)$$

$$\Bigg(1-\frac{d+2\mu-\sqrt{d^2+4\,d\,\mu}}{2\,\mu}\Bigg)^{-2+\alpha}\Bigg(-1+2^{-\alpha b}\Bigg(\frac{d+2\mu-\sqrt{d^2+4\,d\,\mu}}{\mu}\Bigg)^{\alpha b}\Bigg)$$

$$\Bigg(d+\mu\Bigg(1-\frac{d+2\mu-\sqrt{d^2+4\,d\,\mu}}{2\,\mu}\Bigg)^{\alpha}\Bigg)^{2}\Bigg)\Bigg/\Big(2\,a\,(-1+c)\;d\,\beta\,\sqrt{d+4\,\mu}\Big)+$$

$$\Bigg(xo^{-2\,c}\,\alpha^2\,\mu^2\,\Big(2\,\sqrt{d}\;xo\,\mu+\Big(d\,xo-a\,c\,xo^{c}\,\beta\Big)\Big(\sqrt{d}+\sqrt{d+4\,\mu}\Big)\Big)^{2}$$

$$\Bigg(1-\frac{d+2\mu-\sqrt{d^2+4\,d\,\mu}}{2\,\mu}\Bigg)^{2\,(-1+\alpha)}\Bigg(-1+2^{-\alpha b}\Bigg(\frac{d+2\mu-\sqrt{d^2+4\,d\,\mu}}{\mu}\Bigg)^{\alpha b}\Bigg)$$

$$\Bigg(d+\mu\Bigg(1-\frac{d+2\mu-\sqrt{d^2+4\,d\,\mu}}{2\,\mu}\Bigg)^{\alpha}\Bigg)^{2}\Bigg)\Bigg/$$

$$\Big(4\,a^2\,(-1+c)^2\,d^2\,\beta^2\,(d+4\,\mu)\Big)+\Bigg(2^{-2\,\alpha b}\,c\,\alpha b^2\,\Bigg(\frac{d+2\mu-\sqrt{d^2+4\,d\,\mu}}{\mu}\Bigg)^{-2+2\,\alpha b}$$

$$\Big(2\,\sqrt{d}\;xo\,\mu+\Big(d\,xo-a\,c\,xo^{c}\,\beta\Big)\Big(\sqrt{d}+\sqrt{d+4\,\mu}\Big)\Big)^{2}$$

$$\Bigg(-1+2^{-\alpha b}\Bigg(\frac{d+2\mu-\sqrt{d^2+4\,d\,\mu}}{\mu}\Bigg)^{\alpha b}\Bigg)\Bigg(d+\mu\Bigg(1-\frac{d+2\mu-\sqrt{d^2+4\,d\,\mu}}{2\,\mu}\Bigg)^{\alpha}\Bigg)^{2}\Bigg)\Bigg/$$

$$\left((-1+c)\ d^2\ (d+4\mu)\left(xo+\frac{1}{2\ d\ \sqrt{d+4\mu}}\left(2\ \sqrt{d}\ xo\ \mu+\left(d\ xo-a\ c\ xo^c\ \beta\right)\right.\right.\right.$$

$$\left.\left.\left.\left(\sqrt{d}+\sqrt{d+4\mu}\right)\right)\left(-1+2^{-\alpha b}\left(\frac{d+2\mu-\sqrt{d^2+4\ d\ \mu}}{\mu}\right)^{\alpha b}\right)\right)^2\right)+$$

$$\left(2^{-2\ \alpha b}\ c^2\ \alpha b^2\left(\frac{d+2\mu-\sqrt{d^2+4\ d\ \mu}}{\mu}\right)^{-2+2\ \alpha b}\left(2\ \sqrt{d}\ xo\ \mu+\left(d\ xo-a\ c\ xo^c\ \beta\right)\right.\right.$$

$$\left.\left(\sqrt{d}+\sqrt{d+4\mu}\right)\right)^2\left(-1+2^{-\alpha b}\left(\frac{d+2\mu-\sqrt{d^2+4\ d\ \mu}}{\mu}\right)^{\alpha b}\right)$$

$$\left.\left(d+\mu\left(1-\frac{d+2\mu-\sqrt{d^2+4\ d\ \mu}}{2\mu}\right)^{\alpha}\right)^2\right)\Big/\left((-1+c)^2\ d^2\ (d+4\mu)\right.$$

$$\left(xo+\frac{1}{2\ d\ \sqrt{d+4\mu}}\left(2\ \sqrt{d}\ xo\ \mu+\left(d\ xo-a\ c\ xo^c\ \beta\right)\left(\sqrt{d}+\sqrt{d+4\mu}\right)\right)\right.$$

$$\left.\left.\left(-1+2^{-\alpha b}\left(\frac{d+2\mu-\sqrt{d^2+4\ d\ \mu}}{\mu}\right)^{\alpha b}\right)\right)^2\right)-$$

$$\left(2^{1-\alpha b}\ c\ \alpha\ \alpha b\ \mu\left(\frac{d+2\mu-\sqrt{d^2+4\ d\ \mu}}{\mu}\right)^{-1+\alpha b}\left(2\ \sqrt{d}\ xo\ \mu+\left(d\ xo-a\ c\ xo^c\ \beta\right)\right.\right.$$

$$\left.\left(\sqrt{d}+\sqrt{d+4\mu}\right)\right)\left(1-\frac{d+2\mu-\sqrt{d^2+4\ d\ \mu}}{2\mu}\right)^{-1+\alpha}$$

$$\left.\left(-1+2^{-\alpha b}\left(\frac{d+2\mu-\sqrt{d^2+4\ d\ \mu}}{\mu}\right)^{\alpha b}\right)\left(d+\mu\left(1-\frac{d+2\mu-\sqrt{d^2+4\ d\ \mu}}{2\mu}\right)^{\alpha}\right)\right)\Big/$$

$$\left((-1+c)\ d\ \sqrt{d+4\mu}\left(xo+\frac{1}{2\ d\ \sqrt{d+4\mu}}\left(2\ \sqrt{d}\ xo\ \mu+\left(d\ xo-a\ c\ xo^c\ \beta\right)\right.\right.\right.$$

$$\left.\left.\left.\left(\sqrt{d}+\sqrt{d+4\mu}\right)\right)\left(-1+2^{-\alpha b}\left(\frac{d+2\mu-\sqrt{d^2+4\ d\ \mu}}{\mu}\right)^{\alpha b}\right)\right)\right)-$$

$$\left(2^{2-2\ \alpha b}\ c\ \alpha b^2\left(\frac{d+2\mu-\sqrt{d^2+4\ d\ \mu}}{\mu}\right)^{-2+2\ \alpha b}\left(2\ \sqrt{d}\ xo\ \mu+\left(d\ xo-a\ c\ xo^c\ \beta\right)\right.\right.$$

$$\left.\left.\left(\sqrt{d}+\sqrt{d+4\mu}\right)\right)\left(d+\mu\left(1-\frac{d+2\mu-\sqrt{d^2+4\ d\ \mu}}{2\mu}\right)^{\alpha}\right)^2\right)\Big/$$

$$\left((-1+c)\, d\, \sqrt{d+4\,\mu}\, \left(xo+\frac{1}{2\, d\, \sqrt{d+4\,\mu}} \left(2\, \sqrt{d}\; xo\, \mu+\left(d\, xo-a\, c\, xo^{c}\, \beta\right)\right.\right.\right.$$

$$\left.\left.\left.\left(\sqrt{d}+\sqrt{d+4\,\mu}\right)\right)\left(-1+2^{-\alpha b}\left(\frac{d+2\,\mu-\sqrt{d^{2}+4\,d\,\mu}}{\mu}\right)^{\alpha b}\right)\right)\right)-$$

$$\left(2^{1-\alpha b}\, c\, (-1+\alpha b)\, \alpha b \left(\frac{d+2\,\mu-\sqrt{d^{2}+4\,d\,\mu}}{\mu}\right)^{-2+\alpha b}\left(2\,\sqrt{d}\; xo\,\mu+\right.\right.$$

$$\left.\left(d\, xo-a\, c\, xo^{c}\,\beta\right)\left(\sqrt{d}+\sqrt{d+4\,\mu}\right)\right)\left(-1+2^{-\alpha b}\left(\frac{d+2\,\mu-\sqrt{d^{2}+4\,d\,\mu}}{\mu}\right)^{\alpha b}\right)$$

$$\left.\left(d+\mu\left(1-\frac{d+2\,\mu-\sqrt{d^{2}+4\,d\,\mu}}{2\,\mu}\right)^{\alpha}\right)^{2}\right)\Big/\left((-1+c)\, d\, \sqrt{d+4\,\mu}\right.$$

$$\left(xo+\frac{1}{2\, d\, \sqrt{d+4\,\mu}}\left(2\,\sqrt{d}\; xo\,\mu+\left(d\, xo-a\, c\, xo^{c}\,\beta\right)\left(\sqrt{d}+\sqrt{d+4\,\mu}\right)\right)\right.$$

$$\left.\left.\left(-1+2^{-\alpha b}\left(\frac{d+2\,\mu-\sqrt{d^{2}+4\,d\,\mu}}{\mu}\right)^{\alpha b}\right)\right)\right)-$$

$$\left(2^{-\alpha b}\, c\, xo^{-c}\,\alpha\,\alpha b\,\mu\left(\frac{d+2\,\mu-\sqrt{d^{2}+4\,d\,\mu}}{\mu}\right)^{-1+\alpha b}\left(2\,\sqrt{d}\; xo\,\mu+\right.\right.$$

$$\left.\left(d\, xo-a\, c\, xo^{c}\,\beta\right)\left(\sqrt{d}+\sqrt{d+4\,\mu}\right)\right)^{2}\left(1-\frac{d+2\,\mu-\sqrt{d^{2}+4\,d\,\mu}}{2\,\mu}\right)^{-1+\alpha}$$

$$\left.\left(-1+2^{-\alpha b}\left(\frac{d+2\,\mu-\sqrt{d^{2}+4\,d\,\mu}}{\mu}\right)^{\alpha b}\right)\left(d+\mu\left(1-\frac{d+2\,\mu-\sqrt{d^{2}+4\,d\,\mu}}{2\,\mu}\right)^{\alpha}\right)^{2}\right)\Big/$$

$$\left(a\,(-1+c)^{2}\, d^{2}\,\beta\,(d+4\,\mu)\left(xo+\frac{1}{2\, d\,\sqrt{d+4\,\mu}}\left(2\,\sqrt{d}\; xo\,\mu+\left(d\, xo-a\, c\, xo^{c}\,\beta\right)\right.\right.\right.$$

$$\left.\left.\left.\left(\sqrt{d}+\sqrt{d+4\,\mu}\right)\right)\left(-1+2^{-\alpha b}\left(\frac{d+2\,\mu-\sqrt{d^{2}+4\,d\,\mu}}{\mu}\right)^{\alpha b}\right)\right)\right)\Big/$$

$$\left(\left(2^{\alpha b}-\left(\frac{d+2\,\mu-\sqrt{d\,(d+4\,\mu)}}{\mu}\right)^{\alpha b}\right)\left(d+\mu\left(1-\frac{d+2\,\mu-\sqrt{d^{2}+4\,d\,\mu}}{2\,\mu}\right)^{\alpha}\right)^{3}\right)\}\};$$

#### 2.2.3. Initialize the Jacobian for checking convergence stability

In[ ]:=
```
mut[ur_] := μb (1 - ur)^α
```

In[•]:=
```
xm[ut_, ur_] := (xo^(1-c) + a (-1 + c) ut (-1 + ur^αb) β)^(1/(1-c));
B[ut_, ur_] := a * (xm[ut, ur])^c
```

In[•]:=
```
Su[vt_, vr_] :=
  ((α mut[vr])/(1 - vr) * (vt + 1/(d + mut[vr])) - αb * vr^(αb-1) (β (B[vt, vr] c vt)/xm[vt, vr] + 1/(1 - vr^αb))) /.
   {αb → 2, α → 2}
```

In[•]:=
```
St[vt_, vr_] := (c * (β * (1 - vr^αb) * B[vt, vr])/xm[vt, vr] - (d + mut[vr])) /. {αb → 2, α → 2}
```

In[•]:=
```
urESS = (d + 2 μb - Sqrt[d^2 + 4 d * μb])/(2 μb);

utESS =
  (xo^-c (2 Sqrt[d] xo μb + (d xo - a c xo^c β) (Sqrt[d] + Sqrt[d + 4 μb]))) / (2 a (-1 + c) d β Sqrt[d + 4 μb]);
```

In[•]:=
```
JacobEx2[μb_] = D[{Su[vt, vr], St[vt, vr]}, {{vr, vt}}] /. {vr → urESS, vt → utESS}
```

Out[•]=

$$\left\{\left\{\frac{4\,\mu b^2\left(1-\frac{d+2\,\mu b-\sqrt{d^2+4\,d\,\mu b}}{2\,\mu b}\right)^2}{\left(d+\mu b\left(1-\frac{d+2\,\mu b-\sqrt{d^2+4\,d\,\mu b}}{2\,\mu b}\right)^2\right)^2}-\right.\right.$$

$$2\,\mu b\left(\frac{xo^{-c}\left(2\,\sqrt{d}\;xo\,\mu b+\left(d\,xo-a\,c\,xo^c\,\beta\right)\left(\sqrt{d}+\sqrt{d+4\,\mu b}\right)\right)}{2\,a\,(-1+c)\,d\,\beta\,\sqrt{d+4\,\mu b}}+\right.$$

$$\left.\frac{1}{d+\mu b\left(1-\frac{d+2\,\mu b-\sqrt{d^2+4\,d\,\mu b}}{2\,\mu b}\right)^2}\right)-\frac{1}{\mu b}$$

$$\left(d+2\,\mu b-\sqrt{d^2+4\,d\,\mu b}\right)\left(\frac{d+2\,\mu b-\sqrt{d^2+4\,d\,\mu b}}{\mu b\left(1-\frac{\left(d+2\,\mu b-\sqrt{d^2+4\,d\,\mu b}\right)^2}{4\,\mu b^2}\right)^2}+\frac{1}{4\,(1-c)\,d^2\,\mu b\,(d+4\,\mu b)}\,c\,xo^{-2\,c}\right.$$

$$\left(d+2\,\mu b-\sqrt{d^2+4\,d\,\mu b}\right)\left(2\,\sqrt{d}\;xo\,\mu b+\left(d\,xo-a\,c\,xo^c\,\beta\right)\left(\sqrt{d}+\sqrt{d+4\,\mu b}\right)\right)^2$$

$$\left(xo^{1-c}+\frac{1}{2\,d\,\sqrt{d+4\,\mu b}}\,xo^{-c}\left(2\,\sqrt{d}\;xo\,\mu b+\left(d\,xo-a\,c\,xo^c\,\beta\right)\left(\sqrt{d}+\sqrt{d+4\,\mu b}\right)\right)\right.$$

$$\left.\left.\left(-1+\frac{\left(d+2\,\mu b-\sqrt{d^2+4\,d\,\mu b}\right)^2}{4\,\mu b^2}\right)\right)^{-1+\frac{1}{1-c}}$$

$$\left(\left(xo^{1-c}+\frac{1}{2\,d\,\sqrt{d+4\,\mu b}}\,xo^{-c}\,\left(2\,\sqrt{d}\;xo\,\mu b+\left(d\,xo-a\,c\,xo^{c}\,\beta\right)\,\left(\sqrt{d}\,+\sqrt{d+4\,\mu b}\right)\right)\right.\right.$$

$$\left.\left.\left.\left(-1+\frac{\left(d+2\,\mu b-\sqrt{d^{2}+4\,d\,\mu b}\right)^{2}}{4\,\mu b^{2}}\right)\right)^{\frac{1}{1-c}}\right)^{-2+c}\right)-$$

$$2\left(\frac{1}{1-\frac{\left(d+2\,\mu b-\sqrt{d^{2}+4\,d\,\mu b}\right)^{2}}{4\,\mu b^{2}}}+\frac{1}{2\,(-1+c)\,d\,\sqrt{d+4\,\mu b}}\,c\,xo^{-c}\,\left(2\,\sqrt{d}\;xo\,\mu b+\right.\right.$$

$$\left.\left.\left(d\,xo-a\,c\,xo^{c}\,\beta\right)\,\left(\sqrt{d}\,+\sqrt{d+4\,\mu b}\right)\right)\right)$$

$$\left(\left(xo^{1-c}+\frac{1}{2\,d\,\sqrt{d+4\,\mu b}}\,xo^{-c}\,\left(2\,\sqrt{d}\;xo\,\mu b+\left(d\,xo-a\,c\,xo^{c}\,\beta\right)\,\left(\sqrt{d}\,+\sqrt{d+4\,\mu b}\right)\right)\right.\right.$$

$$\left.\left.\left.\left(-1+\frac{\left(d+2\,\mu b-\sqrt{d^{2}+4\,d\,\mu b}\right)^{2}}{4\,\mu b^{2}}\right)\right)^{\frac{1}{1-c}}\right)^{-1+c}\right),$$

$$2\,\mu b\,\left(1-\frac{d+2\,\mu b-\sqrt{d^{2}+4\,d\,\mu b}}{2\,\mu b}\right)-\frac{1}{\mu b}\,\left(d+2\,\mu b-\sqrt{d^{2}+4\,d\,\mu b}\right)$$

$$\left(\frac{1}{2\,(1-c)\,d\,\sqrt{d+4\,\mu b}}\right.$$

$$a\,(-1+c)\,c\,xo^{-c}\,\beta\,\left(2\,\sqrt{d}\;xo\,\mu b+\left(d\,xo-a\,c\,xo^{c}\,\beta\right)\,\left(\sqrt{d}\,+\sqrt{d+4\,\mu b}\right)\right)$$

$$\left(-1+\frac{\left(d+2\,\mu b-\sqrt{d^{2}+4\,d\,\mu b}\right)^{2}}{4\,\mu b^{2}}\right)\,\left(xo^{1-c}+\frac{1}{2\,d\,\sqrt{d+4\,\mu b}}\,xo^{-c}\,\left(2\,\sqrt{d}\;xo\,\mu b+\left(d\,xo-\right.\right.\right.$$

$$\left.\left.\left.a\,c\,xo^{c}\,\beta\right)\,\left(\sqrt{d}\,+\sqrt{d+4\,\mu b}\right)\right)\,\left(-1+\frac{\left(d+2\,\mu b-\sqrt{d^{2}+4\,d\,\mu b}\right)^{2}}{4\,\mu b^{2}}\right)\right)^{-1+\frac{1}{1-c}}$$

$$\left(\left(xo^{1-c}+\frac{1}{2\,d\,\sqrt{d+4\,\mu b}}\,xo^{-c}\,\left(2\,\sqrt{d}\;xo\,\mu b+\left(d\,xo-a\,c\,xo^{c}\,\beta\right)\,\left(\sqrt{d}\,+\sqrt{d+4\,\mu b}\right)\right)\right.\right.$$

$$\left.\left.\left(-1+\frac{\left(d+2\,\mu b-\sqrt{d^{2}+4\,d\,\mu b}\right)^{2}}{4\,\mu b^{2}}\right)\right)^{\frac{1}{1-c}}\right)^{-2+c}+$$

$$a\, c\, \beta \left(\left(xo^{1-c} + \frac{1}{2\, d\, \sqrt{d + 4\, \mu b}}\, xo^{-c} \left(2\, \sqrt{d}\, xo\, \mu b + \left(d\, xo - a\, c\, xo^{c}\, \beta\right) \left(\sqrt{d} + \sqrt{d + 4\, \mu b}\right)\right)\right.\right.$$

$$\left.\left.\left(-1 + \frac{\left(d + 2\, \mu b - \sqrt{d^{2} + 4\, d\, \mu b}\right)^{2}}{4\, \mu b^{2}}\right)\right)^{\frac{1}{1-c}}\right)^{-1+c}\Bigg)\Bigg\},$$

$$\Bigg\{2\, \mu b \left(1 - \frac{d + 2\, \mu b - \sqrt{d^{2} + 4\, d\, \mu b}}{2\, \mu b}\right) + \frac{1}{2\, (1 - c)\, d\, \mu b\, \sqrt{d + 4\, \mu b}}$$

$a$

$(-1 + c)$

$c$

$xo^{-c}$

$\beta$

$\left(d + 2\, \mu b - \sqrt{d^{2} + 4\, d\, \mu b}\right)$

$\left(2\, \sqrt{d}\, xo\, \mu b + \left(d\, xo - a\, c\, xo^{c}\, \beta\right) \left(\sqrt{d} + \sqrt{d + 4\, \mu b}\right)\right)$

$$\left(1 - \frac{\left(d + 2\, \mu b - \sqrt{d^{2} + 4\, d\, \mu b}\right)^{2}}{4\, \mu b^{2}}\right)$$

$$\Bigg(xo^{1-c} +$$

$$\left.\frac{xo^{-c} \left(2\, \sqrt{d}\, xo\, \mu b + \left(d\, xo - a\, c\, xo^{c}\, \beta\right) \left(\sqrt{d} + \sqrt{d + 4\, \mu b}\right)\right) \left(-1 + \frac{\left(d + 2\, \mu b - \sqrt{d^{2} + 4\, d\, \mu b}\right)^{2}}{4\, \mu b^{2}}\right)}{2\, d\, \sqrt{d + 4\, \mu b}}\right)^{-1+[illegible]}$$

$$\left(\left(xo^{1-c} + \frac{1}{2\, d\, \sqrt{d + 4\, \mu b}}\, xo^{-c} \left(2\, \sqrt{d}\, xo\, \mu b + \left(d\, xo - a\, c\, xo^{c}\, \beta\right) \left(\sqrt{d} + \sqrt{d + 4\, \mu b}\right)\right)\right.\right.$$

$$\left.\left.\left(-1 + \frac{\left(d + 2\, \mu b - \sqrt{d^{2} + 4\, d\, \mu b}\right)^{2}}{4\, \mu b^{2}}\right)\right)^{\frac{1}{1-c}}\right)^{-2+c} - \frac{1}{\mu b}\, a\, c\, \beta \left(d + 2\, \mu b - \sqrt{d^{2} + 4\, d\, \mu b}\right)$$

$$\left(\left(xo^{1-c} + \frac{1}{2\, d\, \sqrt{d + 4\, \mu b}}\, xo^{-c} \left(2\, \sqrt{d}\, xo\, \mu b + \left(d\, xo - a\, c\, xo^{c}\, \beta\right) \left(\sqrt{d} + \sqrt{d + 4\, \mu b}\right)\right)\right.\right.$$

$$\left(-1+\frac{\left(d+2\,\mu b-\sqrt{d^2+4\,d\,\mu b}\right)^2}{4\,\mu b^2}\right)\Bigg)\Bigg)^{\frac{1}{1-c}}\Bigg)^{-1+c},$$

$$\frac{1}{1-c}\,a^2\,(-1+c)^2\,c\,\beta^2\left(1-\frac{\left(d+2\,\mu b-\sqrt{d^2+4\,d\,\mu b}\right)^2}{4\,\mu b^2}\right)$$

$$\left(-1+\frac{\left(d+2\,\mu b-\sqrt{d^2+4\,d\,\mu b}\right)^2}{4\,\mu b^2}\right)$$

$$\left(xo^{1-c}+\frac{xo^{-c}\left(2\,\sqrt{d}\,xo\,\mu b+\left(d\,xo-a\,c\,xo^{c}\,\beta\right)\left(\sqrt{d}+\sqrt{d+4\,\mu b}\right)\right)\left(-1+\frac{\left(d+2\,\mu b-\sqrt{d^2+4\,d\,\mu b}\right)^2}{4\,\mu b^2}\right)}{2\,d\,\sqrt{d+4\,\mu b}}\right)^{-1+\frac{1}{1-c}}$$

$$\Bigg(\Bigg(xo^{1-c}+\frac{1}{2\,d\,\sqrt{d+4\,\mu b}}\,xo^{-c}\left(2\,\sqrt{d}\,xo\,\mu b+\left(d\,xo-a\,c\,xo^{c}\,\beta\right)\left(\sqrt{d}+\sqrt{d+4\,\mu b}\right)\right)$$

$$\left(-1+\frac{\left(d+2\,\mu b-\sqrt{d^2+4\,d\,\mu b}\right)^2}{4\,\mu b^2}\right)\Bigg)\Bigg)^{\frac{1}{1-c}}\Bigg)^{-2+c}\}\}$$

#### 2.3. Individual-based simulation code

**Comment about the simulation code:**
In section “Population mapping” we specify two function PopIteration[], where all the components of the simulation are captured and PopDynoFullStat[], which allows to take the time average of the results.

In section “Model Specification” we define most of the parameters.

In section “Code for running the Program” we specify where we write the output and the the Do loop runs the program with different sets of parameters.

The simulations then follow a population composed of an endogenously determined number of individuals (approximately from 1800 to 2800 individuals in our simulations, depending on the parameter values) , where each individual is described by a vector of traits consisting of allocation to repair, allocation to growth and number of deleterious mutations the individual has (see function “PopIteration” in section 2.3.1). In the thinning algorithm, during an iteration a maximum of one event can occur (birth, death, or mutation of an individual, see section “Population mapping” below), we considered that a “generation” consist of a number of iterations equal to the carrying capacity predicted by the analytical model (so that on average one event occurs per individual during one generation) and we carried out the simulation for a fixed number Ngenerations of such “generations” (Ngenerations=3000 in the actual simulations so that for a a carrying capacity of 1000, this corresponds to 3000*2000=6 000 000 iterations). We initialize the simulation with a monomorphic population, where individual age is given by a=1/d, where d is the baseline mortality, allocation to germline maintenance and age at maturity are at their equilibrium values predicted from the analysis and the number of deleterious mutations are set to 0.

The function “PopDynoFullStat”: updates the population from one generation to the next by calling the function “PopInitial” and storing relevant statistics in “data” vector. The “data” vector contains the following information: generation time, time average population size, variance of population size, time average “allocation to maintenance”, variance of “allocation to maintenance”, time average “age at maturity”, variance of “age at maturity”, time average “number of novel mutations”, variance of “number of novel mutations”, and “population state” vector.

In section “Code for running the Program”, we use the “Do” loop to runs the simulation program with different values of the baseline mutation rate and we specify the file where we store the output data. The output of simulations is copied into section 2.4. On average, running the “Do” loop once for the chosen parameter values took about 32-62 hours. In order to reproduce the graph we ran the Do loop three times for a different value of the external mortality rate d.

```
In[ ]:= Clear["Global`*"]
```

#### 2.3.1. Population mapping

```
(* Individual={a,ur,t,nu} where a is age of an individual,
ur is investment into repair, t is swithching time,
and nu is the number of deleterious mutations. The
 population state is an array of such individuals and
 PopIteration determines the change in population state *)

PopIteration = Compile[{{PopPrevious, _Real, 2}},
Block[
   {Pop, PopSize, tbound, MaxRate, Time, PopAged, TargetInd, M1, M2, M3, M4, Θ},

   If[Length[PopPrevious] > 0, PopSize = Length[PopPrevious];
    Pop = PopPrevious, PopSize = 1;
```

```
 Pop = {{0, 0, 0, 0}}];

(* Calculates population size and Max
 transition rate from the traits in the population *)

tbound = Max[Table[Pop⟦i, 3⟧, {i, PopSize}]];

(*MaxRate → intensitiy of the point process*)
MaxRate = ( (a (a (1 - c) tbound + wo^(1-c))^(c/(1-c)) (1 - γ PopSize) Exp[-μr]) - sf nubound) +
   (d + sd nubound) + μb;

(* Calculates time of event of homogeneous process
 and transition rates of a target individual for thinning*)

Time = RandomVariate[ExponentialDistribution[1]] / (MaxRate PopSize);
PopAged =
 Table[{Pop⟦i, 1⟧ + Time, Pop⟦i, 2⟧, Pop⟦i, 3⟧, Pop⟦i, 4⟧}, {i, PopSize}];
TargetInd = RandomChoice[PopAged];
M1 = 1/MaxRate (1 - Pmutation)
   f[TargetInd⟦1⟧, TargetInd⟦2⟧, TargetInd⟦3⟧, TargetInd⟦4⟧, PopSize];
M2 =
  1/MaxRate f[TargetInd⟦1⟧, TargetInd⟦2⟧, TargetInd⟦3⟧, TargetInd⟦4⟧, PopSize];
M3 = M2 + dr[TargetInd⟦4⟧]/MaxRate;
M4 = M3 + μ[TargetInd⟦2⟧]/MaxRate;
Θ = RandomVariate[UniformDistribution[{0, 1}]];

(* Iterates time until an event occurs in the true process*)

While[
 M4 ≤ Θ,
 (Time = RandomVariate[ExponentialDistribution[1]] / (MaxRate PopSize);
  PopAged = Table[{PopAged⟦i, 1⟧ + Time,
     PopAged⟦i, 2⟧, PopAged⟦i, 3⟧, PopAged⟦i, 4⟧}, {i, PopSize}];
  TargetInd = RandomChoice[PopAged];
  M1 = 1/MaxRate (1 - Pmutation)
    f[TargetInd⟦1⟧, TargetInd⟦2⟧, TargetInd⟦3⟧, TargetInd⟦4⟧, PopSize];
  M2 = 1/MaxRate
    f[TargetInd⟦1⟧, TargetInd⟦2⟧, TargetInd⟦3⟧, TargetInd⟦4⟧, PopSize];
```

```
   M3 = M2 + dr[TargetInd[[4]]] / MaxRate;
   M4 = M3 + μ[TargetInd[[2]]] / MaxRate;
   Θ = RandomVariate[UniformDistribution[{0, 1}]]) ];

  (* Construct new populations by selecting the appropriate event *)

  If[M1 > Θ, (*new offspring without mutation*)
   Join[PopAged, {{0, TargetInd[[2]], TargetInd[[3]], TargetInd[[4]]}}],
   If[M1 ≤ Θ < M2, (*new offspring with mutation *)Join[PopAged,
     {{0, TargetInd[[2]] + RandomVariate[NormalDistribution[0, mutationStepx]],
       TargetInd[[3]] + RandomVariate[NormalDistribution[0, mutationStepy]],
       TargetInd[[4]]}}],
    If[M2 ≤ Θ < M3, (*death of target individual*)
     DeleteCases[PopAged, TargetInd, 1, 1],
     (* deleterious mutation in target individual*)
     Join[DeleteCases[PopAged, TargetInd, 1, 1],
      {{TargetInd[[1]], TargetInd[[2]], TargetInd[[3]], TargetInd[[4]] + 1.}}]]]] ]]
```

DeleteCases : DeleteCases called with 4 arguments; 2 or 3 arguments are expected.

DeleteCases : DeleteCases called with 4 arguments; 2 or 3 arguments are expected.

Out[•]= CompiledFunction[ Argument count: 1 Argument types: {{_Real, 2 }} ]

In[•]:=

```
PopDynoStat[data_] := Block[{PopNew, Time}, PopNew = PopIteration[data[[6]]];
   Time = data[[1]] + 1.;
   {Time, Length[PopNew] / Time + (Time - 1.) / Time data[[2]],
    Mean[PopNew[[All, 2]]] / Time + (Time - 1.) / Time data[[3]], Mean[PopNew[[All, 3]]] / Time +
     (Time - 1.) / Time data[[4]], Mean[PopNew[[All, 4]]] / Time + (Time - 1.) / Time data[[5]], PopNew}];

(*The "data" vector contains the following information:generation time,
time average population size,variance of population size,
time average "allocation to maintenance",
variance of "allocation to maintenance",time average "age at maturity",
variance of "age at maturity",time average "number of novel mutations",
variance of "number of novel mutations",and "population state" vector*)
```

```
In[ ]:= PopDynoFullStat[data_] :=
  Block[{PopNew, Time, PopSize, TempoMeanPopSize, MeanUr, MeanTs,
    MeanNu, TempoMeanUr, TempoMeanTs, TempoMeanNu},
   PopNew = PopIteration[data[[11]]];
   Time = data[[1]] + 1;
   PopSize = Length[PopNew];
   TempoMeanPopSize = PopSize/Time + (Time - 1)/Time data[[2]];
   MeanUr = Mean[PopNew[[All, 2]]];
   MeanTs = Mean[PopNew[[All, 3]]];
   MeanNu = Mean[PopNew[[All, 4]]];
   TempoMeanUr = MeanUr/Time + (Time - 1)/Time data[[4]];
   TempoMeanTs = MeanTs/Time + (Time - 1)/Time data[[6]];
   TempoMeanNu = MeanNu/Time + (Time - 1)/Time data[[8]];
   {Time, TempoMeanPopSize,
    1/Time (PopSize^2 - 2 PopSize TempoMeanPopSize + TempoMeanPopSize^2) +
     (Time - 1)/Time data[[3]],
    TempoMeanUr,
    1/Time (MeanUr^2 - 2 MeanUr TempoMeanUr + TempoMeanUr^2) + (Time - 1)/Time data[[5]],
    TempoMeanTs,
    1/Time (MeanTs^2 - 2 MeanTs TempoMeanTs + TempoMeanTs^2) + (Time - 1)/Time data[[7]],
    TempoMeanNu, 1/Time (MeanNu^2 - 2 MeanNu TempoMeanNu + TempoMeanNu^2) +
     (Time - 1)/Time data[[9]], Variance[PopNew[[All, 4]]]/Time + (Time - 1)/Time data[[10]], PopNew}]
```

#### 2.3.2. Model specification

Here we specify parameter values and vital rates as functions of traits

ur : investment into repair
ts : switching time (age at maturity)

tbound : bound on switching time
nubound : bound on number of deleterious mutations

In[ ]:=
```
(* Analytical predictions for a uninvdable strategy
 {ur, ts, nu, mutation rate, popSize} from invasion analysis *)
```

In[ ]:=

$$\text{Sing} = \left\{ur,\ ts,\ \frac{\left(d - \sqrt{d}\ \sqrt{d + 4\,\mu b}\right)^2}{4\,\mu b},\right.$$
$$\frac{1}{\gamma}\left(1 + \left(e^{d\,ts + ts\,(1-ur)^{\alpha}\,\mu b + \mu r}\left(a\,(-1+c)\,ts\,(-1+ur^{\nu}) + wo^{1-c}\right)^{\frac{c}{-1+c}}\left(d + (1-ur)^{\alpha}\,\mu b\right)\right) \Big/ \left(a\,(-1+ur^{\nu})\right)\right)\Big\}\ //.\ \left\{ur \to \frac{d + 2\,\mu b - \sqrt{d}\ \sqrt{d+4\,\mu b}}{2\,\mu b},\right.$$
$$ts \to \left(wo^{-c}\left(2\,\sqrt{d}\ wo\,\mu b + \left(d\,wo - a\,c\,wo^{c}\right)\left(\sqrt{d} + \sqrt{d+4\,\mu b}\right)\right)\right) \Big/ \left(2\,a\,(-1+c)\,d\,\sqrt{d+4\,\mu b}\right)\Big\};$$

```
(* vital rates*)

f=Compile[{{age, _Real},{ur, _Real},{ts,_Real},{nu,_Real},{np,_Real}},
If[age≥ts, If[(Exp[-μr]a(1-ur^ν) (a (1-c) ts (1-ur^ν)+wo^(1-c))^(c/(1-c)) (1-γ np))-sf nu>0,(Exp[-μr]a(
dr=Compile[{{nu,_Real}}, d+sd nu];
μ=Compile[{{ur, _Real}}, μb(1-ur)^α];

(* Mutation parameters for evolving phenotype *)
Pmutation = 0.1;      (* Probability of mutation in the life-history locus *)
mutationStepx = 0.07; (* Standard deviation in mutation step, default value for simulat
mutationStepy = 0.07; (* Standard deviation in mutation step, default value for simulat

(*Imortant to set the bounds*)
tbound=4.;   (* bound on the switching time  *)
nubound=2.; (* bound on the number of deleterious mutations  *)

(*Parameter values*)
a=0.9; (*Power law growth mulpticative factor *)
c=0.75; (*Power law growth scaling factor *)
wo=1.; (*Size at birth*)
μr=10^-10; (*mutation rate at giving birth*)
α=2.; (*efficiency of repair*)
ν=2.; (* efficinecy of reproduction/growth*)
d=0.1625; (*mortality rate*)
sf=sd; (*sd, sf reductions in survival and fecundity from carrying an additional  delet
sd=0.2;
γ=0.00035; (*parameter tuning the intensity of density dependence*)
```

In[ ]:=

##### 2.3.3. Code for running the simulations at equilibrium (data for Figures 4 and 5)

In[ ]:=
```
Res = {}; (*Initialize the array where we collect the results*)
```

```
Do[
 (*Initial condition*)
  initialX = Sing[[1]]; (* initial prop. investment into repair *)
  initialY = Sing[[2]];  (* initial switching time  *)
 initialMut = 0.;    (* initial prop. investment into repair *)
 Nind = IntegerPart[Sing[[5]]];
 Nind = 5; (*Here we try for a small number of individuals
  to check if simulation works, comment this line out *)

 PopInitial = Table[{1 / d, initialX, initialY, initialMut}, {j, Nind}];
 (*Define the number of "generations"*)
 Ngenerations = 3000;
 Ngenerations = 2; (*Here we try for a small number of generations
  to check if the simulation works, comment this line out *)

 If[μb == 0.02, startTime = AbsoluteTime[]];
 (*Start counting time*)

 PhenoVecTime =
  NestList[Nest[PopIteration, #, Nind] &, PopInitial, Ngenerations];
 currentTime = AbsoluteTime[] - startTime;

 AppendTo[Res, {μb, Mean[
     Table[Mean[Flatten[PhenoVecTime[[i]][[All, 2]]]], {i, Length[PhenoVecTime]}]],
    Mean[Table[Mean[Flatten[PhenoVecTime[[i]][[All, 3]]]],
      {i, Length[PhenoVecTime]}]], Mean[Table[
       Mean[Flatten[PhenoVecTime[[i]][[All, 4]]]], {i, Length[PhenoVecTime]}]] // N,
    Mean[Table[Length[PhenoVecTime[[i]]], {i, Length[PhenoVecTime]}]] // N,
    currentTime}];
 Print["Current simulation for param:  μb = ", μb, "  Time (s) = ",
  currentTime], {μb, {0.02, 0.06, 0.1, 0.15, 0.2, 0.25, 0.3, 0.35, 0.4, 0.45, 0.5}}]
(*Define values of baseline mortality
 rate for which we want to simulate results*)
```

```
Current simulation for param:  μb = 0.02  Time (s) = 0.050513
Current simulation for param:  μb = 0.06  Time (s) = 0.075240
Current simulation for param:  μb = 0.1  Time (s) = 0.087518
Current simulation for param:  μb = 0.15  Time (s) = 0.096438
Current simulation for param:  μb = 0.2  Time (s) = 0.106286
Current simulation for param:  μb = 0.25  Time (s) = 0.112476
Current simulation for param:  μb = 0.3  Time (s) = 0.118744
Current simulation for param:  μb = 0.35  Time (s) = 0.125727
Current simulation for param:  μb = 0.4  Time (s) = 0.129884
Current simulation for param:  μb = 0.45  Time (s) = 0.137134
Current simulation for param:  μb = 0.5  Time (s) = 0.141716
```

In[ ]:=
```
(*This allows to track which simulation is
 currently computing and how long it took in seconds *)
```

In[ ]:=

In[ ]:=
```
Print[Res]
```

```
{{0.02, 0.0888891, 12.3067, 0., 7.33333, 0.050513},
 {0.06, 0.222946, 10.419, 0., 7.33333, 0.075240},
 {0.1, 0.300827, 9.30549, 0., 5.33333, 0.087518},
 {0.15, 0.368323, 8.35008, 0., 4.66667, 0.096438},
 {0.2, 0.417544, 7.64069, 0., 8., 0.106286},
 {0.25, 0.455733, 7.07246, 0., 9.33333, 0.112476},
 {0.3, 0.478108, 6.58895, 0., 8.66667, 0.118744},
 {0.35, 0.511244, 6.18017, 0.0608466, 9.33333, 0.125727},
 {0.4, 0.543047, 5.81542, 0., 10., 0.129884},
 {0.45, 0.553091, 5.4837, 0.0555556, 9., 0.137134},
 {0.5, 0.569705, 5.18105, 0., 8.66667, 0.141716}}
```

```
(*The Result in the following format: array of vectors of results res(μb)=
 {μb,ur,ts, nu,N,simulTime(sec)},where the first element of the
 results vector indicated the value of the baseline mutation rate μb,
these Results can be copied to section 2.4 "Simulation results" *)
```

#### 2.3.4. Code for running the simulations of equilibrium (data for Figure 6)

```
(* Mutation parameters for evolving phenotype *)
Pmutation = 0.1;  (* Probability of mutation in the life-history locus *)
mutationStepx = 0.07; (* Standard deviation in mutation step,
default value for simulations 0.07 *)
mutationStepy = 0.5; (* Standard deviation in mutation step,
default value for simulations 0.07 *)

(*Imortant to set the bounds*)
tbound = 4.; (* bound on the switching time  *)
nubound = 2.; (* bound on the number of deleterious mutations  *)

(*Parameter values*)
a = 0.9; (*Power law growth mulpticative fector *)
c = 0.75; (*Power law growth scaling factor *)
wo = 1.; (*Size at birth*)
μr = 10^-10; (*mutation rate at giving birth*)
α = 2.; (*efficiency of repair*)
ν = 2.; (* efficinecy of reproduction/growth*)
d = 0.1250; (*mortality rate*)
sf = sd; (*sd, sf reductions in survival and fecundity
 from carrying an additional  deleterious mutation*)
sd = 0.2;
γ = 0.00035; (*parameter tuning the intensity of density dependence*)
```

```
In[ ]:= Nind = 1500; (* Population size *)
Ngenerations = 3500; (* Number of generations *)
initialMut = 0.000001;
(*Initial mean number of deleterious mutations *)
μb = 0.15;

(* Initial condition for the population *)
initialX1 = 0.1; (*Initial allocation to repair*)
initialY1 = 9;  (*Initial switching time*)

initialX2 = 0.1; (*Initial allocation to repair*)
initialY2 = 14;  (*Initial switching time*)

initialX3 = 0.8; (*Initial allocation to repair*)
initialY3 = 9;  (*Initial switching time*)

initialX4 = 0.8; (*Initial allocation to repair*)
initialY4 = 14;  (*Initial switching time*)
```

In[ ]:=

```
In[ ]:= PopInitial1 = Table[{1 / d, initialX1, initialY1, initialMut}, {j, Nind}];
        PopInitial2 = Table[{1 / d, initialX2, initialY2, initialMut}, {j, Nind}];
        PopInitial3 = Table[{1 / d, initialX3, initialY3, initialMut}, {j, Nind}];
        PopInitial4 = Table[{1 / d, initialX4, initialY4, initialMut}, {j, Nind}];

In[ ]:= PhenoVecTime1 =
          NestList[Nest[PopIteration, #, Nind] &, PopInitial1, Ngenerations];

In[ ]:= PhenoVecTime2 =
          NestList[Nest[PopIteration, #, Nind] &, PopInitial2, Ngenerations];

In[ ]:= PhenoVecTime3 =
          NestList[Nest[PopIteration, #, Nind] &, PopInitial3, Ngenerations];

In[ ]:= PhenoVecTime4 =
          NestList[Nest[PopIteration, #, Nind] &, PopInitial4, Ngenerations];

In[ ]:= (*dataFile="/Users/pavila/Dropbox/Apps/convergenceResult.csv";*)

In[ ]:= (*dataStream = OpenWrite[dataFile , PageWidth → Infinity];*)

In[ ]:= urTemp1 =
         Table[Mean[Flatten[PhenoVecTime1⟦i⟧⟦All, 2⟧]], {i, Length[PhenoVecTime1]}]

In[ ]:= utTemp1 =
         Table[Mean[Flatten[PhenoVecTime1⟦i⟧⟦All, 3⟧]], {i, Length[PhenoVecTime1]}]

In[ ]:= urTemp2 =
         Table[Mean[Flatten[PhenoVecTime2⟦i⟧⟦All, 2⟧]], {i, Length[PhenoVecTime2]}]

In[ ]:= utTemp2 =
         Table[Mean[Flatten[PhenoVecTime2⟦i⟧⟦All, 3⟧]], {i, Length[PhenoVecTime2]}]

In[ ]:= urTemp3 =
         Table[Mean[Flatten[PhenoVecTime3⟦i⟧⟦All, 2⟧]], {i, Length[PhenoVecTime3]}]

In[ ]:= utTemp3 =
         Table[Mean[Flatten[PhenoVecTime3⟦i⟧⟦All, 3⟧]], {i, Length[PhenoVecTime3]}]

In[ ]:= urTemp4 =
         Table[Mean[Flatten[PhenoVecTime4⟦i⟧⟦All, 2⟧]], {i, Length[PhenoVecTime4]}]

In[ ]:= utTemp4 =
         Table[Mean[Flatten[PhenoVecTime4⟦i⟧⟦All, 3⟧]], {i, Length[PhenoVecTime4]}]
```

In[ ]:=
```
threadJoin = Quiet[Re@##] /. Re → List &;
(*Defines a useful function for elementwise join of two arrays*)

uTemp1 = threadJoin[urTemp1, utTemp1, Range[Length[urTemp1]]];
uTemp2 = threadJoin[urTemp2, utTemp2, Range[Length[urTemp2]]];
uTemp3 = threadJoin[urTemp3, utTemp3, Range[Length[urTemp3]]];
uTemp4 = threadJoin[urTemp4, utTemp4, Range[Length[urTemp4]]];
```

In[ ]:=
```
(*PutAppend[uTemp1,dataStream];*)
```

In[ ]:=
```
ListPlot[{uTemp1⟦All, {1, 2}⟧,
   uTemp2⟦All, {1, 2}⟧, uTemp3⟦All, {1, 2}⟧, uTemp4⟦All, {1, 2}⟧},
 PlotRange → {{0.05, 0.9}, {6, 17}}, Joined → True, ColorFunction → "Rainbow"]
```

Out[ ]=

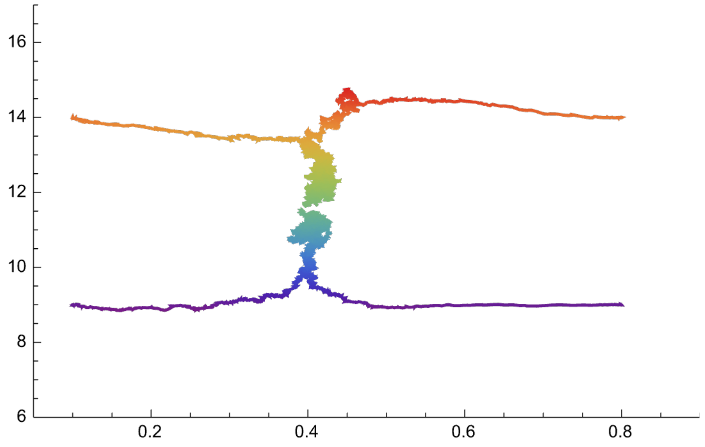


#### 2.4. Simulation results

In[ ]:=
```
(*The Result in the following format: array of vectors of results res(μb)=
 {μb,ur,ts, nu,N,simulTime(sec)} for diffrent
  values of the baseline mutation rate μb,
where the first element of the results vector indicated
 the current value of the baseline mutation rate μb *)
```

#### 2.4.1. Results (d = 0.125)

```
In[]:= DataSimul1 =
  {{0.02`, 0.12080859632233154`, 16.79278864071536`, 0.07339972024754612`,
    2809.418193935355`, 15417.2482775`11.639551859782888},
   {0.06`, 0.2627032565240449`, 14.319338977122218`, 0.15518751334423012`,
    2773.675108297234`, 31485.092955`11.949649973325066},
   {0.1`, 0.346511344443359`, 12.769995934906985`, 0.2023788528407637`,
    2735.2629123625456`, 52513.1805369`12.171813316254454},
   {0.15`, 0.4139652675191035`, 11.612391867685998`, 0.243273330204319`,
    2684.0266577807397`, 78051.2707169`12.343924971501012},
   {0.2`, 0.4632526911221776`, 10.866153622375716`, 0.2711772331531502`,
    2632.8423858713763`, 100636.6238678`12.454301052239268},
   {0.25`, 0.5025584088228097`, 10.084335116436979`, 0.289588408213536`,
    2579.306564478507`, 120533.4795768`12.532652687510156},
   {0.3`, 0.5245337265650566`, 9.435801740316101`, 0.3154726132961478`,
    2524.7154281906032`, 144937.7998656`12.612726658002163},
   {0.35`, 0.5536606090614234`, 9.070435960180943`, 0.32437724912099897`,
    2469.972009330223`, 167822.1278575`12.676394216811218},
   {0.4`, 0.5698643287965464`, 8.431827113118748`, 0.3424995432057324`,
    2413.762412529157`, 186648.9458599`12.722570534993649},
   {0.45`, 0.5917583160974786`, 8.200992291180802`, 0.3453462950441961`,
    2360.2515828057312`, 204766.9022998`12.76280475385549},
   {0.5`, 0.6056073011370291`, 7.999782179416541`, 0.3547699418444243`,
    2310.768077307564`, 223272.2466589`12.800379735949495}};

In[]:=

In[]:= urSimul1 = DataSimul1[[All, {1, 2}]];

In[]:= urSimul1Inter = Interpolation[urSimul1];

In[]:= tsSimul1 = DataSimul1[[All, {1, 3}]];

In[]:= tsSimul1Inter = Interpolation[urSimul1];

In[]:=
  laSimul1 = DataSimul1[[All, {1, 4}]];

In[]:= laSimul1Inter = Interpolation[urSimul1];

In[]:= NSimul1 = DataSimul1[[All, {1, 5}]];

In[]:= NSimul1Inter = Interpolation[urSimul1];

In[]:=
  (*Computes the mutation rate from the data*)

In[]:= muTemp = Flatten[DataSimul1[[All, {1}]] * (1 - DataSimul1[[All, {2}]])^2];
  threadJoin = Quiet[Re@##] /. Re → List &; (*elementwise join*)
  muSimul1 = threadJoin[Flatten[DataSimul1[[All, {1}]]], muTemp];
```

#### 2.4.2. Results (d = 0.1625)

```
In[•]:= DataSimul2 =
  {{0.02`, 0.09920442386280412`, 12.378560333532677`, 0.07682781745337154`,
    2754.676107964012`, 14209.9266723`11.604136830332827},
   {0.06`, 0.22215872528844313`, 10.178753770545576`, 0.16953247303983504`,
    2695.8763745418196`, 28359.1902675`11.904238819887924},
   {0.1`, 0.3002701038615933`, 9.199854507156223`, 0.2298087843274355`,
    2634.5498167277574`, 43572.3694142`12.090756170496983},
   {0.15`, 0.3652445555756124`, 8.255944909277574`, 0.2811082708341462`,
    2557.49116961013`, 59192.0412457`12.223808310431128},
   {0.2`, 0.41579847890619964`, 7.668640626205748`, 0.3162021046192281`,
    2480.7947350883037`, 73861.5474617`12.319963395741844},
   {0.25`, 0.448074412227987`, 7.133280444038305`, 0.3526692015102858`,
    2401.1992669110296`, 87417.5952608`12.393143848963154},
   {0.3`, 0.48438299568604654`, 6.5415903752312445`, 0.3639824739209994`,
    2328.677440853049`, 101313.4700559`12.457212183989931},
   {0.35`, 0.5023036921933003`, 6.203402969049051`, 0.39287345819170033`,
    2254.860046651116`, 115012.8225303`12.512291255099303},
   {0.4`, 0.5324357215830254`, 6.016524564901906`, 0.39723819233643776`,
    2179.068977007664`, 127663.8117258`12.557612800742882},
   {0.45`, 0.5491715732015098`, 5.393139891679337`, 0.40494778938246206`,
    2110.9760079973344`, 139171.529454`12.59509539352869},
   {0.5`, 0.5628892643725261`, 4.987318264298185`, 0.41900530228617694`,
    2039.403532155948`, 149397.827525`12.62588927570919}};

In[•]:=

In[•]:= urSimul2 = DataSimul2⟦All, {1, 2}⟧;

In[•]:= urSimul2Inter = Interpolation[urSimul2];

In[•]:= tsSimul2 = DataSimul2⟦All, {1, 3}⟧;

In[•]:= tsSimul2Inter = Interpolation[urSimul2];

In[•]:=
  laSimul2 = DataSimul2⟦All, {1, 4}⟧;

In[•]:= laSimul2Inter = Interpolation[urSimul2];

In[•]:= NSimul2 = DataSimul2⟦All, {1, 5}⟧;

In[•]:= NSimul2Inter = Interpolation[urSimul2];

In[•]:=
  (*Computes the mutation rate from the data*)

In[•]:= muTemp = Flatten[DataSimul2⟦All, {1}⟧ * (1 - DataSimul2⟦All, {2}⟧)^2];
  threadJoin = Quiet[Re@##] /. Re → List &; (*elementwise join*)
  muSimul2 = threadJoin[Flatten[DataSimul2⟦All, {1}⟧], muTemp];
```

#### 2.4.3. Results (d = 0.2)

```
In[ ]:= DataSimul3 =
  {{0.02`, 0.08629752565650547`, 9.258998631890183`, 0.07824356752730674`,
    2673.492835721426`, 16157.2994437`11.659913767341186},
   {0.06`, 0.19732782568150614`, 8.29083032223525`, 0.18214405372464845`,
    2586.0666444518492`, 31536.2930964`11.950355636756735},
   {0.1`, 0.25995469005381966`, 7.1531006555535805`, 0.25495572100727704`,
    2502.1046317894034`, 43168.329478`12.08671023621006},
   {0.15`, 0.3299886266133877`, 6.166400165246838`, 0.3091743337154376`,
    2399.050983005665`, 54411.8609013`12.187238572652124},
   {0.2`, 0.3743330234957759`, 5.6397044796319875`, 0.35665543193721505`,
    2299.8830389870045`, 65070.8589677`12.264931533170817},
   {0.25`, 0.4186297191310371`, 4.975126671210442`, 0.3764988825540744`,
    2200.4701766077974`, 74559.5224627`12.324048111290416},
   {0.3`, 0.4446919664295791`, 4.571935209363454`, 0.4077912529794138`,
    2108.5201599466845`, 83607.7886216`12.3737917302391},
   {0.35`, 0.4726024580297106`, 4.562918911394726`, 0.43139433270375355`,
    2016.8343885371544`, 91778.8141263`12.414287435373554},
   {0.4`, 0.4976249230948432`, 3.9514428743235466`, 0.4349625499906405`,
    1929.0239920026659`, 99652.898143`12.450034927031327},
   {0.45`, 0.5096749741675464`, 3.764201604413529`, 0.4628479733863716`,
    1843.8890369876708`, 109229.3559612`12.489884366467335},
   {0.5`, 0.5309463347574952`, 3.4197789198285933`, 0.462211286365743`,
    1761.6444518493836`, 117766.4219811`12.52256647387007}};

In[ ]:=

In[ ]:= urSimul3 = DataSimul3[[All, {1, 2}]];

In[ ]:= urSimul3Inter = Interpolation[urSimul3];

In[ ]:= tsSimul3 = DataSimul3[[All, {1, 3}]];

In[ ]:= tsSimul3Inter = Interpolation[urSimul3];

In[ ]:=
  laSimul3 = DataSimul3[[All, {1, 4}]];

In[ ]:= laSimul3Inter = Interpolation[urSimul3];

In[ ]:= NSimul3 = DataSimul3[[All, {1, 5}]];

In[ ]:= NSimul3Inter = Interpolation[urSimul2];

In[ ]:=
  (*Computes the mutation rate from the data*)

In[ ]:= muTemp = Flatten[DataSimul3[[All, {1}]] * (1 - DataSimul3[[All, {2}]])^2];
  threadJoin = Quiet[Re@##] /. Re → List &; (*elementwise join*)
  muSimul3 = threadJoin[Flatten[DataSimul3[[All, {1}]]], muTemp];
```

#### 2.4b. Simulation results (with standard deviations)

```
(*The Result in the following format: array of vectors of results res(μb)=
 {μb,mean(N), std(N), mean(ur), std(ur), mean(ts), std(ts), mean(nu),std(nu),
   simulTime(sec)} for diffrent values of the baseline mutation rate μb,
where the first element of the results vector indicated the
 current value of the baseline mutation rate μb *)
```

##### 2.4.1. Results (d = 0.125)

###### Imported data from csv file

```
In[•]:= DataSimul1 = {{0.02`, 2808.754830113258`, 793.8398802301919`,
     0.134900849875521`, 0.00012541509190665032`, 16.830842502090405`,
     0.002565463618408274`, 0.07155377877111245`, 0.00005872830965438578`,
     0.07181740807800668`, 97887.9782376`12.442274352248463},
    {0.06`, 2774.0066622251834`, 364.2417189332797`, 0.2574236479268348`,
     0.00010388733534314133`, 14.252855605947197`, 0.0007442613665908484`,
     0.15617793340632694`, 0.00016168549796490928`,
     0.15723736116224563`, 180077.6477993`12.707004802707136},
    {0.1`, 2735.4017321785477`, 203.15286693484757`, 0.34819564128582847`,
     0.00006604738047788019`, 12.965247840868896`,
     0.013573658678324932`, 0.2016411709480052`, 0.000216836140652532`,
     0.20366662873472444`, 262640.3942874`12.87090651519987},
    {0.15`, 2685.0299800133243`, 245.4328901808938`, 0.41326419715423723`,
     0.00006545097158577366`, 11.563426940781067`, 0.002153424558965981`,
     0.24284567500260212`, 0.00031070207146567994`,
     0.2462352506449131`, 349362.7656503`12.99482161041564},
    {0.2`, 2631.788807461692`, 389.77446975644614`, 0.4583738314100979`,
     0.00007029223714339156`, 10.708893179251817`,
     0.0011091033820935757`, 0.2765364769773768`, 0.00041436119366715005`,
     0.2816549193793917`, 438151.6102676`13.093169405620259},
    {0.25`, 2579.0093271152564`, 495.83136043930546`, 0.49995987646351403`,
     0.00007860354419738638`, 10.021609002724405`, 0.00201474927708421`,
     0.29203789747653247`, 0.00046255202229046225`,
     0.2972077619638711`, 524438.663625`13.17123969554531},
    {0.3`, 2525.2891405729515`, 638.1143339180934`, 0.5261211510537591`,
     0.0001478158207916008`, 9.528020551377104`,
     0.000621817492800487`, 0.314279162235747`, 0.000665488118609776`,
     0.3214670954908576`, 606976.6216979`13.23471695761386},
    {0.35`, 2471.4750166555627`, 705.8748098671547`, 0.5538367833579493`,
     0.00007385853570277242`, 9.062196434839565`,
     0.001977321219940539`, 0.3244845091838124`, 0.0005788382998155187`,
     0.33223985027209046`, 677562.3073957`13.282494231808341},
    {0.4`, 2417.5649566955362`, 913.1582317152402`, 0.568907189073545`,
     0.000055300820806620056`, 8.501768903874611`,
     0.01163956338714866`, 0.3406476129182799`, 0.0006004006821559078`,
     0.3504292252610172`, 708161.1359287`13.301677082375495},
    {0.45`, 2363.4783477681544`, 976.6776953913908`, 0.5926619690404699`,
     0.00011550048096089256`, 8.044620140824174`,
     0.01769910476168996`, 0.3397009017824008`, 0.0007870449405941597`,
     0.3501349170483164`, 733061.2792736`13.316685273918818},
    {0.5`, 2308.0672884743503`, 1222.7788261376577`, 0.6042607367305617`,
     0.00004812144610104804`, 7.6460232187352695`,
     0.010511263326419776`, 0.35550819682041396`, 0.0006465207752103867`,
     0.3673932530940462`, 786571.121977`13.347282991257144}};
```

#### Save the data

```
In[ ]:=

In[ ]:= NSimul1 = DataSimul1[[All, {1, 2}]];

In[ ]:= NSimul1Inter = Interpolation[NSimul1];

In[ ]:= NWithErr1 =
          Transpose[{DataSimul1[[All, 1]], DataSimul1[[All, 2]], DataSimul1[[All, 3]]}];

In[ ]:= urSimul1 = DataSimul1[[All, {1, 4}]];

In[ ]:= urSimul1Inter = Interpolation[urSimul1];

In[ ]:= urWithErr1 =
          Transpose[{DataSimul1[[All, 1]], DataSimul1[[All, 4]], DataSimul1[[All, 5]]}];

In[ ]:= tsSimul1 = DataSimul1[[All, {1, 6}]];

In[ ]:= tsSimul1Inter = Interpolation[tsSimul1];

In[ ]:= tsWithErr1 =
          Transpose[{DataSimul1[[All, 1]], DataSimul1[[All, 6]], DataSimul1[[All, 7]]}];

In[ ]:=
        laSimul1 = DataSimul1[[All, {1, 8}]];

In[ ]:= laSimul1Inter = Interpolation[laSimul1];

In[ ]:= laWithErr1 =
          Transpose[{DataSimul1[[All, 1]], DataSimul1[[All, 8]], DataSimul1[[All, 9]]}];

In[ ]:=
        (*Computes the mutation rate from the data*)

In[ ]:= muTemp = Flatten[DataSimul1[[All, {1}]] * (1 - DataSimul1[[All, {2}]])^2];
        threadJoin = Quiet[Re@##] /. Re → List &; (*elementwise join*)
        muSimul1 = threadJoin[Flatten[DataSimul1[[All, {1}]]], muTemp];

In[ ]:= urErr1 = ScientificForm[DataSimul1[[All, 5]]]
Out[ ]//ScientificForm=
```

$\{1.25415\times10^{-4}, 1.03887\times10^{-4}, 6.60474\times10^{-5}, 6.5451\times10^{-5}, 7.02922\times10^{-5}, 7.86035\times10^{-5}, 1.47816\times10^{-4}, 7.38585\times10^{-5}, 5.53008\times10^{-5}, 1.155\times10^{-4}, 4.81214\times10^{-5}\}$

```
In[ ]:= tsErr1 = ScientificForm[DataSimul1[[All, 7]]]
Out[ ]//ScientificForm=
```

$\{2.56546\times10^{-3}, 7.44261\times10^{-4}, 1.35737\times10^{-2}, 2.15342\times10^{-3}, 1.1091\times10^{-3}, 2.01475\times10^{-3}, 6.21817\times10^{-4}, 1.97732\times10^{-3}, 1.16396\times10^{-2}, 1.76991\times10^{-2}, 1.05113\times10^{-2}\}$

##### 2.4.2. Results (d = 0.1625)

###### Imported data from csv file

```
In[ ]:= DataSimul2 = {{0.02`, 2755.0433044636907`, 536.5294275622039`,
    0.09903942935561733`, 0.00010331057462534423`, 12.297138227659808`,
    0.0004914115723029283`, 0.0769973953370077`, 0.00007436143406984321`,
    0.07715940294664798`, 87023.5639935`12.391181859065115},
   {0.06`, 2695.502331778814`, 323.32255480880843`, 0.2200711462453549`,
    0.00008320054686602266`, 10.373810235211378`, 0.00502489587619561`,
    0.17239323620192684`, 0.00019058848105624635`,
    0.17336599084035825`, 162509.95748`12.662424970171653},
   {0.1`, 2634.7521652231844`, 340.2727228853022`, 0.2963651969686648`,
    0.0001205965613312057`, 9.388751209257753`, 0.004405569947287283`,
    0.2330702566360764`, 0.0003519245304173914`,
    0.23561468810991942`, 234634.6665308`12.821937171653905},
   {0.15`, 2558.888074616922`, 495.21047729257236`, 0.3654562628552222`,
    0.00009645270160462806`, 8.39900370484576`, 0.005220805099400193`,
    0.28187377344640474`, 0.0004132329843429348`,
    0.2854296026965954`, 295181.8971883`12.921634713204222},
   {0.2`, 2480.8067954696867`, 651.0418914448874`, 0.4123591897592261`,
    0.00010082037798164045`, 7.544514796830182`,
    0.013344856390660544`, 0.3200062548859565`, 0.0005020193367145457`,
    0.3252596858153852`, 358442.5406579`13.005964540458727},
   {0.25`, 2407.228514323784`, 887.1778486431629`, 0.44838395843682777`,
    0.00009954890516295658`, 7.043840765866973`,
    0.0008827515793130204`, 0.3469942156029377`, 0.0005920389515721051`,
    0.3539207117014948`, 432380.693505`13.087411287261434},
   {0.3`, 2328.244503664224`, 1038.7882506699098`, 0.48522222054849173`,
    0.00008509693473545405`, 6.514380111669317`, 0.002314520795875358 7`,
    0.36175480968974677`, 0.0006908317413940796`,
    0.37124674445029243`, 501731.0256975`13.152015950957061},
   {0.35`, 2256.6042638241174`, 1406.8479646910223`, 0.5118285159387631`,
    0.00006945140851971012`, 6.249777835443503`,
    0.00266059191060921 5`, 0.3762422662644141`, 0.000736521072785873`,
    0.38809932762105576`, 554431.2097048`13.195392662828937},
   {0.4`, 2182.008660892738`, 1464.7467295079805`, 0.5315652829902648`,
    0.00005422159808767476 5`, 5.879908944383478`,
    0.00252432026245407 8`, 0.39367867575817045`, 0.0007052909339878725`,
    0.40673864546687344`, 607124.695511`13.234822892167013},
   {0.45`, 2107.7668221185877`, 1687.6669669466319`, 0.5542328194260738`,
    0.00010029948597379469`, 5.506559704216444`, 0.0030313987720422303`,
    0.39909410099005443`, 0.0010137001681308796`,
    0.41333594042647714`, 658097.9561577`13.26983553551795},
   {0.5`, 2038.053964023984`, 1991.0226791809428`, 0.5663426528769431`,
    0.00007343877762362062`, 5.212354656652666`, 0.0019204812304466027`,
    0.41555015117200056`, 0.0009714532329763185`,
    0.4316815056569084`, 680987.5771246`13.284684182889746}};
```

#### Save the data

In[ ]:=

```
In[ ]:= NSimul2 = DataSimul2[[All, {1, 2}]];

In[ ]:= NSimul2Inter = Interpolation[NSimul2];

In[ ]:= NWithErr2 =
          Transpose[{DataSimul2[[All, 1]], DataSimul2[[All, 2]], DataSimul2[[All, 3]]}];

In[ ]:= urSimul2 = DataSimul2[[All, {1, 4}]];

In[ ]:= urSimul2Inter = Interpolation[urSimul2];

In[ ]:= urWithErr2 =
          Transpose[{DataSimul2[[All, 1]], DataSimul2[[All, 4]], DataSimul2[[All, 5]]}];

In[ ]:= tsSimul2 = DataSimul2[[All, {1, 6}]];

In[ ]:= tsSimul2Inter = Interpolation[tsSimul2];

In[ ]:= tsWithErr2 =
          Transpose[{DataSimul2[[All, 1]], DataSimul2[[All, 6]], DataSimul2[[All, 7]]}];

In[ ]:=
        laSimul2 = DataSimul2[[All, {1, 8}]];

In[ ]:= laSimul2Inter = Interpolation[laSimul2];

In[ ]:= laWithErr2 =
          Transpose[{DataSimul2[[All, 1]], DataSimul2[[All, 8]], DataSimul2[[All, 9]]}];

In[ ]:=
        (*Computes the mutation rate from the data*)

In[ ]:= muTemp = Flatten[DataSimul2[[All, {1}]] * (1 - DataSimul2[[All, {2}]])^2];
        threadJoin = Quiet[Re@##] /. Re → List &; (*elementwise join*)
        muSimul2 = threadJoin[Flatten[DataSimul2[[All, {1}]]], muTemp];

In[ ]:= urErr2 = ScientificForm[DataSimul2[[All, 5]]]
```

Out[ ]//ScientificForm=

$\{1.03311\times10^{-4}, 8.32005\times10^{-5}, 1.20597\times10^{-4}, 9.64527\times10^{-5}, 1.0082\times10^{-4}, 9.95489\times10^{-5}, 8.50969\times10^{-5}, 6.94514\times10^{-5}, 5.42216\times10^{-5}, 1.00299\times10^{-4}, 7.34388\times10^{-5}\}$

```
In[ ]:= tsErr2 = ScientificForm[DataSimul2[[All, 7]]]
```

Out[ ]//ScientificForm=

$\{4.91412\times10^{-4}, 5.0249\times10^{-3}, 4.40557\times10^{-3}, 5.22081\times10^{-3}, 1.33449\times10^{-2}, 8.82752\times10^{-4}, 2.31452\times10^{-3}, 2.66059\times10^{-3}, 2.52432\times10^{-3}, 3.0314\times10^{-3}, 1.92048\times10^{-3}\}$

#### 2.4.3. Results (d = 0.2)

##### Imported data from csv file

```
In[ ]:= DataSimul3 = {{0.02`, 2673.2478347768156`, 480.47371888389506`,
      0.08444316281968929`, 0.00015325419939834296`, 9.35495578955712`,
      0.002378812763946473`, 0.07934226649590412`, 0.00008042863577613878`,
      0.07946303746257709`, 78285.8624797`12.345228334079925},
     {0.06`, 2587.6615589606927`, 446.9329896526851`, 0.18657299714445558`,
      0.0001187679025198859`, 7.8865858490276475`, 0.0007488843879577162`,
      0.18566383078563464`, 0.00023507633850044815`,
      0.18690097132628428`, 147865.903351`12.621413034337486},
     {0.1`, 2503.6215856095937`, 618.061503608438`, 0.2700054468449218`,
      0.00008123393969347868`, 7.117643437763498`, 0.0035871495652001224`,
      0.24712799943630748`, 0.00029638997866225246`,
      0.24991289911462383`, 206440.276246`12.76633942504111},
     {0.15`, 2398.8287808127916`, 797.3615524711089`, 0.3291282252204003`,
      0.00022346028715112734`, 6.181428158459358`, 0.0030903404118417656`,
      0.30872246858558305`, 0.0006313043910847502`,
      0.31401577141238196`, 258720.8641797`12.864376446691601},
     {0.2`, 2300.3930712858096`, 1073.8637477844104`, 0.37878852165316296`,
      0.0000685335747874381`, 5.611041553561748`, 0.001497155211118069`,
      0.34999573279790375`, 0.0006304566106273063`,
      0.35592574055704473`, 311389.7923688`12.94484936540642},
     {0.25`, 2201.0612924716856`, 1435.0802546408092`, 0.4141314836677564`,
      0.00018812418144482728`, 5.052717633613999`,
      0.007861091840374531`, 0.38462436970583214`, 0.001066809063789698`,
      0.3946467396966417`, 357614.9040136`13.004960603720479},
     {0.3`, 2107.9187208527646`, 1565.8968534385303`, 0.44774575561336194`,
      0.00008440652213382265`, 4.734198043380007`, 0.0019242890560639027`,
      0.40668850313752636`, 0.0008438172358883622`,
      0.4172680635402238`, 414537.633973`13.069108957778825},
     {0.35`, 2015.8087941372419`, 1711.2011750723543`, 0.46373777098829094`,
      0.00009437176181981719`, 4.298311199518207`,
      0.018729551418060975`, 0.4429431356459253`, 0.0010943713138845449`,
      0.4561894603552144`, 437452.4363544`13.092475833135978},
     {0.4`, 1929.9447035309793`, 2059.0393750130665`, 0.4902247301746488`,
      0.00014596684266175326`, 3.938946437917659`, 0.012645427576222938`,
      0.44645310701841695`, 0.0015042467318397619`,
      0.4596785473537712`, 458387.002783`13.1127772885381},
     {0.45`, 1840.914723517655`, 2331.0011390796735`, 0.511497413918476`,
      0.00013283381439528993`, 3.8950236876740085`,
      0.008195700406704427`, 0.46706048869296896`, 0.0014987813248539764`,
      0.482593273193199`, 481446.1804136`13.134092739046293},
     {0.5`, 1764.338441039307`, 2358.8618063222307`, 0.5247269781273268`,
      0.00008805135185285045`, 3.4525029312098994`,
      0.007391361570435105`, 0.4761027896471976`, 0.0012985395411168337`,
      0.49436715731983866`, 501820.7142695`13.152093577750902}};
```

##### Save the data

```
In[ ]:=

In[ ]:= NSimul3 = DataSimul3[[All, {1, 2}]];

In[ ]:= NSimul3Inter = Interpolation[NSimul3];

In[ ]:= NWithErr3 =
          Transpose[{DataSimul3[[All, 1]], DataSimul3[[All, 2]], DataSimul3[[All, 3]]}];

In[ ]:= urSimul3 = DataSimul3[[All, {1, 4}]];

In[ ]:= urSimul3Inter = Interpolation[urSimul3];

In[ ]:= urWithErr3 =
          Transpose[{DataSimul3[[All, 1]], DataSimul3[[All, 4]], DataSimul3[[All, 5]]}];

In[ ]:= tsSimul3 = DataSimul3[[All, {1, 6}]];

In[ ]:= tsSimul3Inter = Interpolation[tsSimul3];

In[ ]:= tsWithErr3 =
          Transpose[{DataSimul3[[All, 1]], DataSimul3[[All, 6]], DataSimul3[[All, 7]]}];

In[ ]:=
        laSimul3 = DataSimul3[[All, {1, 8}]];

In[ ]:= laSimul3Inter = Interpolation[laSimul3];

In[ ]:= laWithErr3 =
          Transpose[{DataSimul3[[All, 1]], DataSimul3[[All, 8]], DataSimul3[[All, 9]]}];

In[ ]:=
        (*Computes the mutation rate from the data*)

In[ ]:= muTemp = Flatten[DataSimul3[[All, {1}]] * (1 - DataSimul3[[All, {2}]])^2 ];
        threadJoin = Quiet[Re@##] /. Re → List &; (*elementwise join*)
        muSimul3 = threadJoin[Flatten[DataSimul3[[All, {1}]]], muTemp];

In[ ]:= urErr3 = ScientificForm[DataSimul3[[All, 5]]]
Out[ ]//ScientificForm=
```

$\{1.53254\times10^{-4}, 1.18768\times10^{-4}, 8.12339\times10^{-5}, 2.2346\times10^{-4}, 6.85336\times10^{-5}, 1.88124\times10^{-4}, 8.44065\times10^{-5}, 9.43718\times10^{-5}, 1.45967\times10^{-4}, 1.32834\times10^{-4}, 8.80514\times10^{-5}\}$

```
In[ ]:= tsErr3 = ScientificForm[DataSimul3[[All, 7]]]
Out[ ]//ScientificForm=
```

$\{2.37881\times10^{-3}, 7.48884\times10^{-4}, 3.58715\times10^{-3}, 3.09034\times10^{-3}, 1.49716\times10^{-3}, 7.86109\times10^{-3}, 1.92429\times10^{-3}, 1.87296\times10^{-2}, 1.26454\times10^{-2}, 8.1957\times10^{-3}, 7.39136\times10^{-3}\}$

#### 2.5. Graphical illustrations (Fig. 3-6 of the main text)

#### 2.5.1. Initialize parameter data and analytical results for theoretical predictions

In[•]:=

```
Data = {xo → 1.0, a → 0.9, c → 0.75};
DataN = {xo → 1.0, a → 0.9, c → 0.75,
   mr → 0.0, α → 2.0, αb → 2.0, γ → 0.00035, β → 1.0, mf → 0};
```

$$\text{urSin}[d\_, \mu\_] := \frac{d + 2\mu - \sqrt{d}\ \sqrt{d + 4\mu}}{2\mu}$$

$$\text{utSin}[a\_, c\_, d\_, xo\_, \mu\_] := \left(xo^{-c}\left(2\sqrt{d}\ xo\,\mu + \left(d\,xo - a\,c\,xo^{c}\right)\left(\sqrt{d} + \sqrt{d + 4\mu}\right)\right)\right) \Big/ \left(2\,a\,(-1 + c)\,d\sqrt{d + 4\mu}\right)$$

$$\text{NSin}[a\_, c\_, d\_, xo\_, \mu\_, mr\_, \gamma\_, \beta\_] :=$$

$$\frac{1}{\gamma}\Bigg(1 + \Bigg(d\ e^{\frac{xo^{-c}\left(8\sqrt{d}\ xo\,\mu^2 + \left(d\,xo - a\,c\,xo^{c}\,\beta\right)\left(\sqrt{d} + \sqrt{d+4\mu}\right)\left(d - \sqrt{d\,(d+4\mu)}\right) - 2\mu\left(-3\,d^{3/2}\,xo - 2\,d\,xo\sqrt{d+4\mu} - 2\,a\left(-c + (-1+c)\,mf\right)xo^{c}\,\beta\sqrt{d+4\mu} + \sqrt{d}\left(2\,a\,c\,xo^{c}\,\beta + xo\sqrt{d\ [illegible]}\right.\right.\right.}{4\,a\,(-1+c)\,\beta\,\mu\sqrt{d+4\mu}}}$$

$$\mu\left(d + 4\mu - \sqrt{d\,(d + 4\mu)}\right)$$

$$\left(\left(xo^{1-c} + \left(xo^{-c}\left(2\sqrt{d}\ xo\,\mu + \left(d\,xo - a\,c\,xo^{c}\,\beta\right)\left(\sqrt{d} + \sqrt{d + 4\mu}\right)\right)\right.\right.\right.$$

$$\left.\left.\left(-1 + \frac{1}{4\mu^2}\left(d + 2\mu - \sqrt{d\,(d + 4\mu)}\right)^2\right)\right) \Big/ \left(2\,d\sqrt{d + 4\mu}\right)\right)^{\frac{1}{1-c}}\Bigg)^{-c}\Bigg) \Big/$$

$$\left(a\left(d\left(d - \sqrt{d\,(d + 4\mu)}\right) - 2\mu\left(-2\,d + \sqrt{d\,(d + 4\mu)}\right)\right)\right)\Bigg)$$

$$\text{laSin}[d\_, \mu\_, sd\_] := \left(\frac{\left(d - \sqrt{d}\ \sqrt{d + 4\mu}\right)^2}{4\mu} \Big/ sd\right)$$

In[ ]:=

```
(*Uninvadable mutation rate*)
mut[d_, μ_] = μ (1 - urSin[d, μ])^2 ;
(*Uninvdable body size at maturity*)
wm[a_, c_, d_, xo_, μ_] =
  (a (-1 + c) utSin[a, c, d, xo, μ] (-1 + urSin[d, μ]^2) + xo^(1-c))^(1/(1-c)) ;
(*Uninvadable effective fecundity*)
fec[a_, c_, d_, xo_, μ_] = (1 - (urSin[d, μ])^2) * a
   (wm[a, c, d, xo, μ])^c * (1 - γ * NSin[a, c, d, xo, μ, mr, γ, β]) ;

(*Uninvadable fertility*)
fert[a_, c_, d_, xo_, μ_] = (1 - (urSin[d, μ])^2) * a (wm[a, c, d, xo, μ])^c;

(* Classic switching time *)
urSinClassic = 0;
utSinClassic[a_, c_, d_, wo_] := (a c - d wo^(1-c)) / (a d - a c d) ;


NSinClassic[a_, c_, d_, xo_, γ_, β_] =
  (1 - (d E^(-(d xo^(1-c) - a c β)/(a β - a c β)) (((a c β)/d)^(1/(1-c)))^(-c)) / a) / γ ;

(*Classic body size at maturity and fecundity*)
wmSinClassic[a_, c_, d_, xo_] =
  (a (-1 + c) utSinClassic[a, c, d, xo] (-1 + (urSinClassic)^2) + xo^(1-c))^(1/(1-c)) ;
fecSinClassic[a_, c_, d_, xo_, β_] = (1 - (urSinClassic)^2) * a
   (wmSinClassic[a, c, d, xo])^c * (1 - γ * NSinClassic[a, c, d, xo, γ, β]) ;
fertSinClassic[a_, c_, d_, xo_, β_] =
  (1 - (urSinClassic)^2) * a (wmSinClassic[a, c, d, xo])^c ;
```

#### 2.5.2. Graphs for Figure 4 of the main text

In order to produce the graphs sections 2.4.1-2.4.3 and 2.5.1. need to be initialized.

In[ ]:=

```
Show[Plot[{Clip[urSin[0.125, x], {0, 1}],
    Clip[urSin[0.1625, x], {0, 1}], Clip[urSin[0.2, x], {0, 1}]}, {x, 0, 0.515},
  PlotStyle → {{Black, Thickness[0.012]}, {Red, Thickness[0.012]},
    {Orange, Thickness[0.012]}}, PlotRange → {{0, 0.525}, {-0.075, 1}},
  PlotLegends → Placed[{Style["db=0.1250", 16, FontFamily → "Times New Roman"],
     Style["db=0.1625", 16, FontFamily → "Times New Roman"],
     Style["db=0.2000", 16, FontFamily → "Times New Roman"]}, {0.75, 0.25}]],
 ListPlot[urSimul1, PlotMarkers → {Graphics[{Black, Disk[]}], 0.055}],
 ListPlot[urSimul2, PlotMarkers → {Graphics[{Red, Disk[]}], 0.055}],
 ListPlot[urSimul3, PlotMarkers → {Graphics[{Orange, Disk[]}], 0.055}],
 Frame → True,
 FrameLabel → {Style["Baseline mutation rate, μb"],
   Style["Uninvadable investment into repair, u*g"]},
 BaseStyle → {FontFamily → "Times New Roman", 14},
 LabelStyle → {FontFamily → "Times New Roman", 14}, AspectRatio → 0.85]
```

Out[ ]=

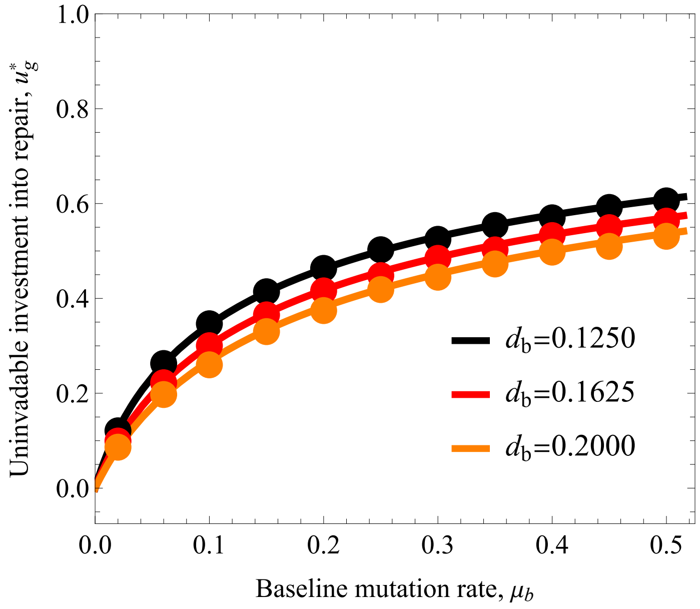

In[ ]:=

```
Show[Plot[{utSin[0.9, 0.75, 0.125, 1, x],
    utSin[0.9, 0.75, 0.1625, 1, x], utSin[0.9, 0.75, 0.2, 1, x]},
   {x, 0, 0.515}, PlotStyle → {{Black, Thickness[0.012]},
     {Red, Thickness[0.012]}, {Orange, Thickness[0.012]}},
   GridLines → {{}, {{utSinClassic[a, c, 0.125, xo] /. DataN, Black},
      {utSinClassic[a, c, 0.1625, xo] /. DataN, Red},
      {utSinClassic[a, c, 0.2, xo] /. DataN, Orange}}},
   GridLinesStyle → Directive[Dashed, Thick],
   PlotRange → {{0, 0.525}, {-0.075, 21}}],
  ListPlot[tsSimul1, PlotMarkers → {Graphics[{Black, Disk[]}], 0.055}],
  ListPlot[tsSimul2, PlotMarkers → {Graphics[{Red, Disk[]}], 0.055}],
  ListPlot[tsSimul3, PlotMarkers → {Graphics[{Orange, Disk[]}], 0.055}],
  Frame → True,
  FrameLabel → {Style["Baseline mutation rate, μb"],
    Style["Uninvadable age at maturity, u*m"]},
  BaseStyle → {FontFamily → "Times New Roman", 14},
  LabelStyle → {FontFamily → "Times New Roman", 14}, AspectRatio → 0.85]
```

Out[ ]=

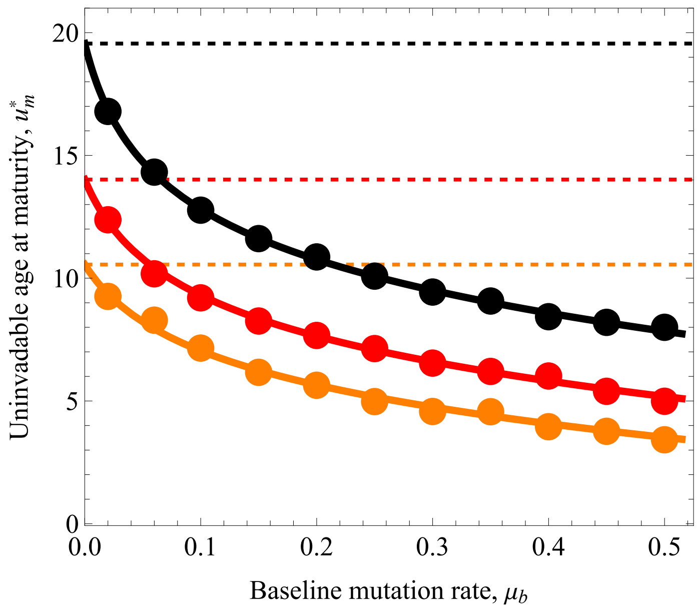

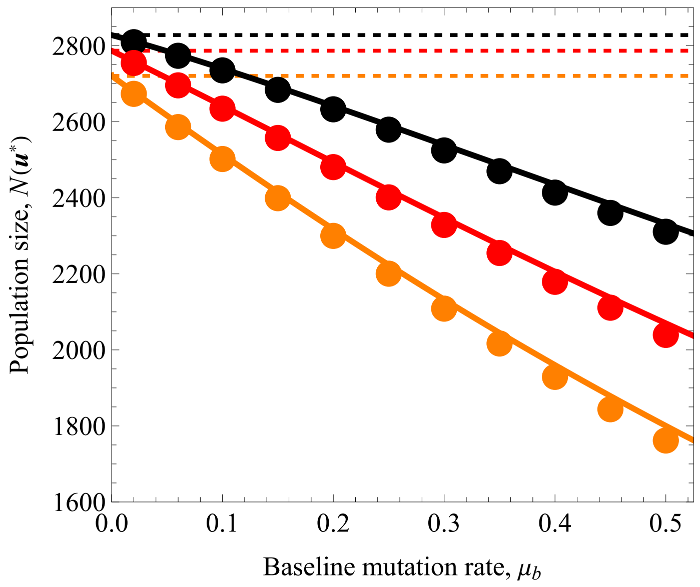


In[ ]:=

```
Show[Plot[{mut[0.125, x], mut[0.1625, x], mut[0.2, x]}, {x, 0, 0.515}, PlotStyle →
    {{Black, Thickness[0.01]}, {Red, Thickness[0.01]}, {Orange, Thickness[0.01]}},
   PlotRange → {{0, 0.525}, {0, 0.15}}],
  ListPlot[muSimul1, PlotMarkers → {Graphics[{Black, Disk[]}], 0.055}],
  ListPlot[muSimul2, PlotMarkers → {Graphics[{Red, Disk[]}], 0.055}],
  ListPlot[muSimul3, PlotMarkers → {Graphics[{Orange, Disk[]}], 0.055}],
  Frame → True,
  FrameLabel → {Style["Baseline mutation rate, μb"],
    Style["Uninvadable mutation rate, μs(ur*)"]},
  BaseStyle → {FontFamily → "Times New Roman", 14},
  LabelStyle → {FontFamily → "Times New Roman", 14}, AspectRatio → 0.85]
```

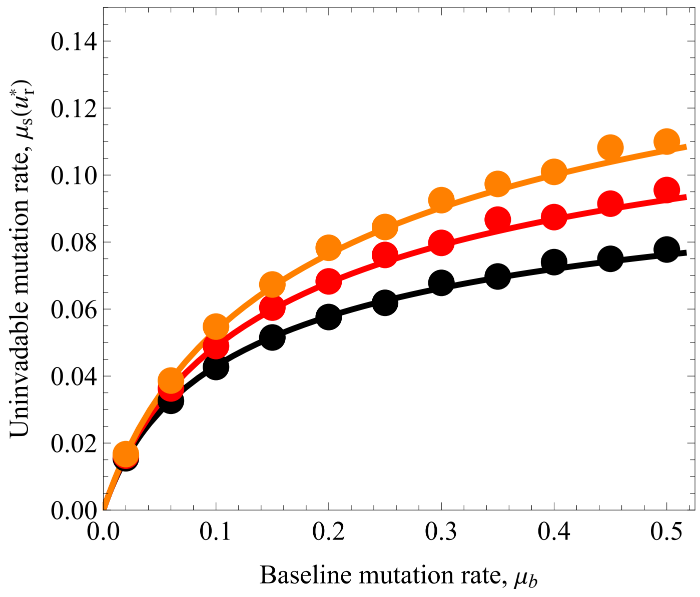


#### 2.5.2b Graphs for mean traits with standard deviations

```
(* Simulations with standard deviations as error bars*)
```

In[ ]:=
```
Needs["ErrorBarPlots`"]
Show[Plot[{Clip[urSin[0.125, x], {0, 1}],
    Clip[urSin[0.1625, x], {0, 1}], Clip[urSin[0.2, x], {0, 1}]}, {x, 0, 0.515},
   PlotStyle → {{Black, Thickness[0.003]}, {Red, Thickness[0.003]},
     {Orange, Thickness[0.003]}}, PlotRange → {{0, 0.525}, {-0.075, 1}},
   PlotLegends → Placed[{Style["db=0.1250", 16, FontFamily → "Times New Roman"],
      Style["db=0.1625", 16, FontFamily → "Times New Roman"],
      Style["db=0.2000", 16, FontFamily → "Times New Roman"]}, {0.75, 0.25}]],
  (*ListPlot[urSimul1, PlotMarkers →{Graphics[{Black,Disk[]}],0.055}],
  ListPlot[urSimul2, PlotMarkers →{Graphics[{Red,Disk[]}],0.055}],
  ListPlot[urSimul3, PlotMarkers →{Graphics[{Orange,Disk[]}],0.055}],*)
  ErrorListPlot[urWithErr1, PlotStyle → Black,
   PlotMarkers → {Graphics[{Black, Disk[]}], 0.001}],
  ErrorListPlot[urWithErr2, PlotStyle → Red,
   PlotMarkers → {Graphics[{Black, Disk[]}], 0.001}],
  ErrorListPlot[urWithErr3, PlotStyle → Orange,
   PlotMarkers → {Graphics[{Black, Disk[]}], 0.001}],
  Frame → True,
  FrameLabel → {Style["Baseline mutation rate, μb"],
    Style["Uninvadable investment into repair, u*g"]},
  BaseStyle → {FontFamily → "Times New Roman", 14},
  LabelStyle → {FontFamily → "Times New Roman", 14}, AspectRatio → 0.85]
```

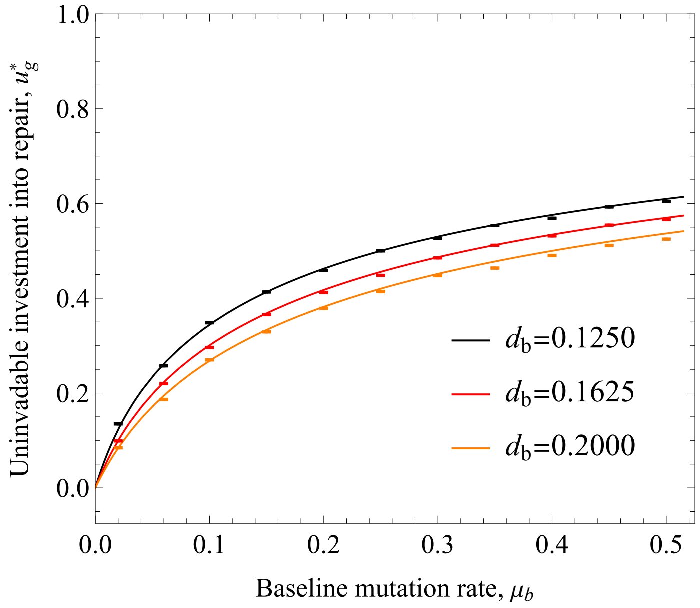

In[ ]:=
```
Needs["ErrorBarPlots`"]
Show[Plot[{utSin[0.9, 0.75, 0.125, 1, x],
    utSin[0.9, 0.75, 0.1625, 1, x], utSin[0.9, 0.75, 0.2, 1, x]},
   {x, 0, 0.515}, PlotStyle → {{Black, Thickness[0.003]},
     {Red, Thickness[0.003]}, {Orange, Thickness[0.003]}},
   GridLines → {{}, {{utSinClassic[a, c, 0.125, xo] /. DataN, Black},
      {utSinClassic[a, c, 0.1625, xo] /. DataN, Red},
      {utSinClassic[a, c, 0.2, xo] /. DataN, Orange}}},
   GridLinesStyle → Directive[Dashed, Thick],
   PlotRange → {{0, 0.525}, {-0.075, 21}}],
  (*ListPlot[tsSimul1, PlotMarkers →{Graphics[{Black,Disk[]}],0.055}],
  ListPlot[tsSimul2, PlotMarkers →{Graphics[{Red,Disk[]}],0.055}],
  ListPlot[tsSimul3, PlotMarkers →{Graphics[{Orange,Disk[]}],0.055}], *)
  ErrorListPlot[tsWithErr1, PlotStyle → Black,
   PlotMarkers → {Graphics[{Black, Disk[]}], 0.001}],
  ErrorListPlot[tsWithErr2, PlotStyle → Red,
   PlotMarkers → {Graphics[{Black, Disk[]}], 0.001}],
  ErrorListPlot[tsWithErr3, PlotStyle → Orange,
   PlotMarkers → {Graphics[{Black, Disk[]}], 0.001}],
  Frame → True,
  FrameLabel → {Style["Baseline mutation rate, μb"],
    Style["Uninvadable age at maturity, u*m"]},
  BaseStyle → {FontFamily → "Times New Roman", 14},
  LabelStyle → {FontFamily → "Times New Roman", 14}, AspectRatio → 0.85]
```

Out[ ]=

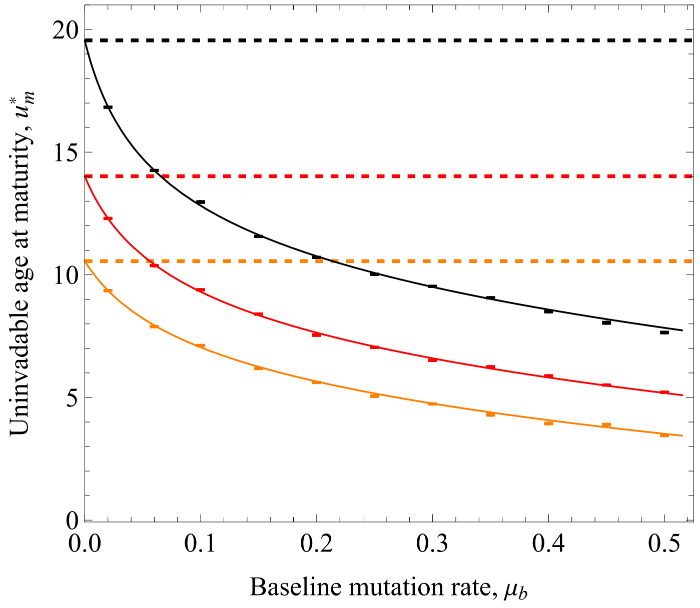

#### 2.5.3. Graphs for Figure 6 ($x_m(\boldsymbol{u}^*)$ body size of maturity and $b_0(\boldsymbol{u}^*, \boldsymbol{u}^*)$ birth rate as a function of $\mu_b$)

In[∘]:=
```
LogPlot[{Evaluate@Table[wm[a, c, d, xo, μ] /. DataN, {d, {0.125, 0.1625, 0.2}}] },
  {μ, 0, 0.515},
  GridLines → {{}, {{wmSinClassic[a, c, 0.125, xo] /. DataN, Black},
     {wmSinClassic[a, c, 0.1625, xo] /. DataN, Red},
     {wmSinClassic[a, c, 0.2, xo] /. DataN, Orange}}},
  PlotStyle → {{Black, Thickness[0.012]},
    {Red, Thickness[0.012]}, {Orange, Thickness[0.012]}},
  PlotRange → {{Automatic, Automatic}, {1, Automatic}},
  GridLinesStyle → Directive[Dashed, Thick],
  Frame → True,
  FrameLabel →
   {Style["Baseline mutation rate, μb"], Style["Body size at maturity, xm(u*)"]},
  BaseStyle → {FontFamily → "Times New Roman", 14},
  LabelStyle → {FontFamily → "Times New Roman", 14}, AspectRatio → 0.8]
```

Out[∘]=

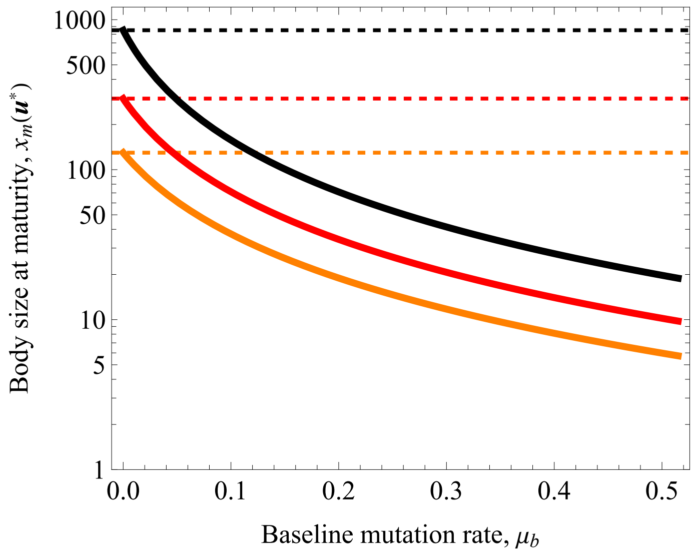

In[ ]:=
```
Plot[{Evaluate@Table[fec[a, c, d, xo, μ] /. DataN, {d, {0.125, 0.1625, 0.20}}] },
  {μ, 0, 0.515},
  PlotStyle → {{Black, Thickness[0.012]},
    {Red, Thickness[0.012]}, {Orange, Thickness[0.012]}},
  PlotRange → {{Automatic, Automatic}, {0.75, Automatic}},
  GridLines → {{}, {{fecSinClassic[a, c, 0.125, xo, β] /. DataN, Black},
     {fecSinClassic[a, c, 0.1625, xo, β] /. DataN, Red},
     {fecSinClassic[a, c, 0.2, xo, β] /. DataN, Orange}}},
  GridLinesStyle → Directive[Dashed, Thick],
  Frame → True,
  FrameLabel → {Style["Baseline mutation rate, μb"],
    Style["Effective birth rate, b0(u*,u*)"]},
  BaseStyle → {FontFamily → "Times New Roman", 14},
  LabelStyle → {FontFamily → "Times New Roman", 14}, AspectRatio → 0.8]
```

Out[ ]=

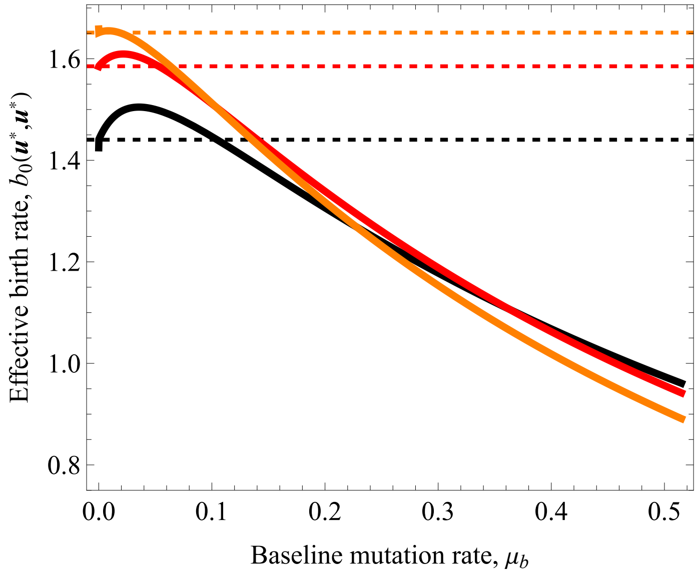


#### 2.5.4. Evaluate uninvadability at the parameter values plotted

In[ ]:=
```
DataN = {xo → 1.0, a → 0.9, c → 0.75,
   mr → 0.0, α → 2.0, αb → 2.0, γ → 0.00035, β → 1.0, mf → 0};
```

In[ ]:=
```
(*Note that the Hessian from section 2.2.2. should be initialized here *)
```

In[ ]:=
```
HuEx2Data = HessianEx2FullParamSave /. DataN;
```

In[ ]:=
```
(*d→0.125, d→0.1625 ,d→0.2*)
```

In[ ]:=
```
Do[Print[NegativeDefiniteMatrixQ[HuEx2Data /. {d → 0.125}]],
 {μ, 0.05, 0.5, 0.05}]
```

```
True
True
True
True
True
True
True
True
True
True
```

In[ ]:=

```
Do[Print[NegativeDefiniteMatrixQ[HuEx2Data /. {d → 0.1625}]],
  {μ, 0.05, 0.5, 0.05}]
```

```
True
True
True
True
True
True
True
True
True
True
```

In[ ]:=

```
Do[Print[NegativeDefiniteMatrixQ[HuEx2Data /. {d → 0.2}]], {μ, 0.05, 0.5, 0.05}]
```

```
True
True
True
True
True
True
True
True
True
True
```

In[ ]:=

```
(* If TRUE→
 the Hessian of Ro is negative definite → The startegies are uninvadable *)
```

#### 2.5.5. Evaluate convergence stability at the parameter values plotted

In[ ]:=
```
(*Check convergence stability,
note that the Jacobian should be initialized here from section 1.2.2. *)
```

In[ ]:=
```
Do[Print[NegativeDefiniteMatrixQ[JacobEx2[μb] /. DataN /. d → 0.1250]],
  {μb, 0.1, 2, 0.1}]
```

```
True
True
True
True
True
True
True
True
True
True
True
True
True
True
True
True
True
True
True
True
```

In[ ]:=
```
Do[Print[NegativeDefiniteMatrixQ[JacobEx2[μb] /. DataN /. d → 0.1625]],
  {μb, 0.1, 2, 0.1}]
```

```
True
True
True
True
True
True
True
True
True
True
True
True
True
True
True
True
True
True
True
True
```

In[ ]:=
```
Do[Print[NegativeDefiniteMatrixQ[JacobEx2[μb] /. DataN /. d → 0.200]],
  {μb, 0.1, 2, 0.1}]
```

```
True
True
True
True
True
True
True
True
True
True
True
True
True
True
True
True
True
True
True
True
```

In[ ]:=
```
(*If TRUE→ Jacobian of selection gradients is negative definite for
   these parameter values → Singular strategy is converegnce stable*)
```

##### 2.5.6. Comment about mean number of mutations

In[ ]:=
```
(*Note that the mean number of mutations in this
 model is not expected to follow Poisson distribution,
because the vital rates (birth rate - death) over indidivuals life
 are not constant. This is because  we have the growth phase. However,
it is interesting to note that it seems that the simulations
 imply that the number of deleterious mutations are smaller
 than the prediction with constant vital rates would imply.   *)
```

In[ ]:=
```
Show[Plot[{laSin[0.125, x, 0.2], laSin[0.1625, x, 0.2], laSin[0.2, x, 0.2]},
  {x, 0, 0.515}, PlotStyle → {{Black, Thickness[0.012]},
    {Red, Thickness[0.012]}, {Orange, Thickness[0.012]}},
  PlotRange → {{0, 0.525}, {-0.075, 1}}],
 ListPlot[laSimul1, PlotMarkers → {Graphics[{Black, Disk[]}], 0.055}],
 ListPlot[laSimul2, PlotMarkers → {Graphics[{Red, Disk[]}], 0.055}],
 ListPlot[laSimul3, PlotMarkers → {Graphics[{Orange, Disk[]}], 0.055}],
 Frame → True,
 FrameLabel → {Style["Baseline mutation rate, μb"],
   Style["The mean number of deleterious mutations, λ(u*)"]},
 BaseStyle → {FontFamily → "Times New Roman", 14},
 LabelStyle → {FontFamily → "Times New Roman", 14}, AspectRatio → 0.85]
```

Out[ ]=

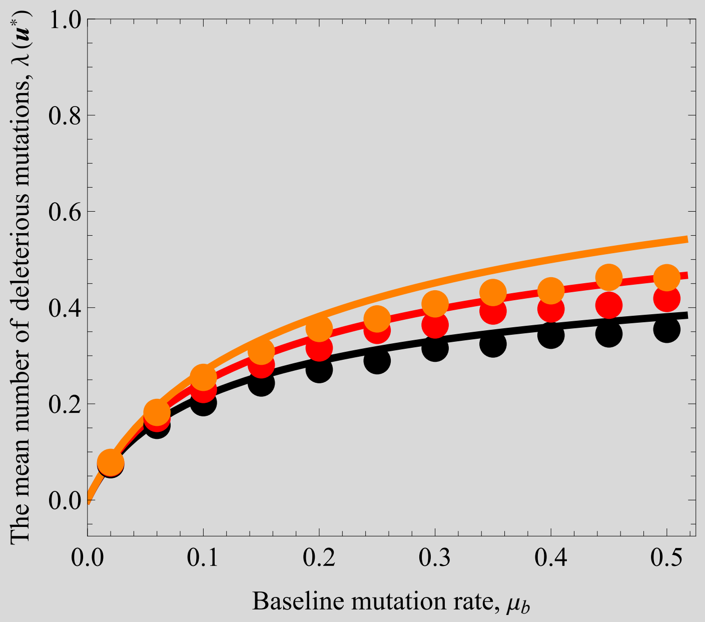


#### 2.5.7. Graphs for Figure 5 (convergence stability) of the main text

In[ ]:=
```
Clear["Global`*"]
```

In[ ]:=
```
mut[ur_] := μb (1 - ur)^α
```

In[ ]:=
```
xm[ut_, ur_] := (xo^(1-c) + a (-1 + c) ut (-1 + ur^αb) β)^(1/(1-c));
B[ut_, ur_] := a * (xm[ut, ur])^c
```

In[ ]:=
```
Su[vt_, vr_] :=
  ((α mut[vr])/(1 - vr) * (vt + 1/(d + mut[vr])) - αb * vr^(αb-1) (β (B[vt, vr] c vt)/(xm[vt, vr]) + 1/(1 - vr^αb))) /.
   {αb → 2, α → 2}
```

In[ ]:=
```
St[vt_, vr_] := (c * (β * (1 - vr^αb) * B[vt, vr])/(xm[vt, vr]) - (d + mut[vr])) /. {αb → 2, α → 2}
```

In[ ]:= $vrSol = \frac{d + 2\,\mu b - \sqrt{d^2 + 4\,d * \mu b}}{2\,\mu b};$

$vtSol = \left(xo^{-c}\left(2\sqrt{d}\;xo\,\mu b + \left(d\,xo - a\,c\,xo^{c}\,\beta\right)\left(\sqrt{d} + \sqrt{d + 4\,\mu b}\right)\right)\right) \Big/ \left(2\,a\,(-1 + c)\,d\,\beta\,\sqrt{d + 4\,\mu b}\right);$

In[ ]:= Data = {xo → 1.0, a → 0.9, c → 0.75, mr → 0.0, α → 2.0, αb → 2.0, γ → 0.00035, β → 1.0, mf → 0, d → 0.1250, μb → 0.15};

In[ ]:= urESS = vrSol /. Data

Out[ ]= 0.4132

In[ ]:= utESS = vtSol /. Data

Out[ ]= 11.6232

In[ ]:= p1 = VectorPlot[{Su[vt, vr] /. Data, 1000 * St[vt, vr] /. Data}, {vr, 0.05, 0.95}, {vt, 6, 17}]

Out[ ]=
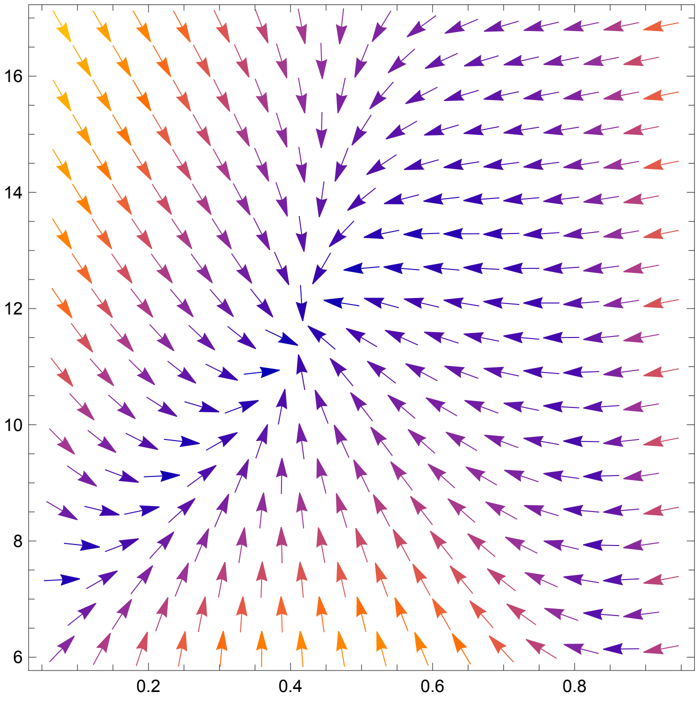


Get Data 1

Join Data and plot

Get Data 2

Join Data and plot

Get Data 3

Join Data and plot

#### Get Data 4

#### Join Data and plot

#### Put it together

In[∘]:= pAnalysis = 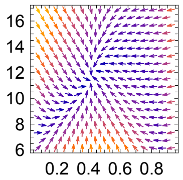 ;



In[∘]:=
```
ESSPoint = Graphics[
   {EdgeForm[Black], Gray, Thickness[.01], Disk[{urESS, utESS}, {0.015, 0.15}]}];
```

In[∘]:=
```
Show[{pAnalysis, pData1, pData2, pData3, pData4, ESSPoint }, FrameLabel →
  {Style["Investment into maintenance, vg"], Style["Age at maturity , vm"]},
 BaseStyle → {FontFamily → "Times New Roman", 14},
 LabelStyle → {FontFamily → "Times New Roman", 14}]
```

Out[∘]=

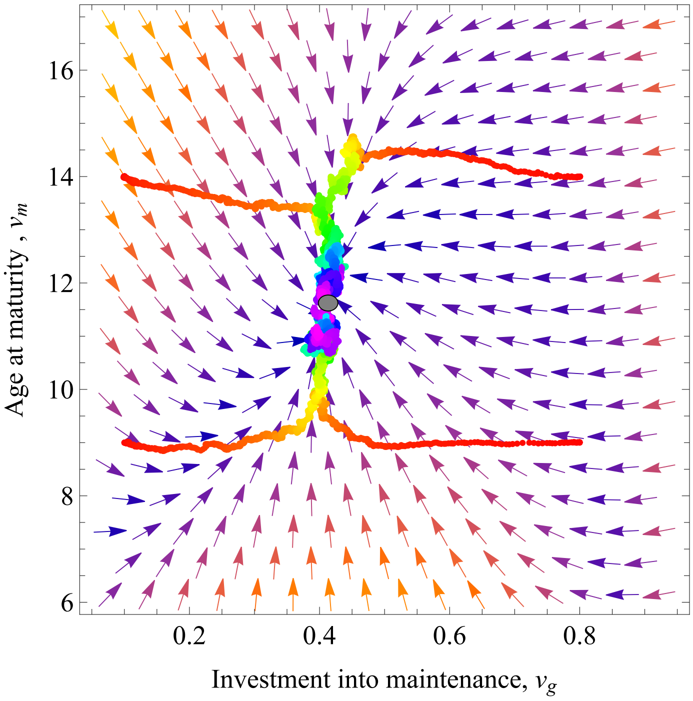

In[ ]:=
```
bar = BarLegend[{{"Rainbow", "Reverse"}, {0, 3500}}]
```

Out[ ]=

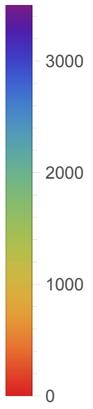

In[•]:=

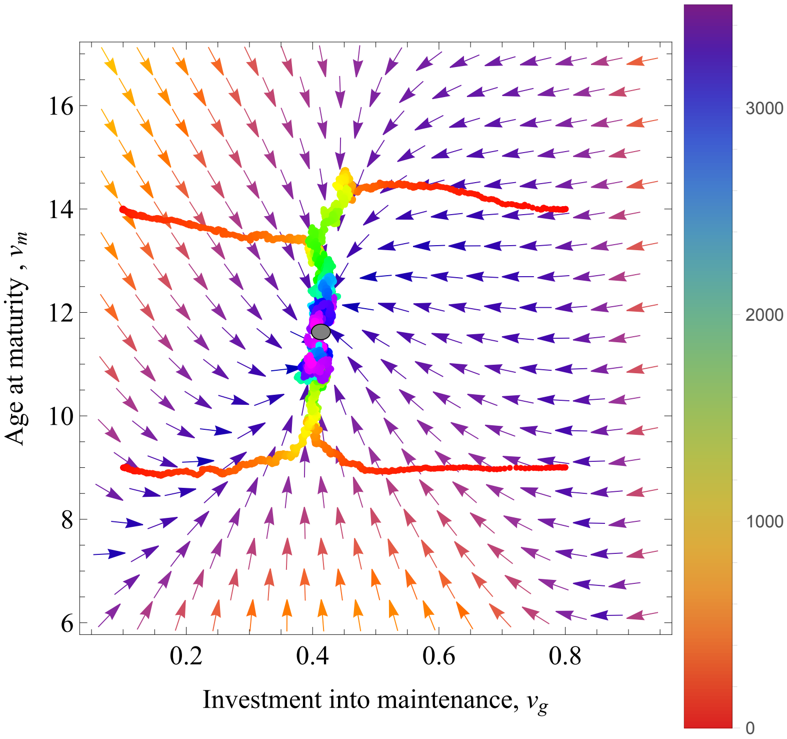
16
14
12
10
8
6
0.2
0.4
0.6
0.8
Age at maturity , $v_m$
Investment into maintenance, $v_g$
3000
2000
1000
0

Out[•]=

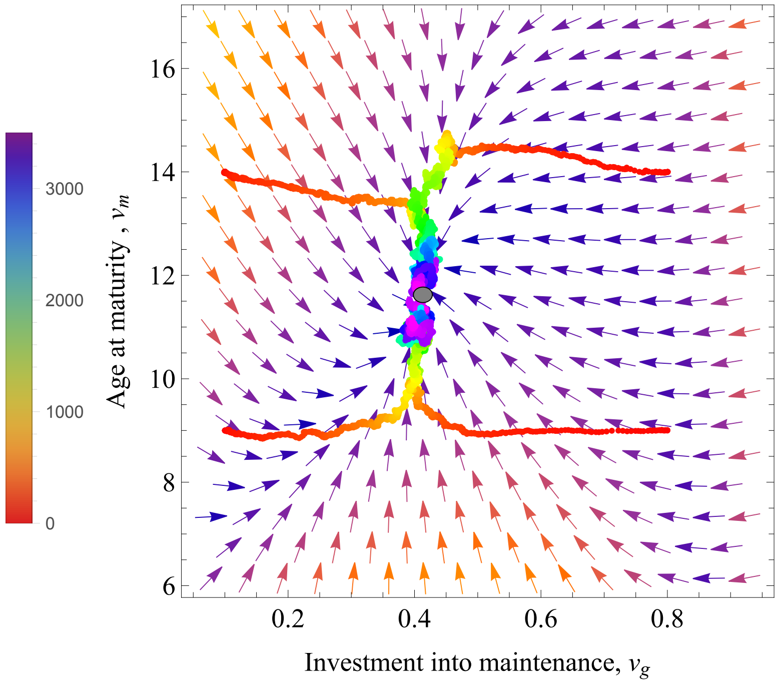
3000
2000
1000
0
16
14
12
10
8
6
0.2
0.4
0.6
0.8
Age at maturity , $v_m$
Investment into maintenance, $v_g$

In[ ]:=

Out[ ]=

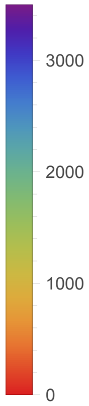
